# Functional Analysis of FRIGIDA Using Naturally Occurring Variation in *Arabidopsis thaliana*

**DOI:** 10.1101/677468

**Authors:** Lei Zhang, José M Jiménez-Gómez

## Abstract

The *FRIGIDA* locus (FRI, AT4G00650) has been extensively studied in *Arabidopsis thaliana* because of its role creating flowering time diversity. The FRI protein regulates flowering induction by binding partner proteins on its N- and C-terminus domains and creating a supercomplex that promotes transcription of the floral repressor FLC. Despite the knowledge accumulated on FRI, the function of the highly conserved central domain of the protein is still unknown. Functional characterization of naturally occurring DNA polymorphisms can provide useful information about the role of a protein and the localization of its operative domains. In the case of FRI, deleterious mutations are positively selected and widespread in nature, making them a powerful tool to study the function of the different domains of the protein. Here we explore natural sequence variation in the FRI locus in more than 1000 Arabidopsis accessions. We identify new mutations predicted to compromise the function of the protein and confirm our predictions by cloning 22 different alleles of FRI and expressing them in a common null genetic background. Our analysis allows us to pinpoint two single amino acid changes in the central domain that render the protein non-functional. We show that these two mutations determine the stability and cellular localization of the FRI protein. In summary, our work makes use of natural variants at the FRI locus to help understanding the function of the central domain of the FRI protein.

## Introduction

Natural populations of *Arabidopsis thaliana* have evolved different life history strategies to ensure reproduction in a diverse range of environmental conditions (Bloomer and Dean, 2017). Winter annual accessions germinate in autumn and integrate low temperature signals during winter (vernalization) that induce floral transition and seed production in the following spring. While winter annual accessions require vernalization to flower, summer annual plants do not. They germinate in the spring or summer, flower and set seeds before winter, thus finishing the entire reproduction cycle within in the same year (Koornneef et al., 2004; Pigliucci et al., 2003). The adoption of these two strategies depends in great part on natural genetic variation at two flowering time genes in the vernalization pathway: *FRIGIDA* (*FRI*) and *FLOWERING LOCUS C* (*FLC*). Winter annual plants typically carry the functional alleles of both genes that activate the vernalization requirement (Johanson et al., 2000; Michaels and Amasino, 1999).

*FLC* is a central floral repressor; it encodes a MADS-box transcription factor that acts as a potent transcriptional repressor of floral inducers such as *FLOWERING LOCUS T* (*FT*) and *SUPPRESSOR OF CONSTANS 1 (SOC1)* (Amasino, 2010; Michaels and Amasino, 1999). Vernalization leads to histone modification and stable epigenetic silencing at the *FLC* locus; therefore enabling the plants to flower even upon returning to the warmth (Angel et al., 2011; De Lucia et al., 2008; Heo and Sung, 2011). The coding sequence of *FLC* is highly conserved among Arabidopsis accessions (Li et al., 2014). Null alleles of *FLC* due to protein truncation or expression suppression have been reported but are rare (Werner et al., 2005). Most polymorphisms in the *FLC* locus are present in non-coding sequences and are associated with different vernalization requirement in terms of duration and temperature (Li et al., 2014). Nevertheless, the effect of variation in *FLC* can only be observed in the presence of a functional allele of *FRI* (Caicedo et al., 2004).

*FRI* plays an essential role to re-set the expression level of *FLC* during embryogenesis via chromatin modification; ensuring that every generation of newly germinated seedlings requires vernalization to flower despite coming from previously vernalized parents (Choi et al., 2009; Sheldon et al., 2008). The FRI protein acts as a scaffold to form a transcription activation complex by interaction with FRIGIDA LIKE 1 (FRL1) at the N-terminal, and SUPPRESSOR OF FRIGIDA 4 (SUF4), FRIGIDA ESSENTIAL 1 (FES1) and FLC EXPRESSOR (FLX) at the C-terminal (Choi et al., 2011). The FRI complex interacts with H3K4 methyltransferase complex COMPASS-like, HISTONE ACETYLTRANSFERASE OF THE MYST FAMLIY1 (HAM1), SWR1 chromatin remodeling complex (SWR1-C), ubiquitin-conjugating enzyme 1 (UBC1) that promotes H2B mono-ubiquitination and cap binding complex (CBC) that binds to 5’ cap of nascent pre-mRNAs (Li et al., 2018). This FRI-containing super complex modifies chromatin structure and promotes formation of the FLC 5’ to 3’ gene loop, establishing a local chromosomal environment at that promotes FLC mRNA transcriptional activation and fast elongation (Li et al., 2018). The *FRI* transcript can be first detected during embryogenesis and continues to be present in all tissues throughout the lifetime of Arabidopsis. This is also true for the FRI protein, although the protein is degraded during cold treatment; suggesting that FRI also plays a role in the down-regulation of *FLC* during vernalization (Hu et al., 2014). *FRI* encodes a 609 amino acid protein and it is the founding member of the FRIGIDA family, which comprises seven members in Arabidopsis and is characterized by a conserved central region (Risk et al., 2010). Attempts on structural analysis of FRI protein found out that only the central region is soluble (Risk et al., 2010). In grape, the crystal structure of the central region of FRI reveals 14 α-helices linked by loops or short one-turn α-helices (Hyun et al., 2016). Despite its high degree of conservation, the specific function of the central domain of FRI remains largely unknown.

Extensive nucleotide diversity has been reported at the *FRI* locus among Arabidopsis accessions from a wide range of latitudes (Hagenblad et al., 2004; Lempe et al., 2005; Shindo et al., 2005) or narrower local collections (Le Corre et al., 2002; Méndez-Vigo et al., 2011). Comparison of the FRI protein sequence from Arabidopsis and its closely related species *A. lyrata* reveals that the ancestral allele of *FRI* is the one found in the accession H51 (*FRI-H51*) (Le Corre et al., 2002). *FRI-H51* encodes a fully functional FRI protein that is able to induce *FLC* expression, thus delaying flowering in the absence of vernalization (Johanson et al., 2000). Most accessions that do not require vernalization to induce flowering carry nonfunctional alleles of *FRI*, which typically contain deleterious mutations such as premature stop codons or large deletions. The two most widespread *FRI* alleles in early flowering accessions are found in the common lab strain Columbia (Col-0) and Landsberg *erecta* (L*er*). *FRI-Col* contains a 16 bp deletion, which causes a premature stop codon after 314 amino acids, hampering the function of the protein (Johanson et al., 2000). *FRI-Ler* contains a 376 bp deletion combined with a 31 bp insertion in the promoter region that removes its translational start and greatly reduces the expression of a truncated transcript lacking 42 amino acids in the N-terminal domain (Schmalenbach et al., 2014). Despite this truncation, the FRI-L*er* protein is able to up-regulate the expression of *FLC* and delay flowering when expressed at wild-type levels (Schmalenbach et al., 2014). Accessions carrying point amino acid mutations relative to the *FRI-H51* allele, especially in the first exon, are widespread in nature; and some of these wild alleles have been shown to confer the vernalization requirement (Gazzani et al., 2003). Because of this, it has been generally accepted that truncated alleles of *FRI* are non-functional, while alleles encoding full-length protein are functional regardless of the presence point mutations (Lempe et al., 2005; Shindo et al., 2005; Werner et al., 2005).

In this work, we analyzed sequence variation in the FRI gene in 1135 Arabidopsis accessions for which re-sequencing data is available (1001 Genomes Consortium, 2016). We define 103 different alleles of FRI and select 22 alleles based on flowering time differences between accessions. We then perform heterologous expression of the selected alleles in a common genetic background and identify two deleterious mutations in the central domain of the protein. Finally, we show that these mutations determine the stability and cellular localization of the FRI protein, providing clues to the function of the central domain of this protein.

## Results

### Sequence variation in the *FRI* gene in natural populations of Arabidopsis

We explored allele diversity at the *FRI* locus using whole genome sequencing data available for 1135 Arabidopsis accessions (1001 Genomes Consortium, 2016). Short reads from all accessions were aligned to a modified Col-0 reference genome where the nonfunctional *FRI-Col-0* allele was substituted by the functional *FRI-H51* allele (Johanson et al., 2000). We then called variants in the *FRI* gene from the aligned reads. Accessions that presented low coverage and/or heterozygous variants were filtered out, leaving us with 1016 accessions for our analyses. After several rounds of manual curation to discard artifacts, we detected a total of 171 mutations (147 SNPs + 24 indels) (Suppl. Material 1). The exact sequence for five large indels could not be obtained using only short read alignments, and was annotated from de novo assemblies of the affected accessions. We annotated all variants with their effect on the FRI protein, resulting in 127 nonsynonymous variants. Among those, 26 variants were labeled as deleterious: 4 large indels, 14 frameshift indels, 7 premature stop codons and 1 deletion of the stop codon. These deleterious variants included the deletion of the start codon present in the *FRI-Ler* allele (Schmalenbach et al., 2014) and the frameshift deletion that defines the *FRI-Col-0* allele (Johanson et al., 2000). Measures of selection using these polymorphisms confirmed previous reports of positive selection on the *FRI* gene and population expansion in Arabidopsis (dn/ds = 3.98, Tajima‘s D = −2.12) (Johanson et al., 2000; Le Corre, 2005; Le Corre et al., 2002; Toomajian et al., 2006). Interestingly, nonsynonymous changes and indels accumulated preferentially outside the central region of the gene (Suppl. Fig. 1), suggesting an essential role of this conserved part of the protein (Risk et al., 2010).

**Figure 1.**
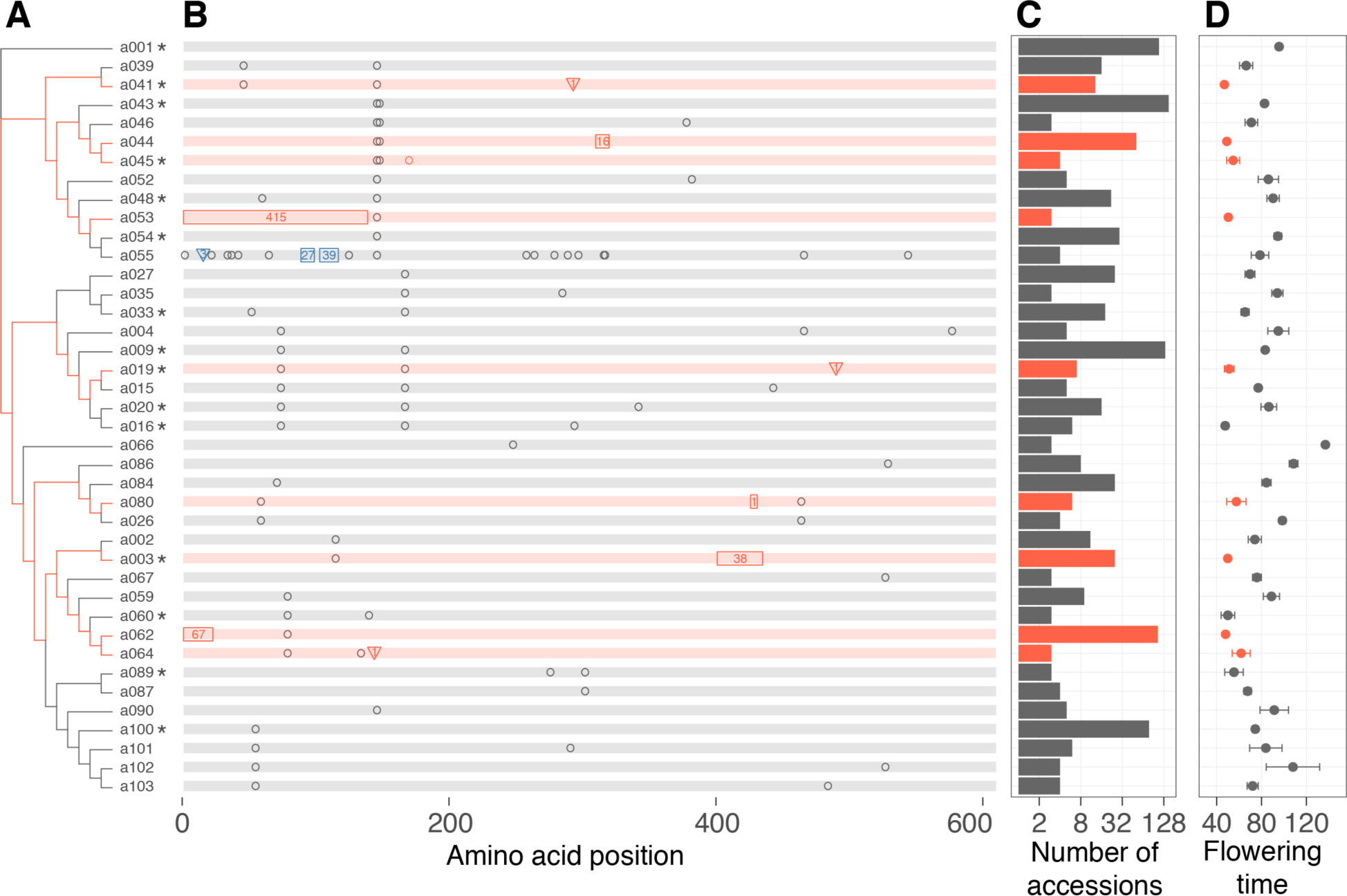
Allelic diversity at the FRI locus in Arabidopsis. Only the 40 most frequent FRI alleles are included. The complete set of alleles can be found in Suppl. Fig. 2. (A) Maximum likelihood tree for the most frequent alleles found. The H51 allele (a001) was used as root. Asterisks to the right of the names indicate alleles selected for functional characterization in this work. (B) Graphical representation of sequence differences between each allele and the a001 allele. Background color represents putatively functional (gray) and non-functional (red) alleles. Black open circles represent nonsynonymous SNPs. Rectangles and triangles represent deletions and insertions respectively, and the numbers inside indicate their length in bp. Blue symbols indicate non-frameshift indels. Red symbols indicate frameshift indels or stop codon gain/losses, all of which are considered as putatively deleterious. (C) Number of accessions (in log2 scale) carrying each allele. Alleles carrying putative deleterious mutations are colored in red. (D) Average flowering time ± standard error of the mean of all accessions carrying each allele. Alleles carrying putative deleterious mutations are indicated in red.

We defined 103 distinct FRI alleles based only on nonsynonymous changes (Fig. 1, Suppl. Fig. 2, Suppl. Material 2). The majority of accessions (61.8%) carried one of six alleles, including the H51 allele (a001), the nonfunctional alleles present in the accessions L*er* (a062) and Col-0 (a044) or another 3 functional alleles (a043, a009 and a100). Putatively deleterious mutations were present in 31 alleles distributed among 245 accessions. Almost half of the alleles (48) were found in a single accession. We used the published classification of accession in 9 STRUCTURE groups based on their genome-wide genotypes to explore the frequencies of FRI alleles across the globe (Suppl. Material 2) (1001 Genomes Consortium, 2016). As expected, each of the genetically distinct groups presented a different allelic profile for FRI, with one or two alleles being very common in each group (Fig. 2). Deleterious alleles were overrepresented among accession from Central Europe, Germany and Western Europe; but underrepresented in all other groups (p<0.01 in Fisher‘s exact test, Suppl. Fig 3). Indeed, most central European accessions carry the *FRI-Ler* allele (a062); most German accessions carry either one of two alleles that are identical except for the 16bp frameshift deletion characteristic of the Col-0 allele (a043 and a044); and western European accessions carry a variety of nonfunctional alleles, including some only found in Western Europe (i.e. a003, a041 and a019). Interestingly, the only relict accession that contained a deleterious allele of FRI was Cvi-0, the southern-most accession in the set, which carries an allele with a premature stop codon that is private to this accession (a021, Suppl. Fig. 2, Fig. 2, Suppl. Material 1).

**Figure 2.**
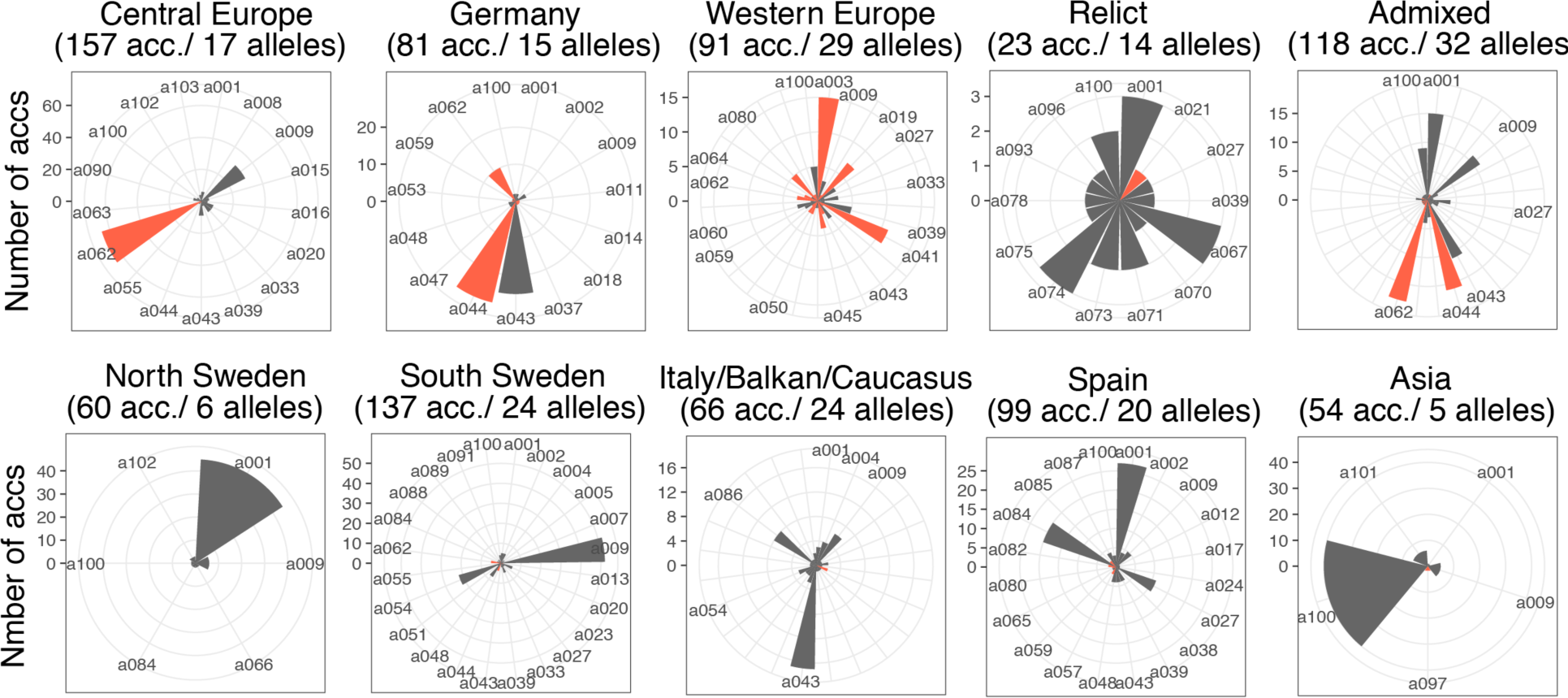
FRI allele frequency and distribution among Arabidopsis populations. The number of accessions carrying individual FRI allele in each STRUCTURE group indicated in (1001 Genomes Consortium; 2016) is represented in circular barplots. Allele names are as in Figure 1. Names from low frequency alleles in groups with more than 25 alleles have been removed, and can be found in Suppl. Material 2. Bars representing putatively non-functional alleles are colored in red.

**Figure 3.**
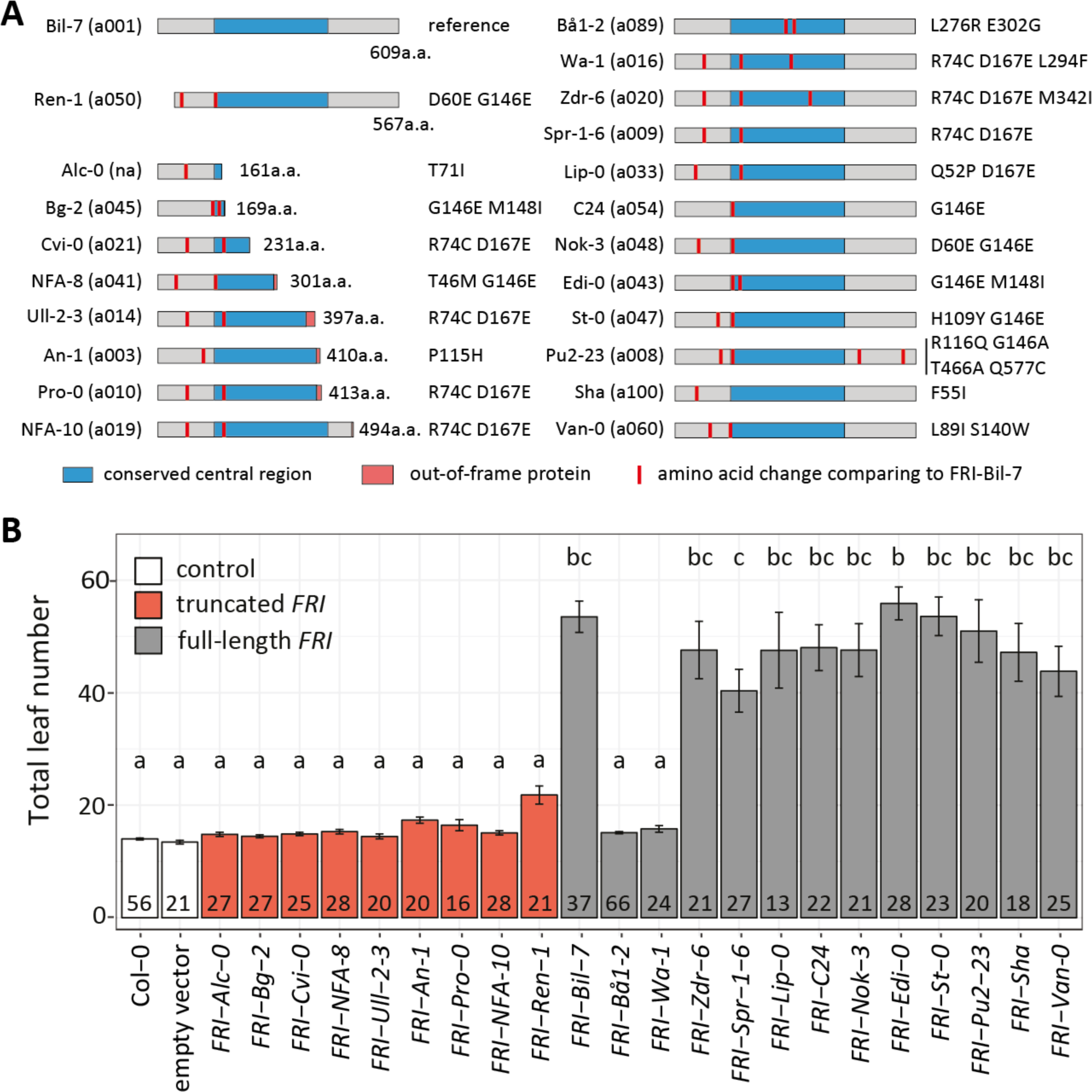
Flowering time of transgenic lines carrying selected alleles of FRI. (A) Schematic representation of selected FRI proteins including mutations relative to the FRI-Bil-7 allele. (B) Flowering time of T_1_ transgenic lines carrying the FRI alleles shown in (A). All plants were grown under long day conditions. Error bars represent the standard error of the mean. The number in each bar indicates the number of plants analyzed per line. Letters on top of each bar represent significance groups as determined by the Tukey HSD test.

We studied the effect of the deleterious mutations in the function of the protein by analyzing their flowering time (1001 Genomes Consortium, 2016). On average, accessions carrying deleterious mutations in FRI flowered significantly earlier than accessions without deleterious mutations (Suppl. Fig. 4A, p<2e-16 in ANOVA). As previously observed (Stinchcombe et al., 2004), flowering time from accessions with functional alleles of FRI strongly correlates with latitude, but this correlation disappears in accessions carrying deleterious FRI alleles (Suppl. Fig. 4B). We then studied flowering time variation among accessions grouped by their FRI allele, considering only those alleles sufficiently represented in the population (alleles present in at least 3 accessions, Fig. 1). All alleles containing deleterious mutations accelerated flowering with respect to alleles labeled as functional. Interestingly, alleles a016, a060 and a089 were associated with early flowering despite lacking deleterious polymorphisms (Fig. 1). These three alleles, together with another 19, were selected to further study the effect of natural polymorphisms in the function of the FRI protein.

**Figure 4.**
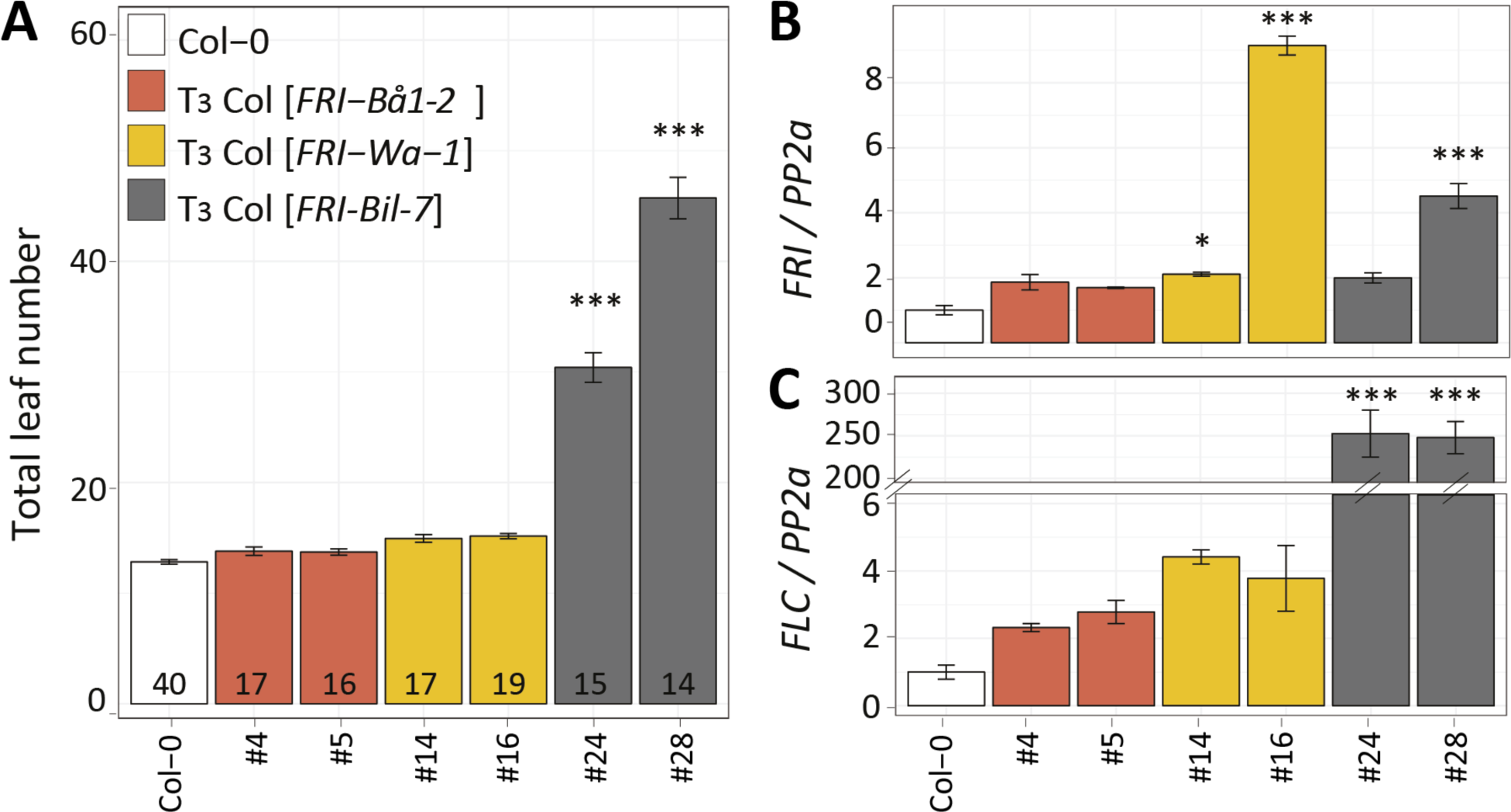
Confirmation of non-functional alleles in T3 transgenic lines. (A) Flowering time of homozygous T_3_ lines carrying single insertions of the FRI-Bå1-2, FRI-Wa-1 or FRI-Bil-7 transgenes in comparison to untransformed Col-0. All plants were grown in the greenhouse under long day conditions. The number in each bar indicates the number of individual plants analyzed. (B, C) Relative expression of FRI (B) and FLC (C) in the transgenic lines shown in (A). The aerial part from 10-day-old seedlings grown in long day conditions was pooled for each replicate, and three biological replicates were used for each line. Expression was normalized to the expression of PP2a and compared to Col-0 using the Tukey HSD test (* p < 0.05; *** p < 0.001). In all cases, error bars represent the standard error of the mean.

### Functional analysis of natural FRI alleles

We cloned and sequenced a region spanning positions −1372bp to +2691bp from the start codon from 22 different alleles of the *FRI* locus (Fig. 3A, Suppl. Material 3). One of the accessions used, Alc-0, carried an allele with a premature termination codon that was not found in the re-sequenced accessions (Fig. 1). Sequence analysis of the promoters of all chosen alleles revealed frequent single nucleotide polymorphisms and small deletions of 1 to 3 nucleotides, and a major deletion in the St-0 allele (from −540 to −470 bp), where no known regulator motif has been described (Suppl. Material 4, Suppl. Fig. 5).

**Figure 5.**
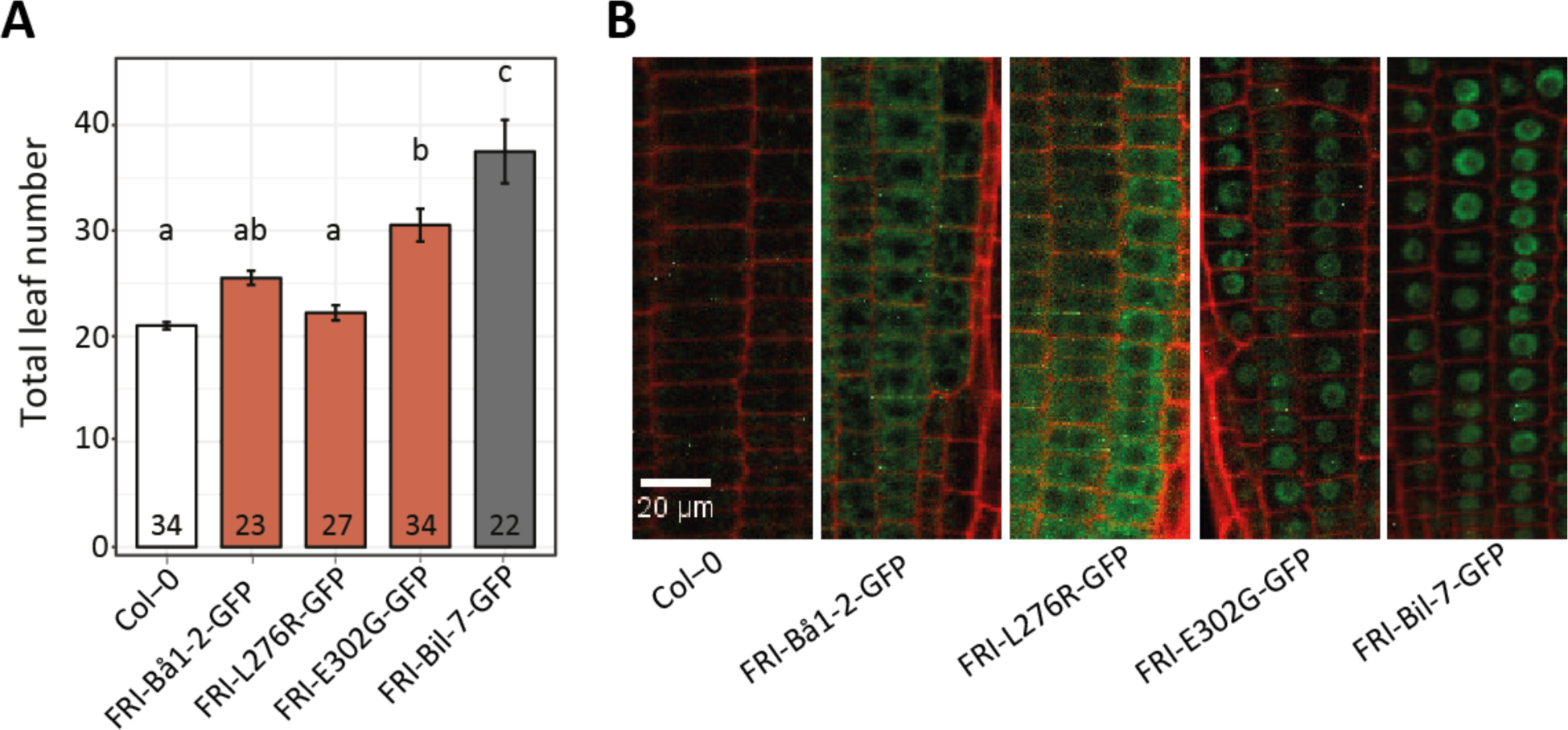
Characterization of amino acid mutations in FRI-Bå1-2 protein. (A) Flowering time of T_1_ transgenic plants and Col-0 control. All transgenic plants were selected on MS agar medium containing kanamycin for 7 days; Col-0 plants were grown on MS agar medium without kanamycin. All plants were transplanted to soil and grown under long day conditions in the greenhouse. Numbers inside each bar indicate the number of individual plants analyzed in each line; letters on top of the box represent the significance groups as determined by Tukey HSD test. (B) Confocal images of root tips of 7-day-old seedlings. The root was treated with propidium iodide, which outlined the cells in red. GFP signals from the FRI proteins could only be observed in the cytoplasm or nuclei of transgenic lines but not in the Col-0 plants. One representative image from each genotype is shown.

We studied functionality of these alleles by measuring flowering time in transgenic lines expressing each allele from its native promoter. To enable comparison, all constructs were transformed in a wild type Col-0 background that carries a null allele of *FRI* and a functional allele of *FLC* (Schmalenbach et al., 2014). We measured flowering time as total leaf number in the T_1_ generation. All lines carrying *FRI* alleles encoding C-terminal truncated proteins flowered at the same time as Col-0 control plants (Fig. 3B). The longest C-terminal truncated allele was that from the accession NFA-10 (a019) that only misses 115 amino acids, suggesting that essential amino acids for the function of FRI are located beyond amino acid 494. Interestingly, lines carrying the N-terminal truncated *FRI-Ren-1* allele flowered significantly later than the control (21.8±1.6 in *FRI-Ren-1* compared to 14.0±0.2 in Col-0, p < 0.001 in t-test between the two groups). This is reminiscent of the slight but significant flowering time delay observed with the *FRI-Ler* allele, where a deletion overlapping the promoter and start codon of the *FRI* gene caused expression at low levels of a functional FRI protein lacking the first 42 amino acids (Schmalenbach et al., 2014). It is therefore likely that the large deletion present in the promoter and N-terminal domain of the *FRI-Ren-1* allele, although different from the one found in *FRI-Ler*, has a similar consequence (Fig 3A, Suppl. Fig. 5).

Most *FRI* alleles encoding full-length proteins significantly delayed flowering when introduced into a Col-0 background (Fig. 3B). Interestingly, from the three putatively functional alleles that were selected because their association with early flowering in Fig. 1, the *FRI-Van-0* (a060) allele failed to delay flowering time in the transgenic lines, indicating that this allele is functional and that the accessions that carry it may contain mutations in additional flowering time genes. On the contrary, nonfunctional alleles of *FRI* were confirmed in lines carrying the *FRI-Bå1-2* (a089) and *FRI-Wa-1* (a016) alleles, which did not flower significantly different from Col-0 (15.1±0.2 and 15.8±0.6 total leaves in the *FRI-Bå1-2* and *FRI-Wa-1* lines respectively versus 14.0±0.2 in Col-0). In order to better understand the causes for this lack of functionality of the *FRI* locus, we further investigated the mutations in the *FRI-Bå1-2* and *FRI-Wa-1* alleles.

### Nonsynonymous polymorphisms disrupt *FRI-Bå1-2* and *FRI-Wa-1*

We first confirmed the lack of function in the *FRI-Bå1-2* and *FRI-Wa-1* alleles in independent T_3_ families that were homozygous for a single copy of the transgene. As observed in the T_1_ generation, T_3_ lines carrying the *FRI-Bå1-2* and *FRI-Wa-1* alleles flowered with a similar leaf number as Col-0 plants, while lines carrying the *FRI-Bil-7* allele flowered with a significantly increased number of leaves (Fig. 4A). One possibility is that early flowering in the *FRI-Bå1-2* and *FRI-Wa-1* transgenic lines is caused by low expression of these alleles triggered by the various mutations found in their regulatory regions (Suppl. Material 4, Suppl. Fig. 5). However, all lines showed either slightly higher or significantly elevated *FRI* transcription levels in comparison to the control Col-0 plants (Fig. 4B), suggesting that the early flowering phenotype of the transgenic lines carrying the *FRI-Bå1-2* and *FRI-Wa-1* alleles is likely caused by coding differences. If that was the case, these alleles, albeit normally expressed, should fail to increase the expression of its main target *FLC*. Indeed, in contrast to the *FRI-Bil-7* allele, the *FRI-Bå1-2* and *FRI-Wa-1* alleles did not significantly up-regulate the expression of *FLC* (Fig. 4C). These results suggest that the aborted functionality of the *FRI-Bå1-2* and *FRI-Wa-1* alleles is due to coding polymorphisms that render the proteins nonfunctional.

In terms of their nonsynonymous mutations, both the FRI-Bå1-2 and FRI-Wa-1 protein carry amino acid changes in the central domain of the FRI protein, whose function has not been characterized in depth despite being highly conserved in the Brassicaceae family (Suppl. Fig. 6). The two mutations present in the FRI-Bå1-2 protein (L276R and E302G) result in dramatic changes in the chemical properties of the residues. The hydrophobic leucine is replaced by the negatively charged glutamic acid and the positively charged arginine is replaced by the hydrophobic glycine. The mutation in the central domain of the FRI-Wa-1 (L294F), replaces the leucine in Bil-7 by a much bulkier phenylalanine. The FRI-Wa-1 allele also carries two amino acid mutations in the N-terminal domain, although these are also present in the functional FRI-Zdr-6 and FRI-Spr1-6 alleles and they are not likely to affect FRI function (Fig. 3). In summary, the amino acid changes found in the central domain of the FRI-Bå1-2 and FRI-Wa-1 proteins could potentially compromise their function and provide clues to the role of this domain in the FRI protein.

**Figure 6.**
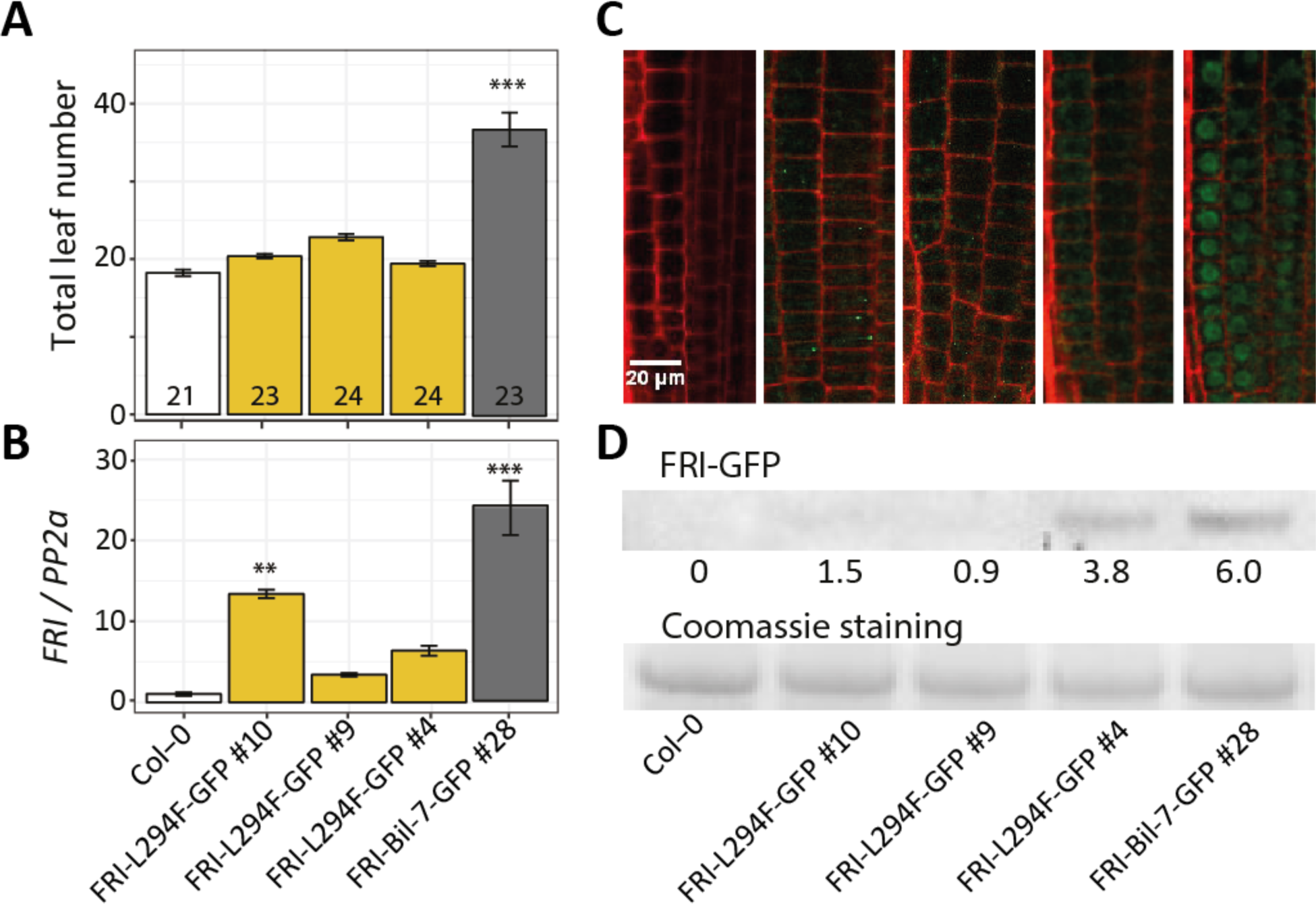
Characterization of L294F amino acid mutation in FRI protein. (A) Flowering time expressed as total leaf number from homozygous T_3_ transgenic lines carrying FRI-L294F-GFP or FRI-Bil-7-GFP. All plants were grown under long day conditions in a greenhouse. The number in each bar indicates the number of plants per line analyzed. Letters in each bar represent significance groups as determined by Tukey HSD test. (B) Expression of FRI in the same genotypes shown in (A). Aerial part was pooled from 10-day-old seedlings grown in long day conditions. Three biological replicates were used per genotype. Expression was normalized to the expression of PP2a and compared to Col-0 (***, p < 0.001) (C) Confocal images of root tips of one-week-old seedlings grown on MS agar medium under long day condition. One representative image from each line is shown. (D) Western blot detection of FRI protein using anti-GFP antibodies. Seedlings from the same experiment shown in (C) were pooled for protein extraction. The numbers represent the intensity of the bands.

### Amino acid mutation L276R prevents the nuclear localization of FRI

We studied the effect of the two mutations present in the FRI-Bå1-2 allele on protein function. To do this, we obtained GFP-tagged versions of the *FRI-Bil-7* and *FRI-Bå1-2* alleles (*FRI-Bil-7-GFP* and *FRI-Bå1-2-GFP*) under the control of their respective native promoter, and performed directed mutagenesis on the *FRI-Bil-7-GFP* allele to include the individual L276R or E302G mutations (*FRI-L276R-GFP* and *FRI-E302G-GFP*). We scored flowering time as total leaf number in the T_1_ generation from Col-0 plants transformed with these constructs (Fig. 5A). Lines carrying the functional *FRI-Bil-7-GFP* transgene flowered significantly later than Col-0 plants and lines transformed with the *FRI-Bå1-2-GFP* allele, indicating that the GFP tag does not interfere with the function of the FRI protein. Among the individual mutations, the lines carrying the *FRI-E302G-GFP* construct flowered significantly later than Col-0, while those carrying the *FRI-L276R-GFP* allele did not. This result suggests that the L276R mutation present in the FRI-Bå1-2 protein is sufficient to prevent its normal function.

We characterized the effect of the L276R mutation on FRI by studying its cellular localization in root tip cells, where wild type FRI accumulates in the nuclei (Kim et al., 2006). The nuclear localization of FRI is essential to its function because the FRI complex delays flowering by causing chromatin modification at the *FLC* locus (Choi et al., 2011; Li et al., 2018). While a strong GFP signal could be observed in the nuclei of the root tip cells in late flowering lines (*FRI-E302G-GFP* and *FRI-Bil-7-GFP*, Fig. 5B), the FRI-Bå1-2-GFP and FRI-L276R-GFP proteins were observed only in the cytoplasm, suggesting that the L276R mutation in the FRI-Bå1-2 allele affects its ability to enter the nucleus.

### Amino acid mutation L294F disrupts the stability of FRI

To study the effect of the L294F mutation in FRI, we created an artificial *FRI-L294F-GFP* allele by introducing the mutation in the *FRI-Bil-7-GFP* construct; which was then transformed into Col-0 plants. Consistent with previous results, T_3_ lines carrying the *FRI-L294F-GFP* construct flowered early, despite presenting a high expression of the transgene; while lines carrying the *FRI-Bil-7-GFP* allele showed both significantly high expression of *FRI* and late flowering (Fig. 6A, 6B). We then studied the localization of both alleles in root tip cells. In contrast to the strong nuclear localized GFP signal in the plants transformed with *FRI-Bil-7-GFP*, we observed no GFP signal in two of the *FRI-L294F-GFP* transgenic lines, and only a weak cytoplasmic GFP signal in the third line (Fig. 6C). Western blot results using the same plant material agreed with the differences in protein abundance observed under the microscope (Fig. 6D). The low accumulation of FRI protein with the L294F mutation suggests that this single amino acid mutation causes instability and degradation of the FRI protein and thus abolishes its function.

## Discussion

The analysis of natural variation in flowering time in Arabidopsis focused on the *FRI* gene from a very early stage, because of its major contribution to this trait (Burn et al., 1993; Clarke and Dean, 1994; Lee et al., 1993). Since its identification (Johanson et al., 2000), multiple studies have surveyed sequence variation at the *FRI* gene in Arabidopsis (Gazzani et al., 2003; Hagenblad et al., 2004; Le Corre et al., 2002; Schmalenbach et al., 2014; Shindo et al., 2005). With the appearance of short read sequencing, it has become possible to investigate the alleles present in hundreds of individuals with reasonable amounts of time and funds, although obtaining an accurate representation of the mutations in a locus still remains challenging. Unsupervised mutation mining in the FRI locus with 1135 re-sequenced Arabidopsis accessions resulted in the identification of 307 variants, with more than one third of them being artifacts that appear form the alignment of short reads to the borders of large indels or rearrangements. After manual curation of variants and de novo assembly of structural variants, we defined 171 naturally occurring mutations in 1016 accessions, including 26 deleterious mutations. As expected, the geographical distribution of these variants was not random, and accessions carrying deleterious *FRI* alleles were underrepresented among southern and northern European accessions but enriched in central and western European accessions (Fig. 2). Evolutionary analyses in the gene confirmed the previously observed genome-wide signatures of population expansion and local positive selection at the *FRI* locus (1001 Genomes Consortium, 2016; Le Corre et al., 2002). When particular mutations can be associated to phenotypic differences in the plants that carry them, they provide an opportunity to investigate the role of the different parts of the protein. In this study, we selected 22 alleles carrying a diversity of mutations with the aim to shed light on the function of the different domains of the protein.

All alleles studied functionally, except *FRI-Bå1-2*, carry mutations in the N-terminal region. This region has been proposed to be under strong selection to generate adaptively significant flowering time variation (Le Corre et al., 2002; Shindo et al., 2005), and it is known to interact with FRL1, which is required for FRI-mediated up-regulation of *FLC* (Choi et al., 2011). In our study, we did not observe significantly different dn/ds ratios for the N-terminal region when compared to the central domain or the C-terminal region, and none of the nonsynonymous polymorphisms tested in this region compromised the function of the protein. However, we observed a large variation in flowering time among the late flowering transgenic lines. It is therefore possible that amino acid substitutions in the N-terminal domain generate subtle differences in protein function that may not be detectable in our study despite our efforts of using large number of independent T_1_ lines to minimize the variation due to transgene insertion position and number. Experiments with better control on the dosage of the transgene or using introgression lines would be better suited to evaluate subtle effects of amino acid substitutions in the N-terminal domain of FRI. One of the studied alleles, *FRI-Ren-1*, presents a partial deletion of the N-terminal domain, presumably forcing translation from a second in-frame start codon at position +127, as it has been described for the *FRI-Ler* allele (Schmalenbach et al., 2014). We have previously shown that the deletion of the first 42 amino acids in FRI-L*er* does not affect flowering time, and accessions carrying the *Ler-FRI* allele show an intermediate flowering phenotype caused by the low expression of the allele due to the deletion in its promoter (Schmalenbach et al., 2014). In this work we hypothesize a similar situation for the *FRI-Ren-1* allele, based on the intermediate flowering time of the transgenic lines expressing this allele (Fig. 3). Interestingly, a previous study showed that deletion of the first 50 amino acids in the N-terminal domain of FRI is sufficient to disrupt binding to FRL1 in a yeast two hybrid system (Choi et al., 2011). The functionality of the *FRI-Ren-1* and *FRI-Ler* alleles suggests that the functional positions in the N-terminal of the FRI protein are located between amino acids 42 and 50. Further work will be needed to explore this hypothesis.

In our study, all alleles presenting truncations in the C-terminal domain were unable to delay flowering time when compared to Col-0, a *fri* null mutant (Fig. 3). A previous study had shown that a truncated FRI protein lacking the last 151 amino acids from the C-terminal domain failed to delay flowering time when expressed in Col-0 (Risk et al., 2010). In fact, Y2H analysis shows that the 150 C-terminal amino acids physically interact with components of the FRI complex such as FLX, SUF4 and FES1 (Choi et al., 2011). In our study, *FRI-NFA-10* encodes a FRI protein with a C-terminal deletion including the last 115 amino acids (Fig. 3). Therefore, the lack of functionality of this allele allows us to redefine the functional domain of the C-terminal region in FRI to its last 115 amino acids.

Despite its conservation across Brassicaceae (Suppl. Fig. 6), the function of the central domain of FRI has remained elusive. We have identified two nonsynonymous mutations in *FRI-Bå1-2* and *FRI-Wa-1* that disrupt the function of the protein. The single mutation from leucine to arginine at residue 276 in FRI-Bå1-2 prevents the nuclear localization of the FRI protein (Fig. 5). We identified five nuclear localization signals (NLS) in the FRI sequence based on the prediction of three bioinformatic tools (King and Guda, 2007; Nair et al., 2003; Sperschneider et al., 2017), but residue 276 is not in the vicinity of any of these NLSs (Suppl. Fig. 7). A recent study hypothesized that the leucine 276 is tightly surrounded by other hydrophobic residues, therefore a mutation to the positively charged arginine may disrupt the core structure of FRI and indirectly affect its ability to enter the nucleus (Hyun et al., 2016). We believe that our study confirms this hypothesis. However, the same study predicted that the mutation at position 302 from positively charged glutamate to glycine would disrupt the protein structure, which we have not been able to confirm through functional studies of the E302G mutation (Fig. 5). Interestingly, another mutation at position 294 abolishes the function of FRI in a different fashion. The change from leucine to a bulky hydrophilic phenylalanine in the tightly folded core of the protein may disrupt its folding and cause its degradation, as indicated by its very low accumulation (Fig. 6).

In terms of their origin, the L294F mutation is present in six accessions, Wa-1, Wil-1, Wil-2, Litva, Tottarp-2 and Est (Suppl. Material 1). Although these accessions are distributed along central Europe and south Sweden, all of them have a clear central European ancestry (Suppl. Fig. 8, Suppl. Material 2), suggesting a central European origin. The mutation L276R is present in three accessions, Bå1-2, Bå4-1 and Bå5-1, all originary from a single location in Southern Sweden (Suppl. Fig. 8, Suppl. Material 1, Suppl. Material 2), which suggest a recent appearance of the a086 allele and contrasts with the low abundance of deleterious alleles among southern Swedish accessions (Fig. 2). Although speculative, it is possible that this recent appearance of accessions that do not need vernalization at such high latitudes could be due to the increase in mean temperatures associated to global warming.

In summary, we have surveyed natural variation in the coding sequence of FRI using data from more than one thousand Arabidopsis accessions, allowing us to define the distribution of nonfunctional alleles in this species and identifying novel variants that can shed light on the function of the protein. According to our results, the central domain of FRI would be implicated both in the stability and nuclear localization of the protein.

## Methods

### Analysis of short reads from 1135 Arabidopsis accessions

We constructed a modified Arabidopsis reference genome by substituting the Col-0 *FRI* allele in the TAIR10 reference sequence by the functional H51 allele and modifying accordingly the available annotation. Short reads for 1135 Arabidopsis accessions (1001 Genomes Consortium, 2016) were downloaded from SRA (SRP056687) and aligned to the modified reference genome using Bowtie 2 version 2.3.4.2 with default parameters (Langmead and Salzberg, 2012). Previous to variant calling, reads with mapping quality lower than 5 were discarded using samtools v1.7 (Li et al., 2009), duplicated reads were removed using Picardtools (http://broadinstitute.github.io/picard) and indels were realigned using GATK IndelRealigner (McKenna et al., 2010). Finally, variants on the *FRI* gene were called simultaneously in all alignments with GATK UnifiedGenotyper (McKenna et al., 2010). Polymorphisms in the *FRI* gene were manually curated using the IGV genome browser (Robinson et al., 2011) to remove artifacts due to low coverage or low quality alignments around large indels. In addition, we removed 111 accessions that presented an average coverage over the *FRI* gene lower than 10, and 8 accessions that presented more than 1 heterozygous variant. We identified 5 large indels for which the precise sequence could not be determined from the short read alignments. The sequence of these indels was obtained from de novo assemblies of the affected accessions (id: 108, 139, 1070, 5236, 6090, 6390, 7427, 8240, 9121 and 9726) using SPAdes with k-mer size set to 33, 55, 77, and 99, and otherwise default parameters (Bankevich et al., 2012). The resulting de novo assembled contigs were screened for the presence of the *FRI* gene using BLAST v2.6.0+ (Camacho et al., 2009) and the *FRI* genomic sequence from the H51 accession. The indel sequence was extracted from the alignment of the positive contigs against the H51 genomic sequence using muscle v3.8.31 (Edgar, 2004). The effect of each polymorphism on the FRI protein was annotated with ANNOVAR (Wang et al., 2010).

The matrix containing 171 polymorphisms and their presence/absence in all 1016 accessions analyzed was used to generate in silico FRI alleles, giving rise to 130 distinct FRI CDS. In order to calculate dn/ds and the average behavior of each codon, we constructed codon alignments of the 1016 alleles by excluding insertions. Average ds/dn value between all alleles and the H51 reference as well as the cumulative codon behavior for indels, nonsynonymous and synonymous variants were calculated using SNAP (http://www.hiv.lanl.gov). Tajima‘s D was calculated for the complete region of the FRI gene using vcftools v0.1.15 (Danecek et al., 2011).

Flowering time and STRUCTURE group membership for each accession was obtained from the 1001 genomes project (1001 Genomes Consortium, 2016)(http://1001genomes.org/tables/1001genomes-FT10-FT16_and_1001genomes-accessions.html).

### Cloning of *FRIGIDA* alleles

The *FRI* locus in all accessions analyzed was amplified using Phusion High-Fidelity DNA Polymerase using primer FRI-clonF (at position −1552, GGAAGCAAATGACCGTAAAATC) and FRI-clonR (position +2946, TCTCAGTGCTGTATAACTACA). The PCR product was digested with BamHI and EcoRI, whose restriction sites are at −1372 and +2691 respectively relative to the annotated start codon of *FRI* in TAIR10. After digestion, the reaction was run in an agarose gel and the expected size band was purified using Gel Extraction Kits (QIAGEN). All the purified *FRI* fragments were introduced into pBinDsRed (kindly provided by Ed Cahoon, University of Nebraska) making use of the BamHI and EcoRI restriction sites.

The *FRI-GFP* fusion protein was generated making use of binary vector pGWB404 (Nakagawa et al., 2007) that contains an in-frame GFP tag downstream of the gateway recombination site. To do this, the *FRI* promoter and coding sequence up to the stop codon (−1372 to +2293) was amplified from the genomic DNA of the accession Bil-7 using the following primers where the AttB sequence for Gateway cloning is marked in parenthesis: FRI-gwF (position −1372) (GGGGACAAGTTTGTACAAAAAAGCAGGCTTA) GATCCCAAAATCTAGTGCCGCCG and FRI-gwR (position +2293) (GGGGACCACTTTGTACAAGAAAGCTGGGTT)TTTGGGGTCTAATGATGAGTTACTGC. Mutations L276R, L294F or E302G were introduced to the amplified DNA fragment of the *FRI-Bil-7* allele by site directed mutagenesis, as described in (Ho et al., 1989), using the following overlapping primers where the mutated nucleotides to introduce the amino acid changes are in parenthesis: mutL276R-F (position +816) TTGCTTTTAC(G)AGTTGCTTGTTTTG and mutL276R-R (position +838) AACAAGCAACT(C)GTAAAAGCAAACC; mutL294-F (position +896) GCTGGATTT(T)ATAAGGATGAGTGG and mutL294-R (position +872) CCACTCATCCTTAT(A)AAATCCAGC; mutE302G-F (position +899) GAATG(G)GATTGCCGGTGCTTTG and mutE302G-R (position +921) CAAAGCACCGGCAATC(C)CATTC. The wild type and mutated *FRI-Bil-7* PCR fragments were introduced into pENTR201 and subsequently pGWB404 binary vector. The alleles were named *FRI-Bil-7-GFP, FRI-L276R-GFP, FRI-L294F-GFP* and *FRI-E302G-GFP* respectively.

Inserts in all final constructs were sequenced and verified by comparison to the expected in silico constructs. Subsequently, all constructs were transformed into *Agrobacterium tumefaciens* strain GV3101 by electroporation. Agrobacteria transformed with pBinDsRed were selected on solid YEP medium containing 50 mg/l kanamycin, 25 mg/l rifampicin and 50 mg/l gentamicin; whereas Agrobacteria transformed with pGWB404 were selected on 100 mg/l spectinomycin, 25 mg/l rifampicin and 50 mg/l gentamicin. Positive colonies were confirmed by colony PCR and cultured in liquid YEP medium containing same antibiotics. All constructs, and an empty pBinDsRed vector used as a negative control, were transformed into Col-0 plants by the floral dip method (Clough and Bent, 1998).

### Selection and phenotyping of transgenic plants

Transgenic seeds carrying pBinDsRed constructs were selected by their red fluorescence under a fluorescence stereomicroscope using monochromatic green light (wavelength ∼ 580 nm). Transgenic seeds selected in this way were directly sown in soil at the greenhouse under long day conditions for flowering time measurement.

Transgenic plants carrying pGWB404 constructs were selected on MS agar containing 50 mg/l kanamycin. Surviving T_1_ plants were transferred to soil and grown under long day condition in the greenhouse for flowering time measurement. As a control, Col-0 seedlings were grown on MS agar without antibiotics and transferred to soil together with the transgenic plants.

Flowering time in all experiments was recorded as the number of days between seed sowing and flowering, and the number of rosette leaves and cauline leaves on the day when the first flower opened. The ratio between rosette and cauline leaves did not show any variation between the different genotypes in our experiment, so only the total leaf number was used. We found a high correlation between total leaf number and days to the opening of the first flower, but differences between genotypes were larger when using leaf number. Consequently, only total leaf number is reported as a measurement of flowering time. Flowering time measurements in the experiment with all T_1_ transgenic plants containing different *FRI* alleles were terminated at end of the 8^th^ week. All plants not flowering at that time were assigned with the highest recorded total leaf number in the experiment.

### Expression analysis using quantitative real-time PCR

For expression analysis, stratified seeds were sown on presoaked soil and grown in a growth chamber in long day conditions for 10 days. Seedlings (leaves plus shoot) were harvested and immediately frozen in liquid nitrogen. Material from 8 to 10 seedlings was pooled for each one of the three biological replicates. Total RNA was extracted using RNeasy Plant Mini Kit (QIAGEN). Contaminating DNA was removed by treatment with TURBO DNA-free™ Kit (Ambion). cDNA was synthesized from 5 µg RNA using Super Script**®** II Reverse Transcriptase in a 20 µl system (Invitrogen). Quantitative RT-PCR was performed on a CFX384 Touch™ Real-Time PCR Detection System using SYBR Green dye (iQ™ SYBR**®** Green Supermix, Biorad) using the following primers: FRI_f - TGCCTGATCGTGGTAAAGGGAAG, FRI_r - GCACCGGCAATCTCATTCGAAC; FLC_f - CCGAACTCATGTTGAAGCTTGTTGAG, FLC_r - CGGAGATTTGTCCAGCAGGTG; PP2A_f - TAACGTGGCCAAAATGATGC, PP2A_r - GTTCTCCACAACCGCTTGGT. The expression of *FRI* and *FLC* were normalized to the expression of *PP2A* and subjected to further statistical analysis.

### Microscopic imaging

Transgenic seeds carrying GFP constructs were sterilized and sown on MS agarose medium and grown vertically under long days for 7 days. At this time, whole seedlings were dipped in a solution of propidium iodide (10 mg/ml) for one minute and rinsed in water. Root tips were cut off and mounted for visualization under confocal microscope (ZEISS LSM800). The GFP signal was visualized under 488 nm light and the propidium iodide (PI) signal was visualized under 514 nm light. The merged image of both GFP and PI signals are presented.

### Western blot assay

Transgenic seeds carrying GFP constructs were sterilized and sown on MS agarose medium and grown under long days for 7 days. Whole seedlings were pooled and pulverized in liquid nitrogen. Total protein was extracted using PEB protein extraction buffer (Agrisera Antibodies) according to the manufacturer‘s instructions. The protein solution was adjusted to equal concentration and separated on a NuPAGE™ 4-12% Bis-Tris Protein Gel (Thermo Fisher Scientific). Each sample was loaded twice on the gel allowing separation of the gel in two parts with identical samples. One part of the gel was subjected to Coomassie staining and the other was used for blotting. Anti-GFP and secondary antibodies (Roche) were used according to the manufacturer‘s protocol. Detection was carried out using the Amersham ECL Prime Western Blotting Detection Reagent (GE healthcare) following the manufacturer‘s protocol and band intensity was analyzed with Image J software.

### FRIGIDA sequences from different species

FRI protein sequences were downloaded from GenBank. The species used and their GeneBank IDs were: *Arabidopsis thaliana* (P0DH90.1), *Boechera stricta* (AFJ66199.1), *Arabidopsis lyrata* (ABY51730.1), *Tarenaya hassleriana* (XP_010531851.1), *Capsella rubella (*AFJ66217.1), *Vitis vinifera* (XP_002283789.1), *Brassica rapa* (NP_001289004.1), *Medicago truncatula* (XP_003602718.2), *Camelina sativa* (XP_010426308.1), *Nelumbo nucifera* (XP_010250405.1), *Arabis alpina* (KFK36753.1). FRI DNA sequences were obtained from the cloned products. All alignment was performed by Clustal Omega on the EMBL-EBI website.

