## Supplemental Figures for "Functional Analysis of FRIGIDA Using Naturally Occurring Variation in *Arabidopsis thaliana*"

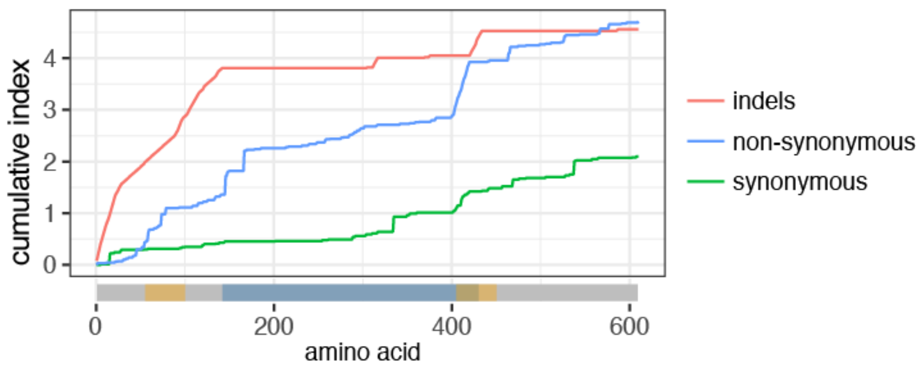


**Supplementary Figure 1.** Cumulative index of indels, synonymous SNPs and non-synonymous SNPs for all pairwise comparisons between all 103 FRI alleles defined in this work (Suppl. Fig. 2). A representation of the canonical FRI protein is depicted in the x-axis, with coiled coil domains highlighted in yellow and the central conserved region highlighted in blue.

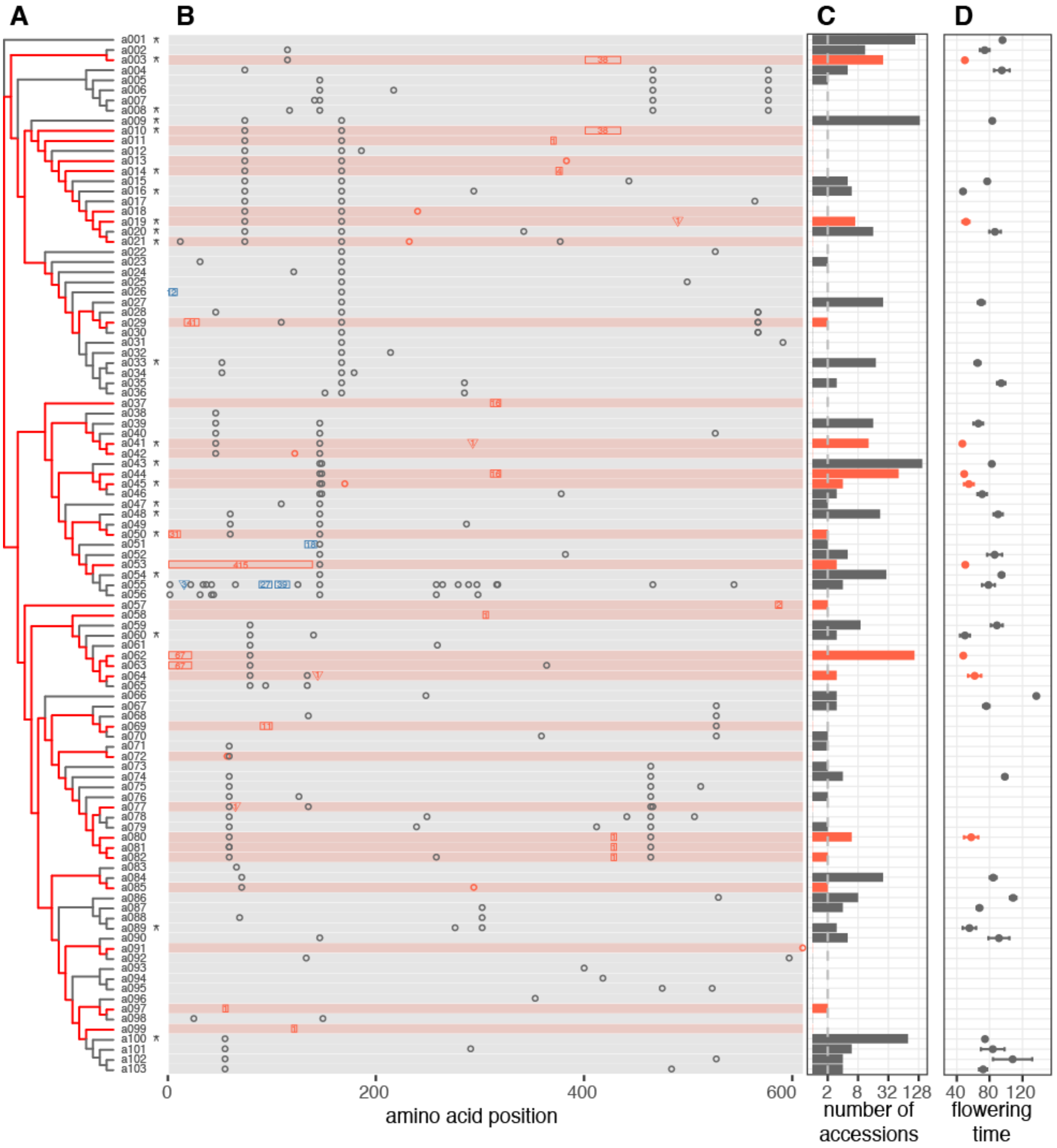


**Supplementary Figure 2.** (A) Maximum likelihood tree for 103 FRI alleles present in the 1135 re-sequenced. The H51 allele (a001) was used as root. Asterisks in the names indicate alleles functionally characterized in this work. (B) Graphical representation of sequence differences between each allele and the H51 allele. Background color represents putatively functional (gray) and non-functional (red) alleles. Black open circles represent nonsynonymous SNPs. Rectangles and triangles represent deletions and insertions respectively, and the numbers inside indicate their length in bp. Blue symbols indicate non-frameshift indels. Red symbols indicate frameshift indels or stop codon gain/losses, all of which are considered as putatively deleterious. (C) Number of accessions (in log2 scale) carrying each allele. The dotted line indicates the threshold of 2. Putatively deleterious alleles are colored in red. (D) Average flowering time ± standard error of the mean of all accessions carrying each allele. Putatively deleterious alleles are indicated in red.

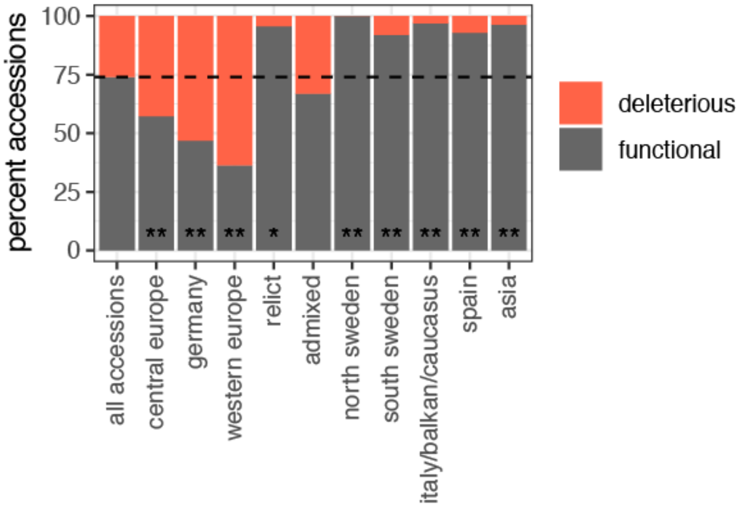


**Supplementary Figure 3.** Percentage of deleterious and non-deleterious FRI alleles in each of the 10 STRUCTURE population groups. The frequency across all accessions analyzed is shown on the leftmost bar. Divergence from this frequency was calculated using a two-sided Fisher's test (* p<0.05, **p<0.01)

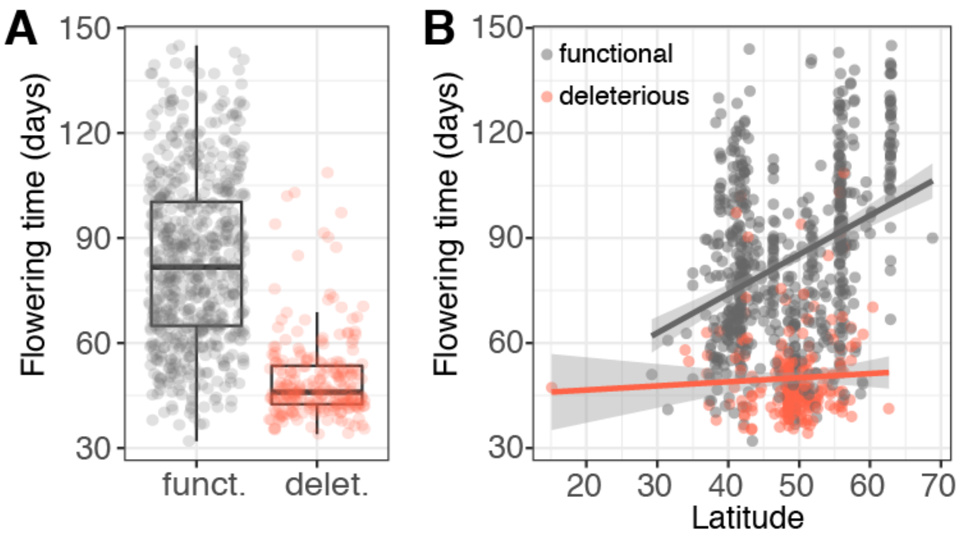


**Supplementary Figure 4.** Flowering time of 1016 Arabidopsis accessions classified by the putative functionality of their FRI allele. Flowering time data at 16 degrees Celsius was obtained from the 1001 genome's project and functionality of the FRI allele as indicated in Supplementary table 2.

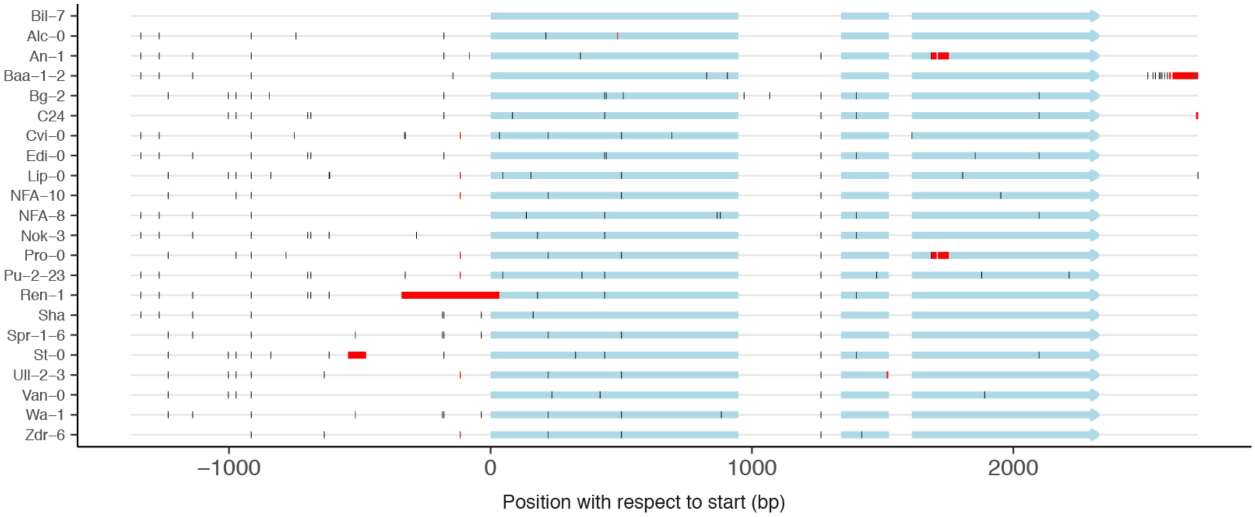


**Supplementary Figure 5.** Graphical representation of polymorphisms found in the FRI alleles cloned and sequenced. All sequences are compared to the fully functional allele in the Bil-7 accession. Light blue blocks represent exons, black tick marks represent SNPs, and red blocks represent deletions.

```
                                         276                 294     302
Arabidopsis_thaliana RLMTEGGLAAAEKMDARGLLLLVACFGVPSNFR--STDLLDLIRMSGSNEIAGALKRSQFL 313
Arabidopsis_lyrata   RLMSEGGLAAAEKMDARGLLLLVACFGVPSNFR--SMDLLDLIRMSGSNEIAGALKRSPFL 316
Capsella_rubella     RLMGEGGLAVAEQIDARGLLLLIACYGVPSDFR--SMDLLDLIRTSGSNEIVGALRRSPFL 314
Brassica_rapa        RLVGEGWLGAAEAMDARGLLLLVACFGIPESFK--SMDLLDLIRQSGTDEIVGALKRSPFL 297
Camelina_sativa      RLIGEGGLAAAELMDARGLLLLIACYGVPSNFR--SMDLLDLILTCGFNEIAGALRRSRFL 328
Arabis_alpina        RLVVEGGLATAEQMDARGLLLLIASFGVPSNFR--NMDLLDLIRKSGSAEIAAALKRSPFL 292
Boechera_stricta     RLMGEGGLAAAEQMDARGLLLLIACFGVPSNFR--SMDLLDLIRTSGSNEIAGALRRSPFL 330
Tarenaya_hassleriana RLMSEGGLANAQTMDARGLLLLIACFGVPSTFGDLDLDLLDLLRQSKPSEISNALRRSAFL 296
Vitis_vinifera       RLINEGGVSNASDIDARGLLLLVASFGIPALFR--NEDLRNLIRLSCPKEISDALRRSRFL 276
Medicago_truncatula  RLVVEGGVAMASEMDARGLTLFLACFGIHGVFR--NEDIANLVRLSKPGEISHLLRVSGSL 319
Nelumbo_nucifera     RLVNEGGVANANSIDARGLLLFIGSFGVPSVFG--NDDLSDLIRLSNPKEIAGALRRSHFL 344
                     **: ** :. *. :***** *::..:*:   *   . *: :*:  .   **   *: *  *
```

**Supplementary Figure 6.** Sequence alignment of FRI conserved central domain. Alignment of FRI protein sequences from species in the Brassicaceae family (in black) and outside (in gray). Positions 276 and 302 are respectively mutated to R and G in FRI-Bå1-2 and are colored in blue. Position 294 is mutated to F in FRI-Wa-1 and is colored in red. Alignment was produced with Clustal Omega.

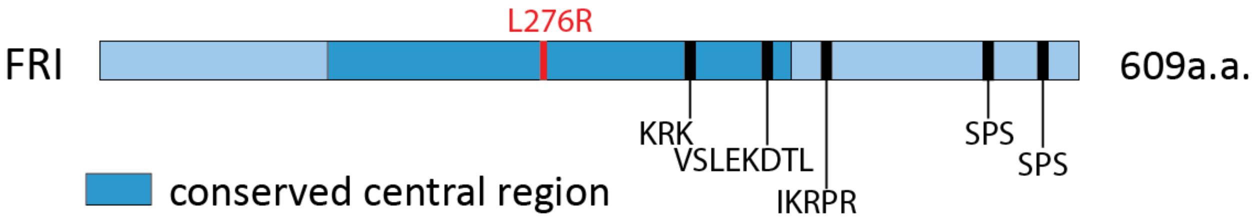


**Supplementary Figure 7.** Relative position of L276R amino acid substitution and the predicted nuclear localization signals (NLS) on FRI protein.

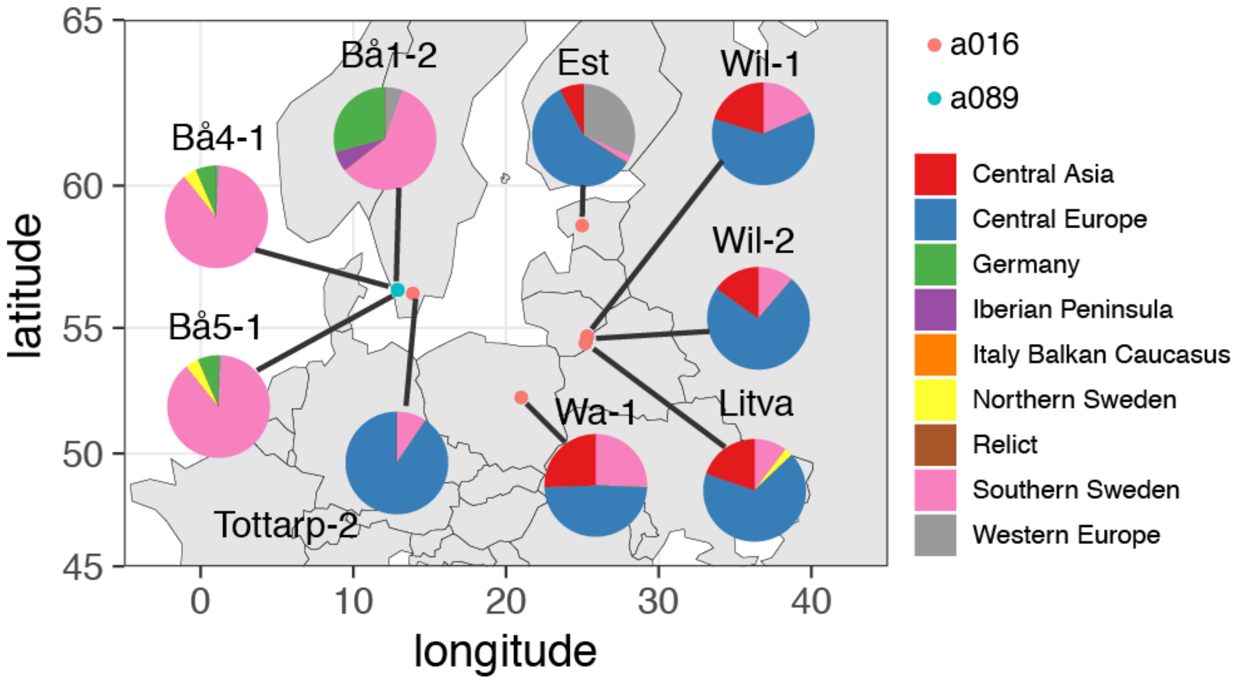


**Supplementary Figure 8.** Geographical distribution of accessions carrying the L276R (a089) and L294F (a016) mutations. The location of the accession as indicated in the 1001 genome's project is represented with dots in the map. The location for accession Litva, for which no location was available, was set to the capital of Lithuania. Genetic components for each accession are represented in the pie charts.
