## Supplemental Material 3 for "Functional Analysis of FRIGIDA Using Naturally Occurring Variation in *Arabidopsis thaliana*"

**Supplementary Material 3.** DNA sequences of all *FRI* alleles cloned in this work.

>FRI_Bil-7
GGATCCCAAAATCTAGTGCCGCCGAGTCCTTTGGGACACAGTTTCGCACTTATCATTCGATTCGGCAAGCAGTAACTCGC
AAAATCCGCACCACCGTGAAACATACTCCGGCGCTGAACGCTAGACGCAAAATCCATATTACTGCTGCAGAAAGACGGAG
AGAACAAGAGACTGACCGATCATAAGAGAAGAGAGCTTCAAGGCTTCAGCTGATGAAGAATCAATGGCGGCGAGAGAGAG
TGAGAATCGAGTAGTTACTGAGAGGAAGATGAAACGTATCTCCTGAGAAAAAACGTAAGAGAGTTCCGTTTCAACAGTAT
TTTTCTCGGCGGTCTTCAGATTTTCCGTCCAACTCTCTCTTTCCTCTTTCAGTTTTTTTTTGCCCAATCACTCGTGGTAG
GATTTTTTTATAAGAAAGTTACGAAAATGCCCTCTCGTGCTTAGAGAGATTCAATTCGTGAACATATAATATGCCGTGAA
TATACCACCAACTCTCTTACTCGTACTCGCATGTGGTGCGTGCAATATACACGTGGTTCGAATCAACGGCTACGTTTTAC
TGATAGGTGGCTCGTTTAGTTTTGCACTGCCTTTGGGTGCGCGTGACCAAGTCATTTTCCAAATGTGTTTTAATATTTGT
GAACTATTATTAATAAAAAAGCGAGCGACGTTGTTCTTTGATGAGGGAACGACCGTACGCTTGGCGACCTTCTCTGATTG
CGTAGAGTCGTTTTCCTTATTGGGCTTACCTAGTACCGTCAGCCCACAATGTTTTCATTTCTTTCCCACCTAAAACTAAA
AGTACTCACAAGTCACAACTTAAACCAAGTACACAAGGATTTTATCATGGGATTATCGTGTTTGAAGACTAAAAAGAGCA
CACCATCACCCCCATTAGTGCAGGTAGAGTAAGACAGTAACTTTTGGGTTCATATTACCGAGCAAGAACCGTTATTTGTG
ATTAGACATGTTATAAACCACTGCTTTAGTGACTATTTAAAACAATATATTACATGTCGTAATCATGCAACCTAACTATG
TTTTCATTAATCAAATACAAAGAATAAAGAGAAAAGTGCGTAGATTCAATTATTTGGCATAGACTCAAAAGAGTGTATAT
ATATCTGACTTTTATTAAATTATTAAACACAAATACATATTTTCATAAGCAAAACTATAAAAGCCCTAAACATATAATGA
TTACCTCAAAGGAAAAAGTCGTTTTCTCCTAATTAAAAGATAGGTTACTTCCTAATTAATATATAATTTATGTGAACTTC
ACAATATACAGTTCAATAAAATTTGGTAATTTGACCGATTTAAGGAGAGTGGAAATTAGGGCTTCTGCAATCTTTTTTCT
TCGCCGCAATCTCATGTCCAATTATCCACCGACGGTGGCGGCGCAACCCACAACGACGGCGAATCCACTGCTGCAGCGAC
ATCAATCTGAACAGCGACGAAGAGAATTACCGAAGATTGTCGAAACAGAGTCTACAAGTATGGACATTACGATCGGTCAA
TCTAAGCAGCCTCAATTTTTGAAATCCATAGACGAATTAGCTGCGTTTTCAGTTGCAGTGGAAACATTCAAACGCCAATT
CGATGATCTTCAGAAGCACATCGAGTCAATCGAAAACGCAATTGATTCCAAACTCGAGAGTAACGGCGTTGTCCTCGCCG
CGCGGAACAATAATTTCCATCAGCCGATGTTATCGCCTCCGCGGAACAATGTATCTGTAGAAACCACCGTCACTGTGAGC
CAACCGTCTCAGGAGATTGTACCGGAGACGTCGAATAAACCGGAGGGGGGACGTATGTGTGAGTTGATGTGTAGCAAAGG
TCTGCGTAAATACATATACGCGAATATCTCTGATCAAGCTAAGTTAATGGAAGAGATTCCTTCAGCTTTGAAATTGGCCA
AGGAGCCAGCGAAGTTTGTATTGGATTGTATTGGCAAGTTTTACTTACAAGGGCGTAGAGCATTTACTAAAGAGTCGCCT
ATGAGCTCTGCGAGACAAGTTTCGCTTCTTATACTGGAGTCTTTTCTTCTAATGCCTGATCGTGGTAAAGGGAAGGTGAA
GATTGAGAGTTGGATTAAAGATGAGGCGGAGACGGCTGCTGTTGCTTGGAGGAAAAGGTTGATGACTGAAGGAGGATTAG
CTGCGGCTGAGAAAATGGATGCAAGGGGTTTGCTTTTACTAGTTGCTTGTTTTGGTGTTCCTTCAAACTTTAGGAGTACA
GATTTGCTGGATTTGATAAGGATGAGTGGTTCGAATGAGATTGCCGGTGCTTTGAAGCGGTCACAGTTTCTTGTCCCTAT
GGTCTCAGGTACCATATTCTGTtCTCACTCGGTGAATTTCATTGCAAAGGTGGTTCCTTTTGTTGACATCATCGACCAAC
ATCAAGTTCCATCTTTGTTTTTCGATAAGCTTGATGGTATAAACTAGGAGAGCACATCAAATATTTAGAGTGCAATGACT
GATTGAGCCAAATCCTAGCTAGAAATTAATCTGGAAAGAACTTGGAACTCTCAACCATAGGTTTTGGTACGAAATTGTTG
CTTGTCAGAACCAAATGATAGGCTATTGCCTTGAAATAGTGTTTCTTGTGGTTTCCAATATTGGAAGTTAAAATCGTATG
ACTTAGCTGTTGGATACTAATTAAGCTTAAGCAATGCCAACTCTAAGAAGTGGTACTTACACAATATTCTATTGGTCATA
GGTATAGTTGAATCAAGTATCAAGCGTGGAATGCATATTGAAGCTCTTGAGATGGTTTATACCTTTGGCATGGAGGATAA
GTTTTCAGCTGCTCTAGTTCTAACTTCATTCTTAAAGATGAGCAAGGAGTCATTTGAGAGGGCAAAACGGAAAGCCCAGT
CACCGCTGGCATTTGTATGAACCCTTCCCTTGCACATTATGTACCTTTATGAACTCTTTATCATCATCTGAGTCTGACCA
TTGATATATTTATTTCTCAACAGAAAGAAGCGGCTACAAAGCAGCTAGCTGTGTTATCATCAGTTATGCAGTGTATGGAG
ACTCACAAGTTAGATCCTGCGAAAGAACTACCAGGATGGCAGATCAAAGAGCAAATTGTTAGCTTGGAGAAAGACACTCT
TCAGCTCGACAAAGAGATGGAAGAGAAAGCAAGATCTCTCAGTTTAATGGAGGAAGCCGCACTTGCCAAGAGAATGTATA
ACCAACAGATAAAACGTCCAAGGTTGTCACCCATGGAAATGCCACCAGTAACTTCTTCATCGTATTCTCCTATCTACCGT
GATAGAAGCTTTCCTAGTCAAAGAGACGATGACCAAGATGAAATATCAGCTCTTGTGAGTAGTTACCTCGGCCCGTCAAC
ATCTTTTCCTCATCGCTCAAGAAGATCCCCGGAATATATGGTTCCACTTCCACATGGTGGGTTAGGAAGAAGTGTATATG
CATATGAACATCTGGCCCCAAATTCATACTCTCCAGGTCACGGACATAGACTTCATCGACAGTACTCTCCGTCTTTGGTT
CACGGACAGAGACATCCACTACAGTACTCTCCTCCAATTCATGGACAACAACAGTTACCATATGGTATACAAAGGGTTTA
CAGACATTCACCATCTGAAGAAAGATATTTGGGTTTATCCAATCAAAGGTCTCCTCGCAGTAACTCATCATTAGACCCCA
AATAGGAGGAATGTAAATTTGTAACAAAGCTTTTTGTTTTTGCTTAAGTTAGTCATTTATTTAACTCCCAACAGTCTCAA
AATTTAATTTAATGTTTGGGGCTTAAGAATGCAAATTTTTTTGCTCCTGTAATTGACATTTAAGATGCTAATGTTATTGC
TTCAGAGGTTTTAGTCAACCTCAGATACATCGATATCACTATCTAAATAGACCTCTGGCTCTTGGTCATCTGGATTCTCT
TCATCTTCTGTCTCTGTTCCTTCTTGTTCTCGTTGCACTGCTCGAGCAATTGCGGATTCCAACCTTGTGCTTACAGTTTC

CCATGACACAAGCTTTTCCATGAATGTATTTATGTCCGCCTTCTTATCTTTCTTGAGGAaGATGAATTC

>FRI_Ren-1
GGATCCCAAAATCTAGTGCCGCCGAGTCCTTTGGGACAGAGTTTCGCACTTATCATTCGATTCGGCAAGCAGTAACTCGC
AAAATCCGCACCACCGTGAAACATACTCTGGCGCTGAACGCTAGACGCAAAATCCATATTACTGCTGCAGAAAGACGGAG
AGAACAAGAGACTGACCGATCATAAGAGAAGAGAGCTTCAAGGCTTCAGCTGATGAAGAATCAATGGCGGCGAGAGTGAG
TGAGAATCGAGTAGTTACTGAGAGGAAGATGAAACGTATCTCCTGAGAAAAAACGTAAGAGAGTTCCGTTTCAACAGTAT
TTTTCTCGGCGGTCTTCAGATTTTCCGTCCAACTCTCTCTTTCCTCTTTCAGTTTTTTTTTGCCCAATCACTCGTGGTAG
GATTTTTTTATAAGAAAGTTACGAAAATGCCCTCTCGTGCTTAGAGAGATTCAATTCGTGCACATATAATATGCCGTGAA
TATACCACCAACTCTCTTACTCGTACTCGCATGTGGTGCGTGCAATATACACGTGGTTCGAATCAACGGCTACGTTTTAC
TGATAGGTGGCTCGTTTAGTTTTGCACTGCCTTTGGGTGCGCGTGACCAAGTCATTTTCCAAATGTGTTTTAATATTTGT
GAACTATTATTAATAAAAAAGCGAGCGACGTTGTTCGTTGATGAGGGAGCGACCGTACGCTTGGCGACCTTCTCTGATTG
CGTAGAGTCGTTTTCCTTATTGGGCTTACCTAGTACCGCCAGCCCACAATGTTTTCATTTCTTTCCCACCTAAAACTAAA
AGTACTCACAAGTCACAACTTAAACCAAGTACACAAGGATTTTATCATGGGATTATCGTGTTTGAAGACTAAAAAGAGCA
CACCATCACCCCCATTAGTGCAGGTAGAGTAAGACAGTAACTTTTGGGTTCATATTACCGAGCAAGAACCGTTATTTGTG
ATTAGACATGTTATAAACCACTGCTTTAGTGACTATTTAAAACAATATATTACATGTCGTAATCATGCAACCTAACCCAC
AACGACGGCGAATCCACTGCTGCAGCGACATCAATCTGAACAGCGACGAAGAGAATTACCGAAGATTGTCGAAACAGAGT
CTACAAGTATGGACATTACGATCGGTCAATCTAAGCAGCCTCAATTTTTGAAATCCATAGAGGAATTAGCTGCGTTTTCA
GTTGCAGTGGAAACATTCAAACGCCAATTCGATGATCTTCAGAAGCACATCGAGTCAATCGAAAACGCAATTGATTCCAA
ACTCGAGAGTAACGGCGTTGTCCTCGCCGCGCGGAACAATAATTTCCATCAGCCGATGTTATCGCCTCCGCGGAACAATG
TATCTGTAGAAACCACCGTCACTGTGAGCCAACCGTCTCAGGAGATTGTACCGGAGACGTCGAATAAACCGGAGGGGGAA
CGTATGTGTGAGTTGATGTGTAGCAAAGGTCTGCGTAAATACATATACGCGAATATCTCTGATCAAGCTAAGTTAATGGA
AGAGATTCCTTCAGCTTTGAAATTGGCCAAGGAGCCAGCGAAGTTTGTATTGGATTGTATTGGCAAGTTTTACTTACAAG
GGCGTAGAGCATTTACTAAAGAGTCGCCTATGAGCTCTGCGAGACAAGTTTCGCTTCTTATACTGGAGTCTTTTCTTCTA
ATGCCTGATCGTGGTAAAGGGAAGGTGAAGATTGAGAGTTGGATTAAAGATGAGGCGGAGACGGCTGCTGTTGCTTGGAG
GAAAAGGTTGATGACTGAAGGAGGATTAGCTGCGGCTGAGAAAATGGATGCAAGGGGTTTGCTTTTACTAGTTGCTTGTT
TTGGTGTTCCTTCAAACTTTAGGAGTACAGATTTGCTGGATTTGATAAGGATGAGTGGTTCGAATGAGATTGCCGGTGCT
TTGAAGCGGTCACAGTTTCTTGTCCCTATGGTCTCAGGTACCATATTCTGTTCTCACTCGGTGAATTTCATTGCAAAGGT
GGTTCCTTTTGTTGACATCATCGACCAACATCAAGTTCCATCTTTGTTTTTCGATAAGCTTGATGGTATAAACTAGGAGA
GCACATCAAATATTTAGAGTGCAATGACTGATTGAGCCAAATCCTAGCTAGAAATTAATCTGGAAAGAACTTGGAACTCT
CAACCATAGGTTTTGGTACGAAATTGTTGCTTGTCAGAACCAAATGATAGGCTATTGCCTTGAAATAGTGTTTCTTGTGG
TTTCCAATATTGGAAGTTAAAATCATATGACTTAGCTGTTGGATACTAATTAAGCTTAAGCAATGCCAACTCTAAGAAGT
GGTACTTACACAATATTCTATTGGTCATAGGTATAGTTGAATCAAGTATCAAGCGTGGAATGCATATTGAAGCTCTTGAA
ATGGTTTATACCTTTGGCATGGAGGATAAGTTTTCAGCTGCTCTAGTTCTAACTTCATTCTTAAAGATGAGCAAGGAGTC
ATTTGAGAGGGCAAAACGGAAAGCCCAGTCACCGCTGGCATTTGTATGAACCCTTCCCTTGCACATTATGTACCTTTATG
AACTCTTTATCATCATCTGAGTCTGACCATTGATATATTTATTTCTCAACAGAAAGAAGCGGCTACAAAGCAGCTAGCTG
TGTTATCATCAGTTATGCAGTGTATGGAGACTCACAAGTTAGATCCTGCGAAAGAACTACCAGGATGGCAGATCAAAGAG
CAAATTGTTAGCTTGGAGAAAGACACTCTTCAGCTCGACAAAGAGATGGAAGAGAAAGCAAGATCTCTCAGTTTAATGGA
GGAAGCCGCACTTGCCAAGAGAATGTATAACCAACAGATAAAACGTCCAAGGTTGTCACCCATGGAAATGCCACCAGTAA
CTTCTTCATCGTATTCTCCTATCTACCGTGATAGAAGCTTTCCTAGTCAAAGAGACGATGACCAAGATGAAATATCAGCT
CTTGTGAGTAGTTACCTCGGCCCGTCAACATCTTTTCCTCATCGCTCAAGAAGATCCCCGGAATATATGGTTCCACTTCC
ACATGGTGGGTTAGGAAGAAGTGTATATGCATATGAACATCTGGCCCCAAATTCATACTCTCCAGGTCACGGACATAGAC
TTCATCGACAGTACTCTCCGTCTTTGGTTCACGGACAGAGACATCCACTACAGTACTCTCCTCCAATTCATGGACAACAA
CAGTTACCATATGGTATACAAAGGGTTTACAGACATTCACCATCTGAAGAAAGATATTTGGGTTtATCCAATCAAAGGTC
TCCTCGCAGTAACTCATCATTAGACCCCAAATAGGAGGaATGTAAATTTGTAACAAAGCTTTTTGTTTTTGCTTAAGTTA
GTCATTTATTTAACTCCCAACAGTCTCAAAATTTAATTTAATGTTTGGGGCTTAAGAATGCAAATTTTTTTGCTCCTGTA
ATTGACATTTAAGATGCTAATGTTATTGCTTCAGAGGTTTTAGTCAACCTCAGATACATCGATATCACTATCTAAATAGA
CCTCTGGCTCTTGGTCATCTGGATTCTCTTCATCTTCTGTCTCTGTTCCTTCTTGTTCTCGTTGCACTGCTCGAGCAATT
GCGGATTCCAACCTTGTGCTTACAGTTTCCCATGACACAaGCTTTTCCATGAATGTATTTATGTCCGCCTTCTTATCTTT
CTTGAGGAAGATGAATTC

>FRI_Alc-0

GGATCCCAAAATCTAGTGCCGCCGAGTCCTTTGGGACAGAGTTTCGCACTTATCATTCGATTCGGCAAGCAGTAACTCGCAAAA
TCCGCACCACCGTGAAACATACTCTGGCGCTGAACGCTAGACGCAAAATCCATATTACTGCTGCAGAAAGACGGAGAGAACA
AGAGACTGACCGATCATAAGAGAAGAGAGCTTCAAGGCTTCAGCTGATGAAGAATCAATGGCGGCGAGAGAGAGTGAGAA
TCGAGTAGTTACTGAGAGGAAGATGAAACGTATCTCCTGAGAAAAAACGTAAGAGAGTTCCGTTTCAACAGTATTTTTCT
CGGCGGTCTTCAGATTTTCCGTCCAACTCTCTCTTTCCTCTTTCAGTTTTTTTTTGCCCAATCACTCGTGGTAGGATTTT
TTTATAAGAAAGTTACGAAAATGCCCTCTCGTGCTTAGAGAGATTCAATTCGTGCACATATAATATGCCGTGAATATACC
ACCAACTCTCTTACTCGTACTCGCATGTGGTGCGTGCAATATACACGTGGTTCGAATCAACGGCTACGTTTTACTGATAG
GTGGCTCGTTTAGTTTTGCACTGCCTTTGGGTGCGCGTGACCAAGTCATTTTCCAAATGTGTTTTGATATTTGTGAACTA
TTATTAATAAAAAAGCGAGCGACGTTGTTCTTTGATGAGGGAACGACCGTACGCTTGGCGACCTTCTCTGATTGCGTAGA
GTCGTTTTCCTTATTGGGCTTACCTAGTACCGTCAGCCCACAATGTTTTCATTTCTTTCCCACCTAAAACTAAAAGTACT
CACAAGTCACAACTTAAACCAAGTACACAAGGATTTTATCATGGGATTATCGTGTTTGAAGACTAAAAAGAGCACACCAT
CACCCCCATTAGTGCAGGTAGAGTAAGACAGTAACTTTTGGGTTCATATTACCGAGCAAGAACCGTTATTTGTGATTAGA
CATGTTATAAACCACTGCTTTAGTGACTATTTAAAACAATATATTACATGTCGTAATCATGCAACCTAACTATGTTTTCA
TTAATCAAATACAAAGAATAAAGAGAAAAGTGCGTAGATTCAATTATTTGGCATAGACTCAAAAGAGTGTATATATATCT
GACTTTTATTAAATTATTAAACACAAATACATATTTTCATAAGCAAAACTATAAAAGCCCTAAACATATAACGATTACCT
CAAAGGAAAAAGTCGTTTTCTCCTAATTAAAAGATAGGTTACTTCCTAATTAATATATAATTTATGTGAACTTCACAATA
TACAGTTCAATAAAATTTGGTAATTTGACCGATTTAAGGAGAGTGGAAATTAGGGCTTCTGCAATCTTTTTTCTTCGCCG
CAATCTCATGTCCAATTATCCACCGACGGTGGCGGCGCAACCCACAACGACGGCGAATCCACTGCTGCAGCGACATCAAT
CTGAACAGCGACGAAGAGAATTACCGAAGATTGTCGAAACAGAGTCTACAAGTATGGACATTACGATCGGTCAATCTAAG
CAGCCTCAATTTTTGAAATCCATAGACGAATTAGCTGCGTTTTCAGTTGCAGTGGAAATATTCAAACGCCAATTCGATGA
TCTTCAGAAGCACATCGAGTCAATCGAAAACGCAATTGATTCCAAACTCGAGAGTAACGGCGTTGTCCTCGCCGCGCGGA
ACAATAATTTCCATCAGCCGATGTTATCGCCTCCGCGGAACAATGTATCTGTAGAAACCACCGTCACTGTGAGCCAACCG
TCTCAGGAGATTGTACCGGAGACGTCGAATAAACCGGAGGGGGGACGTATGTGTGAGTTGATGTGTAGCAAAGGTCTGCG
TAAATACATATAGTCGAATATCTCTGATCAAGCTAAGTTAATGGAAGAGATTCCTTCAGCTTTGAAATTGGCCAAGGAGC
CAGCGAAGTTTGTATTGGATTGTATTGGCAAGTTTACTTACAAGGGCGTAGAGCATTTACTAAAGAGTCGCCTATGAGC
TCTGCGAGACAAGTTTCGCTTCTTATACTGGAGTCTTTTCTTCTAATGCCTGATCGTGGTAAAGGGAAGGTGAAGATTGA
GAGTTGGATTAAAGATGAGGCGGAGACGGCTGCTGTTGCTTGGAGGAAAAGGTTGATGACTGAAGGAGGATTAGCTGCGG
CTGAGAAAATGGATGCAAGGGGTTTGCTTTTACTAGTTGCTTGTTTTGGTGTTCCTTCAAACTTTAGGAGTACAGATTTG
CTGGATTTGATAAGGATGAGTGGTTCGAATGAGATTGCCGGTGCTTTGAAGCGGTCACAGTTTCTTGTCCCTATGGTCTC
AGGTACCATATTCTGTTCTCACTCGGTGAATTTCATTGCAAAGGTGGTTCCTTTTGTTGACATCATCGACCAACATCAAG
TTCCATCTTTGTTTTTCGATAAGCTTGATGGTATAAACTAGGAGAGCACATCAAATATTTAGAGTGCAATGACTGATTGA
GCCAAATCCTAGCTAGAAATTAATCTGGAAAGAACTTGGAACTCTCAACCATAGGTTTTGGTACGAAATTGTTGCTTGTC
AGAACCAAATGATAGGCTATTGCCTTGAAATAGTGTTTCTTGTGGTTTCCAATATTGGAAGTTAAAATCGTATGACTTAG
CTGTTGGATACTAATTAAGCTTAAGCAATGCCAACTCTAAGAAGTGGTACTTACACAATATTCTATTGGTCATAGGTATA
GTTGAATCAAGTATCAAGCGTGGAATGCATATTGAAGCTCTTGAGATGGTTTATACCTTTGGCATGGAGGATAAGTTTTC
AGCTGCTCTAGTTCTAACTTCATTCTTAAAGATGAGCAAGGAGTCATTTGAGAGGGCAAAACGGAAAGCCCAGTCACCGC
TGGCATTTGTATGAACCCTTCCCTTGCACATTATGTACCTTTATGAACTCTTTATCATCATCTGAGTCTGACCATTGATA
TATTTATTTCTCAACAGAAAGAAGCGGCTACAAAGCAGCTAGCTGTGTTATCATCAGTTATGCAGTGTATGGAGACTCAC
AAGTTAGATCCTGCGAAAGAACTACCAGGATGGCAGATCAAAGAGCAAATTGTTAGCTTGGAGAAAGACACTCTTCAGCT
CGACAAAGAGATGGAAGAGAAAGCAAGATCTCTCAGTTTAATGGAGGAAGCCGCACTTGCCAAGAGAATGTATAACCAAC
AGATAAAACGTCCAAGGTTGTCACCCATGGAAATGCCACCAGTAACTTCTTCATCGTATTCTCCTATCTACCGTGATAGA
AGCTTTCCTAGTCAAAGAGACGATGACCAAGATGAAATATCAGCTCTTGTGAGTAGTTACCTCGGCCCGTCAACATCTTT
TCCTCATCGCTCAAGAAGATCCCCGGAATATATGGTTCCACTTCCACATGGTGGGTTAGGAAGAAGTGTATATGCATATG
AACATCTGGCCCCAAATTCATACTCTCCAGGTCACGGACATAGACTTCATCGACAGTACTCTCCGTCTTTGGTTCACGGA
CAGAGACATCCACTACAGTACTCTCCTCCAATTCATGGACAACAACAGTTACCATATGGTATACAAAGGGTTTACAGACA
TTCACCATCTGAAGAAAGATATTTGGGTTTATCCAATCAAAGGTCTCCTCGCAGTAACTCATCATTAGACCCCAAATAGG
AGGAATGTAAATTTGTAACAAAGCTTTTTGTTTTTGCTTAAGTTAGTCATTTATTTAACTCCCAACAGTCTCAAAATTTA
ATTTAATGTTTGGGGCTTAAGAATGCAAATTTTTTTGCTCCTGTAATTGACATTTAAGATGCTAATGTTATTGCTTCAGA
GGTTTTAGTCAACCTCAGATACATCGATATCACTATCTAAATAGACCTCTGGCTCTTGGTCATCTGGATTCTCTTCATCT
TCTGTCTCTGTTCCTTCTTGTTCTCGTTGCACTGCTCGAGCAATTGCGGATTCCAACCTTGTGCTTACAGTTTCCCATGA
CACAAGCTTTTCCATGAATGTATTTATGTCCGCCTTCTTATCTTTCTTGAGGAAGATGAATTC

>FRI_Bg-2

GGATCCCAAAATCTAGTGCCGCCGAGTCCTTTGGGACACAGTTTCGCACTTATCATTCGATTCGGCAAGCAGTAACTCGC
AAAATCCGCACCACCGTGAAACATACTCCGGCGCTGAACGCTAGACGCAAAATCCATATTACAGCTGCAGAAAGACGGAG
AGAACAAGAGACTGACCGATCATAAGAGAAGAGAGCTTCAAGGCTTCAGCTGATGAAGAATCAATGGCGGCGAGAGAGAG
TGAGAATCGAGTAGTTACTGAGAGGAAGATGAAACGTATCTCCTGAGAAAAAACGTAAGAGAGTTCCGTTTCAACAGTAT
TTTTCTCGGCGGTCTTCAGATTTTCCGTCCAACTCTCTCTTTCCTCTTTCAGTTTTTTTTGCCCAATCACTCGTGGTAGG
ATTTTTTATAAGAAAGTTACGAAAATGCCCTCTCGTGCTTAGAGAGATTCAATTCGTGCACATATAATATGCCGTGAATA
TACCACCAACTCTCTTACTCGTACTCGCATGTGGTGCGTGCAATATATACGTGGTTCGAATCAACGGCTACGTTTTACTG
ATAGGTGGCTCGTTTAGTTTTGCACTGCCTTTGGGTGCGCGTGACCAAGTCATTTTCCAAATGTGTTTTAATATTTGTGA
ACTATTATTAATAAAAAAGCGAGCGACGTTGTTCTTTGATGAGGGAACGACCGTACGCTTGGCGACCTTCTCTGATTGCG
TAGAGTCGTTTTCCTTATTGGGCTTACCTAGTACCGTCAGCCCACAATGTTTTCATTTCTTTCCCACCTAAAACTAAAAG
TACTCACAAGTCACAACTTAAACCAAGTACACAAGGATTTTATCATGGGATTATCGTGTTTGAAGACTAAAAAGAGCACA
CCATCACCCCCATTAGTGCAGGTAGAGTAAGACAGTAACTTTTGGGTTCATATTACCGAGCAAGAACCGTTATTTGTGAT
TAGACATGTTATAAACCACTGCTTTAGTGACTATTTAAAACAATATATTACATGTCGTAATCATGCAACCTAACTATGTT
TTCATTAATCAAATACAAAGAATAAAGAGAAAAGTGCGTAGATTCAATTATTTGGCATAGACTCAAAAGAGTGTATATAT
ATCTGACTTTTATTAAATTATTAAACACAAATACATATTTTCATAAGCAAAACTATAAAAGCCCTAAACATATAACGATT
ACCTCAAAGGAAAAAGTCGTTTTCTCCTAATTAAAAGATAGGTTACTTCCTAATTAATATATAATTTATGTGAACTTCAC
AATATACAGTTCAATAAAATTTGGTAATTTGACCGATTTAAGGAGAGTGGAAATTAGGGCTTCTGCAATCTTTTTTCTTC
GCCGCAATCTCATGTCCAATTATCCACCGACGGTGGCGGCGCAACCCACAACGACGGCGAATCCACTGCTGCAGCGACAT
CAATCTGAACAGCGACGAAGAGAATTACCGAAGATTGTCGAAACAGAGTCTACAAGTATGGACATTACGATCGGTCAATC
TAAGCAGCCTCAATTTTTGAAATCCATAGACGAATTAGCTGCGTTTTCAGTTGCAGTGGAAACATTCAAACGCCAATTCG
ATGATCTTCAGAAGCACATCGAGTCAATCGAAAACGCAATTGATTCCAAACTCGAGAGTAACGGCGTTGTCCTCGCCGCG
CGGAACAATAATTTCCATCAGCCGATGTTATCGCCTCCGCGGAACAATGTATCTGTAGAAACCACCGTCACTGTGAGCCA
ACCGTCTCAGGAGATTGTACCGGAGACGTCGAATAAACCGGAGGGGGAACGTATATGTGAGTTGATGTGTAGCAAAGGTC
TGCGTAAATACATATACGCGAATATCTCTGATCAAGCTTAGTTAATGGAAGAGATTCCTTCAGCTTTGAAATTGGCCAAG
GAGCCAGCGAAGTTTGTATTGGATTGTATTGGCAAGTTTTACTTACAAGGGCGTAGAGCATTTACTAAAGAGTCGCCTAT
GAGCTCTGCGAGACAAGTTTCGCTTCTTATACTGGAGTCTTTTCTTCTAATGCCTGATCGTGGTAAAGGGAAGGTGAAGA
TTGAGAGTTGGATTAAAGATGAGGCGGAGACGGCTGCTGTTGCTTGGAGGAAAAGGTTGATGACTGAAGGAGGATTAGCT
GCGGCTGAGAAAATGGATGCAAGGGGTTTGCTTTTACTAGTTGCTTGTTTTGGTGTTCCTTCAAACTTTAGGAGTACAGA
TTTGCTGGATTTGATAAGGATGAGTGGTTCGAATGAGATTGCCGGTGCTTTGAAGCGGTCACAGTTTCTTGTCCCTATGG
TCTCAGGTACCATATTCTGTCTCACTCGGTGAATTTCATTGCAAAGGTGGTTCCTTTTGTTGACATCATCGACCAACATC
AAGTTCCATCTTTGTTTTTCGATAAGCTTGATGGTACAAACTAGGAGAGCACATCAAATATTTAGAGTGCAATGACTGAT
TGAGCCAAATCCTAGCTAGAAATTAATCTGGAAAGAACTTGGAACTCTCAACCATAGGTTTTGGTACGAAATTGTTGCTT
GTCAGAACCAAATGATAGGCTATTGCCTTGAAATAGTGTTTCTTGTGGTTTCCAATATTGGAAGTTAAAATCATATGACT
TAGCTGTTGGATACTAATTAAGCTTAAGCAATGCCAACTCTAAGAAGTGGTACTTACACAATATTCTATTGGTCATAGGT
ATAGTTGAATCAAGTATCAAGCGTGGAATGCATATTGAAGCTCTTGAAATGGTTTATACCTTTGGCATGGAGGATAAGTT
TTCAGCTGCTCTAGTTCTAACTTCATTCTTAAAGATGAGCAAGGAGTCAtTTGAGAGGGCAAAACGGAAAGCCCAGTCAC
CGCTGGCATTTGTATGAACCCTTCCCTTGCACATTATGTACCTTTATGAACTCTTTATCATCATCTGAGTCTGACCATTG
ATATATTTATTTCTCAACAGAAAGAAGCGGCTACAAAGCAGCTAGCTGTGTTATCATCAGTTATGCAGTGTATGGAGACT
CACAAGTTAGATCCTGCGAAAGAACTACCAGGATGGCAGATCAAAGAGCAAATTGTTAGCTTGGAGAAAGACACTCTTCA
GCTCGACAAAGAGATGGAAGAGAAAGCAAGATCTCTCAGTTTAATGGAGGAAGCCGCACTTGCCAAGAGAATGTATAACC
AACAGATAAAACGTCCAAGGTTGTCACCCATGGAAATGCCACCAGTAACTTCTTCATCGTATTCTCCTATCTACCGTGAT
AGAAGCTTTCCTAGTCAAAGAGACGATGACCAAGATGAAATATCAGCTCTTGTGAGTAGTTACCTCGGCCCGTCAACATC
TTTTCCTCATCGCTCAAGAAGATCCCCGGAATATATGGTTCCACTTCCACATGGTGGGTTAGGAAGAAGTGTATATGCAT
ATGAACATCTGGCCCCAAATTCATATTCTCCAGGTCACGGACATAGACTTCATCGACAGTACTCTCCGTCTTTGGTTCAC
GGACAGAGACATCCACTACAGTACTCTCCTCCAATTCATGGACAACAACAGTTACCATATGGTATACAAAGGGTTTACAG
ACATTCACCATCTGAAGAAAGATATTTGGGTTTATCCAATCAAAGGTCTCCTCGCAGTAACTCATCATTAGACCCCAAAT
AGGAGGAATGTAAATTTGTAACAAAGCTTTTTGTTTTTGCTTAAGTTAGTCATTTATTTAACTCCCAACAGTCTCAAAAT
TTAATTTAATGTTTGGGGCTTAAGAATGCAAATTTTTTTGCTCCTGTAATTGACATTTAAGATGCTAATGTTATTGCTTC
AGAGGTTTTAGTCAACCTCAGATACATCGATATCACTATCTAAATAGACCTCTGGCTCTTGGTCATCTGGATTCTCTTCA
TCTTCTGTCTCTGTTCCTTCTTGTTCTCGTTGCACTGCTCGAGCAATTGCGGATTCCAACCTTGTGCTTACAGTTTCCCA
TGACACAAGCTTTTCCATGAATGTATTTATGTCCGCCTTCTTATCTTTCTTGAGGAAGATGAATTC

>FRI_Cvi-0

GGATCCCAAAATCTAGTGCCGCCGAGTCCTTTGGGACAGAGTTTCGCACTTATCATTCGATTCGGCAAGCAGTAACTCGC
AAAATCCGCACCACCGTGAAACATACTCTGGCGCTGAACGCTAGACGCAAAATCCATATTACTGCTGCAGAAAGACGGAG
AGAACAAGAGACTGACCGATCATAAGAGAAGAGAGCTTCAAGGCTTCAGCTGATGAAGAATCAATGGCGGCGAGAGAGAG
TGAGAATCGAGTAGTTACTGAGAGGAAGATGAAACGTATCTCCTGAGAAAAAACGTAAGAGAGTTCCGTTTCAACAGTAT
TTTTCTCGGCGGTCTTCAGATTTTCCGTCCAACTCTCTCTTTCCTCTTTCAGTTTTTTTTTGCCCAATCACTCGTGGTAG
GATTTTTTTATAAGAAAGTTACGAAAATGCCCTCTCGTGCTTAGAGAGATTCAATTCGTGCACATATAATATGCCGTGAA
TATACCACCAACTCTCTTACTCGTACTCGCATGTGGTGCGTGCAATATACACGTGGTTCGAATCAACGGCTACGTTTTAC
TGATAGGTGGCTCGTTTAGTTTTGCACTGCCTTTGGGTGCGCGTGACCAAGTCATTTTCCAAATGGGTTTTAATATTTGT
GAACTATTATTAATAAAAAAGCGAGCGACGTTGTTCTTTGATGAGGGAACGACCGTACGCTTGGCGACCTTCTCTGATTG
CGTAGAGTCGTTTTCCTTATTGGGCTTACCTAGTACCGTCAGCCCACAATGTTTTCATTTCTTTCCCACCTAAAACTAAA
AGTACTCACAAGTCACAACTTAAACCAAGTACACAAGGATTTTATCATGGGATTATCGTGTTTGAAGACTAAAAAGAGCA
CACCATCACCCCCATTAGTGCAGGTAGAGTAAGACAGTAACTTTTGGGTTCATATTACCGAGCAAGAACCGTTATTTGTG
ATTAGACATGTTATAAACCACTGCTTTAGTGACTATTTAAAACAATATATTACATGTCGTAATCATGCAACCTAACTATG
TTTTCACTAAACAAATACAAAGAATAAAGAGAAAAGTGCGTAGATTCAATTATTTGGCATAGACTCAAAAGAGTGTATAT
ATATCTGACTTTTATTAAATTATTAAACACAAATACATATTTTCATAAGCAAAACTATAAAAGCCCTAAACATATAATGA
TTACCTCAAAGGAAAAAGTCGTTTTCTCCTAATTAAAAGATAGGTTACTTCCTAATTAATAGTATATAATTTATGTGAAC
TTCACAATATACAGTTCAATAAAATTTGGTAATTTGACCGATTTAAGGAGAGTGGAAATTAGGGCTTCTGCAATCTTTTT
TCTTCGCCGCAATCTCATGTCCAATTATCCACCGACGGTGGCGGCGCAAACCACAACGACGGCGAATCCACTGCTGCAGC
GACATCAATCTGAACAGCGACGAAGAGAATTACCGAAGATTGTCGAAACAGAGTCTACAAGTATGGACATTACGATCGGT
CAATCTAAGCAGCCTCAATTTTTGAAATCCATAGACGAATTAGCTGCGTTTTCAGTTGCAGTGGAAACATTCAAATGCCA
ATTCGATGATCTTCAGAAGCACATCGAGTCAATCGAAAACGCAATTGATTCCAAACTCGAGAGTAACGGCGTTGTCCTCG
CCGCGCGGAACAATAATTTCCATCAGCCGATGTTATCGCCTCCGCGGAACAATGTATCTGTAGAAACCACCGTCACTGTG
AGCCAACCGTCTCAGGAGATTGTACCGGAGACGTCGAATAAACCGGAGGGGGGACGTATGTGTGAGTTGATGTGTAGCAA
AGGTCTGCGTAAATACATATACGCGAATATCTCTGAACAAGCTAAGTTAATGGAAGAGATTCCTTCAGCTTTGAAATTGG
CCAAGGAGCCAGCGAAGTTTGTATTGGATTGTATTGGCAAGTTTTACTTACAAGGGCGTAGAGCATTTACTAAAGAGTCG
CCTATGAGCTCTGCGAGACAAGTTTCGCTTCTTATACTGGAGTCTTTTCTTCTAATGCCTGATCGTGGTTAAGGGAAGGT
GAAGATTGAGAGTTGGATTAAAGATGAGGCGGAGACGGCTGCTGTTGCTTGGAGGAAAAGGTTGATGACTGAAGGAGGAT
TAGCTGCGGCTGAGAAAATGGATGCAAGGGGTTTGCTTTTACTAGTTGCTTGTTTTGGTGTTCCTTCAAACTTTAGGAGT
ACAGATTTGCTGGATTTGATAAGGATGAGTGGTTCGAATGAGATTGCCGGTGCTTTGAAGCGGTCACAGTTTCTTGTCCC
TATGGTCTCAGGTACCATATTCTGTTCTCACTCGGTGAATTTCATTGCAAAGGTGGTTCCTTTTGTTGACATCATCGACC
AACATCAAGTTCCATCTTTGTTTTTCGATAAGCTTGATGGTATAAACTAGGAGAGCACATCAAATATTTAGAGTGCAATG
ACTGATTGAGCCAAATCCTAGCTAGAAATTAATCTGGAAAGAACTTGGAACTCTCAACCATAGGTTTTGGTACGAAATTG
TTGCTTGTCAGAACCAAATGATAGGCTATTGCCTTGAAATAGTGTTTCTTGTGGTTTCCAATATTGGAAGTTAAAATCAT
ATGACTTAGCTGTTGGATACTAATTAAGCTTAAGCAATGCCAACTCTAAGAAGTGGTACTTACACAATATTCTATTGGTC
ATAGGTATAGTTGAATCAAGTATCAAGCGTGGAATGCATATTGAAGCTCTTGAGATGGTTTATACCTTTGGCATGGAGGA
TAAGTTTTCAGCTGCTCTAGTTCTAACTTCATTCTTAAAGATGAGCAAGGAGTCATTTGAGAGGGCAAAACGGAAAGCCC
AGTCACCGCTGGCATTTGTATGAACCCTTCCCTTGCACATTATGTACCTTTATGAACTCTTTATCATCATCTGAGTCTGA
CCATTGATATATTTATTTCTCAACAGCAAGAAGCGGCTACAAAGCAGCTAGCTGTGTTATCATCAGTTATGCAGTGTATG
GAGACTCACAAGTTAGATCcTGCGAAAGAACTACCAGGATGGCAGATCAAAGAGCAAATTGTTAGCTTGGAGAAAGACAC
TCTTCAGCTCGACAAAGAGATGGAAGAGAAAGCAAGATCTCTCAGTTTAATGGAGGAAGCCGCACTTGCCAAGAGAATGT
ATAACCAACAGATAAAACGTCCAAGGTTGTCACCCATGGAAATGCCACCAGTAACTTCTTCATCGTATTCTCCTATCTAC
CGTGATAGAAGCTTTCCTAGTCAAAGAGACGATGACCAAGATGAAATATCAGCTCTTGTGAGTAGTTACCTCGGCCCGTC
AACATCTTTTCCTCATCGCTCAAGAAGATCCCCGGAATATATGGTTCCACTTCCACATGGTGGGTTAGGAAGAAGTGTAT
ATGCATATGAACATCTGGCCCCAAATTCATACTCTCCAGGTCACGGACATAGACTTCATCGACAGTACTCTCCGTCTTTG
GTTCACGGACAGAGACATCCACTACAGTACTCTCCTCCAATTCATGGACAACAACAGTTACCATATGGTATACAAAGGGt
TTACAGACATTCACCATcTGAAGAAAGATATTTGGGTTTATCCAATCAAAGGTCTCCTCGCAGTAACTCATCATTAGACC
CCAAATAGGAGGAATGTAAATTTGTAACAAAGCTTTTTGTTTTTGCTTAAGTTAGTCATTTATTTAACTCCCAACAGTCT
CAAAATTTAATTTAATGTTTGGGGCTTAAGAATGCAAATTTTTTTGCTCCTGTAATTGACATTTAAGATGCTAATGTTAT
TGCTTCAGAGGTTTTAGTCAACCTCAGATACATCGATATCACTATCTAAATAGACCTCTGGCTCTTGGTCATCTGGATTC
TCTTCATCTTCTGTCTCTGTTCCTTCTTGTTCTCGTTGCACTGCTCGAGCAATTGCGGATTCCAACCTTGTGCTTACAGT
TTCCCATGACACAAGCTTTTCCATGAATGTATTTATGTCCGCCTTCTTATCTTTCTTGAGGAAGATGAATTC

>FRI_NFA-8

GGATCCCAAAATCTAGTGCCGCCGAGTCCTTTGGGACAGAGTTTCGCACTTATCATTCGATTCGGCAAGCAGTAACTCGC
AAAATCCGCACCACCGTGAAACATACTCTGGCGCTGAACGCTAGACGCAAAATCCATATTACTGCTGCAGAAAGACGGAG
AGAACAAGAGACTGACCGATCATAAGAGAAGAGAGCTTCAAGGCTTCAGCTGATGAAGAATCAATGGCGGCGAGAGTGAG
TGAGAATCGAGTAGTTACTGAGAGGAAGATGAAACGTATCTCCTGAGAAAAAACGTAAGAGAGTTCCGTTTCAACAGTAT
TTTTCTCGGCGGTCTTCAGATTTTCCGTCCAACTCTCTCTTTCCTCTTTCAGTTTTTTTTTGCCCAATCACTCGTGGTAG
GATTTTTTTATAAGAAAGTTACGAAAATGCCCTCTCGTGCTTAGAGAGATTCAATTCGTGCACATATAATATGCCGTGAA
TATACCACCAACTCTCTTACTCGTACTCGCATGTGGTGCGTGCAATATACACGTGGTTCGAATCAACGGCTACGTTTTAC
TGATAGGTGGCTCGTTTAGTTTTGCACTGCCTTTGGGTGCGCGTGACCAAGTCATTTTCCAAATGTGTTTTAATATTTGT
GAACTATTATTAATAAAAAAGCGAGCGACGTTGTTCTTTGATGAGGGAACGACCGTACGCTTGGCGACCTTCTCTGATTG
CGTAGAGTCGTTTTCCTTATTGGGCTTACCTAGTACCGTCAGCCCACAATGTTTTCATTTCTTTCCCACCTAAAACTAAA
AGTACTCACAAGTCACAACTTAAACCAAGTACACAAGGATTTTATCATGGGATTATCGTGTTTGAAGACTAAAAAGAGCA
CACCATCACCCCCATTAGTGCAGGTAGAGTAAGACAGTAACTTTTGGGTTCATATTACCGAGCAAGAACCGTTATTTGTG
ATTAGACATGTTATAAACCACTGCTTTAGTGACTATTTAAAACAATATATTACATGTCGTAATCATGCAACCTAACTATG
TTTTCATTAATCAAATACAAAGAATAAAGAGAAAAGTGCGTAGATTCAATTATTTGGCATAGACTCAAAAGAGTGTATAT
ATATCTGACTTTTATTAAATTATTAAACACAAATACATATTTTCATAAGCAAAACTATAAAAGCCCTAAACATATAATGA
TTACCTCAAAGGAAAAAGTCGTTTTCTCCTAATTAAAAGATAGGTTACTTCCTAATTAATATATAATTTATGTGAACTTC
ACAATATACAGTTCAATAAAATTTGGTAATTTGACCGATTTAAGGAGAGTGGAAATTAGGGCTTCTGCAATCTTTTTTCT
TCGCCGCAATCTCATGTCCAATTATCCACCGACGGTGGCGGCGCAACCCACAACGACGGCGAATCCACTGCTGCAGCGAC
ATCAATCTGAACAGCGACGAAGAGAATTACCGAAGATTGTCGAAACAGAGTCTACAAGTATGGACATTATGATCGGTCAA
TCTAAGCAGCCTCAATTTTTGAAATCCATAGACGAATTAGCTGCGTTTTCAGTTGCAGTGGAAACATTCAAACGCCAATT
CGATGATCTTCAGAAGCACATCGAGTCAATCGAAAACGCAATTGATTCCAAACTCGAGAGTAACGGCGTTgTCCTCGCCG
CGCGGAACAATAATTTCCATCAGCCGATGTTATCGCCTCCGCGGAACAATGTATCTGTAGAAACCACCGTCACTGTGAGC
CAACCGTCTCAGGAGATTGTACCGGAGACGTCGAATAAACCGGAGGGGGAACGTATGTGTGAGTTGATGTGTAGCAAAGG
TCTGCGTAAATACATATACGCGAATATCTCTGATCAAGCTAAGTTAATGGAAGAGATTCCTTCAGCTTTGAAATTGGCCA
AGGAGCCAGCGAAGTTTGTATTGGATTGTATTGGCAAGTTTTACTTACAAGGGCGTAGAGCATTTACTAAAGAGTCGCCT
ATGAGCTCTGCGAGACAAGTTTCGCTTCTTATACTGGAGTCTTTTCTTCTAATGCCTGATCGTGGTAAAGGGAAGGTGAA
GATTGAGAGTTGGATTAAAGATGAGGCGGAGACGGCTGCTGTTGCTTGGAGGAAAAGGTTGATGACTGAAGGAGGATTAG
CTGCGGCTGAGAAAATGGATGCAAGGGGTTTGCTTTTACTAGTTGCTTGTTTTGGTGTTCCTTCAAACTTTAGGAGTACG
GATTTGCTGGATTTTGATAAGGATGAGTGGTTCGAATGAGATTGCCGGTGCTTTGAAGCGGTCACAGTTTCTTGTCCCTA
TGGTCTCAGGTACCATATTCTGTTCTCACTCGGTGAATTTCATTGCAAAGGTGGTTCCTTTTGTTGACATCATCGACCAA
CATCAAGTTCCATCTTTGTTTTTCGATAAGCTTGATGGTATAAACTAGGAGAGCACATCAAATATTTAGAGTGCAATGAC
TGATTGAGCCAAATCCTAGCTAGAAATTAATCTGGAAAGAACTTGGAACTCTCAACCATAGGTTTTGGTACGAAATTGTT
GCTTGTCAGAACCAAATGATAGGCTATTGCCTTGAAATAGTGTTTCTTGTGGTTTCCAATATTGGAAGTTAAAATCATAT
GACTTAGCTGTTGGATACTAATTAAGCTTAAGCAATGCCAACTCTAAGAAGTGGTACTTACACAATATTCTATTGGTCAT
AGGTATAGTTGAATCAAGTATCAAGCGTGGAATGCATATTGAAGCTCTTGAAATGGTTTATACCTTTGGCATGGAGGATA
AGTTTTCAGCTGCTCTAGTTCTAACTTCATTCTTAAAGATGAGCAAGGAGTCATTTGAGAGGGCAAAACGGAAAGCCCAG
TCACCGCTGGCATTTGTATGAACCCTTCCCTTGCACATTATGTACCTTTATGAACTCTTTATCATCATCTGAGTCTGACC
ATTGATATATTTATTTCTCAACAGAAAGAAGCGGCTACAAAGCAGCTAGCTGTGTTATCATCAGTTATGCAGTGTATGGA
GACTCACAAGTTAGATCCTGCGAAAGAACTACCAGGATGGCAGATCAAAGAGCAAATTGTTAGCTTGGAGAAAGACACTC
TTCAGCTCGACAAAGAGATGGAAGAGAAAGCAAGATCTCTCAGTTTAATGGAGGAAGCCGCACTTGCCAAGAGAATGTAT
AACCAACAGATAAAACGTCCAAGGTTGTCACCCATGGAAATGCCACCAGTAACTTCTTCATCGTATTCTCCTATCTACCG
TGATAGAAGCTTTCCTAGTCAAAGAGACGATGACCAAGATGAAATATCAGCTCTTGTGAGTAGTTACCTCGGCCCGTCAA
CATCTTTTCCTCATCGCTCAAGAAGATCCCCGGAATATATGGTTCCACTTCCACATGGTGGGTTAGGAAGAAGTGTATAT
GCATATGAACATCTGGCCCCAAATTCATATTCTCCAGGTCACGGACATAGACTTCATCGACAGTACTCTCCGTCTTTGGT
TCACGGACAGAGACATCCACTACAGTACTCTCCTCCAATTCATGGACAACAACAGTTACCATATGGTATACAAAGGGTTT
ACAGACATTCACCATCTGAAGAAAGATATTTGGGTTtATCCAATCAAAGGTCTCCTCGCAGTAACTCATCATTAGACCCC
AAATAGGAGGAATGTAAATTTGTAACAAAGCTTTTTGTTTTTGCTTAAGTTAGTCATTTATTTAACTCCCAACAGTCTCA
AAATTTAATTTAATGTTTGGGGCTTAAGAATGCAAATTTTTTTGCTCCTGTAATTGACATTTAAGATGCTAATGTTATTG
CTTCAGAGGTTTTAGTCAACCTCAGATACATCGATATCACTATCTAAATAGACCTCTGGCTCTTGGTCATCTGGATTCTC
TTCATCTTCTGTCTCTGTTCCTTCTTGTTCTCGTTGCACTGCTCGAGCAATTGCGGATTCCAACCTTGTGCTTACAGTTT
CCCATGACACAAGCTTTTCCATGAATGTATTTATGTCCGCCTTCTTATCTTTCTTGAGGAAGATGAATTC

>FRI_Ull-2-3

GGATCCCAAAATCTAGTGCCGCCGAGTCCTTTGGGACACAGTTTCGCACTTATCATTCGATTCGGCAAGCAGTAACTCGCAAAA
TCCGCACCACCGTGAAACATACTCCGGCGCTGAACGCTAGACGCAAAATCCATATTACAGCTGCAGAAAGACGGAGAGAACA
AGAGACTGACCGATCATAAGAGAAGAGAGCTTCAAGGCTTCAGCTGATGAAGAATCAATGGCGGCGAGAGAGAGTGAGAAT
CGAGTAGTTACTGAGAGGAAGATGAAACGTATCTCCTGAGAAAAAACGTAAGAGAGTTCCGTTTCAACAGTATTTTTCTC
GGCGGTCTTCAGATTTTCCGTCCAACTCTCTCTTTCCTCTTTCAGTTTTTTTGCCCAATCACTCGTGGTAGGATTTTTT
ATAAGAAAGTTACGAAAATGCCCTCTCGTGCTTAGAGAGATTCAATTCGTGCACATATAATATGCCGTGAATATACCACC
AACTCTCTTACTCGTACTCGCATGTGGTGCGTGCAATATACACGTGGTTCGAATCAACGGCTACGTTTTACTGATAGGTG
GCTCGTTTAGTTTTGCACTGCCTTTGGGTGCGCGTGACCAAGTCATTTTCCAAATGTGTTTTAATATTTGTGAACTATTA
TTAATAAAAAAGCGAGCGACGTTGTTCTTTGATGAGGGAACGACCGTACGCTTGGCGACCTTCTCTGATTGCGTAGAGTC
GTTTTCCTTACTGGGCTTACCTAGTACCGTCAGCCCACAATGTTTTCATTTCTTTCCCACCTAAAACTAAAAGTACTCAC
AAGTCACAACTTAAACCAAGTACACAAGGATTTTATCATGGGATTATCGTGTTTGAAGACTAAAAAGAGCACACCATCAC
CCCCATTAGTGCAGGTAGAGTAAGACAGTAACTTTTGGGTTCATATTACCGAGCAAGAACCGTTATTTGTGATTAGACAT
GTTATAAACCACTGCTTTAGTGACTATTTAAAACAATATATTACATGTCGTAATCATGCAACCTAACTATGTTTTCATTA
ATCAAATACAAAGAATAAAGAGAAAAGTGCGTAGATTCAATTATTTGGCATAGACTCAAAAGAGTGTATATATATCTGAC
TTTTATTAAATTATTAAACACAAATACATATTTTCATAAGCAAAACTATAAAAGCCCTAAACATATAATGATTACCTCAA
AGGAAAAAGTCGTTTTCTCCTAATTAAAAGATAGGTTACTTCCTAATTAATAGTATATAATTTATGTGAACTTCACAATA
TACAGTTCAATAAAATTTGGTAATTTGACCGATTTAAGGAGAGTGGAAATTAGGGCTTCTGCAATCTTTTTTCTTCGCCG
CAATCTCATGTCCAATTATCCACCGACGGTGGCGGCGCAACCCACAACGACGGCGAATCCACTGCTGCAGCGACATCAAT
CTGAACAGCGACGAAGAGAATTACCGAAGATTGTCGAAACAGAGTCTACAAGTATGGACATTACGATCGGTCAATCTAAG
CAGCCTCAATTTTTGAAATCCATAGACGAATTAGCTGCGTTTTCAGTTGCAGTGGAAACATTCAAATGCCAATTCGATGA
TCTTCAGAAGCACATCGAGTCAATCGAAAACGCAATTGATTCCAAACTCGAGAGTAACGGCGTTGTCCTCGCCGCGCGGA
ACAATAATTTCCATCAGCCGATGTTATCGCCTCCGCGGAACAATGTATCTGTAGAAACCACCGTCACTGTGAGCCAACCG
TCTCAGGAGATTGTACCGGAGACGTCGAATAAACCGGAGGGGGGACGTATGTGTGAGTTGATGTGTAGCAAAGGTCTGCG
TAAATACATATACGCGAATATCTCTGAACAAGCTAAGTTAATGGAAGAGATTCCTTCAGCTTTGAAATTGGCCAAGGAGC
CAGCGAAGTTTGTATTGGATTGTATTGGCAAGTTTTACTTACAAGGGCGTAGAGCATTTACTAAAGAGTCGCCTATGAGC
TCTGCGAGACAAGTTTCGCTTCTTATACTGGAGTCTTTTCTTCTAATGCCTGATCGTGGTAAAGGGAAGGTGAAGATTGA
GAGTTGGATTAAAGATGAGGCGGAGACGGCTGCTGTTGCTTGGAGGAAAAGGTTGATGACTGAAGGAGGATTAGCTGCGG
CTGAGAAAATGGATGCAAGGGGTTTGCTTTTACTAGTTGCTTGTTTTGGTGTTCCTTCAAACTTTAGGAGTACAGATTTG
CTGGATTTGATAAGGATGAGTGGTTCGAATGAGATTGCCGGTGCTTTGAAGCGGTCACAGTTTCTTGTCCCTATGGTCTC
AGGTACCATATTCTGTtCTCACTCGGTGAATTTCATTGCAAAGGTGGTTCCTTTTGTTGACATCATCGACCAACATCAAG
TTCCATCTTTGTTTTTCGATAAGCTTGATGGTATAAACTAGGAGAGCACATCAAATATTTAGAGTGCAATGACTGATTGA
GCCAAATCCTAGCTAGAAATTAATCTGGAAAGAACTTGGAACTCTCAACCATAGGTTTTGGTACGAAATTGTTGCTTGTC
AGAACCAAATGATAGGCTATTGCCTTGAAATAGTGTTTCTTGTGGTTTCCAATATTGGAAGTTAAAATCATATGACTTAG
CTGTTGGATACTAATTAAGCTTAAGCAATGCCAACTCTAAGAAGTGGTACTTACACAATATTCTATTGGTCATAGGTATA
GTTGAATCAAGTATCAAGCGTGGAATGCATATTGAAGCTCTTGAGATGGTTTATACCTTTGGCATGGAGGATAAGTTTTC
AGCTGCTCTAGTTCTAACTTCATTCTTAAAGATGAGCAAGGAGTCATTTGAGAGGGCAAAACGGAAAGCCCAGTCACCGC
TGTTGTATGAACCCTTCCCTTGCACATTATGTACCTTTATGAACTCTTTATCATCATCTGAGTCTGACCATTGATATATT
TATTTCTCAACAGAAAGAAGCGGCTACAAAGCAGCTAGCTGTGTTATCATCAGTTATGCAGTGTATGGAGACTCACAAGT
TAGATCCTGCGAAAGAACTACCAGGATGGCAGATCAAAGAGCAAATTGTTAGCTTGGAGAAAGACACTCTTCAGCTCGAC
AAAGAGATGGAAGAGAAAGCAAGATCTCTCAGTTTAATGGAGGAAGCCGCACTTGCCAAGAGAATGTATAACCAACAGAT
AAAACGTCCAAGGTTGTCACCCATGGAAATGCCACCAGTAACTTCTTCATCGTATTCTCCTATCTACCGTGATAGAAGCT
TTCCTAGTCAAAGAGACGATGACCAAGATGAAATATCAGCTCTTGTGAGTAGTTACCTCGGCCCGTCAACATCTTTTCCT
CATCGCTCAAGAAGATCCCCGGAATATATGGTTCCACTTCCACATGGTGGGTTAGGAAGAAGTGTATATGCATATGAACA
TCTGGCCCCAAATTCATACTCTCCAGGTCACGGACATAGACTTCATCGACAGTACTCTCCGTCTTTGGTTCACGGACAGA
GACATCCACTACAGTACTCTCCTCCAATTCATGGACAACAACAGTTACCATATGGTATACAAAGGGtTTACAGACATTCA
CCATCTGAAGAAAGATATTTGGGTTTATCCAATCAAAGGTCTCCTCGCAGTAACTCATCATTAGACCCCAAATAGGAGGA
ATGTAAATTTGTAACAAAGCTTTTTGTTTTTGCTTAAGTTAGTCATTTATTTAACTCCCAACAGTCTCAAAATTTAATTT
AATGTTTGGGGCTTAAGAATGCAAATTTTTTTGCTCCTGTAATTGACATTTAAGATGCTAATGTTATTGCTTCAGAGGTT
TTAGTCAACCTCAGATACATCGATATCACTATCTAAATAGACCTCTGGCTCTTGGTCATCTGGATTCTCTTCATCTTCTG
TCTCTGTTCCTTCTTGTTCTCGTTGCACTGCTCGAGCAATTGCGGATTCCAACCTTGTGCTTACAGTTTCCCATGACACA
AGCTTTTCCATGAATGTATTTATGTCCGCCTTCTTATCTTTCTTGAGGAAGATGAATTC

>FRI_An-1

GGATCCCAAAATCTAGTGCCGCCGAGTCCTTTGGGACAGAGTTTCGCACTTATCATTCGATTCGGCAAGCAGTAACTCGC
AAAATCCGCACCACCGTGAAACATACTCTGGCGCTGAACGCTAGACGCAAAATCCATATTACTGCTGCAGAAAGACGGAG
AGAACAAGAGACTGACCGATCATAAGAGAAGAGAGCTTCAAGGCTTCAGCTGATGAAGAATCAATGGCGGCGAGAGTGAG
TGAGAATCGAGTAGTTACTGAGAGGAAGATGAAACGTATCTCCTGAGAAAAAACGTAAGAGAGTTCCGTTTCAACAGTAT
TTTTCTCGGCGGTCTTCAGATTTTCCGTCCAACTCTCTCTTTCCTCTTTCAGTTTTTTTTTGCCCAATCACTCGTGGTAG
GATTTTTTTATAAGAAAGTTACGAAAATGCCCTCTCGTGCTTAGAGAGATTCAATTCGTGCACATATAATATGCCGTGAA
TATACCACCAACTCTCTTACTCGTACTCGCATGTGGTGCGTGCAATATACACGTGGTTCGAATCAACGGCTACGTTTTAC
TGATAGGTGGCTCGTTTAGTTTTGCACTGCCTTTGGGTGCGCGTGACCAAGTCATTTTCCAAATGTGTTTTAATATTTGT
GAACTATTATTAATAAAAAAGCGAGCGACGTTGTTCTTTGATGAGGGAACGACCGTACGCTTGGCGACCTTCTCTGATTG
CGTAGAGTCGTTTTCCTTATTGGGCTTACCTAGTACCGTCAGCCCACAATGTTTTCATTTCTTTCCCACCTAAAACTAAA
AGTACTCACAAGTCACAACTTAAACCAAGTACACAAGGATTTTATCATGGGATTATCGTGTTTGAAGACTAAAAAGAGCA
CACCATCACCCCCATTAGTGCAGGTAGAGTAAGACAGTAACTTTTGGGTTCATATTACCGAGCAAGAACCGTTATTTGTG
ATTAGACATGTTATAAACCACTGCTTTAGTGACTATTTAAAACAATATATTACATGTCGTAATCATGCAACCTAACTATG
TTTTCATTAATCAAATACAAAGAATAAAGAGAAAAGTGCGTAGATTCAATTATTTGGCATAGACTCAAAAGAGTGTATAT
ATATCTGACTTTTATTAAATTATTAAACACAAATACATATTTTCATAAGCAAAACTATAAAAGCCCTAAACATATAACGA
TTACCTCAAAGGAAAAAGTCGTTTTCTCCTAATTAAAAGATAGGTTACTTCCTAATTAATATATAATTTATGTGAACTTC
ACAATATACAGCTCAATAAAATTTGGTAATTTGACCGATTTAAGGAGAGTGGAAATTAGGGCTTCTGCAATCTTTTTTCT
TCGCCGCAATCTCATGTCCAATTATCCACCGACGGTGGCGGCGCAACCCACAACGACGGCGAATCCACTGCTGCAGCGAC
ATCAATCTGAACAGCGACGAAGAGAATTACCGAAGATTGTCGAAACAGAGTCTACAAGTATGGACATTACGATCGGTCAA
TCTAAGCAGCCTCAATTTTTGAAATCCATAGACGAATTAGCTGCGTTTTCAGTTGCAGTGGAAACATTCAAACGCCAATT
CGATGATCTTCAGAAGCACATCGAGTCAATCGAAAACGCAATTGATTCCAAACTCGAGAGTAACGGCGTTGTCCTCGCCG
CGCGGAACAATAATTTCCATCAGCCGATGTTATCGCATCCGCGGAACAATGTATCTGTAGAAACCACCGTCACTGTGAGC
CAACCGTCTCAGGAGATTGTACCGGAGACGTCGAATAAACCGGAGGGGGGACGTATGTGTGAGTTGATGTGTAGCAAAGG
TCTGCGTAAATACATATACGCGAATATCTCTGATCAAGCTAAGTTAATGGAAGAGATTCCTTCAGCTTTGAAATTGGCCA
AGGAGCCAGCGAAGTTTGTATTGGATTGTATTGGCAAGTTTTACTTACAAGGGCGTAGAGCATTTACTAAAGAGTCGCCT
ATGAGCTCTGCGAGACAAGTTTCGCTTCTTATACTGGAGTCTTTTCTTCTAATGCCTGATCGTGGTAAAGGGAAGGTGAA
GATTGAGAGTTGGATTAAAGATGAGGCGGAGACGGCTGCTGTTGCTTGGAGGAAAAGGTTGATGACTGAAGGAGGATTAG
CTGCGGCTGAGAAAATGGATGCAAGGGGTTTGCTTTTACTAGTTGCTTGTTTTGGTGTTCCTTCAAACTTTAGGAGTACA
GATTTGCTGGATTTGATAAGGATGAGTGGTTCGAATGAGATTGCCGGTGCTTTGAAGCGGTCACAGTTTCTTGTCCCTAT
GGTCTCAGGTACCATATTCTGTTCTCACTCGGTGAATTTCATTGCAAAGGTGGTTCCTTTTGTTGACATCATCGACCAAC
ATCAAGTTCCATCTTTGTTTTTCGATAAGCTTGATGGTATAAACTAGGAGAGCACATCAAATATTTAGAGTGCAATGACT
GATTGAGCCAAATCCTAGCTAGAAATTAATCTGGAAAGAACTTGGAACTCTCAACCATAGGTTTTGGTACGAAATTGTTG
CTTGTCAGAACCAAATGATAGGCTATTGCCTTGAAATAGTGTTTCTTGTGGTTTCCAATATTGGAAGTTAAAATCATATG
ACTTAGCTGTTGGATACTAATTAAGCTTAAGCAATGCCAACTCTAAGAAGTGGTACTTACACAATATTCTATTGGTCATA
GGTATAGTTGAATCAAGTATCAAGCGTGGAATGCATATTGAAGCTCTTGAGATGGTTTATACCTTTGGCATGGAGGATAA
GTTTTCAGCTGCTCTAGTTCTAACTTCATTCTTAAAGATGAGCAAGGAGTCATTTGAGAGGGCAAAACGGAAAGCCCAGT
CACCGCTGGCATTTGTATGAACCCTTCCCTTGCACATTATGTACCTTTATGAACTCTTTATCATCATCTGAGTCTGACCA
TTGATATATTTATTTCTCAACAGAAAGAAGCGGCTACAAAGCAGCTAGCTGTGTTATCATCAGTTATGCAGTGTATGGAG
ACTCACAAGTTAGATCTTGCTTTCTCTTCCATCTCTTTGTCGAGCTGAAGAGTGTCTTTCTCCAAGCTAACAATTTGCTC
TCAGTTTAATGGAGGAAGCCGCACTTGCCAAGAGAATGTATAACCAACAGATAAAACGTCCAAGGTTGTCACCCATGGAA
ATGCCACCAGTAACTTCTTCATCGTATTCTCCTATCTACCGTGATAGAAGCTTTCCTAGTCAAAGAGACGATGACCAAGA
TGAAATATCAGCTCTTGTGAGTAGTTACCTCGGCCCGTCAACATCTTTTCCTCATCGCTCAAGAAGATCCCCGGAATATA
TGGTTCCACTTCCACATGGTGGGTTAGGAAGAAGTGTATATGCATATGAACATCTGGCCCCAAATTCATACTCTCCAGGT
CACGGACATAGACTTCATCGACAGTACTCTCCGTCTTTGGTTCACGGACAGAGACATCCACTACAGTACTCTCCTCCAAT
TCATGGACAACAACAGTTACCATATGGTATACAAAGGGTTTACAGACATTCACCATCTGAAGAAAGATATTTGGGTTTAT
CCAATCAAAGGTCTCCTCGCAGTAACTCATCATTAGACCCCAAATAGGAGGAATGTAAATTTGTAACAAAGCTTTTTGTT
TTTGCTTAAGTTAGTCATTTATTTAACTCCCAACAGTCTCAAAATTTAATTTAATGTTTGGGGCTTAAGAATGCAAATTT
TTTTGCTCCTGTAATTGACATTTAAGATGCTAATGTTATTGCTTCAGAGGTTTTAGTCAACCTCAGATACATCGATATCA
CTATCTAAATAGACCTCTGGCTCTTGGTCATCTGGATTCTCTTCATCTTCTGTCTCTGTTCCTTCTTGTTCTCGTTGCAC
TGCTCGAGCAATTGCGGATTCCAACCTTGTGCTTACAGTTTCCCATGACACAAGCTTTTCCATGAATGTATTTATGTCCG
CCTTCTTATCTTTCTTGAGGAAGATGAATTC

>FRI_Pro-0

GGATCCCAAAATCTAGTGCCGCCGAGTCCTTTGGGACACAGTTTCGCACTTATCATTCGATTCGGCAAGCAGTAACTCGC
AAAATCCGCACCACCGTGAAACATACTCCGGCGCTGAACGCTAGACGCAAAATCCATATTACAGCTGCAGAAAGACGGAG
AGAACAAGAGACTGACCGATCATAAGAGAAGAGAGCTTCAAGGCTTCAGCTGATGAAGAATCAATGGCGGCGAGAGAGAG
TGAGAATCGAGTAGTTACTGAGAGGAAGATGAAACGTATCTCCTGAGAAAAAACGTAAGAGAGTTCCGTTTCAACAGTAT
TTTTCTCGGCGGTCTTCAGATTTTCCGTCCAACTCTCTCTTTCCTCTTTCAGTTTTTTTTTGCCCAATCACTCGTGGTAG
GATTTTTTATAAGAAAGTTACGAAAATGCCCTCTCGTGCTTAGAGAGATTCAATTCGTGCACATATAATATGCCGTGAAT
ATACCACCAACTCTCTTACTCGTACTCGCATGTGGTGCGTGCAATATACACGTGGTTCGAATCAACGGCTACGTTTTACT
GATAGGTGGCTCGTTTAGTTTTGCACTGCCTTCGGGTGCGCGTGACCAAGTCATTTTCCAAATGTGTTTTAATATTTGTG
AACTATTATTAATAAAAAAGCGAGCGACGTTGTTCTTTGATGAGGGAACGACCGTACGCTTGGCGACCTTCTCTGATTGC
GTAGAGTCGTTTTCCTTATTGGGCTTACCTAGTACCGTCAGCCCACAATGTTTTCATTTCTTTCCCACCTAAAACTAAAA
GTACTCACAAGTCACAACTTAAACCAAGTACACAAGGATTTTATCATGGGATTATCGTGTTTGAAGACTAAAAAGAGCAC
ACCATCACCCCCATTAGTGCAGGTAGAGTAAGACAGTAACTTTTGGGTTCATATTACCGAGCAAGAACCGTTATTTGTGA
TTAGACATGTTATAAACCACTGCTTTAGTGACTATTTAAAACAATATATTACATGTCGTAATCATGCAACCTAACTATGT
TTTCATTAATCAAATACAAAGAATAAAGAGAAAAGTGCGTAGATTCAATTATTTGGCATAGACTCAAAAGAGTGTATATA
TATCTGACTTTTATTAAATTATTAAACACAAATACATATTTTCATAAGCAAAACTATAAAAGCCCTAAACATATAATGAT
TACCTCAAAGGAAAAAGTCGTTTTCTCCTAATTAAAAGATAGGTTACTTCCTAATTAATAGTATATAATTTATGTGAACT
TCACAATATACAGTTCAATAAAATTTGGTAATTTGACCGATTTAAGGAGAGTGGAAATTAGGGCTTCTGCAATCTTTTTT
CTTCGCCGCAATCTCATGTCCAATTATCCACCGACGGTGGCGGCGCAACCCACAACGACGGCGAATCCACTGCTGCAGCG
ACATCAATCTGAACAGCGACGAAGAGAATTACCGAAGATTGTCGAAACAGAGTCTACAAGTATGGACATTACGATCGGTC
AATCTAAGCAGCCTCAATTTTTGAAATCCATAGACGAATTAGCTGCGTTTTCAGTTGCAGTGGAAACATTCAAATGCCAA
TTCGATGATCTTCAGAAGCACATCGAGTCAATCGAAAACGCAATTGATTCCAAACTCGAGAGTAACGGCGTTGTCCTCGC
CGCGCGGAACAATAATTTCCATCAGCCGATGTTATCGCCTCCGCGGAACAATGTATCTGTAGAAACCACCGTCACTGTGA
GCCAACCGTCTCAGGAGATTGTACCGGAGACGTCGAATAAACCGGAGGGGGGACGTATGTGTGAGTTGATGTGTAGCAAA
GGTCTGCGTAAATACATATACGCGAATATCTCTGAACAAGCTAAGTTAATGGAAGAGATTCCTTCAGCTTTGAAATTGGC
CAAGGAGCCAGCGAAGTTTGTATTGGATTGTATTGGCAAGTTTTACTTACAAGGGCGTAGAGCATTTACTAAAGAGTCGC
CTATGAGCTCTGCGAGACAAGTTTCGCTTCTTATACTGGAGTCTTTTCTTCTAATGCCTGATCGTGGTAAAGGGAAGGTG
AAGATTGAGAGTTGGATTAAAGATGAGGCGGAGACGGCTGCTGTTGCTTGGAGGAAAAGGTTGATGACTGAAGGAGGATT
AGCTGCGGCTGAGAAAATGGATGCAAGGGGTTTGCTTTTACTAGTTGCTTGTTTTGGTGTTCCTTCAAACTTTAGGAGTA
CAGATTTGCTGGATTTGATAAGGATGAGTGGTTCGAATGAGATTGCCGGTGCTTTGAAGCGGTCACAGTTTCTTGTCCCT
ATGGTCTCAGGTACCATATTCTGTTCTCACTCGGTGAATTTCATTGCAAAGGTGGTTCCTTTTGTTGACATCATCGACCA
ACATCAAGTTCCATCTTTGTTTTTCGATAAGCTTGATGGTATAAACTAGGAGAGCACATCAAATATTTAGAGTGCAATGA
CTGATTGAGCCAAATCCTAGCTAGAAATTAATCTGGAAAGAACTTGGAACTCTCAACCATAGGTTTTGGTACGAAATTGT
TGCTTGTCAGAACCAAATGATAGGCTATTGCCTTGAAATAGTGTTTCTTGTGGTTTCCAATATTGGAAGTTAAAATCATA
TGACTTAGCTGTTGGATACTAATTAAGCTTAAGCAATGCCAACTCTAAGAAGTGGTACTTACACAATATTCTATTGGTCA
TAGGTATAGTTGAATCAAGTATCAAGCGTGGAATGCATATTGAAGCTCTTGAGATGGTTTATACCTTTGGCATGGAGGAT
AAGTTTTCAGCTGCTCTAGTTCTAACTTCATTCTTAAAGATGAGCAAGGAGTCATTTGAGAGGGCAAAACGGAAAGCCCA
GTCACCGCTGGCATTTGTATGAACCCTTCCCTTGCACATTATGTACCTTTATGAACTCTTTATCATCATCTGAGTCTGAC
CATTGATATATTTATTTCTCAACAGAAAGAAGCGGCTACAAAGCAGCTAGCTGTGTTATCATCAGTTATGCAGTGTATGG
AGACTCACAAGTTAGATCTTGCTTTCTCTTCCATCTCTTTGTCGAGCTGAAGAGTGTCTTTCTCCAAGCTAACAATTTGC
TCTCAGTTTAATGGAGGAAGCCGCACTTGCCAAGAGAATGTATAACCAACAGATAAAACGTCCAAGGTTGTCACCCATGG
AAATGCCACCAGTAACTTCTTCATCGTATTCTCCTATCTACCGTGATAGAAGCTTTCCTAGTCAAAGAGACGATGACCAA
GATGAAATATCAGCTCTTGTGAGTAGTTACCTCGGCCCGTCAACATCTTTTCCTCATCGCTCAAGAAGATCCCCGGAATA
TATGGTTCCACTTCCACATGGTGGGTTAGGAAGAAGTGTATATGCATATGAACATCTGGCCCCAAATTCATACTCTCCAG
GTCACGGACATAGACTTCATCGACAGTACTCTCCGTCTTTGGTTCACGGACAGAGACATCCACTACAGTACTCTCCTCCA
ATTCATGGACAACAACAGTTACCATATGGTATACAAAGGGTTTACAGACATTCACCATCTGAAGAAAGATATTTGGGTTT
ATCCAATCAAAGGTCTCCTCGCAGTAACTCATCATTAGACCCCAAATAGGAGGaATGTAAATTTGTAACAAAGCTTTTTG
TTTTTGCTTAAGTTAGTCATTTATTTAACTCCCAACAGTCTCAAAATTTAATTTAATGTTTGGGGCTTAAGAATGCAAAT
TTTTTTGCTCCTGTAATTGACATTTAAGATGCTAATGTTATTGCTTCAGAGGTTTTAGTCAACCTCAGATACATCGATAT
CACTATCTAAATAGACCTCTGGCTCTTGGTCATCTGGATTCTCTTCATCTTCTGTCTCTGTTCCTTCTTGTTCTCGTTGC
ACTGCTCGAGCAATTGCGGATTCCAACCTTGTGCTTACAGTTTCCCATGACACAAGCTTTTCCATGAATGTATTTATGTC
CGCCTTCTTATCTTTCTTGAGGAAGATGAATTC

>FRI_NFA-10

GGATCCCAAAATCTAGTGCCGCCGAGTCCTTTGGGACACAGTTTCGCACTTATCATTCGATTCGGCAAGCAGTAACTCGC
AAAATCCGCACCACCGTGAAACATACTCCGGCGCTGAACGCTAGACGCAAAATCCATATTACAGCTGCAGAAAGACGGAG
AGAACAAGAGACTGACCGATCATAAGAGAAGAGAGCTTCAAGGCTTCAGCTGATGAAGAATCAATGGCGGCGAGAGAGAG
TGAGAATCGAGTAGTTACTGAGAGGAAGATGAAACGTATCTCCTGAGAAAAAACGTAAGAGAGTTCCGTTTCAACAGTAT
TTTTCTCGGCGGTCTTCAGATTTTCCGTCCAACTCTCTCTTTCCTCTTTCAGTTTTTTTTTGCCCAATCACTCGTGGTAG
GATTTTTTATAAGAAAGTTACGAAAATGCCCTCTCGTGCTTAGAGAGATTCAATTCGTGCACATATAATATGCCGTGAAT
ATACCACCAACTCTCTTACTCGTACTCGCATGTGGTGCGTGCAATATACACGTGGTTCGAATCAACGGCTACGTTTTACT
GATAGGTGGCTCGTTTAGTTTTGCACTGCCTTTGGGTGCGCGTGACCAAGTCATTTTCCAAATGTGTTTTAATATTTGTG
AACTATTATTAATAAAAAAGCGAGCGACGTTGTTCTTTGATGAGGGAACGACCGTACGCTTGGCGACCTTCTCTGATTGC
GTAGAGTCGTTTTCCTTATTGGGCTTACCTAGTACCGTCAGCCCACAATGTTTTCATTTCTTTCCCACCTAAAACTAAAA
GTACTCACAAGTCACAACTTAAACCAAGTACACAAGGATTTTATCATGGGATTATCGTGTTTGAAGACTAAAAAGAGCAC
ACCATCACCCCCATTAGTGCAGGTAGAGTAAGACAGTAACTTTTGGGTTCATATTACCGAGCAAGAACCGTTATTTGTGA
TTAGACATGTTATAAACCACTGCTTTAGTGACTATTTAAAACAATATATTACATGTCGTAATCATGCAACCTAACTATGT
TTTCATTAATCAAATACAAAGAATAAAGAGAAAAGTGCGTAGATTCAATTATTTGGCATAGACTCAAAAGAGTGTATATA
TATCTGACTTTTATTAAATTATTAAACACAAATACATATTTTCATAAGCAAAACTATAAAAGCCCTAAACATATAATGAT
TACCTCAAAGGAAAAAGTCGTTTTCTCCTAATTAAAAGATAGGTTACTTCCTAATTAATAGTATATAATTTATGTGAACT
TCACAATATACAGTtCAATAAAATTTGGTAATTTGACCGATTTAAGGAGAGTGGAAATTAGGGCTTCTGCAATCTTTTTT
CTTCGCCGCAATCTCATGTCCAATTATCCACCGACGGTGGCGGCGCAACCCACAACGACGGCGAATCCACTGCTGCAGCG
ACATCAATCTGAACAGCGACGAAGAGAATTACCGAAGATTGTCGAAACAGAGTCTACAAGTATGGACATTACGATCGGTC
AATCTAAGCAGCCTCAATTTTTGAAATCCATAGACGAATTAGCTGCGTTTTCAGTTGCAGTGGAAACATTCAAATGCCAA
TTCGATGATCTTCAGAAGCACATCGAGTCAATCGAAAACGCAATTGATTCCAAACTCGAGAGTAACGGCGTTGTCCTCGC
CGCGCGGAACAATAATTTCCATCAGCCGATGTTATCGCCTCCGCGGAACAATGTATCTGTAGAAACCACCGTCACTGTGA
GCCAACCGTCTCAGGAGATTGTACCGGAGACGTCGAATAAACCGGAGGGGGGACGTATGTGTGAGTTGATGTGTAGCAAA
GGTCTGCGTAAATACATATACGCGAATATCTCTGAACAAGCTAAGTTAATGGAAGAGATTCCTTCAGCTTTGAAATTGGC
CAAGGAGCCAGCGAAGTTTGTATTGGATTGTATTGGCAAGTTTTACTTACAAGGGCGTAGAGCATTTACTAAAGAGTCGC
CTATGAGCTCTGCGAGACAAGTTTCGCTTCTTATACTGGAGTCTTTTCTTCTAATGCCTGATCGTGGTAAAGGGAAGGTG
AAGATTGAGAGTTGGATTAAAGATGAGGCGGAGACGGCTGCTGTTGCTTGGAGGAAAAGGTTGATGACTGAAGGAGGATT
AGCTGCGGCTGAGAAAATGGATGCAAGGGGTTTGCTTTTACTAGTTGCTTGTTTTGGTGTTCCTTCAAACTTTAGGAGTA
CAGATTTGCTGGATTTGATAAGGATGAGTGGTTCGAATGAGATTGCCGGTGCTTTGAAGCGGTCACAGTTTCTTGTCCCT
ATGGTCTCAGGTACCATATTCTGTTCTCACTCGGTGAATTTCATTGCAAAGGTGGTTCCTTTTGTTGACATCATCGACCA
ACATCAAGTTCCATCTTTGTTTTTCGATAAGCTTGATGGTATAAACTAGGAGAGCACATCAAATATTTAGAGTGCAATGA
CTGATTGAGCCAAATCCTAGCTAGAAATTAATCTGGAAAGAACTTGGAACTCTCAACCATAGGTTTTGGTACGAAATTGT
TGCTTGTCAGAACCAAATGATAGGCTATTGCCTTGAAATAGTGTTTCTTGTGGTTTCCAATATTGGAAGTTAAAATCATA
TGACTTAGCTGTTGGATACTAATTAAGCTTAAGCAATGCCAACTCTAAGAAGTGGTACTTACACAATATTCTATTGGTCA
TAGGTATAGTTGAATCAAGTATCAAGCGTGGAATGCATATTGAAGCTCTTGAGATGGTTTATACCTTTGGCATGGAGGAT
AAGTTTTCAGCTGCTCTAGTTCTAACTTCATTCTTAAAGATGAGCAAGGAGTCATTTGAGAGGGCAAAACGGAAAGCCCA
GTCACCGCTGGCATTTGTATGAACCCTTCCCTTGCACATTATGTACCTTTATGAACTCTTTATCATCATCTGAGTCTGAC
CATTGATATATTTATTTCTCAACAGAAAGAAGCGGCTACAAAGCAGCTAGCTGTGTTATCATCAGTTATGCAGTGTATGG
AGACTCACAAGTTAGATCCTGCGAAAGAACTACCAGGATGGCAGATCAAAGAGCAAATTGTTAGCTTGGAGAAAGACACT
CTTCAGCTCGACAAAGAGATGGAAGAGAAAGCAAGATCTCTCAGTTTAATGGAGGAAGCCGCACTTGCCAAGAGAATGTA
TAACCAACAGATAAAACGTCCAAGGTTGTCACCCATGGAAATGCCACCAGTAACTTCTTCATCGTATTCTCCTATCTACC
GTGATAGAAGCTTTCCTAGTCAAAGAGACGATGACCAAGATGAAATTATCAGCTCTTGTGAGTAGTTACCTCGGCCCGTC
AACATCTTTTCCTCATCGCTCAAGAAGATCCCCGGAATATATGGTTCCACTTCCACATGGTGGGTTAGGAAGAAGTGTAT
ATGCATATGAACATCTGGCCCCAAATTCATACTCTCCAGGTCACGGACATAGACTTCATCGACAGTACTCTCCGTCTTTG
GTTCACGGACAGAGACATCCACTACAGTACTCTCCTCCAATTCATGGACAACAACAGTTACCATATGGTATACAAAGGGT
TTACAGACATTCACCATCTGAAGAAAGATATTTGGGTTtATCCAATCAAAGGTCTCCTCGCAGTAACTCATCATTAGACC
CCAAATAGGAGGAATGTAAATTTGTAACAAAGCTTTTTGTTTTTGCTTAAGTTAGTCATTTATTTAACTCCCAACAGTCT
CAAAATTTAATTTAATGTTTGGGGCTTAAGAATGCAAATTTTTTTGCTCCTGTAATTGACATTTAAGATGCTAATGTTAT
TGCTTCAGAGGTTTTAGTCAACCTCAGATACATCGATATCACTATCTAAATAGACCTCTGGCTCTTGGTCATCTGGATTC
TCTTCATCTTCTGTCTCTGTTCCTTCTTGTTCTCGTTGCACTGCTCGAGCAATTGCGGATTCCAACCTTGTGCTTACAGT
TTCCCATGACACAAGCTTTTCCATGAATGTATTTATGTCCGCCTTCTTATCTTTCTTGAGGAAGATGAATTC

>FRI_Bå1-2

GGATCCCAAAATCTAGTGCCGCCGAGTCCTTTGGGACAGAGTTTCGCACTTATCATTCGATTCGGCAAGCAGTAACTCGCA
AAATCCGCACCACCGTGAAACATACTCTGGCGCTGAACGCTAGACGCAAAATCCATATTACTGCTGCAGAAAGACGGAGA
GAACAAGAGACTGACCGATCATAAGAGAAGAGAGCTTCAAGGCTTCAGCTGATGAAGAATCAATGGCGGCGAGAGTGAGT
GAGAATCGAGTAGTTACTGAGAGGAAGATGAAACGTATCTCCTGAGAAAAAACGTAAGAGAGTTCCGTTTCAACAGTATT
TTTCTCGGCGGTCTTCAGATTTTCCGTCCAACTCTCTCTTTCCTCTTTCAGTTTTTTTTTGCCCAATCACTCGTGGTAGG
ATTTTTTTATAAGAAAGTTACGAAAATGCCCTCTCGTGCTTAGAGAGATTCAATTCGTGCACATATAATATGCCGTGAAT
ATACCACCAACTCTCTTACTCGTACTCGCATGTGGTGCGTGCAATATACACGTGGTTCGAATCAACGGCTACGTTTTACT
GATAGGTGGCTCGTTTAGTTTTGCACTGCCTTTGGGTGCGCGTGACCAAGTCATTTTCCAAATGTGTTTTAATATTTGTG
AACTATTATTAATAAAAAAGCGAGCGACGTTGTTCTTTGATGAGGGAACGACCGTACGCTTGGCGACCTTCTCTGATTGC
GTAGAGTCGTTTTCCTTATTGGGCTTACCTAGTACCGTCAGCCCACAATGTTTTCATTTCTTTCCCACCTAAAACTAAAA
GTACTCACAAGTCACAACTTAAACCAAGTACACAAGGATTTTATCATGGGATTATCGTGTTTGAAGACTAAAAAGAGCAC
ACCATCACCCCCATTAGTGCAGGTAGAGTAAGACAGTAACTTTTGGGTTCATATTACCGAGCAAGAACCGTTATTTGTGA
TTAGACATGTTATAAACCACTGCTTTAGTGACTATTTAAAACAATATATTACATGTCGTAATCATGCAACCTAACTATGT
TTTCATTAATCAAATACAAAGAATAAAGAGAAAAGTGCGTAGATTCAATTATTTGGCATAGACTCAAAAGAGTGTATATA
TATCTGACTTTTATTAAATTATTAAACACAAATACATATTTTCATAAGCAAAACTATAAAAGCCCTAAACATATAATGAT
TACCTCAAAGGAAAAAGTCGTTTTCTCCTACTTAAAAGATAGGTTACTTCCTAATTAATATATAATTTATGTGAACTTCA
CAATATACAGTTCAATAAAATTTGGTAATTTGACCGATTTAAGGAGAGTGGAAATTAGGGCTTCTGCAATCTTTTTTCTT
CGCCGCAATCTCATGTCCAATTATCCACCGACGGTGGCGGCGCAACCCACAACGACGGCGAATCCACTGCTGCAGCGACA
TCAATCTGAACAGCGACGAAGAGAATTACCGAAGATTGTCGAAACAGAGTCTACAAGTATGGACATTACGATCGGTCAAT
CTAAGCAGCCTCAATTTTTGAAATCCATAGACGAATTAGCTGCGTTTTCAGTTGCAGTGGAAACATTCAAACGCCAATTC
GATGATCTTCAGAAGCACATCGAGTCAATCGAAAACGCAATTGATTCCAAACTCGAGAGTAACGGCGTTGTCCTCGCCGC
GCGGAACAATAATTTCCATCAGCCGATGTTATCGCCTCCGCGGAACAATGTATCTGTAGAAACCACCGTCACTGTGAGCC
AACCGTCTCAGGAGATTGTACCGGAGACGTCGAATAAACCGGAGGGGGGACGTATGTGTGAGTTGATGTGTAGCAAAGGT
CTGCGTAAATACATATACGCGAATATCTCTGATCAAGCTAAGTTAATGGAAGAGATTCCTTCAGCTTTGAAATTGGCCAA
GGAGCCAGCGAAGTTTGTATTGGATTGTATTGGCAAGTTTTACTTACAAGGGCGTAGAGCATTTACTAAAGAGTCGCCTA
TGAGCTCTGCGAGACAAGTTTCGCTTCTTATACTGGAGTCTTTTCTTCTAATGCCTGATCGTGGTAAAGGGAAGGTGAAG
ATTGAGAGTTGGATTAAAGATGAGGCGGAGACGGCTGCTGTTGCTTGGAGGAAAAGGTTGATGACTGAAGGAGGATTAGC
TGCGGCTGAGAAAATGGATGCAAGGGGTTTGCTTTTACGAGTTGCTTGTTTTGGTGTTCCTTCAAACTTTAGGAGTACAG
ATTTGCTGGATTTGATAAGGATGAGTGGTTCGAATGGGATTGCCGGTGCTTTGAAGCGGTCACAGTTTCTTGTCCCTATG
GTCTCAGGTACCATATTCTGTTCTCACTCGGTGAATTTCATTGCAAAGGTGGTTCCTTTTGTTGACATCATCGACCAACA
TCAAGTTCCATCTTTGTTTTTCGATAAGCTTGATGGTATAAACTAGGAGAGCACATCAAATATTTAGAGTGCAATGACTG
ATTGAGCCAAATCCTAGCTAGAAATTAATCTGGAAAGAACTTGGAACTCTCAACCATAGGTTTTGGTACGAAATTGTTGC
TTGTCAGAACCAAATGATAGGCTATTGCCTTGAAATAGTGTTTCTTGTGGTTTCCAATATTGGAAGTTAAAATCGTATGA
CTTAGCTGTTGGATACTAATTAAGCTTAAGCAATGCCAACTCTAAGAAGTGGTACTTACACAATATTCTATTGGTCATAG
GTATAGTTGAATCAAGTATCAAGCGTGGAATGCATATTGAAGCTCTTGAGATGGTTTATACCTTTGGCATGGAGGATAAG
TTTTCAGCTGCTCTAGTTCTAACTTCATTCTTAAAGATGAGCAAGGAGTCATTTGAGAGGGCAAAACGGAAAGCCCAGTC
ACCGCTGGCATTTGTATGAACCCTTCCCTTGCACATTATGTACCTTTATGAACTCTTTATCATCATCTGAGTCTGACCAT
TGATATATTTATTTCTCAACAGAAAGAAGCGGCTACAAAGCAGCTAGCTGTGTTATCATCAGTTATGCAGTGTATGGAGA
CTCACAAGTTAGATCCTGCGAAAGAACTACCAGGATGGCAGATCAAAGAGCAAATTGTTAGCTTGGAGAAAGACACTCTT
CAGCTCGACAAAGAGATGGAAGAGAAAGCAAGATCTCTCAGTTTAATGGAGGAAGCCGCACTTGCCAAGAGAATGTATAA
CCAACAGATAAAACGTCCAAGGTTGTCACCCATGGAAATGCCACCAGTAACTTCTTCATCGTATTCTCCTATCTACCGTG
ATAGAAGCTTTCCTAGTCAAAGAGACGATGACCAAGATGAAATATCAGCTCTTGTGAGTAGTTACCTCGGCCCGTCAACA
TCTTTTCCTCATCGCTCAAGAAGATCCCCGGAATATATGGTTCCACTTCCACATGGTGGGTTAGGAAGAAGTGTATATGC
ATATGAACATCTGGCCCCAAATTCATACTCTCCAGGTCACGGACATAGACTTCATCGACAGTACTCTCCGTCTTTGGTTC
ACGGACAGAGACATCCACTACAGTACTCTCCTCCAATTCATGGACAACAACAGTTACCATATGGTATACAAAGGGTTTAC
AGACATTCACCATCTGAAGAAAGATATTTGGGTTTATCCAATCAAAGGTCTCCTCGCAGTAACTCATCATTAGACCCCAA
ATAGGAGGAATGTAAATTTGTAACAAAGCTTTTTGTTTTTGCTTAAGTTAGTCATTTATTTAACTCCCAACAGTCTCAAA
ATTTAATTTAATGTTTGGGGCTTAAGAATGCAAATTTTTTTGCTCCTGTAATTGACATTTAAGATGCTAATGTTATTGCT
TCAGAGGTTTTAGTCAACCTCAGATACATCGATATCACTATCTAAAATAGACCTCTGGCTCTTGGGTCATCTGGGATTCT
CTTCATCCTCTGGTCTCGGTTCCTTCTTGGTTCTCGTTGGCACTGCCTCGAAGCAATTGGCGGGA

>FRI_Wa-1
GGATCCCAAAATCTAGTGCCGCCGAGTCCTTTGGGACACAGTTTCGCACTTATCATTCGATTCGGCAAGCAGTAACTCGC

AAAATCCGCACCACCGTGAAACATACTCCGGCGCTGAACGCTAGACGCAAAATCCATATTACAGCTGCAGAAAGACGGAG
AGAACAAGAGACTGACCGATCATAAGAGAAGAGAGCTTCAAGGCTTCAGCTGATGAAGAATCAATGGCGGCGAGAGTGAG
TGAGAATCGAGTAGTTACTGAGAGGAAGATGAAACGTATCTCCTGAGAAAAAACGTAAGAGAGTTCCGTTTCAACAGTAT
TTTTCTCGGCGGTCTTCAGATTTTCCGTCCAACTCTCTCTTTCCTCTTTCAGTTTTTTTTTGCCCAATCACTCGTGGTAG
GATTTTTTTATAAGAAAGTTACGAAAATGCCCTCTCGTGCTTAGAGAGATTCAATTCGTGCACATATAATATGCCGTGAA
TATACCACCAACTCTCTTACTCGTACTCGCATGTGGTGCGTGCAATATACACGTGGTTCGAATCAACGGCTACGTTTTAC
TGATAGGTGGCTCGTTTAGTTTTGCACTGCCTTTGGGTGCGCGTGACCAAGTCATTTTCCAAATGTGTTTTAATATTTGT
GAACTATTATTAATAAAAAAGCGAGCGACGTTGTTCTTTGATGAGGGAACGACCGTACGCTTGGCGACCTTCTCTGATTG
CGTAGAGTCGTTTTCCTTATTGGGCTTACCTAGTACCGTCAGCCCACAATGTTTTCATTTCTTTCCCACCTAAAACTAAA
AGTACTCACAAGTCACAACTTAAACCAAGTACACAAGGATTTTATCATGGGATTATCGAGTTTGAAGACTAAAAAGAGCA
CACCATCACCCCCATTAGTGCAGGTAGAGTAAGACAGTAACTTTTGGGTTCATATTACCGAGCAAGAACCGTTATTTGTG
ATTAGACATGTTATAAACCACTGCTTTAGTGACTATTTAAAACAATATATTACATGTCGTAATCATGCAACCTAACTATG
TTTTCATTAATCAAATACAAAGAATAAAGAGAAAAGTGCGTAGATTCAATTATTTGGCATAGACTCAAAAGAGTGTATAT
ATATCTGACTTTTATTAAATTATTAAACACAAATACATATTTTCATAAGCAAAACTATAAAAGCCCTAAAGATATAACGA
TTACCTCAAAGGAAAAAGTCGTTTTCTCCTAATTAAAAGATAGGTTACTTCCTAATTAATATATAATTTATGTGAACTTC
ACAATATACAGTTCAATAAAATTTGGTAATTTGACCGATTTAAGGAGAGTGGAAATTTGGGCTTCTGCAATCTTTTTTCT
TCGCCGCAATCTCATGTCCAATTATCCACCGACGGTGGCGGCGCAACCCACAACGACGGCGAATCCACTGCTGCAGCGAC
ATCAATCTGAACAGCGACGAAGAGAATTACCGAAGATTGTCGAAACAGAGTCTACAAGTATGGACATTACGATCGGTCAA
TCTAAGCAGCCTCAATTTTTGAAATCCATAGACGAATTAGCTGCGTTTTCAGTTGCAGTGGAAACATTCAAATGCCAATT
CGATGATCTTCAGAAGCACATCGAGTCAATCGAAAACGCAATTGATTCCAAACTCGAGAGTAACGGCGTTGTCCTCGCCG
CGCGGAACAATAATTTCCATCAGCCGATGTTATCGCCTCCGCGGAACAATGTATCTGTAGAAACCACCGTCACTGTGAGC
CAACCGTCTCAGGAGATTGTACCGGAGACGTCGAATAAACCGGAGGGGGGACGTATGTGTGAGTTGATGTGTAGCAAAGG
TCTGCGTAAATACATATACGCGAATATCTCTGAACAAGCTAAGTTAATGGAAGAGATTCCTTCAGCTTTGAAATTGGCCA
AGGAGCCAGCGAAGTTTGTATTGGATTGTATTGGCAAGTTTTACTTACAAGGGCGTAGAGCATTTACTAAAGAGTCGCCT
ATGAGCTCTGCGAGACAAGTTTCGCTTCTTATACTGGAGTCTTTTCTTCTAATGCCTGATCGTGGTAAAGGGAAGGTGAA
GATTGAGAGTTGGATTAAAGATGAGGCGGAGACGGCTGCTGTTGCTTGGAGGAAAAGGTTGATGACTGAAGGAGGATTAG
CTGCGGCTGAGAAAATGGATGCAAGGGGTTTGCTTTTACTAGTTGCTTGTTTTGGTGTTCCTTCAAACTTTAGGAGTACA
GATTTGCTGGATTTTATAAGGATGAGTGGTTCGAATGAGATTGCCGGTGCTTTGAAGCGGTCACAGTTTCTTGTCCCTAT
GGTCTCAGGTACCATATTCTGTTCTCACTCGGTGAATTTCATTGCAAAGGTGGTTCCTTTTGTTGACATCATCGACCAAC
ATCAAGTTCCATCTTTGTTTTTCGATAAGCTTGATGGTATAAACTAGGAGAGCACATCAAATATTTAGAGTGCAATGACT
GATTGAGCCAAATCCTAGCTAGAAATTAATCTGGAAAGAACTTGGAACTCTCAACCATAGGTTTTGGTACGAAATTGTTG
CTTGTCAGAACCAAATGATAGGCTATTGCCTTGAAATAGTGTTTCTTGTGGTTTCCAATATTGGAAGTTAAAATCATATG
ACTTAGCTGTTGGATACTAATTAAGCTTAAGCAATGCCAACTCTAAGAAGTGGTACTTACACAATATTCTATTGGTCATA
GGTATAGTTGAATCAAGTATCAAGCGTGGAATGCATATTGAAGCTCTTGAGATGGTTTATACCTTTGGCATGGAGGATAA
GTTTTCAGCTGCTCTAGTTCTAACTTCATTCTTAAAGATGAGCAAGGAGTCATTTGAGAGGGCAAAACGGAAAGCCCAGT
CACCGCTGGCATTTGTATGAACCCTTCCCTTGCACATTATGTACCTTTATGAACTCTTTATCATCATCTGAGTCTGACCA
TTGATATATTTATTTCTCAACAGAAAGAAGCGGCTACAAAGCAGCTAGCTGTGTTATCATCAGTTATGCAGTGTATGGAG
ACTCACAAGTTAGATCCTGCGAAAGAACTACCAGGATGGCAGATCAAAGAGCAAATTGTTAGCTTGGAGAAAGACACTCT
TCAGCTCGACAAAGAGATGGAAGAGAAAGCAAGATCTCTCAGTTTAATGGAGGAAGCCGCACTTGCCAAGAGAATGTATA
ACCAACAGATAAAACGTCCAAGGTTGTCACCCATGGAAATGCCACCAGTAACTTCTTCATCGTATTCTCCTATCTACCGT
GATAGAAGCTTTCCTAGTCAAAGAGACGATGACCAAGATGAAATATCAGCTCTTGTGAGTAGTTACCTCGGCCCGTCAAC
ATCTTTTCCTCATCGCTCAAGAAGATCCCCGGAATATATGGTTCCACTTCCACATGGTGGGTTAGGAAGAAGTGTATATG
CATATGAACATCTGGCCCCAAATTCATACTCTCCAGGTCACGGACATAGACTTCATCGACAGTACTCTCCGTCTTTGGTT
CACGGACAGAGACATCCACTACAGTACTCTCCTCCAATTCATGGACAACAACAGTTACCATATGGTATACAAAGGGTTTA
CAGACATTCACCATCTGAAGAAAGATATTTGGGTTTATCCAATCAAAGGTCTCCTCGCAGTAACTCATCATTAGACCCCA
AATAGGAGGAATGTAAATTTGTAACAAAGCTTTTTGTTTTTGCTTAAGTTAGTCATTTATTTAACTCCCAACAGTCTCAA
AATTTAATTTAATGTTTGGGGCTTAAGAATGCAAATTTTTTTGCTCCTGTAATTGACATTTAAGATGCTAATGTTATTGC
TTCAGAGGTTTTAGTCAACCTCAGATACATCGATATCACTATCTAAATAGACCTCTGGCTCTTGGTCATCTGGATTCTCT
TCATCTTCTGTCTCTGTTCCTTCTTGTTCTCGTTGCACTGCTCGAGCAATTGCGGATTCCAACCTTGTGCTTACAGTTTC
CCATGACACAAGCTTTTCCATGAATGTATTTATGTCCGCCTTCTTATCTTTCTTGAGGAAGATGAATTC

>FRI_Zdr-6
GGATCCCAAAATCTAGTGCCGCCGAGTCCTTTGGGACACAGTTTCGCACTTATCATTCGATTCGGCAAGCAGTAACTCGC

AAAATCCGCACCACCGTGAAACATACTCCGGCGCTGAACGCTAGACGCAAAATCCATATTACTGCTGCAGAAAGACGGAG
AGAACAAGAGACTGACCGATCATAAGAGAAGAGAGCTTCAAGGCTTCAGCTGATGAAGAATCAATGGCGGCGAGAGAGAG
TGAGAATCGAGTAGTTACTGAGAGGAAGATGAAACGTATCTCCTGAGAAAAAACGTAAGAGAGTTCCGTTTCAACAGTAT
TTTTCTCGGCGGTCTTCAGATTTTCCGTCCAACTCTCTCTTTCCTCTTTCAGTTTTTTTTTGCCCAATCACTCGTGGTAG
GATTTTTTTATAAGAAAGTTACGAAAATGCCCTCTCGTGCTTAGAGAGATTCAATTCGTGCACATATAATATGCCGTGAA
TATACCACCAACTCTCTTACTCGTACTCGCATGTGGTGCGTGCAATATACACGTGGTTCGAATCAACGGCTACGTTTTAC
TGATAGGTGGCTCGTTTAGTTTTGCACTGCCTTTGGGTGCGCGTGACCAAGTCATTTTCCAAATGTGTTTTAATATTTGT
GAACTATTATTAATAAAAAAGCGAGCGACGTTGTTCTTTGATGAGGGAACGACCGTACGCTTGGCGACCTTCTCTGATTG
CGTAGAGTCGTTTTCCTTACTGGGCTTACCTAGTACCGTCAGCCCACAATGTTTTCATTTCTTTCCCACCTAAAACTAAA
AGTACTCACAAGTCACAACTTAAACCAAGTACACAAGGATTTTATCATGGGATTATCGTGTTTGAAGACTAAAAAGAGCA
CACCATCACCCCCATTAGTGCAGGTAGAGTAAGACAGTAACTTTTGGGTTCATATTACCGAGCAAGAACCGTTATTTGTG
ATTAGACATGTTATAAACCACTGCTTTAGTGACTATTTAAAACAATATATTACATGTCGTAATCATGCAACCTAACTATG
TTTTCATTAATCAAATACAAAGAATAAAGAGAAAAGTGCGTAGATTCAATTATTTGGCATAGACTCAAAAGAGTGTATAT
ATATCTGACTTTTATTAAATTATTAAACACAAATACATATTTTCATAAGCAAAACTATAAAAGCCCTAAACATATAATGA
TTACCTCAAAGGAAAAAGTCGTTTTCTCCTAATTAAAAGATAGGTTACTTCCTAATTAATAGTATATAATTTATGTGAAC
TTCACAATATACAGTTCAATAAAATTTGGTAATTTGACCGATTTAAGGAGAGTGGAAATTAGGGCTTCTGCAATCTTTTT
TCTTCGCCGCAATCTCATGTCCAATTATCCACCGACGGTGGCGGCGCAACCCACAACGACGGCGAATCCACTGCTGCAGC
GACATCAATCTGAACAGCGACGAAGAGAATTACCGAAGATTGTCGAAACAGAGTCTACAAGTATGGACATTACGATCGGT
CAATCTAAGCAGCCTCAATTTTTGAAATCCATAGACGAATTAGCTGCGTTTTCAGTTGCAGTGGAAACATTCAAATGCCA
ATTCGATGATCTTCAGAAGCACATCGAGTCAATCGAAAACGCAATTGATTCCAAACTCGAGAGTAACGGCGTTGTCCTCG
CCGCGCGGAACAATAATTTCCATCAGCCGATGTTATCGCCTCCGCGGAACAATGTATCTGTAGAAACCACCGTCACTGTG
AGCCAACCGTCTCAGGAGATTGTACCGGAGACGTCGAATAAACCGGAGGGGGGACGTATGTGTGAGTTGATGTGTAGCAA
AGGTCTGCGTAAATACATATACGCGAATATCTCTGAACAAGCTAAGTTAATGGAAGAGATTCCTTCAGCTTTGAAATTGG
CCAAGGAGCCAGCGAAGTTTGTATTGGATTGTATTGGCAAGTTTTACTTACAAGGGCGTAGAGCATTTACTAAAGAGTCG
CCTATGAGCTCTGCGAGACAAGTTTCGCTTCTTATACTGGAGTCTTTTCTTCTAATGCCTGATCGTGGTAAAGGGAAGGT
GAAGATTGAGAGTTGGATTAAAGATGAGGCGGAGACGGCTGCTGTTGCTTGGAGGAAAAGGTTGATGACTGAAGGAGGAT
TAGCTGCGGCTGAGAAAATGGATGCAAGGGGTTTGCTTTTACTAGTTGCTTGTTTTGGTGTTCCTTCAAACTTTAGGAGT
ACAGATTTGCTGGATTTGATAAGGATGAGTGGTTCGAATGAGATTGCCGGTGCTTTGAAGCGGTCACAGTTTCTTGTCCC
TATGGTCTCAGGTACCATATTCTGTtCTCACTCGGTGAATTTCATTGCAAAGGTGGTTCCTTTTGTTGACATCATCGACC
AACATCAAGTTCCATCTTTGTTTTTCGATAAGCTTGATGGTATAAACTAGGAGAGCACATCAAATATTTAGAGTGCAATG
ACTGATTGAGCCAAATCCTAGCTAGAAATTAATCTGGAAAGAACTTGGAACTCTCAACCATAGGTTTTGGTACGAAATTG
TTGCTTGTCAGAACCAAATGATAGGCTATTGCCTTGAAATAGTGTTTCTTGTGGTTTCCAATATTGGAAGTTAAAATCAT
ATGACTTAGCTGTTGGATACTAATTAAGCTTAAGCAATGCCAACTCTAAGAAGTGGTACTTACACAATATTCTATTGGTC
ATAGGTATAGTTGAATCAAGTATCAAGCGTGGAATGCATATTGAAGCTCTTGAGATGGTTTATACCTTTGGCATCGAGGA
TAAGTTTTCAGCTGCTCTAGTTCTAACTTCATTCTTAAAGATGAGCAAGGAGTCATTTGAGAGGGCAAAACGGAAAGCCC
AGTCACCGCTGGCATTTGTATGAACCCTTCCCTTGCACATTATGTACCTTTATGAACTCTTTATCATCATCTGAGTCTGA
CCATTGATATATTTATTTCTCAACAGAAAGAAGCGGCTACAAAGCAGCTAGCTGTGTTATCATCAGTTATGCAGTGTATG
GAGACTCACAAGTTAGATCCTGCGAAAGAACTACCAGGATGGCAGATCAAAGAGCAAATTGTTAGCTTGGAGAAAGACAC
TCTTCAGCTCGACAAAGAGATGGAAGAGAAAGCAAGATCTCTCAGTTTAATGGAGGAAGCCGCACTTGCCAAGAGAATGT
ATAACCAACAGATAAAACGTCCAAGGTTGTCACCCATGGAAATGCCACCAGTAACTTCTTCATCGTATTCTCCTATCTAC
CGTGATAGAAGCTTTCCTAGTCAAAGAGACGATGACCAAGATGAAATATCAGCTCTTGTGAGTAGTTACCTCGGCCCGTC
AACATCTTTTCCTCATCGCTCAAGAAGATCCCCGGAATATATGGTTCCACTTCCACATGGTGGGTTAGGAAGAAGTGTAT
ATGCATATGAACATCTGGCCCCAAATTCATACTCTCCAGGTCACGGACATAGACTTCATCGACAGTACTCTCCGTCTTTG
GTTCACGGACAGAGACATCCACTACAGTACTCTCCTCCAATTCATGGACAACAACAGTTACCATATGGTATACAAAGGGT
TTACAGACATTCACCATCTGAAGAAAGATATTTGGGTTTATCCAATCAAAGGTCTCCTCGCAGTAACTCATCATTAGACC
CCAAATAGGAGGAATGTAAATTTGTAACAAAGCTTTTTGTTTTTGCTTAAGTTAGTCATTTATTTAACTCCCAACAGTCT
CAAAATTTAATTTAATGTTTGGGGCTTAAGAATGCAAATTTTTTTGCTCCTGTAATTGACATTTAAGATGCTAATGTTAT
TGCTTCAGAGGTTTTAGTCAACCTCAGATACATCGATATCACTATCTAAATAGACCTCTGGCTCTTGGTCATCTGGATTC
TCTTCATCTTCTGTCTCTGTTCCTTCTTGTTCTCGTTGCACTGCTCGAGCAATTGCGGATTCCAACCTTGTGCTTACAGT
TTCCCATGACACAAGCTTTTCCATGAATGTATTTATGTCCGCCTTCTTATCTTTCTTGAGGAAGATGAATTC

>FRI_Spr-1-6
GGATCCCAAAATCTAGTGCCGCCGAGTCCTTTGGGACACAGTTTCGCACTTATCATTCGATTCGGCAAGCAGTAACTCGC

AAAATCCGCACCACCGTGAAACATACTCCGGCGCTGAACGCTAGACGCAAAATCCATATTACAGCTGCAGAAAGACGGAG
AGAACAAGAGACTGACCGATCATAAGAGAAGAGAGCTTCAAGGCTTCAGCTGATGAAGAATCAATGGCGGCGAGAGTGAG
TGAGAATCGAGTAGTTACTGAGAGGAAGATGAAACGTATCTCCTGAGAAAAAACGTAAGAGAGTTCCGTTTCAACAGTAT
TTTTCTCGGCGGTCTTCAGATTTTCCGTCCAACTCTCTCTTTCCTCTTTCAGTTTTTTTTTGCCCAATCACTCGTGGTAG
GATTTTTTTATAAGAAAGTTACGAAAATGCCCTCTCGTGCTTAGAGAGATTCAATTCGTGCACATATAATATGCCGTGAA
TATACCACCAACTCTCTTACTCGTACTCGCATGTGGTGCGTGCAATATACACGTGGTTCGAATCAACGGCTACGTTTTAC
TGATAGGTGGCTCGTTTAGTTTTGCACTGCCTTTGGGTGCGCGTGACCAAGTCATTTTCCAAATGTGTTTTAATATTTGT
GAACTATTATTAATAAAAAAGCGAGCGACGTTGTTCTTTGATGAGGGAACGACCGTACGCTTGGCGACCTTCTCTGATTG
CGTAGAGTCGTTTTCCTTATTGGGCTTACCTAGTACCGTCAGCCCACAATGTTTTCATTTCTTTCCCACCTAAAACTAAA
AGTACTCACAAGTCACAACTTAAACCAAGTACACAAGGATTTTATCATGGGATTATCGAGTTTGAAGACTAAAAAGAGCA
CACCATCACCCCCATTAGTGCAGGTAGAGTAAGACAGTAACTTTTGGGTTCATATTACCGAGCAAGAACCGTTATTTGTG
ATTAGACATGTTATAAACCACTGCTTTAGTGACTATTTAAAACAATATATTACATGTCGTAATCATGCAACCTAACTATG
TTTTCATTAATCAAATACAAAGAATAAAGAGAAAAGTGCGTAGATTCAATTATTTGGCATAGACTCAAAAGAGTGTATAT
ATATCTGACTTTTATTAAATTATTAAACACAAATACATATTTTCATAAGCAAAACTATAAAAGCCCTAAAGATATAACGA
TTACCTCAAAGGAAAAAGTCGTTTTCTCCTAATTAAAAGATAGGTTACTTCCTAATTAATATATAATTTATGTGAACTTC
ACAATATACAGTTCAATAAAATTTGGTAATTTGACCGATTTAAGGAGAGTGGAAATTTGGGCTTCTGCAATCTTTTTTCT
TCGCCGCAATCTCATGTCCAATTATCCACCGACGGTGGCGGCGCAACCCACAACGACGGCGAATCCACTGCTGCAGCGAC
ATCAATCTGAACAGCGACGAAGAGAATTACCGAAGATTGTCGAAACAGAGTCTACAAGTATGGACATTACGATCGGTCAA
TCTAAGCAGCCTCAATTTTTGAAATCCATAGACGAATTAGCTGCGTTTTCAGTTGCAGTGGAAACATTCAAATGCCAATT
CGATGATCTTCAGAAGCACATCGAGTCAATCGAAAACGCAATTGATTCCAAACTCGAGAGTAACGGCGTTGTCCTCGCCG
CGCGGAACAATAATTTCCATCAGCCGATGTTATCGCCTCCGCGGAACAATGTATCTGTAGAAACCACCGTCACTGTGAGC
CAACCGTCTCAGGAGATTGTACCGGAGACGTCGAATAAACCGGAGGGGGGACGTATGTGTGAGTTGATGTGTAGCAAAGG
TCTGCGTAAATACATATACGCGAATATCTCTGAACAAGCTAAGTTAATGGAAGAGATTCCTTCAGCTTTGAAATTGGCCA
AGGAGCCAGCGAAGTTTGTATTGGATTGTATTGGCAAGTTTTACTTACAAGGGCGTAGAGCATTTACTAAAGAGTCGCCT
ATGAGCTCTGCGAGACAAGTTTCGCTTCTTATACTGGAGTCTTTTCTTCTAATGCCTGATCGTGGTAAAGGGAAGGTGAA
GATTGAGAGTTGGATTAAAGATGAGGCGGAGACGGCTGCTGTTGCTTGGAGGAAAAGGTTGATGACTGAAGGAGGATTAG
CTGCGGCTGAGAAAATGGATGCAAGGGGTTTGCTTTTACTAGTTGCTTGTTTTGGTGTTCCTTCAAACTTTAGGAGTACA
GATTTGCTGGATTTGATAAGGATGAGTGGTTCGAATGAGATTGCCGGTGCTTTGAAGCGGTCACAGTTTCTTGTCCCTAT
GGTCTCAGGTACCATATTCTGTTCTCACTCGGTGAATTTCATTGCAAAGGTGGTTCCTTTTGTTGACATCATCGACCAAC
ATCAAGTTCCATCTTTGTTTTTCGATAAGCTTGATGGTATAAACTAGGAGAGCACATCAAATATTTAGAGTGCAATGACT
GATTGAGCCAAATCCTAGCTAGAAATTAATCTGGAAAGAACTTGGAACTCTCAACCATAGGTTTTGGTACGAAATTGTTG
CTTGTCAGAACCAAATGATAGGCTATTGCCTTGAAATAGTGTTTCTTGTGGTTTCCAATATTGGAAGTTAAAATCATATG
ACTTAGCTGTTGGATACTAATTAAGCTTAAGCAATGCCAACTCTAAGAAGTGGTACTTACACAATATTCTATTGGTCATA
GGTATAGTTGAATCAAGTATCAAGCGTGGAATGCATATTGAAGCTCTTGAGATGGTTTATACCTTTGGCATGGAGGATAA
GTTTTCAGCTGCTCTAGTTCTAACTTCATTCTTAAAGATGAGCAAGGAGTCATTTGAGAGGGCAAAACGGAAAGCCCAGT
CACCGCTGGCATTTGTATGAACCCTTCCCTTGCACATTATGTACCTTTATGAACTCTTTATCATCATCTGAGTCTGACCA
TTGATATATTTATTTCTCAACAGAAAGAAGCGGCTACAAAGCAGCTAGCTGTGTTATCATCAGTTATGCAGTGTATGGAG
ACTCACAAGTTAGATCCTGCGAAAGAACTACCAGGATGGCAGATCAAAGAGCAAATTGTTAGCTTGGAGAAAGACACTCT
TCAGCTCGACAAAGAGATGGAAGAGAAAGCAAGATCTCTCAGTTTAATGGAGGAAGCCGCACTTGCCAAGAGAATGTATA
ACCAACAGATAAAACGTCCAAGGTTGTCACCCATGGAAATGCCACCAGTAACTTCTTCATCGTATTCTCCTATCTACCGT
GATAGAAGCTTTCCTAGTCAAAGAGACGATGACCAAGATGAAATATCAGCTCTTGTGAGTAGTTACCTCGGCCCGTCAAC
ATCTTTTCCTCATCGCTCAAGAAGATCCCCGGAATATATGGTTCCACTTCCACATGGTGGGTTAGGAAGAAGTGTATATG
CATATGAACATCTGGCCCCAAATTCATACTCTCCAGGTCACGGACATAGACTTCATCGACAGTACTCTCCGTCTTTGGTT
CACGGACAGAGACATCCACTACAGTACTCTCCTCCAATTCATGGACAACAACAGTTACCATATGGTATACAAAGGGTTTA
CAGACATTCACCATCTGAAGAAAGATATTTGGGTTTATCCAATCAAAGGTCTCCTCGCAGTAACTCATCATTAGACCCCA
AATAGGAGGAATGTAAATTTGTAACAAAGCTTTTTGTTTTTGCTTAAGTTAGTCATTTATTTAACTCCCAACAGTCTCAA
AATTTAATTTAATGTTTGGGGCTTAAGAATGCAAATTTTTTTGCTCCTGTAATTGACATTTAAGATGCTAATGTTATTGC
TTCAGAGGTTTTAGTCAACCTCAGATACATCGATATCACTATCTAAATAGACCTCTGGCTCTTGGTCATCTGGATTCTCT
TCATCTTCTGTCTCTGTTCCTTCTTGTTCTCGTTGCACTGCTCGAGCAATTGCGGATTCCAACCTTGTGCTTACAGTTTC
CCATGACACAAGCTTTTCCATGAATGTATTTATGTCCGCCTTCTTATCTTTCTTGAGGAAGATGAATTC

>FRI_Lip-0
GGATCCCAAAATCTAGTGCCGCCGAGTCCTTTGGGACACAGTTTCGCACTTATCATTCGATTCGGCAAGCAGTAACTCGC

AAAATCCGCACCACCGTGAAACATACTCCGGCGCTGAACGCTAGACGCAAAATCCATATTACAGCTGCAGAAAGACGGAG
AGAACAAGAGACTGACCGATCATAAGAGAAGAGAGCTTCAAGGCTTCAGCTGATGAAGAATCAATGGCGGCGAGAGAGAG
TGAGAATCGAGTAGTTACTGAGAGGAAGATGAAACGTATCTCCTGAGAAAAAACGTAAGAGAGTTCCGTTTCAACAGTAT
TTTTCTCGGCGGTCTTCAGATTTTCCGTCCAACTCTCTCTTTCCTCTTTCAGTTTTTTTTGCCCAATCACTCGTGGTAGG
ATTTTTTATAAGAAAGTTACGAAAATGCCCTCTCGTGCTTAGAGAGATTCAATTCGTGCACATATAATATGCCGTGAATA
TACCACCAACTCTCTTACTCGTACTCGCATGTGGTGCGTGCAATATACACGTGATTCGAATCAACGGCTACGTTTTACTG
ATAGGTGGCTCGTTTAGTTTTGCACTGCCTTTGGGTGCGCGTGACCAAGTCATTTTCCAAATGTGTTTTAATATTTGTGA
ACTATTATTAATAAAAAAGCGAGCGACGTTGTTCTTTGATGAGGGAACGACCGTACGCTTGGCGACCTTCTCTGATTGCG
TAGAGTCGTTTTcCTTATTGGGCTTACCTAGTACCGCCTGCCCACAATGTTTTCATTTCTTTCCCACCTAAAACTAAAAG
TACTCACAAGTCACAACTTAAACCAAGTACACAAGGATTTTATCATGGGATTATCGTGTTTGAAGACTAAAAAGAGCACA
CCATCACCCCCATTAGTGCAGGTAGAGTAAGACAGTAACTTTTGGGTTCATATTACCGAGCAAGAACCGTTATTTGTGAT
TAGACATGTTATAAACCACTGCTTTAGTGACTATTTAAAACAATATATTACATGTCGTAATCATGCAACCTAACTATGTT
TTCATTAATCAAATACAAAGAATAAAGAGAAAAGTGCGTAGATTCAATTATTTGGCATAGACTCAAAAGAGTGTATATAT
ATCTGACTTTTATTAAATTATTAAACACAAATACATATTTCATAAGCAAAACTATAAAAGCCCTAAACATATAATGATT
ACCTCAAAGGAAAAAGTCGTTTTCTCCTAATTAAAAGATAGGTTACTTCCTAATTAATAGTATATAATTTATGTGAACTT
CACAATATACAGTTCAATAAAATTTGGTAATTTGACCGATTTAAGGAGAGTGGAAATTAGGGCTTCTGCAATCTTTTTTC
TTCGCCGCAATCTCATGTCCAATTATCCACCGACGGTGGCGGCGCAACCCACAACGACGGCAAATCCACTGCTGCAGCGA
CATCAATCTGAACAGCGACGAAGAGAATTACCGAAGATTGTCGAAACAGAGTCTACAAGTATGGACATTACGATCGGTCA
ATCTAAGCCGCCTCAATTTTTGAAATCCATAGACGAATTAGCTGCGTTTTCAGTTGCAGTGGAAACATTCAAACGCCAAT
TCGATGATCTTCAGAAGCACATCGAGTCAATCGAAAACGCAATTGATTCCAAACTCGAGAGTAACGGCGTTGTCCTCGCC
GCGCGgAACAATAATTTCCATCAGCCGATGTTATCGCCTCCGCGGAACAATGTATCTGTAGAAACCACCGTCACTGTGAG
CCAACCGTCTCAGGAGATTGTACCGGAGACGTCGAATAAACCGGAGGGGGGACGTATGTGTGAGTTGATGTGTAGCAAAG
GTCTGCGTAAATACATATACGCGAATATCTCTGAACAAGCTAAGTTAATGGAAGAGATTCCTTCAGCTTTGAAATTGGCC
AAGGAGCCAGCGAAGTTTGTATTGGATTGTATTGGCAAGTTTTACTTACAAGGGCGTAGAGCATTTACTAAAGAGTCGCC
TATGAGCTCTGCGAGACAAGTTTCGCTTCTTATACTGGAGTCTTTTCTTCTAATGCCTGATCGTGGTAAAGGGAAGGTGA
AGATTGAGAGTTGGATTAAAGATGAGGCGGAGACGGCTGCTGTTGCTTGGAGGAAAAGGTTGATGACTGAAGGAGGATTA
GCTGCGGCTGAGAAAATGGATGCAAGGGGTTTGCTTTTACTAGTTGCTTGTTTTGGTGTTCCTTCAAACTTTAGGAGTAC
AGATTTGCTGGATTTGATAAGGATGAGTGGTTCGAATGAGATTGCCGGTGCTTTGAAGCGGTCACAGTTTCTTGTCCCTA
TGGTCTCAGGTACCATATTCTGTTCTCACTCGGTGAATTTCATTGCAAAGGTGGTTCCTTTTGTTGACATCATCGACCAA
CATCAAGTTCCATCTTTGTTTTTCGATAAGCTTGATGGTATAAACTAGGAGAGCACATCAAATATTTAGAGTGCAATGAC
TGATTGAGCCAAATCCTAGCTAGAAATTAATCTGGAAAGAACTTGGAACTCTCAACCATAGGTTTTGGTACGAAATTGTT
GCTTGTCAGAACCAAATGATAGGCTATTGCCTTGAAATAGTGTTTCTTGTGGTTTCCAATATTGGAAGTTAAAATCATAT
GACTTAGCTGTTGGATACTAATTAAGCTTAAGCAATGCCAACTCTAAGAAGTGGTACTTACACAATATTCTATTGGTCAT
AGGTATAGTTGAATCAAGTATCAAGCGTGGAATGCATATTGAAGCTCTTGAGATGGTTTATACCTTTGGCATGGAGGATA
AGTTTTCAGCTGCTCTAGTTCTAACTTCATTCTTAAAGATGAGCAAGGAGTCATTTGAGAGGGCAAAACGGAAAGCCCAG
TCACCGCTGGCATTTGTATGAACCCTTCCCTTGCACATTATGTACCTTTATGAACTCTTTATCATCATCTGAGTCTGACC
ATTGATATATTTATTTCTCAACAGAAAGAAGCGGCTACAAAGCAGCTAGCTGTGTTATCATCAGTTATGCAGTGTATGGA
GACTCACAAGTTAGATCCTGCGAAAGAACTACCAGGATGGCAGATCAAAGAGCAAATTGTTAGCTTGGAGAAAGACACTC
TTCAGCTCGACAAAGAGATGGAAGAGAAAGCAAGATCTCTCAGTTTAATGGAGGAAGCTGCACTTGCCAAGAGAATGTAT
AACCAACAGATAAAACGTCCAAGGTTGTCACCCATGGAAATGCCACCAGTAACTTCTTCATCGTATTCTCCTATCTACCG
TGATAGAAGCTTTCCTAGTCAAAGAGACGATGACCAAGATGAAATATCAGCTCTTGTGAGTAGTTACCTCGGCCCGTCAA
CATCTTTTCCTCATCGCTCAAGAAGATCCCCGGAATATATGGTTCCACTTCCACATGGTGGGTTAGGAAGAAGTGTATAT
GCATATGAACATCTGGCCCCAAATTCATACTCTCCAGGTCACGGACATAGACTTCATCGACAGTACTCTCCGTCTTTGGT
TCACGGACAGAGACATCCACTACAGTACTCTCCTCCAATTCATGGACAACAACAGTTACCATATGGTATACAAAGGGTTT
ACAGACATTCACCATCTGAAGAAAGATATTTGGGTTTATCCAATCAAAGGTCTCCTCGCAGTAACTCATCATTAGACCCC
AAATAGGAGGAATGTAAATTTGTAACAAAGCTTTTTGTTTTTGCTTAAGTTAGTCATTTATTTAACTCCCAACAGTCTCA
AAATTTAATTTAATGTTTGGGGCTTAAGAATGCAAATTTTTTTGCTCCTGTAATTGACATTTAAGATGCTAATGTTATTG
CTTCAGAGGTTTTAGTCAACCTCAGATACATCGATATCACTATCTAAATAGACCTCTGGCTCTTGGTCATCTGGATTCTC
TTCATCTTCTGTCTCTGTTCCTTCTTGTTCTCGTTGCACTGCTCGAGCAATTGCGGATTCCAACCTTGTGCTTACAGTTT
CCCATGACACAAGCTTTTCCATGAATGTATTTATGTCCGCCTTCTTATCTTTCTTGAGGAAGATGAATT

>FRI_C24
GGATCCCAAAATCTAGTGCCGCCGAGTCCTTTGGGACACAGTTTCGCACTTATCATTCGATTCGGCAAGCAGTAACTCGC

AAAATCCGCACCACCGTGAAACATACTCCGGCGCTGAACGCTAGACGCAAAATCCATATTACTGCTGCAGAAAGACGGAG
AGAACAAGAGACTGACCGATCATAAGAGAAGAGAGCTTCAAGGCTTCAGCTGATGAAGAATCAATGGCGGCGAGAGAGAG
TGAGAATCGAGTAGTTACTGAGAGGAAGATGAAACGTATCTCCTGAGAAAAAACGTAAGAGAGTTCCGTTTCAACAGTAT
TTTTCTCGGCGGTCTTCAGATTTTCCGTCCAACTCTCTCTTTCCTCTTTCAGTTTTTTTTGCCCAATCACTCGTGGTAGG
ATTTTTTATAAGAAAGTTACGAAAATGCCCTCTCGTGCTTAGAGAGATTCAATTCGTGCACATATAATATGCCGTGAATA
TACCACCAACTCTCTTACTCGTACTCGCATGTGGTGCGTGCAATATACACGTGGTTCGAATCAACGGCTACGTTTTACTG
ATAGGTGGCTCGTTTAGTTTTGCACTGCCTTTGGGTGCGCGTGACCAAGTCATTTTCCAAATGTGTTTTAATATTTGTGA
ACTATTATTAATAAAAAAGCGAGCGACGTTGTTCGTTGATGAGGGAGCGACCGTACGCTTGGCGACCTTCTCTGATTGCG
TAGAGTCGTTTTCCTTATTGGGCTTACCTAGTACCGTCAGCCCACAATGTTTTCATTTCTTTCCCACCTAAAACTAAAAG
TACTCACAAGTCACAACTTAAACCAAGTACACAAGGATTTTATCATGGGATTATCGTGTTTGAAGACTAAAAAGAGCACA
CCATCACCCCCATTAGTGCAGGTAGAGTAAGACAGTAACTTTTGGGTTCATATTACCGAGCAAGAACCGTTATTTGTGAT
TAGACATGTTATAAACCACTGCTTTAGTGACTATTTAAAACAATATATTACATGTCGTAATCATGCAACCTAACTATGTT
TTCATTAATCAAATACAAAGAATAAAGAGAAAAGTGCGTAGATTCAATTATTTGGCATAGACTCAAAAGAGTGTATATAT
ATCTGACTTTTATTAAATTATTAAACACAAATACATATTTTCATAAGCAAAACTATAAAAGCCCTAAACATATAACGATT
ACCTCAAAGGAAAAAGTCGTTTTCTCCTAATTAAAAGATAGGTTACTTCCTAATTAATATATAATTTATGTGAACTTCAC
AATATACAGTTCAATAAAATTTGGTAATTTGACCGATTTAAGGAGAGTGGAAATTAGGGCTTCTGCAATCTTTTTTCTTC
GCCGCAATCTCATGTCCAATTATCCACCGACGGTGGCGGCGCAACCCACAACGACGGCGAATCCACTGCTGCAGCGACAT
CAATCTGAACAGCGCCGAAGAGAATTACCGAAGATTGTCGAAACAGAGTCTACAAGTATGGACATTACGATCGGTCAATC
TAAGCAGCCTCAATTTTTGAAATCCATAGACGAATTAGCTGCGTTTTCAGTTGCAGTGGAAACATTCAAACGCCAATTCG
ATGATCTTCAGAAGCACATCGAGTCAATCGAAAACGCAATTGATTCCAAACTCGAGAGTAACGGCGTtGTCCTCGCCGCG
CGgAACAATAATTTCCATCAGCCGATGTTATCGCCTCCGCGGAACAATGTATCTGTAGAAACCACCGTCACTGTGAGCCA
ACCGTCTCAGGAGATTGTACCGGAGACGTCGAATAAACCGGAGGGGGAACGTATGTGTGAGTTGATGTGTAGCAAAGGTC
TGCGTAAATACATATACGCGAATATCTCTGATCAAGCTAAGTTAATGGAAGAGATTCCTTCAGCTTTGAAATTGGCCAAG
GAGCCAGCGAAGTTTGTATTGGATTGTATTGGCAAGTTTTACTTACAAGGGCGTAGAGCATTTACTAAAGAGTCGCCTAT
GAGCTCTGCGAGACAAGTTTCGCTTCTTATACTGGAGTCTTTTCTTCTAATGCCTGATCGTGGTAAAGGGAAGGTGAAGA
TTGAGAGTTGGATTAAAGATGAGGCGGAGACGGCTGCTGTTGCTTGGAGGAAAAGGTTGATGACTGAAGGAGGATTAGCT
GCGGCTGAGAAAATGGATGCAAGGGGTTTGCTTTTACTAGTTGCTTGTTTTGGTGTTCCTTCAAACTTTAGGAGTACAGA
TTTGCTGGATTTGATAAGGATGAGTGGTTCGAATGAGATTGCCGGTGCTTTGAAGCGGTCACAGTTTCTTGTCCCTATGG
TCTCAGGTACCATATTCTGTTCTCACTCGGTGAATTTCATTGCAAAGGTGGTTCCTTTTGTTGACATCATCGACCAACAT
CAAGTTCCATCTTTGTTTTTCGATAAGCTTGATGGTATAAACTAGGAGAGCACATCAAATATTTAGAGTGCAATGACTGA
TTGAGCCAAATCCTAGCTAGAAATTAATCTGGAAAGAACTTGGAACTCTCAACCATAGGTTTTGGTACGAAATTGTTGCT
TGTCAGAACCAAATGATAGGCTATTGCCTTGAAATAGTGTTTCTTGTGGTTTCCAATATTGGAAGTTAAAATCATATGAC
TTAGCTGTTGGATACTAATTAAGCTTAAGCAATGCCAACTCTAAGAAGTGGTACTTACACAATATTCTATTGGTCATAGG
TATAGTTGAATCAAGTATCAAGCGTGGAATGCATATTGAAGCTCTTGAAATGGTTTATACCTTTGGCATGGAGGATAAGT
TTTCAGCTGCTCTAGTTCTAACTTCATTCTTAAAGATGAGCAAGGAGTCATTTGAGAGGGCAAAACGGAAAGCCCAGTCA
CCGCTGGCATTTGTATGAACCCTTCCCTTGCACATTATGTACCTTTATGAACTCTTTATCATCATCTGAGTCTGACCATT
GATATATTTATTTCTCAACAGAAAGAAGCGGCTACAAAGCAGCTAGCTGTGTTATCATCAGTTATGCAGTGTATGGAGAC
TCACAAGTTAGATCCTGCGAAAGAACTACCAGGATGGCAGATCAAAGAGCAAATTGTTAGCTTGGAGAAAGACACTCTTC
AGCTCGACAAAGAGATGGAAGAGAAAGCAAGATCTCTCAGTTTAATGGAGGAAGCCGCACTTGCCAAGAGAATGTATAAC
CAACAGATAAAACGTCCAAGGTTGTCACCCATGGAAATGCCACCAGTAACTTCTTCATCGTATTCTCCTATCTACCGTGA
TAGAAGCTTTCCTAGTCAAAGAGACGATGACCAAGATGAAATATCAGCTCTTGTGAGTAGTTACCTCGGCCCGTCAACAT
CTTTTCCTCATCGCTCAAGAAGATCCCCGGAATATATGGTTCCACTTCCACATGGTGGGTTAGGAAGAAGTGTATATGCA
TATGAACATCTGGCCCCAAATTCATATTCTCCAGGTCACGGACATAGACTTCATCGACAGTACTCTCCGTCTTTGGTTCA
CGGACAGAGACATCCACTACAGTACTCTCCTCCAATTCATGGACAACAACAGTTACCATATGGTATACAAAGGGTTTACA
GACATTCACCATCTGAAGAAAGATATTTGGGTTTATCCAATCAAAGGTCTCCTCGCAGTAACTCATCATTAGACCCCAAA
TAGGAGGAATGTAAATTTGTAACAAAGCTTTTTGTTTTTGCTTAAGTTAGTCATTTATTTAACTCCCAACAGTCTCAAAA
TTTAATTTAATGTTTGGGGCTTAAGAATGCAAATTTTTTTGCTCCTGTAATTGACATTTAAGATGCTAATGTTATTGCTT
CAGAGGTTTTAGTCAACCTCAGATACATCGATATCACTATCTAAATAGACCTCTGGCTCTTGGTCATCTGGATTCTCTTC
ATCTTCTGTCTCTGTTCCTTCTTGTTCTCGTTGCACTGCTCGAGCAATTGCGGATTCCAACCTTGTGCTTACAGTTTCCC
ATGACACAAGCTTTTCCATGAATGTATTTATGTCCGCCTTCTTATCTTTCTTGAGGAAGA

>FRI_Nok-3
GGATCCCAAAATCTAGTGCCGCCGAGTCCTTTGGGACAGAGTTTCGCACTTATCATTCGATTCGGCAAGCAGTAACTCGC

AAAATCCGCACCACCGTGAAACATACTCTGGCGCTGAACGCTAGACGCAAAATCCATATTACTGCTGCAGAAAGACGGAG
AGAACAAGAGACTGACCGATCATAAGAGAAGAGAGCTTCAAGGCTTCAGCTGATGAAGAATCAATGGCGGCGAGAGTGAG
TGAGAATCGAGTAGTTACTGAGAGGAAGATGAAACGTATCTCCTGAGAAAAAACGTAAGAGAGTTCCGTTTCAACAGTAT
TTTTCTCGGCGGTCTTCAGATTTTCCGTCCAACTCTCTCTTTCCTCTTTCAGTTTTTTTTTGCCCAATCACTCGTGGTAG
GATTTTTTTATAAGAAAGTTACGAAAATGCCCTCTCGTGCTTAGAGAGATTCAATTCGTGCACATATAATATGCCGTGAA
TATACCACCAACTCTCTTACTCGTACTCGCATGTGGTGCGTGCAATATACACGTGGTTCGAATCAACGGCTACGTTTTAC
TGATAGGTGGCTCGTTTAGTTTTGCACTGCCTTTGGGTGCGCGTGACCAAGTCATTTTCCAAATGTGTTTTAATATTTGT
GAACTATTATTAATAAAAAAGCGAGCGACGTTGTTCGTTGATGAGGGAGCGACCGTACGCTTGGCGACCTTCTCTGATTG
CGTAGAGTCGTTTTCCTTATTGGGCTTACCTAGTACCGCCAGCCCACAATGTTTTCATTTCTTTCCCACCTAAAACTAAA
AGTACTCACAAGTCACAACTTAAACCAAGTACACAAGGATTTTATCATGGGATTATCGTGTTTGAAGACTAAAAAGAGCA
CACCATCACCCCCATTAGTGCAGGTAGAGTAAGACAGTAACTTTTGGGTTCATATTACCGAGCAAGAACCGTTATTTGTG
ATTAGACATGTTATAAACCACTGCTTTAGTGACTATTTAAAACAATATATTACATGTCGTAATCATGCAACCTAACTATG
TTTTCATTAATCAAATACAAAGAATAAAGAGAAAAGTGCGTAGATTCAATTACTTGGCATAGACTCAAAAGAGTGTATAT
ATATCTGACTTTTATTAAATTATTAAACACAAATACATATTTTCATAAGCAAAACTATAAAAGCCCTAAACATATAATGA
TTACCTCAAAGGAAAAAGTCGTTTTCTCCTAATTAAAAGATAGGTTACTTCCTAATTAATATATAATTTATGTGAACTTC
ACAATATACAGTTCAATAAAATTTGGTAATTTGACCGATTTAAGGAGAGTGGAAATTAGGGCTTCTGCAATCTTTTTTCT
TCGCCGCAATCTCATGTCCAATTATCCACCGACGGTGGCGGCGCAACCCACAACGACGGCGAATCCACTGCTGCAGCGAC
ATCAATCTGAACAGCGACGAAGAGAATTACCGAAGATTGTCGAAACAGAGTCTACAAGTATGGACATTACGATCGGTCAA
TCTAAGCAGcCTCAATTTTTGAAATCCATAGAGGAATTAGCTGCGTTTTCAGTTGCAGTGGAAACATTCAAACGCCAATT
CGATGATCTTCAGAAGCACATCGAGTCAATCGAAAACGCAATTGATTCCAAACTCGAGAGTAACGGCGTTGTCCTCGCCG
CGCGGAACAATAATTTCCATCAGCCGATGTTATCGCCTCCGCGGAACAATGTATCTGTAGAAACCACCGTCACTGTGAGC
CAACCGTCTCAGGAGATTGTACCGGAGACGTCGAATAAACCGGAGGGGGAACGTATGTGTGAGTTGATGTGTAGCAAAGG
TCTGCGTAAATACATATACGCGAATATCTCTGATCAAGCTAAGTTAATGGAAGAGATTCCTTCAGCTTTGAAATTGGCCA
AGGAGCCAGCGAAGTTTGTATTGGATTGTATTGGCAAGTTTTACTTACAAGGGCGTAGAGCATTTACTAAAGAGTCGCCT
ATGAGCTCTGCGAGACAAGTTTCGCTTCTTATACTGGAGTCTTTTCTTCTAATGCCTGATCGTGGTAAAGGGAAGGTGAA
GATTGAGAGTTGGATTAAAGATGAGGCGGAGACGGCTGCTGTTGCTTGGAGGAAAAGGTTGATGACTGAAGGAGGATTAG
CTGCGGCTGAGAAAATGGATGCAAGGGGTTTGCTTTTACTAGTTGCTTGTTTTGGTGTTCCTTCAAACTTTAGGAGTACA
GATTTGCTGGATTTGATAAGGATGAGTGGTTCGAATGAGATTGCCGGTGCTTTGAAGCGGTCACAGTTTCTTGTCCCTAT
GGTCTCAGGTACCATATTCTGTTCTCACTCGGTGAATTTCATTGCAAAGGTGGTTCCTTTTGTTGACATCATCGACCAAC
ATCAAGTTCCATCTTTGTTTTTCGATAAGCTTGATGGTATAAACTAGGAGAGCACATCAAATATTTAGAGTGCAATGACT
GATTGAGCCAAATCCTAGCTAGAAATTAATCTGGAAAGAACTTGGAACTCTCAACCATAGGTTTTGGTACGAAATTGTTG
CTTGTCAGAACCAAATGATAGGCTATTGCCTTGAAATAGTGTTTCTTGTGGTTTCCAATATTGGAAGTTAAAATCATATG
ACTTAGCTGTTGGATACTAATTAAGCTTAAGCAATGCCAACTCTAAGAAGTGGTACTTACACAATATTCTATTGGTCATA
GGTATAGTTGAATCAAGTATCAAGCGTGGAATGCATATTGAAGCTCTTGAAATGGTTTATACCTTTGGCATGGAGGATAA
GTTTTCAGCTGCTCTAGTTCTAACTTCATTCTTAAAGATGAGCAAGGAGTCATTTGAGAGGGCAAAACGGAAAGCCCAGT
CACCGCTGGCATTTGTATGAACCCTTCCCTTGCACATTATGTACCTTTATGAACTCTTTATCATCATCTGAGTCTGACCA
TTGATATATTTATTTCTCAACAGAAAGAAGCGGCTACAAAGCAGCTAGCTGTGTTATCATCAGTTATGCAGTGTATGGAG
ACTCACAAGTTAGATCCTGCGAAAGAACTACCAGGATGGCAGATCAAAGAGCAAATTGTTAGCTTGGAGAAAGACACTCT
TCAGCTCGACAAAGAGATGGAAGAGAAAGCAAGATCTCTCAGTTTAATGGAGGAAGCCGCACTTGCCAAGAGAATGTATA
ACCAACAGATAAAACGTCCAAGGTTGTCACCCATGGAAATGCCACCAGTAACTTCTTCATCGTATTCTCCTATCTACCGT
GATAGAAGCTTTCCTAGTCAAAGAGACGATGACCAAGATGAAATATCAGCTCTTGTGAGTAGTTACCTCGGCCCGTCAAC
ATCTTTTCCTCATCGCTCAAGAAGATCCCCGGAATATATGGTTCCACTTCCACATGGTGGGTTAGGAAGAAGTGTATATG
CATATGAACATCTGGCCCCAAATTCATACTCTCCAGGTCACGGACATAGACTTCATCGACAGTACTCTCCGTCTTTGGTT
CACGGACAGAGACATCCACTACAGTACTCTCCTCCAATTCATGGACAACAACAGTTACCATATGGTATACAAAGGGTTTA
CAGACATTCACCATCTGAAGAAAGATATTTGGGTTtATCCAATCAAAGGTCTCCTCGCAGTAACTCATCATTAGACCCCA
AATAGGAGGaATGTAAATTTGTAACAAAGCTTTTTGTTTTTGCTTAAGTTAGTCATTTATTTAACTCCCAACAGTCTCAA
AATTTAATTTAATGTTTGGGGCTTAAGAATGCAAATTTTTTTGCTCCTGTAATTGACATTTAAGATGCTAATGTTATTGC
TTCAGAGGTTTTAGTCAACCTCAGATACATCGATATCACTATCTAAATAGACCTCTGGCTCTTGGTCATCTGGATTCTCT
TCATCTTCTGTCTCTGTTCCTTCTTGTTCTCGTTGCACTGCTCGAGCAATTGCGGATTCCAACCTTGTGCTTACAGTTTC
CCATGACACAAGCTTTTCCATGAATGTATTTATGTCCGCCTTCTTATCTTTCTTGAGGAAGATGAATTC

>FRI_Edi-0
GGATCCCAAAATCTAGTGCCGCCGAGTCCTTTGGGACAGAGTTTCGCACTTATCATTCGATTCGGCAAGCAGTAACTCGC

AAAATCCGCACCACCGTGAAACATACTCTGGCGCTGAACGCTAGACGCAAAATCCATATTACTGCTGCAGAAAGACGGAG
AGAACAAGAGACTGACCGATCATAAGAGAAGAGAGCTTCAAGGCTTCAGCTGATGAAGAATCAATGGCGGCGAGAGTGAG
TGAGAATCGAGTAGTTACTGAGAGGAAGATGAAACGTATCTCCTGAGAAAAAACGTAAGAGAGTTCCGTTTCAACAGTAT
TTTTCTCGGCGGTCTTCAGATTTTCCGTCCAACTCTCTCTTTCCTCTTTCAGTTTTTTTTTGCCCAATCACTCGTGGTAG
GATTTTTTTATAAGAAAGTTACGAAAATGCCCTCTCGTGCTTAGAGAGATTCAATTCGTGCACATATAATATGCCGTGAA
TATACCACCAACTCTCTTACTCGTACTCGCATGTGGTGCGTGCAATATACACGTGGTTCGAATCAACGGCTACGTTTTAC
TGATAGGTGGCTCGTTTAGTTTTGCACTGCCTTTGGGTGCGCGTGACCAAGTCATTTTCCAAATGTGTTTTAATATTTGT
GAACTATTATTAATAAAAAAGCGAGCGACGTTGTTCGTTGATGAGGGAGCGACCGTACGCTTGGCGACCTTCTCTGATTG
CGTAGAGTCGTTTTCCTTATTGGGCTTACCTAGTACCGTCAGCCCACAATGTTTTCATTTCTTTCCCACCTAAAACTAAA
AGTACTCACAAGTCACAACTTAAACCAAGTACACAAGGATTTTATCATGGGATTATCGTGTTTGAAGACTAAAAAGAGCA
CACCATCACCCCCATTAGTGCAGGTAGAGTAAGACAGTAACTTTTGGGTTCATATTACCGAGCAAGAACCGTTATTTGTG
ATTAGACATGTTATAAACCACTGCTTTAGTGACTATTTAAAACAATATATTACATGTCGTAATCATGCAACCTAACTATG
TTTTCATTAATCAAATACAAAGAATAAAGAGAAAAGTGCGTAGATTCAATTATTTGGCATAGACTCAAAAGAGTGTATAT
ATATCTGACTTTTATTAAATTATTAAACACAAATACATATTTTCATAAGCAAAACTATAAAAGCCCTAAACATATAACGA
TTACCTCAAAGGAAAAAGTCGTTTTCTCCTAATTAAAAGATAGGTTACTTCCTAATTAATATATAATTTATGTGAACTTC
ACAATATACAGTTCAATAAAATTTGGTAATTTGACCGATTTAAGGAGAGTGGAAATTAGGGCTTCTGCAATCTTTTTTCT
TCGCCGCAATCTCATGTCCAATTATCCACCGACGGTGGCGGCGCAACCCACAACGACGGCGAATCCACTGCTGCAGCGAC
ATCAATCTGAACAGCGACGAAGAGAATTACCGAAGATTGTCGAAACAGAGTCTACAAGTATGGACATTACGATCGGTCAA
TCTAAGCAGCCTCAATTTTTGAAATCCATAGACGAATTAGCTGCGTTTTCAGTTGCAGTGGAAACATTCAAACGCCAATT
CGATGATCTTCAGAAGCACATCGAGTCAATCGAAAACGCAATTGATTCCAAACTCGAGAGTAACGGCGTTGTCCTCGCCG
CGCGGAACAATAATTTCCATCAGCCGATGTTATCGCCTCCGCGGaACAATGTATCTGTAGAAACCACCGTCACTGTGAGC
CAACCGTCTCAGGAGATTGTACCGGAGACGTCGAATAAACCGGAGGGGGAACGTATATGTGAGTTGATGTGTAGCAAAGG
TCTGCGTAAATACATATACGCGAATATCTCTGATCAAGCTAAGTTAATGGAAGAGATTCCTTCAGCTTTGAAATTGGCCA
AGGAGCCAGCGAAGTTTGTATTGGATTGTATTGGCAAGTTTTACTTACAAGGGCGTAGAGCATTTACTAAAGAGTCGCCT
ATGAGCTCTGCGAGACAAGTTTCGCTTCTTATACTGGAGTCTTTTCTTCTAATGCCTGATCGTGGTAAAGGGAAGGTGAA
GATTGAGAGTTGGATTAAAGATGAGGCGGAGACGGCTGCTGTTGCTTGGAGGAAAAGGTTGATGACTGAAGGAGGATTAG
CTGCGGCTGAGAAAATGGATGCAAGGGGTTTGCTTTTACTAGTTGCTTGTTTTGGTGTTCCTTCAAACTTTAGGAGTACA
GATTTGCTGGATTTGATAAGGATGAGTGGTTCGAATGAGATTGCCGGTGCTTTGAAGCGGTCACAGTTTCTTGTCCCTAT
GGTCTCAGGTACCATATTCTGTTCTCACTCGGTGAATTTCATTGCAAAGGTGGTTCCTTTTGTTGACATCATCGACCAAC
ATCAAGTTCCATCTTTGTTTTTCGATAAGCTTGATGGTATAAACTAGGAGAGCACATCAAATATTTAGAGTGCAATGACT
GATTGAGCCAAATCCTAGCTAGAAATTAATCTGGAAAGAACTTGGAACTCTCAACCATAGGTTTTGGTACGAAATTGTTG
CTTGTCAGAACCAAATGATAGGCTATTGCCTTGAAATAGTGTTTCTTGTGGTTTCCAATATTGGAAGTTAAAATCATATG
ACTTAGCTGTTGGATACTAATTAAGCTTAAGCAATGCCAACTCTAAGAAGTGGTACTTACACAATATTCTATTGGTCATA
GGTATAGTTGAATCAAGTATCAAGCGTGGAATGCATATTGAAGCTCTTGAAATGGTTTATACCTTTGGCATGGAGGATAA
GTTTTCAGCTGCTCTAGTTCTAACTTCATTCTTAAAGATGAGCAAGGAGTCATTTGAGAGGGCAAAACGGAAAGCCCAGT
CACCGCTGGCATTTGTATGAACCCTTCCCTTGCACATTATGTACCTTTATGAACTCTTTATCATCATCTGAGTCTGACCA
TTGATATATTTATTTCTCAACAGAAAGAAGCGGCTACAAAGCAGCTAGCTGTGTTATCATCAGTTATGCAGTGTATGGAG
ACTCACAAGTTAGATCCTGCGAAAGAACTACCAGGATGGCAGATCAAAGAGCAAATTGTTAGCTTGGAGAAAGACACTCT
TCAGCTCGACAAAGAGATGGAAGAGAAAGCAAGATCTCTCAGTTTAATGGAGGAAGCCGCACTTGCCAAGAGAATGTATA
ACCAACAGATAAAACGTCCAAGGTTATCACCCATGGAAATGCCACCAGTAACTTCTTCATCGTATTCTCCTATCTACCGT
GATAGAAGCTTTCCTAGTCAAAGAGACGATGACCAAGATGAAATATCAGCTCTTGTGAGTAGTTACCTCGGCCCGTCAAC
ATCTTTTCCTCATCGCTCAAGAAGATCCCCGGAATATATGGTTCCACTTCCACATGGTGGGTTAGGAAGAAGTGTATATG
CATATGAACATCTGGCCCCAAATTCATATTCTCCAGGTCACGGACATAGACTTCATCGACAGTACTCTCCGTCTTTGGTT
CACGGACAGAGACATCCACTACAGTACTCTCCTCCAATTCATGGACAACAACAGTTACCATATGGTATACAAAGGGTTTA
CAGACATTCACCATCTGAAGAAAGATATTTGGGTTtATCCAATCAAAGGTCTCCTCGCAGTAACTCATCATTAGACCCCA
AATAGGAGGaATGTAAATTTGTAACAAAGCTTTTTGTTTTTGCTTAAGTTAGTCATTTATTTAACTCCCAACAGTCTCAA
AATTTAATTTAATGTTTGGGGCTTAAGAATGCAAATTTTTTTGCTCCTGTAATTGACATTTAAGATGCTAATGTTATTGC
TTCAGAGGTTTTAGTCAACCTCAGATACATCGATATCACTATCTAAATAGACCTCTGGCTCTTGGTCATCTGGATTCTCT
TCATCTTCTGTCTCTGTTCCTTCTTGTTCTCGTTGCACTGCTCGAGCAATTGCGGATTCCAACCTTGTGCTTACAGTTTC
CCATGACACAAGCTTTTCCATGAATGTATTTATGTCCGCCTTCTTATCTTTCTTGAGGAAGATGAATTC

>FRI_St-0
GGATCCCAAAATCTAGTGCCGCCGAGTCCTTTGGGACACAGTTTCGCACTTATCATTCGATTCGGCAAGCAGTAACTCGC

AAAATCCGCACCACCGTGAAACATACTCCGGCGCTGAACGCTAGACGCAAAATCCATATTACAGCTGCAGAAAGACGGAG
AGAACAAGAGACTGACCGATCATAAGAGAAGAGAGCTTCAAGGCTTCAGCTGATGAAGAATCAATGGCGGCGAGAGAGAG
TGAGAATCGAGTAGTTACTGAGAGGAAGATGAAACGTATCTCCTGAGAAAAAACGTAAGAGAGTTCCGTTTCAACAGTAT
TTTTCTCGGCGGTCTTCAGATTTTCCGTCCAACTCTCTCTTTCCTCTTTCAGTTTTTTTTGCCCAATCACTCGTGGTAGG
ATTTTTTATAAGAAAGTTACGAAAATGCCCTCTCGTGCTTAGAGAGATTCAATTCGTGCACATATAATATGCCGTGAATA
TACCACCAACTCTCTTACTCGTACTCGCATGTGGTGCGTGCAATATACACGTGATTCGAATCAACGGCTACGTTTTACTG
ATAGGTGGCTCGTTTAGTTTTGCACTGCCTTTGGGTGCGCGTGACCAAGTCATTTTCCAAATGTGTTTTAATATTTGTGA
ACTATTATTAATAAAAAAGCGAGCGACGTTGTTCTTTGATGAGGGAACGACCGTACGCTTGGCGACCTTCTCTGATTGCG
TAGAGTCGTTTTCCTTATTGGGCTTACCTAGTACCGCCAGCCCACAATGTTTTCATTTCTTTCCCACCTAAAACTAAAAG
TACTCACAAGTCACAACTTAAACCAAGTACAGGTAGAGTAAGACAGTAACTTTTGGGTTCATATTACCGAGCAAGAACCG
TTATTTGTGATTAGACATGTTATAAACCACTGCTTTAGTGACTATTTAAAACAATATATTACATGTCGTAATCATGCAAC
CTAACTATGTTTTCATTAATCAAATACAAAGAATAAAGAGAAAAGTGCGTAGATTCAATTATTTGGCATAGACTCAAAAG
AGTGTATATATATCTGACTTTTATTAAATTATTAAACACAAATACATATTTTCATAAGCAAAACTATAAAAGCCCTAAAC
ATATAACGATTACCTCAAAGGAAAAAGTCGTTTTCTCCTAATTAAAAGATAGGTTACTTCCTAATTAATATATATAATTTAT
GTGAACTTCACAATATACAGTTCAATAAAATTTGGTAATTTGACCGATTTAAGGAGAGTGGAAATTAGGGCTTCTGCAAT
CTTTTTTCTTCGCCGCAATCTCATGTCCAATTATCCACCGACGGTGGCGGCGCAACCCACAACGACGGCGAATCCACTGC
TGCAGCGACATCAATCTGAACAGCGACGAAGAGAATTACCGAAGATTGTCGAAACAGAGTCTACAAGTATGGACATTACG
ATCGGTCAATCTAAGCAGcCTCAATTTTTGAAATCCATAGACGAATTAGCTGCGTTTTCAGTTGCAGTGGAAACATTCAA
ACGCCAATTCGATGATCTTCAGAAGCACATCGAGTCAATCGAAAACGCAATTGATTCCAAACTCGAGAGTAACGGCGTTG
TCCTCGCCGCGCGGAACAATAATTTCTATCAGCCGATGTTATCGCCTCCGCGGAACAATGTATCTGTAGAAACCACCGTC
ACTGTGAGCCAACCGTCTCAGGAGATTGTACCGGAGACGTCGAATAAACCGGAGGGGGAACGTATGTGTGAGTTGATGTG
TAGCAAAGGTCTGCGTAAATACATATACGCGAATATCTCTGATCAAGCTAAGTTAATGGAAGAGATTCCTTCAGCTTTGA
AATTGGCCAAGGAGCCAGCGAAGTTTGTATTGGATTGTATTGGCAAGTTTTACTTACAAGGGCGTAGAGCATTTACTAAA
GAGTCGCCTATGAGCTCTGCGAGACAAGTTTCGCTTCTTATACTGGAGTCTTTTCTTCTAATGCCTGATCGTGGTAAAGG
GAAGGTGAAGATTGAGAGTTGGATTAAAGATGAGGCGGAGACGGCTGCTGTTGCTTGGAGGAAAAGGTTGATGACTGAAG
GAGGATTAGCTGCGGCTGAGAAAATGGATGCAAGGGGTTTGCTTTTACTAGTTGCTTGTTTTGGTGTTCCTTCAAACTTT
AGGAGTACAGATTTGCTGGATTTGATAAGGATGAGTGGTTCGAATGAGATTGCCGGTGCTTTGAAGCGGTCACAGTTTCT
TGTCCCTATGGTCTCAGGTACCATATTCTGTtCTCACTCGGTGAATTTCATTGCAAAGGTGGTTCCTTTTGTTGACATCA
TCGACCAACATCAAGTTCCATCTTTGTTTTTCGATAAGCTTGATGGTATAAACTAGGAGAGCACATCAAATATTTAGAGT
GCAATGACTGATTGAGCCAAATCCTAGCTAGAAATTAATCTGGAAAGAACTTGGAACTCTCAACCATAGGTTTTGGTACG
AAATTGTTGCTTGTCAGAACCAAATGATAGGCTATTGCCTTGAAATAGTGTTTCTTGTGGTTTCCAATATTGGAAGTTAA
AATCATATGACTTAGCTGTTGGATACTAATTAAGCTTAAGCAATGCCAACTCTAAGAAGTGGTACTTACACAATATTCTA
TTGGTCATAGGTATAGTTGAATCAAGTATCAAGCGTGGAATGCATATTGAAGCTCTTGAAATGGTTTATACCTTTGGCAT
GGAGGATAAGTTTTCAGCTGCTCTAGTTCTAACTTCATTCTTAAAGATGAGCAAGGAGTCATTTGAGAGGGCAAAACGGA
AAGCCCAGTCACCGCTGGCATTTGTATGAACCCTTCCCTTGCACATTATGTACCTTTATGAACTCTTTATCATCATCTGA
GTCTGACCATTGATATATTTATTTCTCAACAGAAAGAAGCGGCTACAAAGCAGCTAGCTGTGTTATCATCAGTTATGCAG
TGTATGGAGACTCACAAGTTAGATCCTGCGAAAGAACTACCAGGATGGCAGATCAAAGAGCAAATTGTTAGCTTGGAGAA
AGACACTCTTCAGCTCGACAAAGAGATGGAAGAGAAAGCAAGATCTCTCAGTTTAATGGAGGAAGCCGCACTTGCCAAGA
GAATGTATAACCAACAGATAAAACGTCCAAGGTTGTCACCCATGGAAATGCCACCAGTAACTTCTTCATCGTATTCTCCT
ATCTACCGTGATAGAAGCTTTCCTAGTCAAAGAGACGATGACCAAGATGAAATATCAGCTCTTGTGAGTAGTTACCTCGG
CCCGTCAACATCTTTTCCTCATCGCTCAAGAAGATCCCCGGAATATATGGTTCCACTTCCACATGGTGGGTTAGGAAGAA
GTGTATATGCATATGAACATCTGGCCCCAAATTCATATTCTCCAGGTCACGGACATAGACTTCATCGACAGTACTCTCCG
TCTTTGGTTCACGGACAGAGACATCCACTACAGTACTCTCCTCCAATTCATGGACAACAACAGTTACCATATGGTATACA
AAGGGTTTACAGACATTCACCATCTGAAGAAAGATATTTGGGTTTATCCAATCAAAGGTCTCCTCGCAGTAACTCATCAT
TAGACCCCAAATAGGAGGaATGTAAATTTGTAACAAAGCTTTTTGTTTTTGCTTAAGTTAGTCATTTATTTAACTCCCAA
CAGTCTCAAAATTTAATTTAATGTTTGGGGCTTAAGAATGCAAATTTTTTTGCTCCTGTAATTGACATTTAAGATGCTAA
TGTTATTGCTTCAGAGGTTTTAGTCAACCTCAGATACATCGATATCACTATCTAAATAGACCTCTGGCTCTTGGTCATCT
GGATTCTCTTCATCTTCTGTCTCTGTTCCTTCTTGTTCTCGTTGCACTGCTCGAGCAATTGCGGATTCCAACCTTGTGCT
TACAGTTTCCCATGACACAAGCTTTTCCATGAATGTATTTATGTCCGCCTTCTTATCTTTCTTGAGGAAGATGAATTC

>FRI_Pu-2-23
GGATCCCAAAATCTAGTGCCGCCGAGTCCTTTGGGACAGAGTTTCGCACTTATCATTCGATTCGGCAAGCAGTAACTCGC
AAAATCCGCACCACCGTGAAACATACTCTGGCGCTGAACGCTAGACGCAAAATCCATATTACTGCTGCAGAAAGACGGAG

AGAACAAGAGACTGACCGATCATAAGAGAAGAGAGCTTCAAGGCTTCAGCTGATGAAGAATCAATGGCGGCGAGAGAGAG
TGAGAATCGAGTAGTTACTGAGAGGAAGATGAAACGTATCTCCTGAGAAAAAACGTAAGAGAGTTCCGTTTCAACAGTAT
TTTTCTCGGCGGTCTTCAGATTTTCCGTCCAACTCTCTCTTTCCTCTTTCAGTTTTTTTTTGCCCAATCACTCGTGGTAG
GATTTTTTTATAAGAAAGTTACGAAAATGCCCTCTCGTGCTTAGAGAGATTCAATTCGTGCACATATAATATGCCGTGAA
TATACCACCAACTCTCTTACTCGTACTCGCATGTGGTGCGTGCAATATACACGTGGTTCGAATCAACGGCTACGTTTTAC
TGATAGGTGGCTCGTTTAGTTTTGCACTGCCTTTGGGTGCGCGTGACCAAGTCATTTTCCAAATGTGTTTTAATATTTGT
GAACTATTATTAATAAAAAAGCGAGCGACGTTGTTCGTTGATGAGGGAGCGACCGTACGCTTGGCGACCTTCTCTGATTG
CGTAGAGTCGTTTTCCTTATTGGGCTTACCTAGTACCGTCAGCCCACAATGTTTTCATTTCTTTCCCACCTAAAACTAAA
AGTACTCACAAGTCACAACTTAAACCAAGTACACAAGGATTTTATCATGGGATTATCGTGTTTGAAGACTAAAAAGAGCA
CACCATCACCCCCATTAGTGCAGGTAGAGTAAGACAGTAACTTTTGGGTTCATATTACCGAGCAAGAACCGTTATTTGTG
ATTAGACATGTTATAAACCACTGCTTTAGTGACTATTTAAAACAATATATTACATGTCGTAATCATGCAACCTAACTATG
TTTTCATTAGTCAAATACAAAGAATAAAGAGAAAAGTGCGTAGATTCAATTATTTGGCATAGACTCAAAAGAGTGTATAT
ATATCTGACTTTTATTAAATTATTAAACACAAATACATATTTTCATAAGCAAAACTATAAAAGCCCTAAACATATAATGA
TTACCTCAAAGGAAAAAGTCGTTTTCTCCTAATTAAAAGATAGGTTACTTCCTAATTAATAGTATATAATTTATGTGAAC
TTCACAATATACAGTTCAATAAAATTTGGTAATTTGACCGATTTAAGGAGAGTGGAAATTAGGGCTTCTGCAATCTTTTT
TCTTCGCCGCAATCTCATGTCCAATTATCCACCGACGGTGGCGGCGCAACCCACAACGACGGCAAATCCACTGCTGCAGC
GACATCAATCTGAACAGCGACGAAGAGAATTACCGAAGATTGTCGAAACAGAGTCTACAAGTATGGACATTACGATCGGT
CAATCTAAGCAGCCTCAATTTTTGAAATCCATAGACGAATTAGCTGCGTTTTCAGTTGCAGTGGAAACATTCAAACGCCA
ATTCGATGATCTTCAGAAGCACATCGAGTCAATCGAAAACGCAATTGATTCCAAACTCGAGAGTAACGGCGTTGTCCTCG
CCGCGCGGAACAATAATTTCCATCAGCCGATGTTATCGCCTCCGCAGAACAATGTATCTGTAGAAACCACCGTCACTGTG
AGCCAACCGTCTCAGGAGATTGTACCGGAGACGTCGAATAAACCGGAGGGGGCACGTATGTGTGAGTTGATGTGTAGCAA
AGGTCTGCGTAAATACATATACGCGAATATCTCTGATCAAGCTAAGTTAATGGAAGAGATTCCTTCAGCTTTGAAATTGG
CCAAGGAGCCAGCGAAGTTTGTATTGGATTGTATTGGCAAGTTTTACTTACAAGGGCGTAGAGCATTTACTAAAGAGTCG
CCTATGAGCTCTGCGAGACAAGTTTCGCTTCTTATACTGGAGTCTTTTCTTCTAATGCCTGATCGTGGTAAAGGGAAGGT
GAAGATTGAGAGTTGGATTAAAGATGAGGCGGAGACGGCTGCTGTTGCTTGGAGGAAAAGGTTGATGACTGAAGGAGGAT
TAGCTGCGGCTGAGAAAATGGATGCAAGGGGTTTGCTTTTACTAGTTGCTTGTTTTGGTGTTCCTTCAAACTTTAGGAGT
ACAGATTTGCTGGATTTGATAAGGATGAGTGGTTCGAATGAGATTGCCGGTGCTTTGAAGCGGTCACAGTTTCTTGTCCC
TATGGTCTCAGGTACCATATTCTGTTCTCACTCGGTGAATTTCATTGCAAAGGTGGTTCCTTTTGTTGACATCATCGACC
AACATCAAGTTCCATCTTTGTTTTTCGATAAGCTTGATGGTATAAACTAGGAGAGCACATCAAATATTTAGAGTGCAATG
ACTGATTGAGCCAAATCCTAGCTAGAAATTAATCTGGAAAGAACTTGGAACTCTCAACCATAGGTTTTGGTACGAAATTG
TTGCTTGTCAGAACCAAATGATAGGCTATTGCCTTGAAATAGTGTTTCTTGTGGTTTCCAATATTGGAAGTTAAAATCAT
ATGACTTAGCTGTTGGATACTAATTAAGCTTAAGCAATGCCAACTCTAAGAAGTGGTACTTACACAATATTCTATTGGTC
ATAGGTATAGTTGAATCAAGTATCAAGCGTGGAATGCATATTGAAGCTCTTGAGATGGTTTATACCTTTGGCATGGAGGA
TAAGTTTTCAGCTGCTCTAGTTCTAACTTCATTCTTAAAGATGAGCAAGGAATCATTTGAGAGGGCAAAACGGAAAGCCC
AGTCACCGCTGGCATTTGTATGAACCCTTCCCTTGCACATTATGTACCTTTATGAACTCTTTATCATCATCTGAGTCTGA
CCATTGATATATTTATTTCTCAACAGAAAGAAGCGGCTACAAAGCAGCTAGCTGTGTTATCATCAGTTATGCAGTGTATG
GAGACTCACAAGTTAGATCCTGCGAAAGAACTACCAGGATGGCAGATCAAAGAGCAAATTGTTAGCTTGGAGAAAGACAC
TCTTCAGCTCGACAAAGAGATGGAAGAGAAAGCAAGATCTCTCAGTTTAATGGAGGAAGCCGCACTTGCCAAGAGAATGT
ATAACCAACAGATAAAACGTCCAAGGTTGTCACCCATGGAAATGCCACCAGTAGCTTCTTCATCGTATTCTCCTATCTAC
CGTGATAGAAGCTTTCCTAGTCAAAGAGACGATGACCAAGATGAAATATCAGCTCTTGTGAGTAGTTACCTCGGCCCGTC
AACATCTTTTCCTCATCGCTCAAGAAGATCCCCGGAATATATGGTTCCACTTCCACATGGTGGGTTAGGAAGAAGTGTAT
ATGCATATGAACATCTGGCCCCAAATTCATACTCTCCAGGTCACGGACATAGACTTCATCGACAGTACTCTCCGTCTTTG
GTTCACGGACAGAGACATCCACTACAGTACTCTCCTCCAATTCATGGACAACAACAGTTACCATATTGTATACAAAGGGT
TTACAGACATTCACCATCTGAAGAAAGATATTTGGGTTTATCCAATCAAAGGTCTCCTCGCAGTAACTCATCATTAGACC
CCAAATAGGAGGAATGTAAATTTGTAACAAAGCTTTTTGTTTTTGCTTAAGTTAGTCATTTATTTAACTCCCAACAGTCT
CAAAATTTAATTTAATGTTTGGGGCTTAAGAATGCAAATTTTTTTGCTCCTGTAATTGACATTTAAGATGCTAATGTTAT
TGCTTCAGAGGTTTTAGTCAACCTCAGATACATCGATATCACTATCTAAATAGACCTCTGGCTCTTGGTCATCTGGATTC
TCTTCATCTTCTGTCTCTGTTCCTTCTTGTTCTCGTTGCACTGCTCGAGCAATTGCGGATTCCAACCTTGTGCTTACAGT
TTCCCATGACACAAGCTTTTCCATGAATGTATTTATGTCCGCCTTCTTATCTTTCTTGAGGAAGATGAATTC

>FRI_Sha

GGATCCCAAAATCTAGTGCCGCCGAGTCCTTTGGGACAGAGTTTCGCACTTATCATTCGATTCGGCAAGCAGTAACTCGCAAAA
TCCGCACCACCGTGAAACATACTCTGGCGCTGAACGCTAGACGCAAAATCCATATTACTGCTGCAGAAAGACGGAGAGAACAA
GAGACTGACCGATCATAAGAGAAGAGAGCTTCAAGGCTTCAGCTGATGAAGAATCAATGGCGGCGAGAGTGAGTGAGAATC
GAGTAGTTACTGAGAGGAAGATGAAACGTATCTCCTGAGAAAAAACGTAAGAGAGTTCCGTTTCAACAGTATTTTTCTCGGCG
GTCTTCAGATTTTCCGTCCAACTCTCTCTTTCCTCTTTCAGTTTTTTTTTGCCCAATCACTCGTGGTAGGATTTTTTTATAAGAAAG
TTACGAAAATGCCCTCTCGTGCTTAGAGAGATTCAATTCGTGCACATATAATATGCCGTGAATATACCACCAACTCTCTTACTCG
TACTCGCATGTGGTGCGTGCAATATACACGTGGTTCGAATCAACGGCTACGTTTTACTGATAGGTGGCTCGTTTAGTTTTGCAC
TGCCTTTGGGTGCGCGTGACCAAGTCATTTTCCAAATGTGTTTTAATATTTGTGAACTATTATTAATAAAAAAGCGAGCGACGTT
GTTCTTTGATGAGGGAACGACCGTACGCTTGGCGACCTTCTCTGATTGCGTAGAGTCGTTTTCCTTATTGGGCTTACCTAGTACC
GTCAGCCCACAATGTTTTCATTTCTTTCCCACCTAAAACTAAAAGTACTCACAAGTCACAACTTAAACCAAGTACACAAGGATTT
TATCATGGGATTATCGTGTTTGAAGACTAAAAAGAGCACACCATCACCCCCATTAGTGCAGGTAGAGTAAGACAGTAACTTTT
GGGTTCATATTACCGAGCAAGAACCGTTATTTGTGATTAGACATGTTATAAACCACTGCTTTAGTGACTATTTAAAACAATATAT
TACATGTCGTAATCATGCAACCTAACTATGTTTTCATTAATCAAATACAAAGAATAAAGAGAAAAGTGCGTAGATTCAATTATTT
GGCATAGACTCAAAAGAGTGTATATATATCTGACTTTTATTAAATTATTAAACACAAATACATATTTTCATAAGCAAAACTATAA
AAGCCCTAAAGATATAACGATTACCTCAAAGGAAAAAGTCGTTTTCTCCTAATTAAAAGATAGGTTACTTCCTAATTAATATATA
ATTTATGTGAACTTCACAATATACAGTTCAATAAAATTTGGTAATTTGACCGATTTAAGGAGAGTGGAAATTTGGGCTTCTGCA
ATCTTTTTTCTTCGCCGCAATCTCATGTCCAATTATCCACCGACGGTGGCGGCGCAACCCACAACGACGGCGAATCCACTGCTG
CAGCGACATCAATCTGAACAGCGACGAAGAGAATTACCGAAGATTGTCGAAACAGAGTCTACAAGTATGGACATTACGATCG
GTCAATCTAAGCAGCCTCAAATTTTGAAATCCATAGACGAATTAGCTGCGTTTTCAGTTGCAGTGGAAACATTCAAACGCCAAT
TCGATGATCTTCAGAAGCACATCGAGTCAATCGAAAACGCAATTGATTCCAAACTCGAGAGTAACGGCGTTGTCCTCGCCGCG
CGGAACAATAATTTCCATCAGCCGATGTTATCGCCTCCGCGGAACAATGTATCTGTAGAAACCACCGTCACTGTGAGCCAACCG
TCTCAGGAGATTGTACCGGAGACGTCGAATAAACCGGAGGGGGGACGTATGTGTGAGTTGATGTGTAGCAAAGGTCTGCGT
AAATACATATACGCGAATATCTCTGATCAAGCTAAGTTAATGGAAGAGATTCCTTCAGCTTTGAAATTGGCCAAGGAGCC
AGCGAAGTTTGTATTGGATTGTATTGGCAAGTTTTACTTACAAGGGCGTAGAGCATTTACTAAAGAGTCGCCTATGAGCT
CTGCGAGACAAGTTTCGCTTCTTATACTGGAGTCTTTTCTTCTAATGCCTGATCGTGGTAAAGGGAAGGTGAAGATTGAG
AGTTGGATTAAAGATGAGGCGGAGACGGCTGCTGTTGCTTGGAGGAAAAGGTTGATGACTGAAGGAGGATTAGCTGCGGC
TGAGAAAATGGATGCAAGGGGTTTGCTTTTACTAGTTGCTTGTTTTGGTGTTCCTTCAAACTTTAGGAGTACAGATTTGC
TGGATTTGATAAGGATGAGTGGTTCGAATGAGATTGCCGGTGCTTTGAAGCGGTCACAGTTTCTTGTCCCTATGGTCTCA
GGTACCATATTCTGTTCTCACTCGGTGAATTTCATTGCAAAGGTGGTTCCTTTTGTTGACATCATCGACCAACATCAAGT
TCCATCTTTGTTTTTCGATAAGCTTGATGGTATAAACTAGGAGAGCACATCAAATATTTAGAGTGCAATGACTGATTGAG
CCAAATCCTAGCTAGAAATTAATCTGGAAAGAACTTGGAACTCTCAACCATAGGTTTTGGTACGAAATTGTTGCTTGTCA
GAACCAAATGATAGGCTATTGCCTTGAAATAGTGTTTCTTGTGGTTTCCAATATTGGAAGTTAAAATCATATGACTTAGC
TGTTGGATACTAATTAAGCTTAAGCAATGCCAACTCTAAGAAGTGGTACTTACACAATATTCTATTGGTCATAGGTATAG
TTGAATCAAGTATCAAGCGTGGAATGCATATTGAAGCTCTTGAGATGGTTTATACCTTTGGCATGGAGGATAAGTTTTCA
GCTGCTCTAGTTCTAACTTCATTCTTAAAGATGAGCAAGGAGTCATTTGAGAGGGCAAAACGGAAAGCCCAGTCACCGCT
GGCATTTGTATGAACCCTTCCCTTGCACATTATGTACCTTTATGAACTCTTTATCATCATCTGAGTCTGACCATTGATAT
ATTTATTTCTCAACAGAAAGAAGCGGCTACAAAGCAGCTAGCTGTGTTATCATCAGTTATGCAGTGTATGGAGACTCACA
AGTTAGATCCTGCGAAAGAACTACCAGGATGGCAGATCAAAGAGCAAATTGTTAGCTTGGAGAAAGACACTCTTCAGCTC
GACAAAGAGATGGAAGAGAAAGCAAGATCTCTCAGTTTAATGGAGGAAGCCGCACTTGCCAAGAGAATGTATAACCAACA
GATAAAACGTCCAAGGTTGTCACCCATGGAAATGCCACCAGTAACTTCTTCATCGTATTCTCCTATCTACCGTGATAGAA
GCTTTCCTAGTCAAAGAGACGATGACCAAGATGAAATATCAGCTCTTGTGAGTAGTTACCTCGGCCCGTCAACATCTTTT
CCTCATCGCTCAAGAAGATCCCCGGAATATATGGTTCCACTTCCACATGGTGGGTTAGGAAGAAGTGTATATGCATATGA
ACATCTGGCCCCAAATTCATACTCTCCAGGTCACGGACATAGACTTCATCGACAGTACTCTCCGTCTTTGGTTCACGGAC
AGAGACATCCACTACAGTACTCTCCTCCAATTCATGGACAACAACAGTTACCATATGGTATACAAAGGGTTTACAGACAT
TCACCATCTGAAGAAAGATATTTGGGTTTATCCAATCAAAGGTCTCCTCGCAGTAACTCATCATTAGACCCCAAATAGGA
GGAATGTAAATTTGTAACAAAGCTTTTTGTTTTTGCTTAAGTTAGTCATTTATTTAACTCCCAACAGTCTCAAAATTTAA
TTTAATGTTTGGGGCTTAAGAATGCAAATTTTTTTGCTCCTGTAATTGACATTTAAGATGCTAATGTTATTGCTTCAGAG
GTTTTAGTCAACCTCAGATACATCGATATCACTATCTAAATAGACCTCTGGCTCTTGGTCATCTGGATTCTCTTCATCTT
CTGTCTCTGTTCCTTCTTGTTCTCGTTGCACTGCTCGAGCAATTGCGGATTCCAACCTTGTGCTTACAGTTTCCCATGAC
ACAAGCTTTTCCATGAATGTATTTATGTCCGCCTTCTTATCTTTCTTGAGGAAGATGAATTC

>FRI_Van-0
GGATCCCAAAATCTAGTGCCGCCGAGTCCTTTGGGACACAGTTTCGCACTTATCATTCGATTCGGCAAGCAGTAACTCGC

AAAATCCGCACCACCGTGAAACATACTCCGGCGCTGAACGCTAGACGCAAAATCCATATTACAGCTGCAGAAAGACGGAG
AGAACAAGAGACTGACCGATCATAAGAGAAGAGAGCTTCAAGGCTTCAGCTGATGAAGAATCAATGGCGGCGAGAGAGAG
TGAGAATCGAGTAGTTACTGAGAGGAAGATGAAACGTATCTCCTGAGAAAAAACGTAAGAGAGTTCCGTTTCAACAGTAT
TTTTCTCGGCGGTCTTCAGATTTTCCGTCCAACTCTCTCTTTCCTCTTTCAGTTTTTTTTGCCCAATCACTCGTGGTAGG
ATTTTTTATAAGAAAGTTACGAAAATGCCCTCTCGTGCTTAGAGAGATTCAATTCGTGCACATATAATATGCCGTGAATA
TACCACCAACTCTCTTACTCGTACTCGCATGTGGTGCGTGCAATATACACGTGGTTCGAATCAACGGCTACGTTTTACTG
ATAGGTGGCTCGTTTAGTTTTGCACTGCCTTTGGGTGCGCGTGACCAAGTCATTTTCCAAATGTGTTTTAATATTTGTGA
ACTATTATTAATAAAAAAGCGAGCGACGTTGTTCTTTGATGAGGGAACGACCGTACGCTTGGCGACCTTCTCTGATTGCG
TAGAGTCGTTTTCCTTATTGGGCTTACCTAGTACCGTCAGCCCACAATGTTTTCATTTCTTTCCCACCTAAAACTAAAAG
TACTCACAAGTCACAACTTAAACCAAGTACACAAGGATTTTATCATGGGATTATCGTGTTTGAAGACTAAAAAGAGCACA
CCATCACCCCCATTAGTGCAGGTAGAGTAAGACAGTAACTTTTGGGTTCATATTACCGAGCAAGAACCGTTATTTGTGAT
TAGACATGTTATAAACCACTGCTTTAGTGACTATTTAAAACAATATATTACATGTCGTAATCATGCAACCTAACTATGTT
TTCATTAATCAAATACAAAGAATAAAGAGAAAAGTGCGTAGATTCAATTATTTGGCATAGACTCAAAAGAGTGTATATAT
ATCTGACTTTTATTAAATTATTAAACACAAATACATATTTTCATAAGCAAAACTATAAAAGCCCTAAACATATAATGATT
ACCTCAAAGGAAAAAGTCGTTTTCTCCTAATTAAAAGATAGGTTACTTCCTAATTAATATATAATTTATGTGAACTTCAC
AATATACAGTTCAATAAAATTTGGTAATTTGACCGATTTAAGGAGAGTGGAAATTAGGGCTTCTGCAATCTTTTTTCTTC
GCCGCAATCTCATGTCCAATTATCCACCGACGGTGGCGGCGCAACCCACAACGACGGCGAATCCACTGCTGCAGCGACAT
CAATCTGAACAGCGACGAAGAGAATTACCGAAGATTGTCGAAACAGAGTCTACAAGTATGGACATTACGATCGGTCAATC
TAAGCAGCCTCAATTTTTGAAATCCATAGACGAATTAGCTGCGTTTTCAGTTGCAGTGGAAACATTCAAACGCCAATTCG
ATGATATTCAGAAGCACATCGAGTCAATCGAAAACGCAATTGATTCCAAACTCGAGAGTAACGGCGTTGTCCTCGCCGCG
CGGAACAATAATTTCCATCAGCCGATGTTATCGCCTCCGCGGAACAATGTATCTGTAGAAACCACCGTCACTGTGAGCCA
ACCGTCTCAGGAGATTGTACCGGAGACGTGGAATAAACCGGAGGGGGGACGTATGTGTGAGTTGATGTGTAGCAAAGGTC
TGCGTAAATACATATACGCGAATATCTCTGATCAAGCTAAGTTAATGGAAGAGATTCCTTCAGCTTTGAAATTGGCCAAG
GAGCCAGCGAAGTTTGTATTGGATTGTATTGGCAAGTTTTACTTACAAGGGCGTAGAGCATTTACTAAAGAGTCGCCTAT
GAGCTCTGCGAGACAAGTTTCGCTTCTTATACTGGAGTCTTTTCTTCTAATGCCTGATCGTGGTAAAGGGAAGGTGAAGA
TTGAGAGTTGGATTAAAGATGAGGCGGAGACGGCTGCTGTTGCTTGGAGGAAAAGGTTGATGACTGAAGGAGGATTAGCT
GCGGCTGAGAAAATGGATGCAAGGGGTTTGCTTTTACTAGTTGCTTGTTTTGGTGTTCCTTCAAACTTTAGGAGTACAGA
TTTGCTGGATTTGATAAGGATGAGTGGTTCGAATGAGATTGCCGGTGCTTTGAAGCGGTCACAGTTTCTTGTCCCTATGG
TCTCAGGTACCATATTCTGTtCTCACTCGGTGAATTTCATTGCAAAGGTGGTTCCTTTTGTTGACATCATCGACCAACAT
CAAGTTCCATCTTTGTTTTTCGATAAGCTTGATGGTATAAACTAGGAGAGCACATCAAATATTTAGAGTGCAATGACTGA
TTGAGCCAAATCCTAGCTAGAAATTAATCTGGAAAGAACTTGGAACTCTCAACCATAGGTTTTGGTACGAAATTGTTGCT
TGTCAGAACCAAATGATAGGCTATTGCCTTGAAATAGTGTTTCTTGTGGTTTCCAATATTGGAAGTTAAAATCATATGAC
TTAGCTGTTGGATACTAATTAAGCTTAAGCAATGCCAACTCTAAGAAGTGGTACTTACACAATATTCTATTGGTCATAGG
TATAGTTGAATCAAGTATCAAGCGTGGAATGCATATTGAAGCTCTTGAGATGGTTTATACCTTTGGCATGGAGGATAAGT
TTTCAGCTGCTCTAGTTCTAACTTCATTCTTAAAGATGAGCAAGGAGTCATTTGAGAGGGCAAAACGGAAAGCCCAGTCA
CCGCTGGCATTTGTATGAACCCTTCCCTTGCACATTATGTACCTTTATGAACTCTTTATCATCATCTGAGTCTGACCATT
GATATATTTATTTCTCAACAGAAAGAAGCGGCTACAAAGCAGCTAGCTGTGTTATCATCAGTTATGCAGTGTATGGAGAC
TCACAAGTTAGATCCTGCGAAAGAACTACCAGGATGGCAGATCAAAGAGCAAATTGTTAGCTTGGAGAAAGACACTCTTC
AGCTCGACAAAGAGATGGAAGAGAAAGCAAGATCTCTCAGTTTAATGGAGGAAGCCGCACTTGCCAAGAGAATGTATAAC
CAACAGATAAAACGTCCAAGGTTGTCACCCATGGAAATGCCACCAGTAACTTCTTCATCATATTCTCCTATCTACCGTGA
TAGAAGCTTTCCTAGTCAAAGAGACGATGACCAAGATGAAATATCAGCTCTTGTGAGTAGTTACCTCGGCCCGTCAACAT
CTTTTCCTCATCGCTCAAGAAGATCCCCGGAATATATGGTTCCACTTCCACATGGTGGGTTAGGAAGAAGTGTATATGCA
TATGAACATCTGGCCCCAAATTCATACTCTCCAGGTCACGGACATAGACTTCATCGACAGTACTCTCCGTCTTTGGTTCA
CGGACAGAGACATCCACTACAGTACTCTCCTCCAATTCATGGACAACAACAGTTACCATATGGTATACAAAGGGTTTACA
GACATTCACCATCTGAAGAAAGATATTTGGGTTTATCCAATCAAAGGTCTCCTCGCAGTAACTCATCATTAGACCCCAAA
TAGGAGGAATGTAAATTTGTAACAAAGCTTTTTGTTTTTGCTTAAGTTAGTCATTTATTTAACTCCCAACAGTCTCAAAA
TTTAATTTAATGTTTGGGGCTTAAGAATGCAAATTTTTTTGCTCCTGTAATTGACATTTAAGATGCTAATGTTATTGCTT
CAGAGGTTTTAGTCAACCTCAGATACATCGATATCACTATCTAAATAGACCTCTGGCTCTTGGTCATCTGGATTCTCTTC
ATCTTCTGTCTCTGTTCCTTCTTGTTCTCGTTGCACTGCTCGAGCAATTGCGGATTCCAACCTTGTGCTTACAGTTTCCC
ATGACACAAGCTTTTCCATGAATGTATTTATGTCCGCCTTCTTATCTTTCTTGAGGAAGATGAATTC
