## Supplemental Material 4 for "Functional Analysis of FRIGIDA Using Naturally Occurring Variation in *Arabidopsis thaliana*"

**Supplementary Material 4.** ClustalW alignment of *FRI* alleles cloned and sequenced in this work.

CLUSTAL O(1.2.4) multiple sequence alignment

```
FRI_An-1         GGATCCCAAAATCTAGTGCCGCCGAGTCCTTTGGGACAGAGTTTCGCACTTATCATTCGA         60
FRI_Pro-0        GGATCCCAAAATCTAGTGCCGCCGAGTCCTTTGGGACACAGTTTCGCACTTATCATTCGA         60
FRI_Bål-2        GGATCCCAAAATCTAGTGCCGCCGAGTCCTTTGGGACAGAGTTTCGCACTTATCATTCGA         60
FRI_St-0         GGATCCCAAAATCTAGTGCCGCCGAGTCCTTTGGGACACAGTTTCGCACTTATCATTCGA         60
FRI_Bg-2         GGATCCCAAAATCTAGTGCCGCCGAGTCCTTTGGGACACAGTTTCGCACTTATCATTCGA         60
FRI_C24          GGATCCCAAAATCTAGTGCCGCCGAGTCCTTTGGGACACAGTTTCGCACTTATCATTCGA         60
FRI_Van-0        GGATCCCAAAATCTAGTGCCGCCGAGTCCTTTGGGACACAGTTTCGCACTTATCATTCGA         60
FRI_Ull-2-3      GGATCCCAAAATCTAGTGCCGCCGAGTCCTTTGGGACACAGTTTCGCACTTATCATTCGA         60
FRI_Lip-0        GGATCCCAAAATCTAGTGCCGCCGAGTCCTTTGGGACACAGTTTCGCACTTATCATTCGA         60
FRI_NFA-8        GGATCCCAAAATCTAGTGCCGCCGAGTCCTTTGGGACAGAGTTTCGCACTTATCATTCGA         60
FRI_Edi-0        GGATCCCAAAATCTAGTGCCGCCGAGTCCTTTGGGACAGAGTTTCGCACTTATCATTCGA         60
FRI_Ren-1        GGATCCCAAAATCTAGTGCCGCCGAGTCCTTTGGGACAGAGTTTCGCACTTATCATTCGA         60
FRI_Nok-3        GGATCCCAAAATCTAGTGCCGCCGAGTCCTTTGGGACAGAGTTTCGCACTTATCATTCGA         60
FRI_Sha          GGATCCCAAAATCTAGTGCCGCCGAGTCCTTTGGGACAGAGTTTCGCACTTATCATTCGA         60
FRI_Wa-1         GGATCCCAAAATCTAGTGCCGCCGAGTCCTTTGGGACACAGTTTCGCACTTATCATTCGA         60
FRI_Spr-1-6      GGATCCCAAAATCTAGTGCCGCCGAGTCCTTTGGGACACAGTTTCGCACTTATCATTCGA         60
FRI_Bil-7        GGATCCCAAAATCTAGTGCCGCCGAGTCCTTTGGGACACAGTTTCGCACTTATCATTCGA         60
FRI_Alc-0        GGATCCCAAAATCTAGTGCCGCCGAGTCCTTTGGGACAGAGTTTCGCACTTATCATTCGA         60
FRI_Pu-2-23      GGATCCCAAAATCTAGTGCCGCCGAGTCCTTTGGGACAGAGTTTCGCACTTATCATTCGA         60
FRI_NFA-10       GGATCCCAAAATCTAGTGCCGCCGAGTCCTTTGGGACACAGTTTCGCACTTATCATTCGA         60
FRI_Cvi-0        GGATCCCAAAATCTAGTGCCGCCGAGTCCTTTGGGACAGAGTTTCGCACTTATCATTCGA         60
FRI_Zdr-6        GGATCCCAAAATCTAGTGCCGCCGAGTCCTTTGGGACACAGTTTCGCACTTATCATTCGA         60
                 ************************************** *********************

FRI_An-1         TTCGGCAAGCAGTAACTCGCAAAATCCGCACCACCGTGAAACATACTCTGGCGCTGAACG         120
FRI_Pro-0        TTCGGCAAGCAGTAACTCGCAAAATCCGCACCACCGTGAAACATACTCCGGCGCTGAACG         120
FRI_Bål-2        TTCGGCAAGCAGTAACTCGCAAAATCCGCACCACCGTGAAACATACTCTGGCGCTGAACG         120
FRI_St-0         TTCGGCAAGCAGTAACTCGCAAAATCCGCACCACCGTGAAACATACTCCGGCGCTGAACG         120
FRI_Bg-2         TTCGGCAAGCAGTAACTCGCAAAATCCGCACCACCGTGAAACATACTCCGGCGCTGAACG         120
FRI_C24          TTCGGCAAGCAGTAACTCGCAAAATCCGCACCACCGTGAAACATACTCCGGCGCTGAACG         120
FRI_Van-0        TTCGGCAAGCAGTAACTCGCAAAATCCGCACCACCGTGAAACATACTCCGGCGCTGAACG         120
FRI_Ull-2-3      TTCGGCAAGCAGTAACTCGCAAAATCCGCACCACCGTGAAACATACTCCGGCGCTGAACG         120
FRI_Lip-0        TTCGGCAAGCAGTAACTCGCAAAATCCGCACCACCGTGAAACATACTCCGGCGCTGAACG         120
FRI_NFA-8        TTCGGCAAGCAGTAACTCGCAAAATCCGCACCACCGTGAAACATACTCTGGCGCTGAACG         120
FRI_Edi-0        TTCGGCAAGCAGTAACTCGCAAAATCCGCACCACCGTGAAACATACTCTGGCGCTGAACG         120
FRI_Ren-1        TTCGGCAAGCAGTAACTCGCAAAATCCGCACCACCGTGAAACATACTCTGGCGCTGAACG         120
FRI_Nok-3        TTCGGCAAGCAGTAACTCGCAAAATCCGCACCACCGTGAAACATACTCTGGCGCTGAACG         120
FRI_Sha          TTCGGCAAGCAGTAACTCGCAAAATCCGCACCACCGTGAAACATACTCTGGCGCTGAACG         120
FRI_Wa-1         TTCGGCAAGCAGTAACTCGCAAAATCCGCACCACCGTGAAACATACTCCGGCGCTGAACG         120
FRI_Spr-1-6      TTCGGCAAGCAGTAACTCGCAAAATCCGCACCACCGTGAAACATACTCCGGCGCTGAACG         120
FRI_Bil-7        TTCGGCAAGCAGTAACTCGCAAAATCCGCACCACCGTGAAACATACTCCGGCGCTGAACG         120
FRI_Alc-0        TTCGGCAAGCAGTAACTCGCAAAATCCGCACCACCGTGAAACATACTCTGGCGCTGAACG         120
FRI_Pu-2-23      TTCGGCAAGCAGTAACTCGCAAAATCCGCACCACCGTGAAACATACTCTGGCGCTGAACG         120
FRI_NFA-10       TTCGGCAAGCAGTAACTCGCAAAATCCGCACCACCGTGAAACATACTCCGGCGCTGAACG         120
FRI_Cvi-0        TTCGGCAAGCAGTAACTCGCAAAATCCGCACCACCGTGAAACATACTCTGGCGCTGAACG         120
FRI_Zdr-6        TTCGGCAAGCAGTAACTCGCAAAATCCGCACCACCGTGAAACATACTCCGGCGCTGAACG         120
                 ************************************************ ***********

FRI_An-1         CTAGACGCAAAATCCATATTACTGCTGCAGAAAGACGGAGAGAACAAGAGACTGACCGAT         180
FRI_Pro-0        CTAGACGCAAAATCCATATTACAGCTGCAGAAAGACGGAGAGAACAAGAGACTGACCGAT         180
FRI_Bål-2        CTAGACGCAAAATCCATATTACTGCTGCAGAAAGACGGAGAGAACAAGAGACTGACCGAT         180
FRI_St-0         CTAGACGCAAAATCCATATTACAGCTGCAGAAAGACGGAGAGAACAAGAGACTGACCGAT         180
FRI_Bg-2         CTAGACGCAAAATCCATATTACAGCTGCAGAAAGACGGAGAGAACAAGAGACTGACCGAT         180
FRI_C24          CTAGACGCAAAATCCATATTACTGCTGCAGAAAGACGGAGAGAACAAGAGACTGACCGAT         180
FRI_Van-0        CTAGACGCAAAATCCATATTACAGCTGCAGAAAGACGGAGAGAACAAGAGACTGACCGAT         180
FRI_Ull-2-3      CTAGACGCAAAATCCATATTACAGCTGCAGAAAGACGGAGAGAACAAGAGACTGACCGAT         180
FRI_Lip-0        CTAGACGCAAAATCCATATTACAGCTGCAGAAAGACGGAGAGAACAAGAGACTGACCGAT         180
FRI_NFA-8        CTAGACGCAAAATCCATATTACTGCTGCAGAAAGACGGAGAGAACAAGAGACTGACCGAT         180
FRI_Edi-0        CTAGACGCAAAATCCATATTACTGCTGCAGAAAGACGGAGAGAACAAGAGACTGACCGAT         180
FRI_Ren-1        CTAGACGCAAAATCCATATTACTGCTGCAGAAAGACGGAGAGAACAAGAGACTGACCGAT         180
FRI_Nok-3        CTAGACGCAAAATCCATATTACTGCTGCAGAAAGACGGAGAGAACAAGAGACTGACCGAT         180
FRI_Sha          CTAGACGCAAAATCCATATTACTGCTGCAGAAAGACGGAGAGAACAAGAGACTGACCGAT         180
FRI_Wa-1         CTAGACGCAAAATCCATATTACAGCTGCAGAAAGACGGAGAGAACAAGAGACTGACCGAT         180
FRI_Spr-1-6      CTAGACGCAAAATCCATATTACAGCTGCAGAAAGACGGAGAGAACAAGAGACTGACCGAT         180
FRI_Bil-7        CTAGACGCAAAATCCATATTACTGCTGCAGAAAGACGGAGAGAACAAGAGACTGACCGAT         180
FRI_Alc-0        CTAGACGCAAAATCCATATTACTGCTGCAGAAAGACGGAGAGAACAAGAGACTGACCGAT         180
```

```
FRI_Pu-2-23      CTAGACGCAAAATCCATATTACTGCTGCAGAAAGACGGAGAGAACAAGAGACTGACCGAT      180
FRI_NFA-10       CTAGACGCAAAATCCATATTACAGCTGCAGAAAGACGGAGAGAACAAGAGACTGACCGAT      180
FRI_Cvi-0        CTAGACGCAAAATCCATATTACTGCTGCAGAAAGACGGAGAGAACAAGAGACTGACCGAT      180
FRI_Zdr-6        CTAGACGCAAAATCCATATTACTGCTGCAGAAAGACGGAGAGAACAAGAGACTGACCGAT      180
                 ********************** *************************************

FRI_An-1         CATAAGAGAAGAGAGCTTCAAGGCTTCAGCTGATGAAGAATCAATGGCGGCGAGAGTGAG      240
FRI_Pro-0        CATAAGAGAAGAGAGCTTCAAGGCTTCAGCTGATGAAGAATCAATGGCGGCGAGAGAGAG      240
FRI_Bå1-2        CATAAGAGAAGAGAGCTTCAAGGCTTCAGCTGATGAAGAATCAATGGCGGCGAGAGTGAG      240
FRI_St-0         CATAAGAGAAGAGAGCTTCAAGGCTTCAGCTGATGAAGAATCAATGGCGGCGAGAGAGAG      240
FRI_Bg-2         CATAAGAGAAGAGAGCTTCAAGGCTTCAGCTGATGAAGAATCAATGGCGGCGAGAGAGAG      240
FRI_C24          CATAAGAGAAGAGAGCTTCAAGGCTTCAGCTGATGAAGAATCAATGGCGGCGAGAGAGAG      240
FRI_Van-0        CATAAGAGAAGAGAGCTTCAAGGCTTCAGCTGATGAAGAATCAATGGCGGCGAGAGAGAG      240
FRI_Ull-2-3      CATAAGAGAAGAGAGCTTCAAGGCTTCAGCTGATGAAGAATCAATGGCGGCGAGAGAGAG      240
FRI_Lip-0        CATAAGAGAAGAGAGCTTCAAGGCTTCAGCTGATGAAGAATCAATGGCGGCGAGAGAGAG      240
FRI_NFA-8        CATAAGAGAAGAGAGCTTCAAGGCTTCAGCTGATGAAGAATCAATGGCGGCGAGAGTGAG      240
FRI_Edi-0        CATAAGAGAAGAGAGCTTCAAGGCTTCAGCTGATGAAGAATCAATGGCGGCGAGAGTGAG      240
FRI_Ren-1        CATAAGAGAAGAGAGCTTCAAGGCTTCAGCTGATGAAGAATCAATGGCGGCGAGAGTGAG      240
FRI_Nok-3        CATAAGAGAAGAGAGCTTCAAGGCTTCAGCTGATGAAGAATCAATGGCGGCGAGAGTGAG      240
FRI_Sha          CATAAGAGAAGAGAGCTTCAAGGCTTCAGCTGATGAAGAATCAATGGCGGCGAGAGTGAG      240
FRI_Wa-1         CATAAGAGAAGAGAGCTTCAAGGCTTCAGCTGATGAAGAATCAATGGCGGCGAGAGTGAG      240
FRI_Spr-1-6      CATAAGAGAAGAGAGCTTCAAGGCTTCAGCTGATGAAGAATCAATGGCGGCGAGAGTGAG      240
FRI_Bil-7        CATAAGAGAAGAGAGCTTCAAGGCTTCAGCTGATGAAGAATCAATGGCGGCGAGAGAGAG      240
FRI_Alc-0        CATAAGAGAAGAGAGCTTCAAGGCTTCAGCTGATGAAGAATCAATGGCGGCGAGAGAGAG      240
FRI_Pu-2-23      CATAAGAGAAGAGAGCTTCAAGGCTTCAGCTGATGAAGAATCAATGGCGGCGAGAGAGAG      240
FRI_NFA-10       CATAAGAGAAGAGAGCTTCAAGGCTTCAGCTGATGAAGAATCAATGGCGGCGAGAGAGAG      240
FRI_Cvi-0        CATAAGAGAAGAGAGCTTCAAGGCTTCAGCTGATGAAGAATCAATGGCGGCGAGAGAGAG      240
FRI_Zdr-6        CATAAGAGAAGAGAGCTTCAAGGCTTCAGCTGATGAAGAATCAATGGCGGCGAGAGAGAG      240
                 ******************************************************** ***

FRI_An-1         TGAGAATCGAGTAGTTACTGAGAGGAAGATGAAACGTATCTCCTGAGAAAAAACGTAAGA      300
FRI_Pro-0        TGAGAATCGAGTAGTTACTGAGAGGAAGATGAAACGTATCTCCTGAGAAAAAACGTAAGA      300
FRI_Bå1-2        TGAGAATCGAGTAGTTACTGAGAGGAAGATGAAACGTATCTCCTGAGAAAAAACGTAAGA      300
FRI_St-0         TGAGAATCGAGTAGTTACTGAGAGGAAGATGAAACGTATCTCCTGAGAAAAAACGTAAGA      300
FRI_Bg-2         TGAGAATCGAGTAGTTACTGAGAGGAAGATGAAACGTATCTCCTGAGAAAAAACGTAAGA      300
FRI_C24          TGAGAATCGAGTAGTTACTGAGAGGAAGATGAAACGTATCTCCTGAGAAAAAACGTAAGA      300
FRI_Van-0        TGAGAATCGAGTAGTTACTGAGAGGAAGATGAAACGTATCTCCTGAGAAAAAACGTAAGA      300
FRI_Ull-2-3      TGAGAATCGAGTAGTTACTGAGAGGAAGATGAAACGTATCTCCTGAGAAAAAACGTAAGA      300
FRI_Lip-0        TGAGAATCGAGTAGTTACTGAGAGGAAGATGAAACGTATCTCCTGAGAAAAAACGTAAGA      300
FRI_NFA-8        TGAGAATCGAGTAGTTACTGAGAGGAAGATGAAACGTATCTCCTGAGAAAAAACGTAAGA      300
FRI_Edi-0        TGAGAATCGAGTAGTTACTGAGAGGAAGATGAAACGTATCTCCTGAGAAAAAACGTAAGA      300
FRI_Ren-1        TGAGAATCGAGTAGTTACTGAGAGGAAGATGAAACGTATCTCCTGAGAAAAAACGTAAGA      300
FRI_Nok-3        TGAGAATCGAGTAGTTACTGAGAGGAAGATGAAACGTATCTCCTGAGAAAAAACGTAAGA      300
FRI_Sha          TGAGAATCGAGTAGTTACTGAGAGGAAGATGAAACGTATCTCCTGAGAAAAAACGTAAGA      300
FRI_Wa-1         TGAGAATCGAGTAGTTACTGAGAGGAAGATGAAACGTATCTCCTGAGAAAAAACGTAAGA      300
FRI_Spr-1-6      TGAGAATCGAGTAGTTACTGAGAGGAAGATGAAACGTATCTCCTGAGAAAAAACGTAAGA      300
FRI_Bil-7        TGAGAATCGAGTAGTTACTGAGAGGAAGATGAAACGTATCTCCTGAGAAAAAACGTAAGA      300
FRI_Alc-0        TGAGAATCGAGTAGTTACTGAGAGGAAGATGAAACGTATCTCCTGAGAAAAAACGTAAGA      300
FRI_Pu-2-23      TGAGAATCGAGTAGTTACTGAGAGGAAGATGAAACGTATCTCCTGAGAAAAAACGTAAGA      300
FRI_NFA-10       TGAGAATCGAGTAGTTACTGAGAGGAAGATGAAACGTATCTCCTGAGAAAAAACGTAAGA      300
FRI_Cvi-0        TGAGAATCGAGTAGTTACTGAGAGGAAGATGAAACGTATCTCCTGAGAAAAAACGTAAGA      300
FRI_Zdr-6        TGAGAATCGAGTAGTTACTGAGAGGAAGATGAAACGTATCTCCTGAGAAAAAACGTAAGA      300
                 ************************************************************

FRI_An-1         GAGTTCCGTTTCAACAGTATTTTTCTCGGCGGTCTTCAGATTTTCCGTCCAACTCTCTCT      360
FRI_Pro-0        GAGTTCCGTTTCAACAGTATTTTTCTCGGCGGTCTTCAGATTTTCCGTCCAACTCTCTCT      360
FRI_Bå1-2        GAGTTCCGTTTCAACAGTATTTTTCTCGGCGGTCTTCAGATTTTCCGTCCAACTCTCTCT      360
FRI_St-0         GAGTTCCGTTTCAACAGTATTTTTCTCGGCGGTCTTCAGATTTTCCGTCCAACTCTCTCT      360
FRI_Bg-2         GAGTTCCGTTTCAACAGTATTTTTCTCGGCGGTCTTCAGATTTTCCGTCCAACTCTCTCT      360
FRI_C24          GAGTTCCGTTTCAACAGTATTTTTCTCGGCGGTCTTCAGATTTTCCGTCCAACTCTCTCT      360
FRI_Van-0        GAGTTCCGTTTCAACAGTATTTTTCTCGGCGGTCTTCAGATTTTCCGTCCAACTCTCTCT      360
FRI_Ull-2-3      GAGTTCCGTTTCAACAGTATTTTTCTCGGCGGTCTTCAGATTTTCCGTCCAACTCTCTCT      360
FRI_Lip-0        GAGTTCCGTTTCAACAGTATTTTTCTCGGCGGTCTTCAGATTTTCCGTCCAACTCTCTCT      360
FRI_NFA-8        GAGTTCCGTTTCAACAGTATTTTTCTCGGCGGTCTTCAGATTTTCCGTCCAACTCTCTCT      360
FRI_Edi-0        GAGTTCCGTTTCAACAGTATTTTTCTCGGCGGTCTTCAGATTTTCCGTCCAACTCTCTCT      360
FRI_Ren-1        GAGTTCCGTTTCAACAGTATTTTTCTCGGCGGTCTTCAGATTTTCCGTCCAACTCTCTCT      360
FRI_Nok-3        GAGTTCCGTTTCAACAGTATTTTTCTCGGCGGTCTTCAGATTTTCCGTCCAACTCTCTCT      360
FRI_Sha          GAGTTCCGTTTCAACAGTATTTTTCTCGGCGGTCTTCAGATTTTCCGTCCAACTCTCTCT      360
FRI_Wa-1         GAGTTCCGTTTCAACAGTATTTTTCTCGGCGGTCTTCAGATTTTCCGTCCAACTCTCTCT      360
FRI_Spr-1-6      GAGTTCCGTTTCAACAGTATTTTTCTCGGCGGTCTTCAGATTTTCCGTCCAACTCTCTCT      360
FRI_Bil-7        GAGTTCCGTTTCAACAGTATTTTTCTCGGCGGTCTTCAGATTTTCCGTCCAACTCTCTCT      360
```

```
FRI_Alc-0       GAGTTCCGTTTCAACAGTATTTTTCTCGGCGGTCTTCAGATTTTCCGTCCAACTCTCTCT      360
FRI_Pu-2-23     GAGTTCCGTTTCAACAGTATTTTTCTCGGCGGTCTTCAGATTTTCCGTCCAACTCTCTCT      360
FRI_NFA-10      GAGTTCCGTTTCAACAGTATTTTTCTCGGCGGTCTTCAGATTTTCCGTCCAACTCTCTCT      360
FRI_Cvi-0       GAGTTCCGTTTCAACAGTATTTTTCTCGGCGGTCTTCAGATTTTCCGTCCAACTCTCTCT      360
FRI_Zdr-6       GAGTTCCGTTTCAACAGTATTTTTCTCGGCGGTCTTCAGATTTTCCGTCCAACTCTCTCT      360
                ************************************************************

FRI_An-1        TTCCTCTTTCAGTTTTTTTTTGCCCAATCACTCGTGGTAGGATTTTTTTATAAGAAAGTT      420
FRI_Pro-0       TTCCTCTTTCAGTTTTTTTTTGCCCAATCACTCGTGGTAGGATTT-TTTATAAGAAAGTT      419
FRI_Bå1-2       TTCCTCTTTCAGTTTTTTTTTGCCCAATCACTCGTGGTAGGATTTTTTTATAAGAAAGTT      420
FRI_St-0        TTCCTCTTTCAGTTTTTTTTGCC-CAATCACTCGTGGTAGG-ATTTTTTATAAGAAAGTT      418
FRI_Bg-2        TTCCTCTTTCAGTTTTTTTTGCC-CAATCACTCGTGGTAGG-ATTTTTTATAAGAAAGTT      418
FRI_C24         TTCCTCTTTCAGTTTTTTTTGCC-CAATCACTCGTGGTAGG-ATTTTTTATAAGAAAGTT      418
FRI_Van-0       TTCCTCTTTCAGTTTTTTTTGCC-CAATCACTCGTGGTAGG-ATTTTTTATAAGAAAGTT      418
FRI_Ull-2-3     TTCCTCTTTCAGTTTTTTTTGC-CCAATCACTCGTGGTAGG-ATTTTTTATAAGAAAGTT      418
FRI_Lip-0       TTCCTCTTTCAGTTTTTTTTGC-CCAATCACTCGTGGTAGG-ATTTTTTATAAGAAAGTT      418
FRI_NFA-8       TTCCTCTTTCAGTTTTTTTTTGCCCAATCACTCGTGGTAGGATTTTTTTATAAGAAAGTT      420
FRI_Edi-0       TTCCTCTTTCAGTTTTTTTTTGCCCAATCACTCGTGGTAGGATTTTTTTATAAGAAAGTT      420
FRI_Ren-1       TTCCTCTTTCAGTTTTTTTTTGCCCAATCACTCGTGGTAGGATTTTTTTATAAGAAAGTT      420
FRI_Nok-3       TTCCTCTTTCAGTTTTTTTTTGCCCAATCACTCGTGGTAGGATTTTTTTATAAGAAAGTT      420
FRI_Sha         TTCCTCTTTCAGTTTTTTTTTGCCCAATCACTCGTGGTAGGATTTTTTTATAAGAAAGTT      420
FRI_Wa-1        TTCCTCTTTCAGTTTTTTTTTGCCCAATCACTCGTGGTAGGATTTTTTTATAAGAAAGTT      420
FRI_Spr-1-6     TTCCTCTTTCAGTTTTTTTTTGCCCAATCACTCGTGGTAGGATTTTTTTATAAGAAAGTT      420
FRI_Bil-7       TTCCTCTTTCAGTTTTTTTTTGCCCAATCACTCGTGGTAGGATTTTTTTATAAGAAAGTT      420
FRI_Alc-0       TTCCTCTTTCAGTTTTTTTTTGCCCAATCACTCGTGGTAGGATTTTTTTATAAGAAAGTT      420
FRI_Pu-2-23     TTCCTCTTTCAGTTTTTTTTTGCCCAATCACTCGTGGTAGGATTTTTTTATAAGAAAGTT      420
FRI_NFA-10      TTCCTCTTTCAGTTTTTTTTTGCCCAATCACTCGTGGTAGGATTT-TTTATAAGAAAGTT      419
FRI_Cvi-0       TTCCTCTTTCAGTTTTTTTTTGCCCAATCACTCGTGGTAGGATTTTTTTATAAGAAAGTT      420
FRI_Zdr-6       TTCCTCTTTCAGTTTTTTTTTGCCCAATCACTCGTGGTAGGATTTTTTTATAAGAAAGTT      420
                ********************    *****************  ** **************

FRI_An-1        ACGAAAATGCCCTCTCGTGCTTAGAGAGATTCAATTCGTGCACATATAATATGCCGTGAA      480
FRI_Pro-0       ACGAAAATGCCCTCTCGTGCTTAGAGAGATTCAATTCGTGCACATATAATATGCCGTGAA      479
FRI_Bå1-2       ACGAAAATGCCCTCTCGTGCTTAGAGAGATTCAATTCGTGCACATATAATATGCCGTGAA      480
FRI_St-0        ACGAAAATGCCCTCTCGTGCTTAGAGAGATTCAATTCGTGCACATATAATATGCCGTGAA      478
FRI_Bg-2        ACGAAAATGCCCTCTCGTGCTTAGAGAGATTCAATTCGTGCACATATAATATGCCGTGAA      478
FRI_C24         ACGAAAATGCCCTCTCGTGCTTAGAGAGATTCAATTCGTGCACATATAATATGCCGTGAA      478
FRI_Van-0       ACGAAAATGCCCTCTCGTGCTTAGAGAGATTCAATTCGTGCACATATAATATGCCGTGAA      478
FRI_Ull-2-3     ACGAAAATGCCCTCTCGTGCTTAGAGAGATTCAATTCGTGCACATATAATATGCCGTGAA      478
FRI_Lip-0       ACGAAAATGCCCTCTCGTGCTTAGAGAGATTCAATTCGTGCACATATAATATGCCGTGAA      478
FRI_NFA-8       ACGAAAATGCCCTCTCGTGCTTAGAGAGATTCAATTCGTGCACATATAATATGCCGTGAA      480
FRI_Edi-0       ACGAAAATGCCCTCTCGTGCTTAGAGAGATTCAATTCGTGCACATATAATATGCCGTGAA      480
FRI_Ren-1       ACGAAAATGCCCTCTCGTGCTTAGAGAGATTCAATTCGTGCACATATAATATGCCGTGAA      480
FRI_Nok-3       ACGAAAATGCCCTCTCGTGCTTAGAGAGATTCAATTCGTGCACATATAATATGCCGTGAA      480
FRI_Sha         ACGAAAATGCCCTCTCGTGCTTAGAGAGATTCAATTCGTGCACATATAATATGCCGTGAA      480
FRI_Wa-1        ACGAAAATGCCCTCTCGTGCTTAGAGAGATTCAATTCGTGCACATATAATATGCCGTGAA      480
FRI_Spr-1-6     ACGAAAATGCCCTCTCGTGCTTAGAGAGATTCAATTCGTGCACATATAATATGCCGTGAA      480
FRI_Bil-7       ACGAAAATGCCCTCTCGTGCTTAGAGAGATTCAATTCGTGAACATATAATATGCCGTGAA      480
FRI_Alc-0       ACGAAAATGCCCTCTCGTGCTTAGAGAGATTCAATTCGTGCACATATAATATGCCGTGAA      480
FRI_Pu-2-23     ACGAAAATGCCCTCTCGTGCTTAGAGAGATTCAATTCGTGCACATATAATATGCCGTGAA      480
FRI_NFA-10      ACGAAAATGCCCTCTCGTGCTTAGAGAGATTCAATTCGTGCACATATAATATGCCGTGAA      479
FRI_Cvi-0       ACGAAAATGCCCTCTCGTGCTTAGAGAGATTCAATTCGTGCACATATAATATGCCGTGAA      480
FRI_Zdr-6       ACGAAAATGCCCTCTCGTGCTTAGAGAGATTCAATTCGTGCACATATAATATGCCGTGAA      480
                **************************************** *******************

FRI_An-1        TATACCACCAACTCTCTTACTCGTACTCGCATGTGGTGCGTGCAATATACACGTGGTTCG      540
FRI_Pro-0       TATACCACCAACTCTCTTACTCGTACTCGCATGTGGTGCGTGCAATATACACGTGGTTCG      539
FRI_Bå1-2       TATACCACCAACTCTCTTACTCGTACTCGCATGTGGTGCGTGCAATATACACGTGGTTCG      540
FRI_St-0        TATACCACCAACTCTCTTACTCGTACTCGCATGTGGTGCGTGCAATATACACGTGATTCG      538
FRI_Bg-2        TATACCACCAACTCTCTTACTCGTACTCGCATGTGGTGCGTGCAATATATACGTGGTTCG      538
FRI_C24         TATACCACCAACTCTCTTACTCGTACTCGCATGTGGTGCGTGCAATATACACGTGGTTCG      538
FRI_Van-0       TATACCACCAACTCTCTTACTCGTACTCGCATGTGGTGCGTGCAATATACACGTGGTTCG      538
FRI_Ull-2-3     TATACCACCAACTCTCTTACTCGTACTCGCATGTGGTGCGTGCAATATACACGTGGTTCG      538
FRI_Lip-0       TATACCACCAACTCTCTTACTCGTACTCGCATGTGGTGCGTGCAATATACACGTGATTCG      538
FRI_NFA-8       TATACCACCAACTCTCTTACTCGTACTCGCATGTGGTGCGTGCAATATACACGTGGTTCG      540
FRI_Edi-0       TATACCACCAACTCTCTTACTCGTACTCGCATGTGGTGCGTGCAATATACACGTGGTTCG      540
FRI_Ren-1       TATACCACCAACTCTCTTACTCGTACTCGCATGTGGTGCGTGCAATATACACGTGGTTCG      540
FRI_Nok-3       TATACCACCAACTCTCTTACTCGTACTCGCATGTGGTGCGTGCAATATACACGTGGTTCG      540
FRI_Sha         TATACCACCAACTCTCTTACTCGTACTCGCATGTGGTGCGTGCAATATACACGTGGTTCG      540
FRI_Wa-1        TATACCACCAACTCTCTTACTCGTACTCGCATGTGGTGCGTGCAATATACACGTGGTTCG      540
FRI_Spr-1-6     TATACCACCAACTCTCTTACTCGTACTCGCATGTGGTGCGTGCAATATACACGTGGTTCG      540
```

```
FRI_Bil-7       TATACCACCAACTCTCTTACTCGTACTCGCATGTGGTGCGTGCAATATACACGTGGTTCG    540
FRI_Alc-0       TATACCACCAACTCTCTTACTCGTACTCGCATGTGGTGCGTGCAATATACACGTGGTTCG    540
FRI_Pu-2-23     TATACCACCAACTCTCTTACTCGTACTCGCATGTGGTGCGTGCAATATACACGTGGTTCG    540
FRI_NFA-10      TATACCACCAACTCTCTTACTCGTACTCGCATGTGGTGCGTGCAATATACACGTGGTTCG    539
FRI_Cvi-0       TATACCACCAACTCTCTTACTCGTACTCGCATGTGGTGCGTGCAATATACACGTGGTTCG    540
FRI_Zdr-6       TATACCACCAACTCTCTTACTCGTACTCGCATGTGGTGCGTGCAATATACACGTGGTTCG    540
                ************************************************* ***** ****

FRI_An-1        AATCAACGGCTACGTTTTACTGATAGGTGGCTCGTTTAGTTTTGCACTGCCTTTGGGTGC    600
FRI_Pro-0       AATCAACGGCTACGTTTTACTGATAGGTGGCTCGTTTAGTTTTGCACTGCCTTCGGGTGC    599
FRI_Bål-2       AATCAACGGCTACGTTTTACTGATAGGTGGCTCGTTTAGTTTTGCACTGCCTTTGGGTGC    600
FRI_St-0        AATCAACGGCTACGTTTTACTGATAGGTGGCTCGTTTAGTTTTGCACTGCCTTTGGGTGC    598
FRI_Bg-2        AATCAACGGCTACGTTTTACTGATAGGTGGCTCGTTTAGTTTTGCACTGCCTTTGGGTGC    598
FRI_C24         AATCAACGGCTACGTTTTACTGATAGGTGGCTCGTTTAGTTTTGCACTGCCTTTGGGTGC    598
FRI_Van-0       AATCAACGGCTACGTTTTACTGATAGGTGGCTCGTTTAGTTTTGCACTGCCTTTGGGTGC    598
FRI_Ull-2-3     AATCAACGGCTACGTTTTACTGATAGGTGGCTCGTTTAGTTTTGCACTGCCTTTGGGTGC    598
FRI_Lip-0       AATCAACGGCTACGTTTTACTGATAGGTGGCTCGTTTAGTTTTGCACTGCCTTTGGGTGC    598
FRI_NFA-8       AATCAACGGCTACGTTTTACTGATAGGTGGCTCGTTTAGTTTTGCACTGCCTTTGGGTGC    600
FRI_Edi-0       AATCAACGGCTACGTTTTACTGATAGGTGGCTCGTTTAGTTTTGCACTGCCTTTGGGTGC    600
FRI_Ren-1       AATCAACGGCTACGTTTTACTGATAGGTGGCTCGTTTAGTTTTGCACTGCCTTTGGGTGC    600
FRI_Nok-3       AATCAACGGCTACGTTTTACTGATAGGTGGCTCGTTTAGTTTTGCACTGCCTTTGGGTGC    600
FRI_Sha         AATCAACGGCTACGTTTTACTGATAGGTGGCTCGTTTAGTTTTGCACTGCCTTTGGGTGC    600
FRI_Wa-1        AATCAACGGCTACGTTTTACTGATAGGTGGCTCGTTTAGTTTTGCACTGCCTTTGGGTGC    600
FRI_Spr-1-6     AATCAACGGCTACGTTTTACTGATAGGTGGCTCGTTTAGTTTTGCACTGCCTTTGGGTGC    600
FRI_Bil-7       AATCAACGGCTACGTTTTACTGATAGGTGGCTCGTTTAGTTTTGCACTGCCTTTGGGTGC    600
FRI_Alc-0       AATCAACGGCTACGTTTTACTGATAGGTGGCTCGTTTAGTTTTGCACTGCCTTTGGGTGC    600
FRI_Pu-2-23     AATCAACGGCTACGTTTTACTGATAGGTGGCTCGTTTAGTTTTGCACTGCCTTTGGGTGC    600
FRI_NFA-10      AATCAACGGCTACGTTTTACTGATAGGTGGCTCGTTTAGTTTTGCACTGCCTTTGGGTGC    599
FRI_Cvi-0       AATCAACGGCTACGTTTTACTGATAGGTGGCTCGTTTAGTTTTGCACTGCCTTTGGGTGC    600
FRI_Zdr-6       AATCAACGGCTACGTTTTACTGATAGGTGGCTCGTTTAGTTTTGCACTGCCTTTGGGTGC    600
                ***************************************************** ******

FRI_An-1        GCGTGACCAAGTCATTTTCCAAATGTGTTTTAATATTTGTGAACTATTATTAATAAAAAA    660
FRI_Pro-0       GCGTGACCAAGTCATTTTCCAAATGTGTTTTAATATTTGTGAACTATTATTAATAAAAAA    659
FRI_Bål-2       GCGTGACCAAGTCATTTTCCAAATGTGTTTTAATATTTGTGAACTATTATTAATAAAAAA    660
FRI_St-0        GCGTGACCAAGTCATTTTCCAAATGTGTTTTAATATTTGTGAACTATTATTAATAAAAAA    658
FRI_Bg-2        GCGTGACCAAGTCATTTTCCAAATGTGTTTTAATATTTGTGAACTATTATTAATAAAAAA    658
FRI_C24         GCGTGACCAAGTCATTTTCCAAATGTGTTTTAATATTTGTGAACTATTATTAATAAAAAA    658
FRI_Van-0       GCGTGACCAAGTCATTTTCCAAATGTGTTTTAATATTTGTGAACTATTATTAATAAAAAA    658
FRI_Ull-2-3     GCGTGACCAAGTCATTTTCCAAATGTGTTTTAATATTTGTGAACTATTATTAATAAAAAA    658
FRI_Lip-0       GCGTGACCAAGTCATTTTCCAAATGTGTTTTAATATTTGTGAACTATTATTAATAAAAAA    658
FRI_NFA-8       GCGTGACCAAGTCATTTTCCAAATGTGTTTTAATATTTGTGAACTATTATTAATAAAAAA    660
FRI_Edi-0       GCGTGACCAAGTCATTTTCCAAATGTGTTTTAATATTTGTGAACTATTATTAATAAAAAA    660
FRI_Ren-1       GCGTGACCAAGTCATTTTCCAAATGTGTTTTAATATTTGTGAACTATTATTAATAAAAAA    660
FRI_Nok-3       GCGTGACCAAGTCATTTTCCAAATGTGTTTTAATATTTGTGAACTATTATTAATAAAAAA    660
FRI_Sha         GCGTGACCAAGTCATTTTCCAAATGTGTTTTAATATTTGTGAACTATTATTAATAAAAAA    660
FRI_Wa-1        GCGTGACCAAGTCATTTTCCAAATGTGTTTTAATATTTGTGAACTATTATTAATAAAAAA    660
FRI_Spr-1-6     GCGTGACCAAGTCATTTTCCAAATGTGTTTTAATATTTGTGAACTATTATTAATAAAAAA    660
FRI_Bil-7       GCGTGACCAAGTCATTTTCCAAATGTGTTTTAATATTTGTGAACTATTATTAATAAAAAA    660
FRI_Alc-0       GCGTGACCAAGTCATTTTCCAAATGTGTTTTGATATTTGTGAACTATTATTAATAAAAAA    660
FRI_Pu-2-23     GCGTGACCAAGTCATTTTCCAAATGTGTTTTAATATTTGTGAACTATTATTAATAAAAAA    660
FRI_NFA-10      GCGTGACCAAGTCATTTTCCAAATGTGTTTTAATATTTGTGAACTATTATTAATAAAAAA    659
FRI_Cvi-0       GCGTGACCAAGTCATTTTCCAAATGGGTTTTAATATTTGTGAACTATTATTAATAAAAAA    660
FRI_Zdr-6       GCGTGACCAAGTCATTTTCCAAATGTGTTTTAATATTTGTGAACTATTATTAATAAAAAA    660
                ************************* ***** ****************************

FRI_An-1        GCGAGCGACGTTGTTCTTTGATGAGGGAACGACCGTACGCTTGGCGACCTTCTCTGATTG    720
FRI_Pro-0       GCGAGCGACGTTGTTCTTTGATGAGGGAACGACCGTACGCTTGGCGACCTTCTCTGATTG    719
FRI_Bål-2       GCGAGCGACGTTGTTCTTTGATGAGGGAACGACCGTACGCTTGGCGACCTTCTCTGATTG    720
FRI_St-0        GCGAGCGACGTTGTTCTTTGATGAGGGAACGACCGTACGCTTGGCGACCTTCTCTGATTG    718
FRI_Bg-2        GCGAGCGACGTTGTTCTTTGATGAGGGAACGACCGTACGCTTGGCGACCTTCTCTGATTG    718
FRI_C24         GCGAGCGACGTTGTTCGTTGATGAGGGAGCGACCGTACGCTTGGCGACCTTCTCTGATTG    718
FRI_Van-0       GCGAGCGACGTTGTTCTTTGATGAGGGAACGACCGTACGCTTGGCGACCTTCTCTGATTG    718
FRI_Ull-2-3     GCGAGCGACGTTGTTCTTTGATGAGGGAACGACCGTACGCTTGGCGACCTTCTCTGATTG    718
FRI_Lip-0       GCGAGCGACGTTGTTCTTTGATGAGGGAACGACCGTACGCTTGGCGACCTTCTCTGATTG    718
FRI_NFA-8       GCGAGCGACGTTGTTCTTTGATGAGGGAACGACCGTACGCTTGGCGACCTTCTCTGATTG    720
FRI_Edi-0       GCGAGCGACGTTGTTCGTTGATGAGGGAGCGACCGTACGCTTGGCGACCTTCTCTGATTG    720
FRI_Ren-1       GCGAGCGACGTTGTTCGTTGATGAGGGAGCGACCGTACGCTTGGCGACCTTCTCTGATTG    720
FRI_Nok-3       GCGAGCGACGTTGTTCGTTGATGAGGGAGCGACCGTACGCTTGGCGACCTTCTCTGATTG    720
FRI_Sha         GCGAGCGACGTTGTTCTTTGATGAGGGAACGACCGTACGCTTGGCGACCTTCTCTGATTG    720
FRI_Wa-1        GCGAGCGACGTTGTTCTTTGATGAGGGAACGACCGTACGCTTGGCGACCTTCTCTGATTG    720
```

```
FRI_Spr-1-6      GCGAGCGACGTTGTTCTTTGATGAGGGAACGACCGTACGCTTGGCGACCTTCTCTGATTG      720
FRI_Bil-7        GCGAGCGACGTTGTTCTTTGATGAGGGAACGACCGTACGCTTGGCGACCTTCTCTGATTG      720
FRI_Alc-0        GCGAGCGACGTTGTTCTTTGATGAGGGAACGACCGTACGCTTGGCGACCTTCTCTGATTG      720
FRI_Pu-2-23      GCGAGCGACGTTGTTCGTTGATGAGGGAGCGACCGTACGCTTGGCGACCTTCTCTGATTG      720
FRI_NFA-10       GCGAGCGACGTTGTTCTTTGATGAGGGAACGACCGTACGCTTGGCGACCTTCTCTGATTG      719
FRI_Cvi-0        GCGAGCGACGTTGTTCTTTGATGAGGGAACGACCGTACGCTTGGCGACCTTCTCTGATTG      720
FRI_Zdr-6        GCGAGCGACGTTGTTCTTTGATGAGGGAACGACCGTACGCTTGGCGACCTTCTCTGATTG      720
                 **************** *********** *******************************

FRI_An-1         CGTAGAGTCGTTTTCCTTATTGGGCTTACCTAGTACCGTCAGCCCACAATGTTTTCATTT      780
FRI_Pro-0        CGTAGAGTCGTTTTCCTTATTGGGCTTACCTAGTACCGTCAGCCCACAATGTTTTCATTT      779
FRI_Bål-2        CGTAGAGTCGTTTTCCTTATTGGGCTTACCTAGTACCGTCAGCCCACAATGTTTTCATTT      780
FRI_St-0         CGTAGAGTCGTTTTCCTTATTGGGCTTACCTAGTACCGCCAGCCCACAATGTTTTCATTT      778
FRI_Bg-2         CGTAGAGTCGTTTTCCTTATTGGGCTTACCTAGTACCGTCAGCCCACAATGTTTTCATTT      778
FRI_C24          CGTAGAGTCGTTTTCCTTATTGGGCTTACCTAGTACCGTCAGCCCACAATGTTTTCATTT      778
FRI_Van-0        CGTAGAGTCGTTTTCCTTATTGGGCTTACCTAGTACCGTCAGCCCACAATGTTTTCATTT      778
FRI_Ull-2-3      CGTAGAGTCGTTTTCCTTACTGGGCTTACCTAGTACCGTCAGCCCACAATGTTTTCATTT      778
FRI_Lip-0        CGTAGAGTCGTTTTcCTTATTGGGCTTACCTAGTACCGCCTGCCCACAATGTTTTCATTT      778
FRI_NFA-8        CGTAGAGTCGTTTTCCTTATTGGGCTTACCTAGTACCGTCAGCCCACAATGTTTTCATTT      780
FRI_Edi-0        CGTAGAGTCGTTTTCCTTATTGGGCTTACCTAGTACCGTCAGCCCACAATGTTTTCATTT      780
FRI_Ren-1        CGTAGAGTCGTTTTCCTTATTGGGCTTACCTAGTACCGCCAGCCCACAATGTTTTCATTT      780
FRI_Nok-3        CGTAGAGTCGTTTTCCTTATTGGGCTTACCTAGTACCGCCAGCCCACAATGTTTTCATTT      780
FRI_Sha          CGTAGAGTCGTTTTCCTTATTGGGCTTACCTAGTACCGTCAGCCCACAATGTTTTCATTT      780
FRI_Wa-1         CGTAGAGTCGTTTTCCTTATTGGGCTTACCTAGTACCGTCAGCCCACAATGTTTTCATTT      780
FRI_Spr-1-6      CGTAGAGTCGTTTTCCTTATTGGGCTTACCTAGTACCGTCAGCCCACAATGTTTTCATTT      780
FRI_Bil-7        CGTAGAGTCGTTTTCCTTATTGGGCTTACCTAGTACCGTCAGCCCACAATGTTTTCATTT      780
FRI_Alc-0        CGTAGAGTCGTTTTCCTTATTGGGCTTACCTAGTACCGTCAGCCCACAATGTTTTCATTT      780
FRI_Pu-2-23      CGTAGAGTCGTTTTCCTTATTGGGCTTACCTAGTACCGTCAGCCCACAATGTTTTCATTT      780
FRI_NFA-10       CGTAGAGTCGTTTTCCTTATTGGGCTTACCTAGTACCGTCAGCCCACAATGTTTTCATTT      779
FRI_Cvi-0        CGTAGAGTCGTTTTCCTTATTGGGCTTACCTAGTACCGTCAGCCCACAATGTTTTCATTT      780
FRI_Zdr-6        CGTAGAGTCGTTTTCCTTACTGGGCTTACCTAGTACCGTCAGCCCACAATGTTTTCATTT      780
                 ******************* ****************** * *******************

FRI_An-1         CTTTCCCACCTAAAACTAAAAGTACTCACAAGTCACAACTTAAACCAAGTACACAAGGAT      840
FRI_Pro-0        CTTTCCCACCTAAAACTAAAAGTACTCACAAGTCACAACTTAAACCAAGTACACAAGGAT      839
FRI_Bål-2        CTTTCCCACCTAAAACTAAAAGTACTCACAAGTCACAACTTAAACCAAGTACACAAGGAT      840
FRI_St-0         CTTTCCCACCTAAAACTAAAAGTACTCACAAGTCACAACTTAAACCAAGTA---------      829
FRI_Bg-2         CTTTCCCACCTAAAACTAAAAGTACTCACAAGTCACAACTTAAACCAAGTACACAAGGAT      838
FRI_C24          CTTTCCCACCTAAAACTAAAAGTACTCACAAGTCACAACTTAAACCAAGTACACAAGGAT      838
FRI_Van-0        CTTTCCCACCTAAAACTAAAAGTACTCACAAGTCACAACTTAAACCAAGTACACAAGGAT      838
FRI_Ull-2-3      CTTTCCCACCTAAAACTAAAAGTACTCACAAGTCACAACTTAAACCAAGTACACAAGGAT      838
FRI_Lip-0        CTTTCCCACCTAAAACTAAAAGTACTCACAAGTCACAACTTAAACCAAGTACACAAGGAT      838
FRI_NFA-8        CTTTCCCACCTAAAACTAAAAGTACTCACAAGTCACAACTTAAACCAAGTACACAAGGAT      840
FRI_Edi-0        CTTTCCCACCTAAAACTAAAAGTACTCACAAGTCACAACTTAAACCAAGTACACAAGGAT      840
FRI_Ren-1        CTTTCCCACCTAAAACTAAAAGTACTCACAAGTCACAACTTAAACCAAGTACACAAGGAT      840
FRI_Nok-3        CTTTCCCACCTAAAACTAAAAGTACTCACAAGTCACAACTTAAACCAAGTACACAAGGAT      840
FRI_Sha          CTTTCCCACCTAAAACTAAAAGTACTCACAAGTCACAACTTAAACCAAGTACACAAGGAT      840
FRI_Wa-1         CTTTCCCACCTAAAACTAAAAGTACTCACAAGTCACAACTTAAACCAAGTACACAAGGAT      840
FRI_Spr-1-6      CTTTCCCACCTAAAACTAAAAGTACTCACAAGTCACAACTTAAACCAAGTACACAAGGAT      840
FRI_Bil-7        CTTTCCCACCTAAAACTAAAAGTACTCACAAGTCACAACTTAAACCAAGTACACAAGGAT      840
FRI_Alc-0        CTTTCCCACCTAAAACTAAAAGTACTCACAAGTCACAACTTAAACCAAGTACACAAGGAT      840
FRI_Pu-2-23      CTTTCCCACCTAAAACTAAAAGTACTCACAAGTCACAACTTAAACCAAGTACACAAGGAT      840
FRI_NFA-10       CTTTCCCACCTAAAACTAAAAGTACTCACAAGTCACAACTTAAACCAAGTACACAAGGAT      839
FRI_Cvi-0        CTTTCCCACCTAAAACTAAAAGTACTCACAAGTCACAACTTAAACCAAGTACACAAGGAT      840
FRI_Zdr-6        CTTTCCCACCTAAAACTAAAAGTACTCACAAGTCACAACTTAAACCAAGTACACAAGGAT      840
                 ***************************************************

FRI_An-1         TTTATCATGGGATTATCGTGTTTGAAGACTAAAAAGAGCACACCATCACCCCCATTAGTG      900
FRI_Pro-0        TTTATCATGGGATTATCGTGTTTGAAGACTAAAAAGAGCACACCATCACCCCCATTAGTG      899
FRI_Bål-2        TTTATCATGGGATTATCGTGTTTGAAGACTAAAAAGAGCACACCATCACCCCCATTAGTG      900
FRI_St-0         ------------------------------------------------------------      829
FRI_Bg-2         TTTATCATGGGATTATCGTGTTTGAAGACTAAAAAGAGCACACCATCACCCCCATTAGTG      898
FRI_C24          TTTATCATGGGATTATCGTGTTTGAAGACTAAAAAGAGCACACCATCACCCCCATTAGTG      898
FRI_Van-0        TTTATCATGGGATTATCGTGTTTGAAGACTAAAAAGAGCACACCATCACCCCCATTAGTG      898
FRI_Ull-2-3      TTTATCATGGGATTATCGTGTTTGAAGACTAAAAAGAGCACACCATCACCCCCATTAGTG      898
FRI_Lip-0        TTTATCATGGGATTATCGTGTTTGAAGACTAAAAAGAGCACACCATCACCCCCATTAGTG      898
FRI_NFA-8        TTTATCATGGGATTATCGTGTTTGAAGACTAAAAAGAGCACACCATCACCCCCATTAGTG      900
FRI_Edi-0        TTTATCATGGGATTATCGTGTTTGAAGACTAAAAAGAGCACACCATCACCCCCATTAGTG      900
FRI_Ren-1        TTTATCATGGGATTATCGTGTTTGAAGACTAAAAAGAGCACACCATCACCCCCATTAGTG      900
FRI_Nok-3        TTTATCATGGGATTATCGTGTTTGAAGACTAAAAAGAGCACACCATCACCCCCATTAGTG      900
FRI_Sha          TTTATCATGGGATTATCGTGTTTGAAGACTAAAAAGAGCACACCATCACCCCCATTAGTG      900
```

```
FRI_Wa-1        TTTATCATGGGATTATCGAGTTTGAAGACTAAAAAGAGCACACCATCACCCCCATTAGTG    900
FRI_Spr-1-6     TTTATCATGGGATTATCGAGTTTGAAGACTAAAAAGAGCACACCATCACCCCCATTAGTG    900
FRI_Bil-7       TTTATCATGGGATTATCGTGTTTGAAGACTAAAAAGAGCACACCATCACCCCCATTAGTG    900
FRI_Alc-0       TTTATCATGGGATTATCGTGTTTGAAGACTAAAAAGAGCACACCATCACCCCCATTAGTG    900
FRI_Pu-2-23     TTTATCATGGGATTATCGTGTTTGAAGACTAAAAAGAGCACACCATCACCCCCATTAGTG    900
FRI_NFA-10      TTTATCATGGGATTATCGTGTTTGAAGACTAAAAAGAGCACACCATCACCCCCATTAGTG    899
FRI_Cvi-0       TTTATCATGGGATTATCGTGTTTGAAGACTAAAAAGAGCACACCATCACCCCCATTAGTG    900
FRI_Zdr-6       TTTATCATGGGATTATCGTGTTTGAAGACTAAAAAGAGCACACCATCACCCCCATTAGTG    900


FRI_An-1        CAGGTAGAGTAAGACAGTAACTTTTGGGTTCATATTACCGAGCAAGAACCGTTATTTGTG    960
FRI_Pro-0       CAGGTAGAGTAAGACAGTAACTTTTGGGTTCATATTACCGAGCAAGAACCGTTATTTGTG    959
FRI_Bål-2       CAGGTAGAGTAAGACAGTAACTTTTGGGTTCATATTACCGAGCAAGAACCGTTATTTGTG    960
FRI_St-0        CAGGTAGAGTAAGACAGTAACTTTTGGGTTCATATTACCGAGCAAGAACCGTTATTTGTG    889
FRI_Bg-2        CAGGTAGAGTAAGACAGTAACTTTTGGGTTCATATTACCGAGCAAGAACCGTTATTTGTG    958
FRI_C24         CAGGTAGAGTAAGACAGTAACTTTTGGGTTCATATTACCGAGCAAGAACCGTTATTTGTG    958
FRI_Van-0       CAGGTAGAGTAAGACAGTAACTTTTGGGTTCATATTACCGAGCAAGAACCGTTATTTGTG    958
FRI_Ull-2-3     CAGGTAGAGTAAGACAGTAACTTTTGGGTTCATATTACCGAGCAAGAACCGTTATTTGTG    958
FRI_Lip-0       CAGGTAGAGTAAGACAGTAACTTTTGGGTTCATATTACCGAGCAAGAACCGTTATTTGTG    958
FRI_NFA-8       CAGGTAGAGTAAGACAGTAACTTTTGGGTTCATATTACCGAGCAAGAACCGTTATTTGTG    960
FRI_Edi-0       CAGGTAGAGTAAGACAGTAACTTTTGGGTTCATATTACCGAGCAAGAACCGTTATTTGTG    960
FRI_Ren-1       CAGGTAGAGTAAGACAGTAACTTTTGGGTTCATATTACCGAGCAAGAACCGTTATTTGTG    960
FRI_Nok-3       CAGGTAGAGTAAGACAGTAACTTTTGGGTTCATATTACCGAGCAAGAACCGTTATTTGTG    960
FRI_Sha         CAGGTAGAGTAAGACAGTAACTTTTGGGTTCATATTACCGAGCAAGAACCGTTATTTGTG    960
FRI_Wa-1        CAGGTAGAGTAAGACAGTAACTTTTGGGTTCATATTACCGAGCAAGAACCGTTATTTGTG    960
FRI_Spr-1-6     CAGGTAGAGTAAGACAGTAACTTTTGGGTTCATATTACCGAGCAAGAACCGTTATTTGTG    960
FRI_Bil-7       CAGGTAGAGTAAGACAGTAACTTTTGGGTTCATATTACCGAGCAAGAACCGTTATTTGTG    960
FRI_Alc-0       CAGGTAGAGTAAGACAGTAACTTTTGGGTTCATATTACCGAGCAAGAACCGTTATTTGTG    960
FRI_Pu-2-23     CAGGTAGAGTAAGACAGTAACTTTTGGGTTCATATTACCGAGCAAGAACCGTTATTTGTG    960
FRI_NFA-10      CAGGTAGAGTAAGACAGTAACTTTTGGGTTCATATTACCGAGCAAGAACCGTTATTTGTG    959
FRI_Cvi-0       CAGGTAGAGTAAGACAGTAACTTTTGGGTTCATATTACCGAGCAAGAACCGTTATTTGTG    960
FRI_Zdr-6       CAGGTAGAGTAAGACAGTAACTTTTGGGTTCATATTACCGAGCAAGAACCGTTATTTGTG    960
                ************************************************************

FRI_An-1        ATTAGACATGTTATAAACCACTGCTTTAGTGACTATTTAAAACAATATATTACATGTCGT    1020
FRI_Pro-0       ATTAGACATGTTATAAACCACTGCTTTAGTGACTATTTAAAACAATATATTACATGTCGT    1019
FRI_Bål-2       ATTAGACATGTTATAAACCACTGCTTTAGTGACTATTTAAAACAATATATTACATGTCGT    1020
FRI_St-0        ATTAGACATGTTATAAACCACTGCTTTAGTGACTATTTAAAACAATATATTACATGTCGT    949
FRI_Bg-2        ATTAGACATGTTATAAACCACTGCTTTAGTGACTATTTAAAACAATATATTACATGTCGT    1018
FRI_C24         ATTAGACATGTTATAAACCACTGCTTTAGTGACTATTTAAAACAATATATTACATGTCGT    1018
FRI_Van-0       ATTAGACATGTTATAAACCACTGCTTTAGTGACTATTTAAAACAATATATTACATGTCGT    1018
FRI_Ull-2-3     ATTAGACATGTTATAAACCACTGCTTTAGTGACTATTTAAAACAATATATTACATGTCGT    1018
FRI_Lip-0       ATTAGACATGTTATAAACCACTGCTTTAGTGACTATTTAAAACAATATATTACATGTCGT    1018
FRI_NFA-8       ATTAGACATGTTATAAACCACTGCTTTAGTGACTATTTAAAACAATATATTACATGTCGT    1020
FRI_Edi-0       ATTAGACATGTTATAAACCACTGCTTTAGTGACTATTTAAAACAATATATTACATGTCGT    1020
FRI_Ren-1       ATTAGACATGTTATAAACCACTGCTTTAGTGACTATTTAAAACAATATATTACATGTCGT    1020
FRI_Nok-3       ATTAGACATGTTATAAACCACTGCTTTAGTGACTATTTAAAACAATATATTACATGTCGT    1020
FRI_Sha         ATTAGACATGTTATAAACCACTGCTTTAGTGACTATTTAAAACAATATATTACATGTCGT    1020
FRI_Wa-1        ATTAGACATGTTATAAACCACTGCTTTAGTGACTATTTAAAACAATATATTACATGTCGT    1020
FRI_Spr-1-6     ATTAGACATGTTATAAACCACTGCTTTAGTGACTATTTAAAACAATATATTACATGTCGT    1020
FRI_Bil-7       ATTAGACATGTTATAAACCACTGCTTTAGTGACTATTTAAAACAATATATTACATGTCGT    1020
FRI_Alc-0       ATTAGACATGTTATAAACCACTGCTTTAGTGACTATTTAAAACAATATATTACATGTCGT    1020
FRI_Pu-2-23     ATTAGACATGTTATAAACCACTGCTTTAGTGACTATTTAAAACAATATATTACATGTCGT    1020
FRI_NFA-10      ATTAGACATGTTATAAACCACTGCTTTAGTGACTATTTAAAACAATATATTACATGTCGT    1019
FRI_Cvi-0       ATTAGACATGTTATAAACCACTGCTTTAGTGACTATTTAAAACAATATATTACATGTCGT    1020
FRI_Zdr-6       ATTAGACATGTTATAAACCACTGCTTTAGTGACTATTTAAAACAATATATTACATGTCGT    1020
                ************************************************************

FRI_An-1        AATCATGCAACCTAACTATGTTTTCATTAATCAAATACAAAGAATAAAGAGAAAAGTGCG    1080
FRI_Pro-0       AATCATGCAACCTAACTATGTTTTCATTAATCAAATACAAAGAATAAAGAGAAAAGTGCG    1079
FRI_Bål-2       AATCATGCAACCTAACTATGTTTTCATTAATCAAATACAAAGAATAAAGAGAAAAGTGCG    1080
FRI_St-0        AATCATGCAACCTAACTATGTTTTCATTAATCAAATACAAAGAATAAAGAGAAAAGTGCG    1009
FRI_Bg-2        AATCATGCAACCTAACTATGTTTTCATTAATCAAATACAAAGAATAAAGAGAAAAGTGCG    1078
FRI_C24         AATCATGCAACCTAACTATGTTTTCATTAATCAAATACAAAGAATAAAGAGAAAAGTGCG    1078
FRI_Van-0       AATCATGCAACCTAACTATGTTTTCATTAATCAAATACAAAGAATAAAGAGAAAAGTGCG    1078
FRI_Ull-2-3     AATCATGCAACCTAACTATGTTTTCATTAATCAAATACAAAGAATAAAGAGAAAAGTGCG    1078
FRI_Lip-0       AATCATGCAACCTAACTATGTTTTCATTAATCAAATACAAAGAATAAAGAGAAAAGTGCG    1078
FRI_NFA-8       AATCATGCAACCTAACTATGTTTTCATTAATCAAATACAAAGAATAAAGAGAAAAGTGCG    1080
FRI_Edi-0       AATCATGCAACCTAACTATGTTTTCATTAATCAAATACAAAGAATAAAGAGAAAAGTGCG    1080
FRI_Ren-1       AATCATGCAACC------------------------------------------------    1032
FRI_Nok-3       AATCATGCAACCTAACTATGTTTTCATTAATCAAATACAAAGAATAAAGAGAAAAGTGCG    1080
```

```
FRI_Sha         AATCATGCAACCTAACTATGTTTTCATTAATCAAATACAAAGAATAAAGAGAAAAGTGCG    1080
FRI_Wa-1        AATCATGCAACCTAACTATGTTTTCATTAATCAAATACAAAGAATAAAGAGAAAAGTGCG    1080
FRI_Spr-1-6     AATCATGCAACCTAACTATGTTTTCATTAATCAAATACAAAGAATAAAGAGAAAAGTGCG    1080
FRI_Bil-7       AATCATGCAACCTAACTATGTTTTCATTAATCAAATACAAAGAATAAAGAGAAAAGTGCG    1080
FRI_Alc-0       AATCATGCAACCTAACTATGTTTTCATTAATCAAATACAAAGAATAAAGAGAAAAGTGCG    1080
FRI_Pu-2-23     AATCATGCAACCTAACTATGTTTTCATTAGTCAAATACAAAGAATAAAGAGAAAAGTGCG    1080
FRI_NFA-10      AATCATGCAACCTAACTATGTTTTCATTAATCAAATACAAAGAATAAAGAGAAAAGTGCG    1079
FRI_Cvi-0       AATCATGCAACCTAACTATGTTTTCACTAAACAAATACAAAGAATAAAGAGAAAAGTGCG    1080
FRI_Zdr-6       AATCATGCAACCTAACTATGTTTTCATTAATCAAATACAAAGAATAAAGAGAAAAGTGCG    1080
                ************

FRI_An-1        TAGATTCAATTATTTGGCATAGACTCAAAAGAGTGTATATATATCTGACTTTTATTAAAT    1140
FRI_Pro-0       TAGATTCAATTATTTGGCATAGACTCAAAAGAGTGTATATATATCTGACTTTTATTAAAT    1139
FRI_Bål-2       TAGATTCAATTATTTGGCATAGACTCAAAAGAGTGTATATATATCTGACTTTTATTAAAT    1140
FRI_St-0        TAGATTCAATTATTTGGCATAGACTCAAAAGAGTGTATATATATCTGACTTTTATTAAAT    1069
FRI_Bg-2        TAGATTCAATTATTTGGCATAGACTCAAAAGAGTGTATATATATCTGACTTTTATTAAAT    1138
FRI_C24         TAGATTCAATTATTTGGCATAGACTCAAAAGAGTGTATATATATCTGACTTTTATTAAAT    1138
FRI_Van-0       TAGATTCAATTATTTGGCATAGACTCAAAAGAGTGTATATATATCTGACTTTTATTAAAT    1138
FRI_Ull-2-3     TAGATTCAATTATTTGGCATAGACTCAAAAGAGTGTATATATATCTGACTTTTATTAAAT    1138
FRI_Lip-0       TAGATTCAATTATTTGGCATAGACTCAAAAGAGTGTATATATATCTGACTTTTATTAAAT    1138
FRI_NFA-8       TAGATTCAATTATTTGGCATAGACTCAAAAGAGTGTATATATATCTGACTTTTATTAAAT    1140
FRI_Edi-0       TAGATTCAATTATTTGGCATAGACTCAAAAGAGTGTATATATATCTGACTTTTATTAAAT    1140
FRI_Ren-1       ------------------------------------------------------------    1032
FRI_Nok-3       TAGATTCAATTACTTGGCATAGACTCAAAAGAGTGTATATATATCTGACTTTTATTAAAT    1140
FRI_Sha         TAGATTCAATTATTTGGCATAGACTCAAAAGAGTGTATATATATCTGACTTTTATTAAAT    1140
FRI_Wa-1        TAGATTCAATTATTTGGCATAGACTCAAAAGAGTGTATATATATCTGACTTTTATTAAAT    1140
FRI_Spr-1-6     TAGATTCAATTATTTGGCATAGACTCAAAAGAGTGTATATATATCTGACTTTTATTAAAT    1140
FRI_Bil-7       TAGATTCAATTATTTGGCATAGACTCAAAAGAGTGTATATATATCTGACTTTTATTAAAT    1140
FRI_Alc-0       TAGATTCAATTATTTGGCATAGACTCAAAAGAGTGTATATATATCTGACTTTTATTAAAT    1140
FRI_Pu-2-23     TAGATTCAATTATTTGGCATAGACTCAAAAGAGTGTATATATATCTGACTTTTATTAAAT    1140
FRI_NFA-10      TAGATTCAATTATTTGGCATAGACTCAAAAGAGTGTATATATATCTGACTTTTATTAAAT    1139
FRI_Cvi-0       TAGATTCAATTATTTGGCATAGACTCAAAAGAGTGTATATATATCTGACTTTTATTAAAT    1140
FRI_Zdr-6       TAGATTCAATTATTTGGCATAGACTCAAAAGAGTGTATATATATCTGACTTTTATTAAAT    1140


FRI_An-1        TATTAAACACAAATACATATTTTCATAAGCAAAACTATAAAAGCCCTAAACATATAACGA    1200
FRI_Pro-0       TATTAAACACAAATACATATTTTCATAAGCAAAACTATAAAAGCCCTAAACATATAATGA    1199
FRI_Bål-2       TATTAAACACAAATACATATTTTCATAAGCAAAACTATAAAAGCCCTAAACATATAATGA    1200
FRI_St-0        TATTAAACACAAATACATATTTTCATAAGCAAAACTATAAAAGCCCTAAACATATAACGA    1129
FRI_Bg-2        TATTAAACACAAATACATATTTTCATAAGCAAAACTATAAAAGCCCTAAACATATAACGA    1198
FRI_C24         TATTAAACACAAATACATATTTTCATAAGCAAAACTATAAAAGCCCTAAACATATAACGA    1198
FRI_Van-0       TATTAAACACAAATACATATTTTCATAAGCAAAACTATAAAAGCCCTAAACATATAATGA    1198
FRI_Ull-2-3     TATTAAACACAAATACATATTTTCATAAGCAAAACTATAAAAGCCCTAAACATATAATGA    1198
FRI_Lip-0       TATTAAACACAAATACATATTTTCATAAGCAAAACTATAAAAGCCCTAAACATATAATGA    1198
FRI_NFA-8       TATTAAACACAAATACATATTTTCATAAGCAAAACTATAAAAGCCCTAAACATATAATGA    1200
FRI_Edi-0       TATTAAACACAAATACATATTTTCATAAGCAAAACTATAAAAGCCCTAAACATATAACGA    1200
FRI_Ren-1       ------------------------------------------------------------    1032
FRI_Nok-3       TATTAAACACAAATACATATTTTCATAAGCAAAACTATAAAAGCCCTAAACATATAATGA    1200
FRI_Sha         TATTAAACACAAATACATATTTTCATAAGCAAAACTATAAAAGCCCTAAAGATATAACGA    1200
FRI_Wa-1        TATTAAACACAAATACATATTTTCATAAGCAAAACTATAAAAGCCCTAAAGATATAACGA    1200
FRI_Spr-1-6     TATTAAACACAAATACATATTTTCATAAGCAAAACTATAAAAGCCCTAAAGATATAACGA    1200
FRI_Bil-7       TATTAAACACAAATACATATTTTCATAAGCAAAACTATAAAAGCCCTAAACATATAATGA    1200
FRI_Alc-0       TATTAAACACAAATACATATTTTCATAAGCAAAACTATAAAAGCCCTAAACATATAACGA    1200
FRI_Pu-2-23     TATTAAACACAAATACATATTTTCATAAGCAAAACTATAAAAGCCCTAAACATATAATGA    1200
FRI_NFA-10      TATTAAACACAAATACATATTTTCATAAGCAAAACTATAAAAGCCCTAAACATATAATGA    1199
FRI_Cvi-0       TATTAAACACAAATACATATTTTCATAAGCAAAACTATAAAAGCCCTAAACATATAATGA    1200
FRI_Zdr-6       TATTAAACACAAATACATATTTTCATAAGCAAAACTATAAAAGCCCTAAACATATAATGA    1200


FRI_An-1        TTACCTCAAAGGAAAAAGTCGTTTTCTCCTAATTAAAAGATAGGTTACTTCCTAATTA--    1258
FRI_Pro-0       TTACCTCAAAGGAAAAAGTCGTTTTCTCCTAATTAAAAGATAGGTTACTTCCTAATTAAT    1259
FRI_Bål-2       TTACCTCAAAGGAAAAAGTCGTTTTCTCCTACTTAAAAGATAGGTTACTTCCTAATTA--    1258
FRI_St-0        TTACCTCAAAGGAAAAAGTCGTTTTCTCCTAATTAAAAGATAGGTTACTTCCTAATTA--    1187
FRI_Bg-2        TTACCTCAAAGGAAAAAGTCGTTTTCTCCTAATTAAAAGATAGGTTACTTCCTAATTA--    1256
FRI_C24         TTACCTCAAAGGAAAAAGTCGTTTTCTCCTAATTAAAAGATAGGTTACTTCCTAATTA--    1256
FRI_Van-0       TTACCTCAAAGGAAAAAGTCGTTTTCTCCTAATTAAAAGATAGGTTACTTCCTAATTA--    1256
FRI_Ull-2-3     TTACCTCAAAGGAAAAAGTCGTTTTCTCCTAATTAAAAGATAGGTTACTTCCTAATTAAT    1258
FRI_Lip-0       TTACCTCAAAGGAAAAAGTCGTTTTCTCCTAATTAAAAGATAGGTTACTTCCTAATTAAT    1258
FRI_NFA-8       TTACCTCAAAGGAAAAAGTCGTTTTCTCCTAATTAAAAGATAGGTTACTTCCTAATTA--    1258
FRI_Edi-0       TTACCTCAAAGGAAAAAGTCGTTTTCTCCTAATTAAAAGATAGGTTACTTCCTAATTA--    1258
FRI_Ren-1       ------------------------------------------------------------    1032
```

```
FRI_Nok-3       TTACCTCAAAGGAAAAAGTCGTTTTCTCCTAATTAAAAGATAGGTTACTTCCTAATTA--        1258
FRI_Sha         TTACCTCAAAGGAAAAAGTCGTTTTCTCCTAATTAAAAGATAGGTTACTTCCTAATTA--        1258
FRI_Wa-1        TTACCTCAAAGGAAAAAGTCGTTTTCTCCTAATTAAAAGATAGGTTACTTCCTAATTA--        1258
FRI_Spr-1-6     TTACCTCAAAGGAAAAAGTCGTTTTCTCCTAATTAAAAGATAGGTTACTTCCTAATTA--        1258
FRI_Bil-7       TTACCTCAAAGGAAAAAGTCGTTTTCTCCTAATTAAAAGATAGGTTACTTCCTAATTA--        1258
FRI_Alc-0       TTACCTCAAAGGAAAAAGTCGTTTTCTCCTAATTAAAAGATAGGTTACTTCCTAATTA--        1258
FRI_Pu-2-23     TTACCTCAAAGGAAAAAGTCGTTTTCTCCTAATTAAAAGATAGGTTACTTCCTAATTAAT        1260
FRI_NFA-10      TTACCTCAAAGGAAAAAGTCGTTTTCTCCTAATTAAAAGATAGGTTACTTCCTAATTAAT        1259
FRI_Cvi-0       TTACCTCAAAGGAAAAAGTCGTTTTCTCCTAATTAAAAGATAGGTTACTTCCTAATTAAT        1260
FRI_Zdr-6       TTACCTCAAAGGAAAAAGTCGTTTTCTCCTAATTAAAAGATAGGTTACTTCCTAATTAAT        1260


FRI_An-1        -ATATATAATTTATGTGAACTTCACAATATACAGCTCAATAAAATTTGGTAATTTGACCG        1317
FRI_Pro-0       AGTATATAATTTATGTGAACTTCACAATATACAGTTCAATAAAATTTGGTAATTTGACCG        1319
FRI_Bål-2       -ATATATAATTTATGTGAACTTCACAATATACAGTTCAATAAAATTTGGTAATTTGACCG        1317
FRI_St-0        -ATATATAATTTATGTGAACTTCACAATATACAGTTCAATAAAATTTGGTAATTTGACCG        1246
FRI_Bg-2        -ATATATAATTTATGTGAACTTCACAATATACAGTTCAATAAAATTTGGTAATTTGACCG        1315
FRI_C24         -ATATATAATTTATGTGAACTTCACAATATACAGTTCAATAAAATTTGGTAATTTGACCG        1315
FRI_Van-0       -ATATATAATTTATGTGAACTTCACAATATACAGTTCAATAAAATTTGGTAATTTGACCG        1315
FRI_Ull-2-3     AGTATATAATTTATGTGAACTTCACAATATACAGTTCAATAAAATTTGGTAATTTGACCG        1318
FRI_Lip-0       AGTATATAATTTATGTGAACTTCACAATATACAGTTCAATAAAATTTGGTAATTTGACCG        1318
FRI_NFA-8       -ATATATAATTTATGTGAACTTCACAATATACAGTTCAATAAAATTTGGTAATTTGACCG        1317
FRI_Edi-0       -ATATATAATTTATGTGAACTTCACAATATACAGTTCAATAAAATTTGGTAATTTGACCG        1317
FRI_Ren-1       ------------------------------------------------------------        1032
FRI_Nok-3       -ATATATAATTTATGTGAACTTCACAATATACAGTTCAATAAAATTTGGTAATTTGACCG        1317
FRI_Sha         -ATATATAATTTATGTGAACTTCACAATATACAGTTCAATAAAATTTGGTAATTTGACCG        1317
FRI_Wa-1        -ATATATAATTTATGTGAACTTCACAATATACAGTTCAATAAAATTTGGTAATTTGACCG        1317
FRI_Spr-1-6     -ATATATAATTTATGTGAACTTCACAATATACAGTTCAATAAAATTTGGTAATTTGACCG        1317
FRI_Bil-7       -ATATATAATTTATGTGAACTTCACAATATACAGTTCAATAAAATTTGGTAATTTGACCG        1317
FRI_Alc-0       -ATATATAATTTATGTGAACTTCACAATATACAGTTCAATAAAATTTGGTAATTTGACCG        1317
FRI_Pu-2-23     AGTATATAATTTATGTGAACTTCACAATATACAGTTCAATAAAATTTGGTAATTTGACCG        1320
FRI_NFA-10      AGTATATAATTTATGTGAACTTCACAATATACAGTtCAATAAAATTTGGTAATTTGACCG        1319
FRI_Cvi-0       AGTATATAATTTATGTGAACTTCACAATATACAGTTCAATAAAATTTGGTAATTTGACCG        1320
FRI_Zdr-6       AGTATATAATTTATGTGAACTTCACAATATACAGTTCAATAAAATTTGGTAATTTGACCG        1320


FRI_An-1        ATTTAAGGAGAGTGGAAATTAGGGCTTCTGCAATCTTTTTTCTTCGCCGCAATCTCATGT        1377
FRI_Pro-0       ATTTAAGGAGAGTGGAAATTAGGGCTTCTGCAATCTTTTTTCTTCGCCGCAATCTCATGT        1379
FRI_Bål-2       ATTTAAGGAGAGTGGAAATTAGGGCTTCTGCAATCTTTTTTCTTCGCCGCAATCTCATGT        1377
FRI_St-0        ATTTAAGGAGAGTGGAAATTAGGGCTTCTGCAATCTTTTTTCTTCGCCGCAATCTCATGT        1306
FRI_Bg-2        ATTTAAGGAGAGTGGAAATTAGGGCTTCTGCAATCTTTTTTCTTCGCCGCAATCTCATGT        1375
FRI_C24         ATTTAAGGAGAGTGGAAATTAGGGCTTCTGCAATCTTTTTTCTTCGCCGCAATCTCATGT        1375
FRI_Van-0       ATTTAAGGAGAGTGGAAATTAGGGCTTCTGCAATCTTTTTTCTTCGCCGCAATCTCATGT        1375
FRI_Ull-2-3     ATTTAAGGAGAGTGGAAATTAGGGCTTCTGCAATCTTTTTTCTTCGCCGCAATCTCATGT        1378
FRI_Lip-0       ATTTAAGGAGAGTGGAAATTAGGGCTTCTGCAATCTTTTTTCTTCGCCGCAATCTCATGT        1378
FRI_NFA-8       ATTTAAGGAGAGTGGAAATTAGGGCTTCTGCAATCTTTTTTCTTCGCCGCAATCTCATGT        1377
FRI_Edi-0       ATTTAAGGAGAGTGGAAATTAGGGCTTCTGCAATCTTTTTTCTTCGCCGCAATCTCATGT        1377
FRI_Ren-1       ------------------------------------------------------------        1032
FRI_Nok-3       ATTTAAGGAGAGTGGAAATTAGGGCTTCTGCAATCTTTTTTCTTCGCCGCAATCTCATGT        1377
FRI_Sha         ATTTAAGGAGAGTGGAAATTTGGGCTTCTGCAATCTTTTTTCTTCGCCGCAATCTCATGT        1377
FRI_Wa-1        ATTTAAGGAGAGTGGAAATTTGGGCTTCTGCAATCTTTTTTCTTCGCCGCAATCTCATGT        1377
FRI_Spr-1-6     ATTTAAGGAGAGTGGAAATTTGGGCTTCTGCAATCTTTTTTCTTCGCCGCAATCTCATGT        1377
FRI_Bil-7       ATTTAAGGAGAGTGGAAATTAGGGCTTCTGCAATCTTTTTTCTTCGCCGCAATCTCATGT        1377
FRI_Alc-0       ATTTAAGGAGAGTGGAAATTAGGGCTTCTGCAATCTTTTTTCTTCGCCGCAATCTCATGT        1377
FRI_Pu-2-23     ATTTAAGGAGAGTGGAAATTAGGGCTTCTGCAATCTTTTTTCTTCGCCGCAATCTCATGT        1380
FRI_NFA-10      ATTTAAGGAGAGTGGAAATTAGGGCTTCTGCAATCTTTTTTCTTCGCCGCAATCTCATGT        1379
FRI_Cvi-0       ATTTAAGGAGAGTGGAAATTAGGGCTTCTGCAATCTTTTTTCTTCGCCGCAATCTCATGT        1380
FRI_Zdr-6       ATTTAAGGAGAGTGGAAATTAGGGCTTCTGCAATCTTTTTTCTTCGCCGCAATCTCATGT        1380
                                                                    Start codon


FRI_An-1        CCAATTATCCACCGACGGTGGCGGCGCAACCCACAACGACGGCGAATCCACTGCTGCAGC        1437
FRI_Pro-0       CCAATTATCCACCGACGGTGGCGGCGCAACCCACAACGACGGCGAATCCACTGCTGCAGC        1439
FRI_Bål-2       CCAATTATCCACCGACGGTGGCGGCGCAACCCACAACGACGGCGAATCCACTGCTGCAGC        1437
FRI_St-0        CCAATTATCCACCGACGGTGGCGGCGCAACCCACAACGACGGCGAATCCACTGCTGCAGC        1366
FRI_Bg-2        CCAATTATCCACCGACGGTGGCGGCGCAACCCACAACGACGGCGAATCCACTGCTGCAGC        1435
FRI_C24         CCAATTATCCACCGACGGTGGCGGCGCAACCCACAACGACGGCGAATCCACTGCTGCAGC        1435
FRI_Van-0       CCAATTATCCACCGACGGTGGCGGCGCAACCCACAACGACGGCGAATCCACTGCTGCAGC        1435
FRI_Ull-2-3     CCAATTATCCACCGACGGTGGCGGCGCAACCCACAACGACGGCGAATCCACTGCTGCAGC        1438
FRI_Lip-0       CCAATTATCCACCGACGGTGGCGGCGCAACCCACAACGACGGCAAATCCACTGCTGCAGC        1438
FRI_NFA-8       CCAATTATCCACCGACGGTGGCGGCGCAACCCACAACGACGGCGAATCCACTGCTGCAGC        1437
```

```
FRI_Edi-0        CCAATTATCCACCGACGGTGGCGGCGCAACCCACAACGACGGCGAATCCACTGCTGCAGC        1437
FRI_Ren-1        --------------------------TAACCCACAACGACGGCGAATCCACTGCTGCAGC        1066
FRI_Nok-3        CCAATTATCCACCGACGGTGGCGGCGCAACCCACAACGACGGCGAATCCACTGCTGCAGC        1437
FRI_Sha          CCAATTATCCACCGACGGTGGCGGCGCAACCCACAACGACGGCGAATCCACTGCTGCAGC        1437
FRI_Wa-1         CCAATTATCCACCGACGGTGGCGGCGCAACCCACAACGACGGCGAATCCACTGCTGCAGC        1437
FRI_Spr-1-6      CCAATTATCCACCGACGGTGGCGGCGCAACCCACAACGACGGCGAATCCACTGCTGCAGC        1437
FRI_Bil-7        CCAATTATCCACCGACGGTGGCGGCGCAACCCACAACGACGGCGAATCCACTGCTGCAGC        1437
FRI_Alc-0        CCAATTATCCACCGACGGTGGCGGCGCAACCCACAACGACGGCGAATCCACTGCTGCAGC        1437
FRI_Pu-2-23      CCAATTATCCACCGACGGTGGCGGCGCAACCCACAACGACGGCAAATCCACTGCTGCAGC        1440
FRI_NFA-10       CCAATTATCCACCGACGGTGGCGGCGCAACCCACAACGACGGCGAATCCACTGCTGCAGC        1439
FRI_Cvi-0        CCAATTATCCACCGACGGTGGCGGCGCAAACCACAACGACGGCGAATCCACTGCTGCAGC        1440
FRI_Zdr-6        CCAATTATCCACCGACGGTGGCGGCGCAACCCACAACGACGGCGAATCCACTGCTGCAGC        1440
                                            ** ************* ****************

FRI_An-1         GACATCAATCTGAACAGCGACGAAGAGAATTACCGAAGATTGTCGAAACAGAGTCTACAA        1497
FRI_Pro-0        GACATCAATCTGAACAGCGACGAAGAGAATTACCGAAGATTGTCGAAACAGAGTCTACAA        1499
FRI_Bål-2        GACATCAATCTGAACAGCGACGAAGAGAATTACCGAAGATTGTCGAAACAGAGTCTACAA        1497
FRI_St-0         GACATCAATCTGAACAGCGACGAAGAGAATTACCGAAGATTGTCGAAACAGAGTCTACAA        1426
FRI_Bg-2         GACATCAATCTGAACAGCGACGAAGAGAATTACCGAAGATTGTCGAAACAGAGTCTACAA        1495
FRI_C24          GACATCAATCTGAACAGCGCCGAAGAGAATTACCGAAGATTGTCGAAACAGAGTCTACAA        1495
FRI_Van-0        GACATCAATCTGAACAGCGACGAAGAGAATTACCGAAGATTGTCGAAACAGAGTCTACAA        1495
FRI_Ull-2-3      GACATCAATCTGAACAGCGACGAAGAGAATTACCGAAGATTGTCGAAACAGAGTCTACAA        1498
FRI_Lip-0        GACATCAATCTGAACAGCGACGAAGAGAATTACCGAAGATTGTCGAAACAGAGTCTACAA        1498
FRI_NFA-8        GACATCAATCTGAACAGCGACGAAGAGAATTACCGAAGATTGTCGAAACAGAGTCTACAA        1497
FRI_Edi-0        GACATCAATCTGAACAGCGACGAAGAGAATTACCGAAGATTGTCGAAACAGAGTCTACAA        1497
FRI_Ren-1        GACATCAATCTGAACAGCGACGAAGAGAATTACCGAAGATTGTCGAAACAGAGTCTACAA        1126
FRI_Nok-3        GACATCAATCTGAACAGCGACGAAGAGAATTACCGAAGATTGTCGAAACAGAGTCTACAA        1497
FRI_Sha          GACATCAATCTGAACAGCGACGAAGAGAATTACCGAAGATTGTCGAAACAGAGTCTACAA        1497
FRI_Wa-1         GACATCAATCTGAACAGCGACGAAGAGAATTACCGAAGATTGTCGAAACAGAGTCTACAA        1497
FRI_Spr-1-6      GACATCAATCTGAACAGCGACGAAGAGAATTACCGAAGATTGTCGAAACAGAGTCTACAA        1497
FRI_Bil-7        GACATCAATCTGAACAGCGACGAAGAGAATTACCGAAGATTGTCGAAACAGAGTCTACAA        1497
FRI_Alc-0        GACATCAATCTGAACAGCGACGAAGAGAATTACCGAAGATTGTCGAAACAGAGTCTACAA        1497
FRI_Pu-2-23      GACATCAATCTGAACAGCGACGAAGAGAATTACCGAAGATTGTCGAAACAGAGTCTACAA        1500
FRI_NFA-10       GACATCAATCTGAACAGCGACGAAGAGAATTACCGAAGATTGTCGAAACAGAGTCTACAA        1499
FRI_Cvi-0        GACATCAATCTGAACAGCGACGAAGAGAATTACCGAAGATTGTCGAAACAGAGTCTACAA        1500
FRI_Zdr-6        GACATCAATCTGAACAGCGACGAAGAGAATTACCGAAGATTGTCGAAACAGAGTCTACAA        1500
                 ******************* ****************************************

FRI_An-1         GTATGGACATTACGATCGGTCAATCTAAGCAGCCTCAATTTTTGAAATCCATAGACGAAT        1557
FRI_Pro-0        GTATGGACATTACGATCGGTCAATCTAAGCAGCCTCAATTTTTGAAATCCATAGACGAAT        1559
FRI_Bål-2        GTATGGACATTACGATCGGTCAATCTAAGCAGCCTCAATTTTTGAAATCCATAGACGAAT        1557
FRI_St-0         GTATGGACATTACGATCGGTCAATCTAAGCAGcCTCAATTTTTGAAATCCATAGACGAAT        1486
FRI_Bg-2         GTATGGACATTACGATCGGTCAATCTAAGCAGCCTCAATTTTTGAAATCCATAGACGAAT        1555
FRI_C24          GTATGGACATTACGATCGGTCAATCTAAGCAGCCTCAATTTTTGAAATCCATAGACGAAT        1555
FRI_Van-0        GTATGGACATTACGATCGGTCAATCTAAGCAGCCTCAATTTTTGAAATCCATAGACGAAT        1555
FRI_Ull-2-3      GTATGGACATTACGATCGGTCAATCTAAGCAGCCTCAATTTTTGAAATCCATAGACGAAT        1558
FRI_Lip-0        GTATGGACATTACGATCGGTCAATCTAAGCCGCCTCAATTTTTGAAATCCATAGACGAAT        1558
FRI_NFA-8        GTATGGACATTATGATCGGTCAATCTAAGCAGCCTCAATTTTTGAAATCCATAGACGAAT        1557
FRI_Edi-0        GTATGGACATTACGATCGGTCAATCTAAGCAGCCTCAATTTTTGAAATCCATAGACGAAT        1557
FRI_Ren-1        GTATGGACATTACGATCGGTCAATCTAAGCAGCCTCAATTTTTGAAATCCATAGACGAAT        1186
FRI_Nok-3        GTATGGACATTACGATCGGTCAATCTAAGCAGcCTCAATTTTTGAAATCCATAGAGGAAT        1557
FRI_Sha          GTATGGACATTACGATCGGTCAATCTAAGCAGCCTCAAATTTTGAAATCCATAGACGAAT        1557
FRI_Wa-1         GTATGGACATTACGATCGGTCAATCTAAGCAGCCTCAATTTTTGAAATCCATAGACGAAT        1557
FRI_Spr-1-6      GTATGGACATTACGATCGGTCAATCTAAGCAGCCTCAATTTTTGAAATCCATAGACGAAT        1557
FRI_Bil-7        GTATGGACATTACGATCGGTCAATCTAAGCAGCCTCAATTTTTGAAATCCATAGACGAAT        1557
FRI_Alc-0        GTATGGACATTACGATCGGTCAATCTAAGCAGCCTCAATTTTTGAAATCCATAGACGAAT        1557
FRI_Pu-2-23      GTATGGACATTACGATCGGTCAATCTAAGCAGCCTCAATTTTTGAAATCCATAGACGAAT        1560
FRI_NFA-10       GTATGGACATTACGATCGGTCAATCTAAGCAGCCTCAATTTTTGAAATCCATAGACGAAT        1559
FRI_Cvi-0        GTATGGACATTACGATCGGTCAATCTAAGCAGCCTCAATTTTTGAAATCCATAGACGAAT        1560
FRI_Zdr-6        GTATGGACATTACGATCGGTCAATCTAAGCAGCCTCAATTTTTGAAATCCATAGACGAAT        1560
                 ************ ***************** ******* **************** ****
```

**Start codon in Ren-1**

```
FRI_An-1         TAGCTGCGTTTTCAGTTGCAGTGGAAACATTCAAACGCCAATTCGATGATCTTCAGAAGC        1617
FRI_Pro-0        TAGCTGCGTTTTCAGTTGCAGTGGAAACATTCAAATGCCAATTCGATGATCTTCAGAAGC        1619
FRI_Bål-2        TAGCTGCGTTTTCAGTTGCAGTGGAAACATTCAAACGCCAATTCGATGATCTTCAGAAGC        1617
FRI_St-0         TAGCTGCGTTTTCAGTTGCAGTGGAAACATTCAAACGCCAATTCGATGATCTTCAGAAGC        1546
FRI_Bg-2         TAGCTGCGTTTTCAGTTGCAGTGGAAACATTCAAACGCCAATTCGATGATCTTCAGAAGC        1615
FRI_C24          TAGCTGCGTTTTCAGTTGCAGTGGAAACATTCAAACGCCAATTCGATGATCTTCAGAAGC        1615
FRI_Van-0        TAGCTGCGTTTTCAGTTGCAGTGGAAACATTCAAACGCCAATTCGATGATATTCAGAAGC        1615
FRI_Ull-2-3      TAGCTGCGTTTTCAGTTGCAGTGGAAACATTCAAATGCCAATTCGATGATCTTCAGAAGC        1618
```

```
FRI_Lip-0       TAGCTGCGTTTTCAGTTGCAGTGGAAACATTCAAACGCCAATTCGATGATCTTCAGAAGC      1618
FRI_NFA-8       TAGCTGCGTTTTCAGTTGCAGTGGAAACATTCAAACGCCAATTCGATGATCTTCAGAAGC      1617
FRI_Edi-0       TAGCTGCGTTTTCAGTTGCAGTGGAAACATTCAAACGCCAATTCGATGATCTTCAGAAGC      1617
FRI_Ren-1       TAGCTGCGTTTTCAGTTGCAGTGGAAACATTCAAACGCCAATTCGATGATCTTCAGAAGC      1246
FRI_Nok-3       TAGCTGCGTTTTCAGTTGCAGTGGAAACATTCAAACGCCAATTCGATGATCTTCAGAAGC      1617
FRI_Sha         TAGCTGCGTTTTCAGTTGCAGTGGAAACATTCAAACGCCAATTCGATGATCTTCAGAAGC      1617
FRI_Wa-1        TAGCTGCGTTTTCAGTTGCAGTGGAAACATTCAAATGCCAATTCGATGATCTTCAGAAGC      1617
FRI_Spr-1-6     TAGCTGCGTTTTCAGTTGCAGTGGAAACATTCAAATGCCAATTCGATGATCTTCAGAAGC      1617
FRI_Bil-7       TAGCTGCGTTTTCAGTTGCAGTGGAAACATTCAAACGCCAATTCGATGATCTTCAGAAGC      1617
FRI_Alc-0       TAGCTGCGTTTTCAGTTGCAGTGGAAATATTCAAACGCCAATTCGATGATCTTCAGAAGC      1617
FRI_Pu-2-23     TAGCTGCGTTTTCAGTTGCAGTGGAAACATTCAAACGCCAATTCGATGATCTTCAGAAGC      1620
FRI_NFA-10      TAGCTGCGTTTTCAGTTGCAGTGGAAACATTCAAATGCCAATTCGATGATCTTCAGAAGC      1619
FRI_Cvi-0       TAGCTGCGTTTTCAGTTGCAGTGGAAACATTCAAATGCCAATTCGATGATCTTCAGAAGC      1620
FRI_Zdr-6       TAGCTGCGTTTTCAGTTGCAGTGGAAACATTCAAATGCCAATTCGATGATCTTCAGAAGC      1620
                *************************** ******* ************** *********

FRI_An-1        ACATCGAGTCAATCGAAAACGCAATTGATTCCAAACTCGAGAGTAACGGCGTTGTCCTCG      1677
FRI_Pro-0       ACATCGAGTCAATCGAAAACGCAATTGATTCCAAACTCGAGAGTAACGGCGTTGTCCTCG      1679
FRI_Bål-2       ACATCGAGTCAATCGAAAACGCAATTGATTCCAAACTCGAGAGTAACGGCGTTGTCCTCG      1677
FRI_St-0        ACATCGAGTCAATCGAAAACGCAATTGATTCCAAACTCGAGAGTAACGGCGTTGTCCTCG      1606
FRI_Bg-2        ACATCGAGTCAATCGAAAACGCAATTGATTCCAAACTCGAGAGTAACGGCGTTGTCCTCG      1675
FRI_C24         ACATCGAGTCAATCGAAAACGCAATTGATTCCAAACTCGAGAGTAACGGCGTtGTCCTCG      1675
FRI_Van-0       ACATCGAGTCAATCGAAAACGCAATTGATTCCAAACTCGAGAGTAACGGCGTTGTCCTCG      1675
FRI_Ull-2-3     ACATCGAGTCAATCGAAAACGCAATTGATTCCAAACTCGAGAGTAACGGCGTTGTCCTCG      1678
FRI_Lip-0       ACATCGAGTCAATCGAAAACGCAATTGATTCCAAACTCGAGAGTAACGGCGTTGTCCTCG      1678
FRI_NFA-8       ACATCGAGTCAATCGAAAACGCAATTGATTCCAAACTCGAGAGTAACGGCGTTgTCCTCG      1677
FRI_Edi-0       ACATCGAGTCAATCGAAAACGCAATTGATTCCAAACTCGAGAGTAACGGCGTTGTCCTCG      1677
FRI_Ren-1       ACATCGAGTCAATCGAAAACGCAATTGATTCCAAACTCGAGAGTAACGGCGTTGTCCTCG      1306
FRI_Nok-3       ACATCGAGTCAATCGAAAACGCAATTGATTCCAAACTCGAGAGTAACGGCGTTGTCCTCG      1677
FRI_Sha         ACATCGAGTCAATCGAAAACGCAATTGATTCCAAACTCGAGAGTAACGGCGTTGTCCTCG      1677
FRI_Wa-1        ACATCGAGTCAATCGAAAACGCAATTGATTCCAAACTCGAGAGTAACGGCGTTGTCCTCG      1677
FRI_Spr-1-6     ACATCGAGTCAATCGAAAACGCAATTGATTCCAAACTCGAGAGTAACGGCGTTGTCCTCG      1677
FRI_Bil-7       ACATCGAGTCAATCGAAAACGCAATTGATTCCAAACTCGAGAGTAACGGCGTTGTCCTCG      1677
FRI_Alc-0       ACATCGAGTCAATCGAAAACGCAATTGATTCCAAACTCGAGAGTAACGGCGTTGTCCTCG      1677
FRI_Pu-2-23     ACATCGAGTCAATCGAAAACGCAATTGATTCCAAACTCGAGAGTAACGGCGTTGTCCTCG      1680
FRI_NFA-10      ACATCGAGTCAATCGAAAACGCAATTGATTCCAAACTCGAGAGTAACGGCGTTGTCCTCG      1679
FRI_Cvi-0       ACATCGAGTCAATCGAAAACGCAATTGATTCCAAACTCGAGAGTAACGGCGTTGTCCTCG      1680
FRI_Zdr-6       ACATCGAGTCAATCGAAAACGCAATTGATTCCAAACTCGAGAGTAACGGCGTTGTCCTCG      1680
                ************************************************************

FRI_An-1        CCGCGCGGAACAATAATTTCCATCAGCCGATGTTATCGCATCCGCGGAACAATGTATCTG      1737
FRI_Pro-0       CCGCGCGGAACAATAATTTCCATCAGCCGATGTTATCGCCTCCGCGGAACAATGTATCTG      1739
FRI_Bål-2       CCGCGCGGAACAATAATTTCCATCAGCCGATGTTATCGCCTCCGCGGAACAATGTATCTG      1737
FRI_St-0        CCGCGCGGAACAATAATTTCTATCAGCCGATGTTATCGCCTCCGCGGAACAATGTATCTG      1666
FRI_Bg-2        CCGCGCGGAACAATAATTTCCATCAGCCGATGTTATCGCCTCCGCGGAACAATGTATCTG      1735
FRI_C24         CCGCGCGgAACAATAATTTCCATCAGCCGATGTTATCGCCTCCGCGGAACAATGTATCTG      1735
FRI_Van-0       CCGCGCGGAACAATAATTTCCATCAGCCGATGTTATCGCCTCCGCGGAACAATGTATCTG      1735
FRI_Ull-2-3     CCGCGCGGAACAATAATTTCCATCAGCCGATGTTATCGCCTCCGCGGAACAATGTATCTG      1738
FRI_Lip-0       CCGCGCGgAACAATAATTTCCATCAGCCGATGTTATCGCCTCCGCGGAACAATGTATCTG      1738
FRI_NFA-8       CCGCGCGGAACAATAATTTCCATCAGCCGATGTTATCGCCTCCGCGGAACAATGTATCTG      1737
FRI_Edi-0       CCGCGCGGAACAATAATTTCCATCAGCCGATGTTATCGCCTCCGCGGaACAATGTATCTG      1737
FRI_Ren-1       CCGCGCGGAACAATAATTTCCATCAGCCGATGTTATCGCCTCCGCGGAACAATGTATCTG      1366
FRI_Nok-3       CCGCGCGGAACAATAATTTCCATCAGCCGATGTTATCGCCTCCGCGGAACAATGTATCTG      1737
FRI_Sha         CCGCGCGGAACAATAATTTCCATCAGCCGATGTTATCGCCTCCGCGGAACAATGTATCTG      1737
FRI_Wa-1        CCGCGCGGAACAATAATTTCCATCAGCCGATGTTATCGCCTCCGCGGAACAATGTATCTG      1737
FRI_Spr-1-6     CCGCGCGGAACAATAATTTCCATCAGCCGATGTTATCGCCTCCGCGGAACAATGTATCTG      1737
FRI_Bil-7       CCGCGCGGAACAATAATTTCCATCAGCCGATGTTATCGCCTCCGCGGAACAATGTATCTG      1737
FRI_Alc-0       CCGCGCGGAACAATAATTTCCATCAGCCGATGTTATCGCCTCCGCGGAACAATGTATCTG      1737
FRI_Pu-2-23     CCGCGCGGAACAATAATTTCCATCAGCCGATGTTATCGCCTCCGCAGAACAATGTATCTG      1740
FRI_NFA-10      CCGCGCGGAACAATAATTTCCATCAGCCGATGTTATCGCCTCCGCGGAACAATGTATCTG      1739
FRI_Cvi-0       CCGCGCGGAACAATAATTTCCATCAGCCGATGTTATCGCCTCCGCGGAACAATGTATCTG      1740
FRI_Zdr-6       CCGCGCGGAACAATAATTTCCATCAGCCGATGTTATCGCCTCCGCGGAACAATGTATCTG      1740
                ******************** ****************** ***** **************

FRI_An-1        TAGAAACCACCGTCACTGTGAGCCAACCGTCTCAGGAGATTGTACCGGAGACGTCGAATA      1797
FRI_Pro-0       TAGAAACCACCGTCACTGTGAGCCAACCGTCTCAGGAGATTGTACCGGAGACGTCGAATA      1799
FRI_Bål-2       TAGAAACCACCGTCACTGTGAGCCAACCGTCTCAGGAGATTGTACCGGAGACGTCGAATA      1797
FRI_St-0        TAGAAACCACCGTCACTGTGAGCCAACCGTCTCAGGAGATTGTACCGGAGACGTCGAATA      1726
FRI_Bg-2        TAGAAACCACCGTCACTGTGAGCCAACCGTCTCAGGAGATTGTACCGGAGACGTCGAATA      1795
FRI_C24         TAGAAACCACCGTCACTGTGAGCCAACCGTCTCAGGAGATTGTACCGGAGACGTCGAATA      1795
FRI_Van-0       TAGAAACCACCGTCACTGTGAGCCAACCGTCTCAGGAGATTGTACCGGAGACGTGGAATA      1795
```

```
FRI_Ull-2-3      TAGAAACCACCGTCACTGTGAGCCAACCGTCTCAGGAGATTGTACCGGAGACGTCGAATA        1798
FRI_Lip-0        TAGAAACCACCGTCACTGTGAGCCAACCGTCTCAGGAGATTGTACCGGAGACGTCGAATA        1798
FRI_NFA-8        TAGAAACCACCGTCACTGTGAGCCAACCGTCTCAGGAGATTGTACCGGAGACGTCGAATA        1797
FRI_Edi-0        TAGAAACCACCGTCACTGTGAGCCAACCGTCTCAGGAGATTGTACCGGAGACGTCGAATA        1797
FRI_Ren-1        TAGAAACCACCGTCACTGTGAGCCAACCGTCTCAGGAGATTGTACCGGAGACGTCGAATA        1426
FRI_Nok-3        TAGAAACCACCGTCACTGTGAGCCAACCGTCTCAGGAGATTGTACCGGAGACGTCGAATA        1797
FRI_Sha          TAGAAACCACCGTCACTGTGAGCCAACCGTCTCAGGAGATTGTACCGGAGACGTCGAATA        1797
FRI_Wa-1         TAGAAACCACCGTCACTGTGAGCCAACCGTCTCAGGAGATTGTACCGGAGACGTCGAATA        1797
FRI_Spr-1-6      TAGAAACCACCGTCACTGTGAGCCAACCGTCTCAGGAGATTGTACCGGAGACGTCGAATA        1797
FRI_Bil-7        TAGAAACCACCGTCACTGTGAGCCAACCGTCTCAGGAGATTGTACCGGAGACGTCGAATA        1797
FRI_Alc-0        TAGAAACCACCGTCACTGTGAGCCAACCGTCTCAGGAGATTGTACCGGAGACGTCGAATA        1797
FRI_Pu-2-23      TAGAAACCACCGTCACTGTGAGCCAACCGTCTCAGGAGATTGTACCGGAGACGTCGAATA        1800
FRI_NFA-10       TAGAAACCACCGTCACTGTGAGCCAACCGTCTCAGGAGATTGTACCGGAGACGTCGAATA        1799
FRI_Cvi-0        TAGAAACCACCGTCACTGTGAGCCAACCGTCTCAGGAGATTGTACCGGAGACGTCGAATA        1800
FRI_Zdr-6        TAGAAACCACCGTCACTGTGAGCCAACCGTCTCAGGAGATTGTACCGGAGACGTCGAATA        1800
                 ****************************************************** *****

FRI_An-1         AACCGGAGGGGGGACGTATGTGTGAGTTGATGTGTAGCAAAGGTCTGCGTAAATACATAT        1857
FRI_Pro-0        AACCGGAGGGGGGACGTATGTGTGAGTTGATGTGTAGCAAAGGTCTGCGTAAATACATAT        1859
FRI_Bål-2        AACCGGAGGGGGGACGTATGTGTGAGTTGATGTGTAGCAAAGGTCTGCGTAAATACATAT        1857
FRI_St-0         AACCGGAGGGGGAACGTATGTGTGAGTTGATGTGTAGCAAAGGTCTGCGTAAATACATAT        1786
FRI_Bg-2         AACCGGAGGGGGAACGTATATGTGAGTTGATGTGTAGCAAAGGTCTGCGTAAATACATAT        1855
FRI_C24          AACCGGAGGGGGAACGTATGTGTGAGTTGATGTGTAGCAAAGGTCTGCGTAAATACATAT        1855
FRI_Van-0        AACCGGAGGGGGGACGTATGTGTGAGTTGATGTGTAGCAAAGGTCTGCGTAAATACATAT        1855
FRI_Ull-2-3      AACCGGAGGGGGGACGTATGTGTGAGTTGATGTGTAGCAAAGGTCTGCGTAAATACATAT        1858
FRI_Lip-0        AACCGGAGGGGGGACGTATGTGTGAGTTGATGTGTAGCAAAGGTCTGCGTAAATACATAT        1858
FRI_NFA-8        AACCGGAGGGGGAACGTATGTGTGAGTTGATGTGTAGCAAAGGTCTGCGTAAATACATAT        1857
FRI_Edi-0        AACCGGAGGGGGAACGTATATGTGAGTTGATGTGTAGCAAAGGTCTGCGTAAATACATAT        1857
FRI_Ren-1        AACCGGAGGGGGAACGTATGTGTGAGTTGATGTGTAGCAAAGGTCTGCGTAAATACATAT        1486
FRI_Nok-3        AACCGGAGGGGGAACGTATGTGTGAGTTGATGTGTAGCAAAGGTCTGCGTAAATACATAT        1857
FRI_Sha          AACCGGAGGGGGGACGTATGTGTGAGTTGATGTGTAGCAAAGGTCTGCGTAAATACATAT        1857
FRI_Wa-1         AACCGGAGGGGGGACGTATGTGTGAGTTGATGTGTAGCAAAGGTCTGCGTAAATACATAT        1857
FRI_Spr-1-6      AACCGGAGGGGGGACGTATGTGTGAGTTGATGTGTAGCAAAGGTCTGCGTAAATACATAT        1857
FRI_Bil-7        AACCGGAGGGGGGACGTATGTGTGAGTTGATGTGTAGCAAAGGTCTGCGTAAATACATAT        1857
FRI_Alc-0        AACCGGAGGGGGGACGTATGTGTGAGTTGATGTGTAGCAAAGGTCTGCGTAAATACATAT        1857
FRI_Pu-2-23      AACCGGAGGGGGCACGTATGTGTGAGTTGATGTGTAGCAAAGGTCTGCGTAAATACATAT        1860
FRI_NFA-10       AACCGGAGGGGGGACGTATGTGTGAGTTGATGTGTAGCAAAGGTCTGCGTAAATACATAT        1859
FRI_Cvi-0        AACCGGAGGGGGGACGTATGTGTGAGTTGATGTGTAGCAAAGGTCTGCGTAAATACATAT        1860
FRI_Zdr-6        AACCGGAGGGGGGACGTATGTGTGAGTTGATGTGTAGCAAAGGTCTGCGTAAATACATAT        1860
                 ************ ****** ****************************************

FRI_An-1         ACGCGAATATCTCTGATCAAGCTAAGTTAATGGAAGAGATTCCTTCAGCTTTGAAATTGG        1917
FRI_Pro-0        ACGCGAATATCTCTGAACAAGCTAAGTTAATGGAAGAGATTCCTTCAGCTTTGAAATTGG        1919
FRI_Bål-2        ACGCGAATATCTCTGATCAAGCTAAGTTAATGGAAGAGATTCCTTCAGCTTTGAAATTGG        1917
FRI_St-0         ACGCGAATATCTCTGATCAAGCTAAGTTAATGGAAGAGATTCCTTCAGCTTTGAAATTGG        1846
FRI_Bg-2         ACGCGAATATCTCTGATCAAGCTTAGTTAATGGAAGAGATTCCTTCAGCTTTGAAATTGG        1915
FRI_C24          ACGCGAATATCTCTGATCAAGCTAAGTTAATGGAAGAGATTCCTTCAGCTTTGAAATTGG        1915
FRI_Van-0        ACGCGAATATCTCTGATCAAGCTAAGTTAATGGAAGAGATTCCTTCAGCTTTGAAATTGG        1915
FRI_Ull-2-3      ACGCGAATATCTCTGAACAAGCTAAGTTAATGGAAGAGATTCCTTCAGCTTTGAAATTGG        1918
FRI_Lip-0        ACGCGAATATCTCTGAACAAGCTAAGTTAATGGAAGAGATTCCTTCAGCTTTGAAATTGG        1918
FRI_NFA-8        ACGCGAATATCTCTGATCAAGCTAAGTTAATGGAAGAGATTCCTTCAGCTTTGAAATTGG        1917
FRI_Edi-0        ACGCGAATATCTCTGATCAAGCTAAGTTAATGGAAGAGATTCCTTCAGCTTTGAAATTGG        1917
FRI_Ren-1        ACGCGAATATCTCTGATCAAGCTAAGTTAATGGAAGAGATTCCTTCAGCTTTGAAATTGG        1546
FRI_Nok-3        ACGCGAATATCTCTGATCAAGCTAAGTTAATGGAAGAGATTCCTTCAGCTTTGAAATTGG        1917
FRI_Sha          ACGCGAATATCTCTGATCAAGCTAAGTTAATGGAAGAGATTCCTTCAGCTTTGAAATTGG        1917
FRI_Wa-1         ACGCGAATATCTCTGAACAAGCTAAGTTAATGGAAGAGATTCCTTCAGCTTTGAAATTGG        1917
FRI_Spr-1-6      ACGCGAATATCTCTGAACAAGCTAAGTTAATGGAAGAGATTCCTTCAGCTTTGAAATTGG        1917
FRI_Bil-7        ACGCGAATATCTCTGATCAAGCTAAGTTAATGGAAGAGATTCCTTCAGCTTTGAAATTGG        1917
FRI_Alc-0        AGTCGAATATCTCTGATCAAGCTAAGTTAATGGAAGAGATTCCTTCAGCTTTGAAATTGG        1917
FRI_Pu-2-23      ACGCGAATATCTCTGATCAAGCTAAGTTAATGGAAGAGATTCCTTCAGCTTTGAAATTGG        1920
FRI_NFA-10       ACGCGAATATCTCTGAACAAGCTAAGTTAATGGAAGAGATTCCTTCAGCTTTGAAATTGG        1919
FRI_Cvi-0        ACGCGAATATCTCTGAACAAGCTAAGTTAATGGAAGAGATTCCTTCAGCTTTGAAATTGG        1920
FRI_Zdr-6        ACGCGAATATCTCTGAACAAGCTAAGTTAATGGAAGAGATTCCTTCAGCTTTGAAATTGG        1920
                 *  ************* ****** ************************************

FRI_An-1         CCAAGGAGCCAGCGAAGTTTGTATTGGATTGTATTGGCAAGTTTTACTTACAAGGGCGTA        1977
FRI_Pro-0        CCAAGGAGCCAGCGAAGTTTGTATTGGATTGTATTGGCAAGTTTTACTTACAAGGGCGTA        1979
FRI_Bål-2        CCAAGGAGCCAGCGAAGTTTGTATTGGATTGTATTGGCAAGTTTTACTTACAAGGGCGTA        1977
FRI_St-0         CCAAGGAGCCAGCGAAGTTTGTATTGGATTGTATTGGCAAGTTTTACTTACAAGGGCGTA        1906
FRI_Bg-2         CCAAGGAGCCAGCGAAGTTTGTATTGGATTGTATTGGCAAGTTTTACTTACAAGGGCGTA        1975
FRI_C24          CCAAGGAGCCAGCGAAGTTTGTATTGGATTGTATTGGCAAGTTTTACTTACAAGGGCGTA        1975
```

```
FRI_Van-0       CCAAGGAGCCAGCGAAGTTTGTATTGGATTGTATTGGCAAGTTTTACTTACAAGGGCGTA    1975
FRI_Ull-2-3     CCAAGGAGCCAGCGAAGTTTGTATTGGATTGTATTGGCAAGTTTTACTTACAAGGGCGTA    1978
FRI_Lip-0       CCAAGGAGCCAGCGAAGTTTGTATTGGATTGTATTGGCAAGTTTTACTTACAAGGGCGTA    1978
FRI_NFA-8       CCAAGGAGCCAGCGAAGTTTGTATTGGATTGTATTGGCAAGTTTTACTTACAAGGGCGTA    1977
FRI_Edi-0       CCAAGGAGCCAGCGAAGTTTGTATTGGATTGTATTGGCAAGTTTTACTTACAAGGGCGTA    1977
FRI_Ren-1       CCAAGGAGCCAGCGAAGTTTGTATTGGATTGTATTGGCAAGTTTTACTTACAAGGGCGTA    1606
FRI_Nok-3       CCAAGGAGCCAGCGAAGTTTGTATTGGATTGTATTGGCAAGTTTTACTTACAAGGGCGTA    1977
FRI_Sha         CCAAGGAGCCAGCGAAGTTTGTATTGGATTGTATTGGCAAGTTTTACTTACAAGGGCGTA    1977
FRI_Wa-1        CCAAGGAGCCAGCGAAGTTTGTATTGGATTGTATTGGCAAGTTTTACTTACAAGGGCGTA    1977
FRI_Spr-1-6     CCAAGGAGCCAGCGAAGTTTGTATTGGATTGTATTGGCAAGTTTTACTTACAAGGGCGTA    1977
FRI_Bil-7       CCAAGGAGCCAGCGAAGTTTGTATTGGATTGTATTGGCAAGTTTTACTTACAAGGGCGTA    1977
FRI_Alc-0       CCAAGGAGCCAGCGAAGTTTGTATTGGATTGTATTGGCAAGTTTTACTTACAAGGGCGTA    1977
FRI_Pu-2-23     CCAAGGAGCCAGCGAAGTTTGTATTGGATTGTATTGGCAAGTTTTACTTACAAGGGCGTA    1980
FRI_NFA-10      CCAAGGAGCCAGCGAAGTTTGTATTGGATTGTATTGGCAAGTTTTACTTACAAGGGCGTA    1979
FRI_Cvi-0       CCAAGGAGCCAGCGAAGTTTGTATTGGATTGTATTGGCAAGTTTTACTTACAAGGGCGTA    1980
FRI_Zdr-6       CCAAGGAGCCAGCGAAGTTTGTATTGGATTGTATTGGCAAGTTTTACTTACAAGGGCGTA    1980
                ************************************************************

FRI_An-1        GAGCATTTACTAAAGAGTCGCCTATGAGCTCTGCGAGACAAGTTTCGCTTCTTATACTGG    2037
FRI_Pro-0       GAGCATTTACTAAAGAGTCGCCTATGAGCTCTGCGAGACAAGTTTCGCTTCTTATACTGG    2039
FRI_Bål-2       GAGCATTTACTAAAGAGTCGCCTATGAGCTCTGCGAGACAAGTTTCGCTTCTTATACTGG    2037
FRI_St-0        GAGCATTTACTAAAGAGTCGCCTATGAGCTCTGCGAGACAAGTTTCGCTTCTTATACTGG    1966
FRI_Bg-2        GAGCATTTACTAAAGAGTCGCCTATGAGCTCTGCGAGACAAGTTTCGCTTCTTATACTGG    2035
FRI_C24         GAGCATTTACTAAAGAGTCGCCTATGAGCTCTGCGAGACAAGTTTCGCTTCTTATACTGG    2035
FRI_Van-0       GAGCATTTACTAAAGAGTCGCCTATGAGCTCTGCGAGACAAGTTTCGCTTCTTATACTGG    2035
FRI_Ull-2-3     GAGCATTTACTAAAGAGTCGCCTATGAGCTCTGCGAGACAAGTTTCGCTTCTTATACTGG    2038
FRI_Lip-0       GAGCATTTACTAAAGAGTCGCCTATGAGCTCTGCGAGACAAGTTTCGCTTCTTATACTGG    2038
FRI_NFA-8       GAGCATTTACTAAAGAGTCGCCTATGAGCTCTGCGAGACAAGTTTCGCTTCTTATACTGG    2037
FRI_Edi-0       GAGCATTTACTAAAGAGTCGCCTATGAGCTCTGCGAGACAAGTTTCGCTTCTTATACTGG    2037
FRI_Ren-1       GAGCATTTACTAAAGAGTCGCCTATGAGCTCTGCGAGACAAGTTTCGCTTCTTATACTGG    1666
FRI_Nok-3       GAGCATTTACTAAAGAGTCGCCTATGAGCTCTGCGAGACAAGTTTCGCTTCTTATACTGG    2037
FRI_Sha         GAGCATTTACTAAAGAGTCGCCTATGAGCTCTGCGAGACAAGTTTCGCTTCTTATACTGG    2037
FRI_Wa-1        GAGCATTTACTAAAGAGTCGCCTATGAGCTCTGCGAGACAAGTTTCGCTTCTTATACTGG    2037
FRI_Spr-1-6     GAGCATTTACTAAAGAGTCGCCTATGAGCTCTGCGAGACAAGTTTCGCTTCTTATACTGG    2037
FRI_Bil-7       GAGCATTTACTAAAGAGTCGCCTATGAGCTCTGCGAGACAAGTTTCGCTTCTTATACTGG    2037
FRI_Alc-0       GAGCATTTACTAAAGAGTCGCCTATGAGCTCTGCGAGACAAGTTTCGCTTCTTATACTGG    2037
FRI_Pu-2-23     GAGCATTTACTAAAGAGTCGCCTATGAGCTCTGCGAGACAAGTTTCGCTTCTTATACTGG    2040
FRI_NFA-10      GAGCATTTACTAAAGAGTCGCCTATGAGCTCTGCGAGACAAGTTTCGCTTCTTATACTGG    2039
FRI_Cvi-0       GAGCATTTACTAAAGAGTCGCCTATGAGCTCTGCGAGACAAGTTTCGCTTCTTATACTGG    2040
FRI_Zdr-6       GAGCATTTACTAAAGAGTCGCCTATGAGCTCTGCGAGACAAGTTTCGCTTCTTATACTGG    2040
                ************************************************************

FRI_An-1        AGTCTTTTCTTCTAATGCCTGATCGTGGTAAAGGGAAGGTGAAGATTGAGAGTTGGATTA    2097
FRI_Pro-0       AGTCTTTTCTTCTAATGCCTGATCGTGGTAAAGGGAAGGTGAAGATTGAGAGTTGGATTA    2099
FRI_Bål-2       AGTCTTTTCTTCTAATGCCTGATCGTGGTAAAGGGAAGGTGAAGATTGAGAGTTGGATTA    2097
FRI_St-0        AGTCTTTTCTTCTAATGCCTGATCGTGGTAAAGGGAAGGTGAAGATTGAGAGTTGGATTA    2026
FRI_Bg-2        AGTCTTTTCTTCTAATGCCTGATCGTGGTAAAGGGAAGGTGAAGATTGAGAGTTGGATTA    2095
FRI_C24         AGTCTTTTCTTCTAATGCCTGATCGTGGTAAAGGGAAGGTGAAGATTGAGAGTTGGATTA    2095
FRI_Van-0       AGTCTTTTCTTCTAATGCCTGATCGTGGTAAAGGGAAGGTGAAGATTGAGAGTTGGATTA    2095
FRI_Ull-2-3     AGTCTTTTCTTCTAATGCCTGATCGTGGTAAAGGGAAGGTGAAGATTGAGAGTTGGATTA    2098
FRI_Lip-0       AGTCTTTTCTTCTAATGCCTGATCGTGGTAAAGGGAAGGTGAAGATTGAGAGTTGGATTA    2098
FRI_NFA-8       AGTCTTTTCTTCTAATGCCTGATCGTGGTAAAGGGAAGGTGAAGATTGAGAGTTGGATTA    2097
FRI_Edi-0       AGTCTTTTCTTCTAATGCCTGATCGTGGTAAAGGGAAGGTGAAGATTGAGAGTTGGATTA    2097
FRI_Ren-1       AGTCTTTTCTTCTAATGCCTGATCGTGGTAAAGGGAAGGTGAAGATTGAGAGTTGGATTA    1726
FRI_Nok-3       AGTCTTTTCTTCTAATGCCTGATCGTGGTAAAGGGAAGGTGAAGATTGAGAGTTGGATTA    2097
FRI_Sha         AGTCTTTTCTTCTAATGCCTGATCGTGGTAAAGGGAAGGTGAAGATTGAGAGTTGGATTA    2097
FRI_Wa-1        AGTCTTTTCTTCTAATGCCTGATCGTGGTAAAGGGAAGGTGAAGATTGAGAGTTGGATTA    2097
FRI_Spr-1-6     AGTCTTTTCTTCTAATGCCTGATCGTGGTAAAGGGAAGGTGAAGATTGAGAGTTGGATTA    2097
FRI_Bil-7       AGTCTTTTCTTCTAATGCCTGATCGTGGTAAAGGGAAGGTGAAGATTGAGAGTTGGATTA    2097
FRI_Alc-0       AGTCTTTTCTTCTAATGCCTGATCGTGGTAAAGGGAAGGTGAAGATTGAGAGTTGGATTA    2097
FRI_Pu-2-23     AGTCTTTTCTTCTAATGCCTGATCGTGGTAAAGGGAAGGTGAAGATTGAGAGTTGGATTA    2100
FRI_NFA-10      AGTCTTTTCTTCTAATGCCTGATCGTGGTAAAGGGAAGGTGAAGATTGAGAGTTGGATTA    2099
FRI_Cvi-0       AGTCTTTTCTTCTAATGCCTGATCGTGGTTAAGGGAAGGTGAAGATTGAGAGTTGGATTA    2100
FRI_Zdr-6       AGTCTTTTCTTCTAATGCCTGATCGTGGTAAAGGGAAGGTGAAGATTGAGAGTTGGATTA    2100
                ***************************** ******************************

FRI_An-1        AAGATGAGGCGGAGACGGCTGCTGTTGCTTGGAGGAAAAGGTTGATGACTGAAGGAGGAT    2157
FRI_Pro-0       AAGATGAGGCGGAGACGGCTGCTGTTGCTTGGAGGAAAAGGTTGATGACTGAAGGAGGAT    2159
FRI_Bål-2       AAGATGAGGCGGAGACGGCTGCTGTTGCTTGGAGGAAAAGGTTGATGACTGAAGGAGGAT    2157
FRI_St-0        AAGATGAGGCGGAGACGGCTGCTGTTGCTTGGAGGAAAAGGTTGATGACTGAAGGAGGAT    2086
FRI_Bg-2        AAGATGAGGCGGAGACGGCTGCTGTTGCTTGGAGGAAAAGGTTGATGACTGAAGGAGGAT    2155
```

```
FRI_C24         AAGATGAGGCGGAGACGGCTGCTGTTGCTTGGAGGAAAAGGTTGATGACTGAAGGAGGAT      2155
FRI_Van-0       AAGATGAGGCGGAGACGGCTGCTGTTGCTTGGAGGAAAAGGTTGATGACTGAAGGAGGAT      2155
FRI_Ull-2-3     AAGATGAGGCGGAGACGGCTGCTGTTGCTTGGAGGAAAAGGTTGATGACTGAAGGAGGAT      2158
FRI_Lip-0       AAGATGAGGCGGAGACGGCTGCTGTTGCTTGGAGGAAAAGGTTGATGACTGAAGGAGGAT      2158
FRI_NFA-8       AAGATGAGGCGGAGACGGCTGCTGTTGCTTGGAGGAAAAGGTTGATGACTGAAGGAGGAT      2157
FRI_Edi-0       AAGATGAGGCGGAGACGGCTGCTGTTGCTTGGAGGAAAAGGTTGATGACTGAAGGAGGAT      2157
FRI_Ren-1       AAGATGAGGCGGAGACGGCTGCTGTTGCTTGGAGGAAAAGGTTGATGACTGAAGGAGGAT      1786
FRI_Nok-3       AAGATGAGGCGGAGACGGCTGCTGTTGCTTGGAGGAAAAGGTTGATGACTGAAGGAGGAT      2157
FRI_Sha         AAGATGAGGCGGAGACGGCTGCTGTTGCTTGGAGGAAAAGGTTGATGACTGAAGGAGGAT      2157
FRI_Wa-1        AAGATGAGGCGGAGACGGCTGCTGTTGCTTGGAGGAAAAGGTTGATGACTGAAGGAGGAT      2157
FRI_Spr-1-6     AAGATGAGGCGGAGACGGCTGCTGTTGCTTGGAGGAAAAGGTTGATGACTGAAGGAGGAT      2157
FRI_Bil-7       AAGATGAGGCGGAGACGGCTGCTGTTGCTTGGAGGAAAAGGTTGATGACTGAAGGAGGAT      2157
FRI_Alc-0       AAGATGAGGCGGAGACGGCTGCTGTTGCTTGGAGGAAAAGGTTGATGACTGAAGGAGGAT      2157
FRI_Pu-2-23     AAGATGAGGCGGAGACGGCTGCTGTTGCTTGGAGGAAAAGGTTGATGACTGAAGGAGGAT      2160
FRI_NFA-10      AAGATGAGGCGGAGACGGCTGCTGTTGCTTGGAGGAAAAGGTTGATGACTGAAGGAGGAT      2159
FRI_Cvi-0       AAGATGAGGCGGAGACGGCTGCTGTTGCTTGGAGGAAAAGGTTGATGACTGAAGGAGGAT      2160
FRI_Zdr-6       AAGATGAGGCGGAGACGGCTGCTGTTGCTTGGAGGAAAAGGTTGATGACTGAAGGAGGAT      2160
                ************************************************************

FRI_An-1        TAGCTGCGGCTGAGAAAATGGATGCAAGGGGTTTGCTTTTACTAGTTGCTTGTTTTGGTG      2217
FRI_Pro-0       TAGCTGCGGCTGAGAAAATGGATGCAAGGGGTTTGCTTTTACTAGTTGCTTGTTTTGGTG      2219
FRI_Bå1-2       TAGCTGCGGCTGAGAAAATGGATGCAAGGGGTTTGCTTTTACGAGTTGCTTGTTTTGGTG      2217
FRI_St-0        TAGCTGCGGCTGAGAAAATGGATGCAAGGGGTTTGCTTTTACTAGTTGCTTGTTTTGGTG      2146
FRI_Bg-2        TAGCTGCGGCTGAGAAAATGGATGCAAGGGGTTTGCTTTTACTAGTTGCTTGTTTTGGTG      2215
FRI_C24         TAGCTGCGGCTGAGAAAATGGATGCAAGGGGTTTGCTTTTACTAGTTGCTTGTTTTGGTG      2215
FRI_Van-0       TAGCTGCGGCTGAGAAAATGGATGCAAGGGGTTTGCTTTTACTAGTTGCTTGTTTTGGTG      2215
FRI_Ull-2-3     TAGCTGCGGCTGAGAAAATGGATGCAAGGGGTTTGCTTTTACTAGTTGCTTGTTTTGGTG      2218
FRI_Lip-0       TAGCTGCGGCTGAGAAAATGGATGCAAGGGGTTTGCTTTTACTAGTTGCTTGTTTTGGTG      2218
FRI_NFA-8       TAGCTGCGGCTGAGAAAATGGATGCAAGGGGTTTGCTTTTACTAGTTGCTTGTTTTGGTG      2217
FRI_Edi-0       TAGCTGCGGCTGAGAAAATGGATGCAAGGGGTTTGCTTTTACTAGTTGCTTGTTTTGGTG      2217
FRI_Ren-1       TAGCTGCGGCTGAGAAAATGGATGCAAGGGGTTTGCTTTTACTAGTTGCTTGTTTTGGTG      1846
FRI_Nok-3       TAGCTGCGGCTGAGAAAATGGATGCAAGGGGTTTGCTTTTACTAGTTGCTTGTTTTGGTG      2217
FRI_Sha         TAGCTGCGGCTGAGAAAATGGATGCAAGGGGTTTGCTTTTACTAGTTGCTTGTTTTGGTG      2217
FRI_Wa-1        TAGCTGCGGCTGAGAAAATGGATGCAAGGGGTTTGCTTTTACTAGTTGCTTGTTTTGGTG      2217
FRI_Spr-1-6     TAGCTGCGGCTGAGAAAATGGATGCAAGGGGTTTGCTTTTACTAGTTGCTTGTTTTGGTG      2217
FRI_Bil-7       TAGCTGCGGCTGAGAAAATGGATGCAAGGGGTTTGCTTTTACTAGTTGCTTGTTTTGGTG      2217
FRI_Alc-0       TAGCTGCGGCTGAGAAAATGGATGCAAGGGGTTTGCTTTTACTAGTTGCTTGTTTTGGTG      2217
FRI_Pu-2-23     TAGCTGCGGCTGAGAAAATGGATGCAAGGGGTTTGCTTTTACTAGTTGCTTGTTTTGGTG      2220
FRI_NFA-10      TAGCTGCGGCTGAGAAAATGGATGCAAGGGGTTTGCTTTTACTAGTTGCTTGTTTTGGTG      2219
FRI_Cvi-0       TAGCTGCGGCTGAGAAAATGGATGCAAGGGGTTTGCTTTTACTAGTTGCTTGTTTTGGTG      2220
FRI_Zdr-6       TAGCTGCGGCTGAGAAAATGGATGCAAGGGGTTTGCTTTTACTAGTTGCTTGTTTTGGTG      2220
                ****************************************** *****************

FRI_An-1        TTCCTTCAAACTTTAGGAGTACAGATTTGCTGGAT-TTGATAAGGATGAGTGGTTCGAAT      2276
FRI_Pro-0       TTCCTTCAAACTTTAGGAGTACAGATTTGCTGGAT-TTGATAAGGATGAGTGGTTCGAAT      2278
FRI_Bå1-2       TTCCTTCAAACTTTAGGAGTACAGATTTGCTGGAT-TTGATAAGGATGAGTGGTTCGAAT      2276
FRI_St-0        TTCCTTCAAACTTTAGGAGTACAGATTTGCTGGAT-TTGATAAGGATGAGTGGTTCGAAT      2205
FRI_Bg-2        TTCCTTCAAACTTTAGGAGTACAGATTTGCTGGAT-TTGATAAGGATGAGTGGTTCGAAT      2274
FRI_C24         TTCCTTCAAACTTTAGGAGTACAGATTTGCTGGAT-TTGATAAGGATGAGTGGTTCGAAT      2274
FRI_Van-0       TTCCTTCAAACTTTAGGAGTACAGATTTGCTGGAT-TTGATAAGGATGAGTGGTTCGAAT      2274
FRI_Ull-2-3     TTCCTTCAAACTTTAGGAGTACAGATTTGCTGGAT-TTGATAAGGATGAGTGGTTCGAAT      2277
FRI_Lip-0       TTCCTTCAAACTTTAGGAGTACAGATTTGCTGGAT-TTGATAAGGATGAGTGGTTCGAAT      2277
FRI_NFA-8       TTCCTTCAAACTTTAGGAGTACGGATTTGCTGGATTTTGATAAGGATGAGTGGTTCGAAT      2277
FRI_Edi-0       TTCCTTCAAACTTTAGGAGTACAGATTTGCTGGAT-TTGATAAGGATGAGTGGTTCGAAT      2276
FRI_Ren-1       TTCCTTCAAACTTTAGGAGTACAGATTTGCTGGAT-TTGATAAGGATGAGTGGTTCGAAT      1905
FRI_Nok-3       TTCCTTCAAACTTTAGGAGTACAGATTTGCTGGAT-TTGATAAGGATGAGTGGTTCGAAT      2276
FRI_Sha         TTCCTTCAAACTTTAGGAGTACAGATTTGCTGGAT-TTGATAAGGATGAGTGGTTCGAAT      2276
FRI_Wa-1        TTCCTTCAAACTTTAGGAGTACAGATTTGCTGGAT-TTTATAAGGATGAGTGGTTCGAAT      2276
FRI_Spr-1-6     TTCCTTCAAACTTTAGGAGTACAGATTTGCTGGAT-TTGATAAGGATGAGTGGTTCGAAT      2276
FRI_Bil-7       TTCCTTCAAACTTTAGGAGTACAGATTTGCTGGAT-TTGATAAGGATGAGTGGTTCGAAT      2276
FRI_Alc-0       TTCCTTCAAACTTTAGGAGTACAGATTTGCTGGAT-TTGATAAGGATGAGTGGTTCGAAT      2276
FRI_Pu-2-23     TTCCTTCAAACTTTAGGAGTACAGATTTGCTGGAT-TTGATAAGGATGAGTGGTTCGAAT      2279
FRI_NFA-10      TTCCTTCAAACTTTAGGAGTACAGATTTGCTGGAT-TTGATAAGGATGAGTGGTTCGAAT      2278
FRI_Cvi-0       TTCCTTCAAACTTTAGGAGTACAGATTTGCTGGAT-TTGATAAGGATGAGTGGTTCGAAT      2279
FRI_Zdr-6       TTCCTTCAAACTTTAGGAGTACAGATTTGCTGGAT-TTGATAAGGATGAGTGGTTCGAAT      2279
                ********************** ************ ** *********************

FRI_An-1        GAGATTGCCGGTGCTTTGAAGCGGTCACAGTTTCTTGTCCCTATGGTCTCAGGTACCATA      2336
FRI_Pro-0       GAGATTGCCGGTGCTTTGAAGCGGTCACAGTTTCTTGTCCCTATGGTCTCAGGTACCATA      2338
FRI_Bå1-2       GGGATTGCCGGTGCTTTGAAGCGGTCACAGTTTCTTGTCCCTATGGTCTCAGGTACCATA      2336
FRI_St-0        GAGATTGCCGGTGCTTTGAAGCGGTCACAGTTTCTTGTCCCTATGGTCTCAGGTACCATA      2265
```

```
FRI_Bg-2        GAGATTGCCGGTGCTTTGAAGCGGTCACAGTTTCTTGTCCCTATGGTCTCAGGTACCATA    2334
FRI_C24         GAGATTGCCGGTGCTTTGAAGCGGTCACAGTTTCTTGTCCCTATGGTCTCAGGTACCATA    2334
FRI_Van-0       GAGATTGCCGGTGCTTTGAAGCGGTCACAGTTTCTTGTCCCTATGGTCTCAGGTACCATA    2334
FRI_Ull-2-3     GAGATTGCCGGTGCTTTGAAGCGGTCACAGTTTCTTGTCCCTATGGTCTCAGGTACCATA    2337
FRI_Lip-0       GAGATTGCCGGTGCTTTGAAGCGGTCACAGTTTCTTGTCCCTATGGTCTCAGGTACCATA    2337
FRI_NFA-8       GAGATTGCCGGTGCTTTGAAGCGGTCACAGTTTCTTGTCCCTATGGTCTCAGGTACCATA    2337
FRI_Edi-0       GAGATTGCCGGTGCTTTGAAGCGGTCACAGTTTCTTGTCCCTATGGTCTCAGGTACCATA    2336
FRI_Ren-1       GAGATTGCCGGTGCTTTGAAGCGGTCACAGTTTCTTGTCCCTATGGTCTCAGGTACCATA    1965
FRI_Nok-3       GAGATTGCCGGTGCTTTGAAGCGGTCACAGTTTCTTGTCCCTATGGTCTCAGGTACCATA    2336
FRI_Sha         GAGATTGCCGGTGCTTTGAAGCGGTCACAGTTTCTTGTCCCTATGGTCTCAGGTACCATA    2336
FRI_Wa-1        GAGATTGCCGGTGCTTTGAAGCGGTCACAGTTTCTTGTCCCTATGGTCTCAGGTACCATA    2336
FRI_Spr-1-6     GAGATTGCCGGTGCTTTGAAGCGGTCACAGTTTCTTGTCCCTATGGTCTCAGGTACCATA    2336
FRI_Bil-7       GAGATTGCCGGTGCTTTGAAGCGGTCACAGTTTCTTGTCCCTATGGTCTCAGGTACCATA    2336
FRI_Alc-0       GAGATTGCCGGTGCTTTGAAGCGGTCACAGTTTCTTGTCCCTATGGTCTCAGGTACCATA    2336
FRI_Pu-2-23     GAGATTGCCGGTGCTTTGAAGCGGTCACAGTTTCTTGTCCCTATGGTCTCAGGTACCATA    2339
FRI_NFA-10      GAGATTGCCGGTGCTTTGAAGCGGTCACAGTTTCTTGTCCCTATGGTCTCAGGTACCATA    2338
FRI_Cvi-0       GAGATTGCCGGTGCTTTGAAGCGGTCACAGTTTCTTGTCCCTATGGTCTCAGGTACCATA    2339
FRI_Zdr-6       GAGATTGCCGGTGCTTTGAAGCGGTCACAGTTTCTTGTCCCTATGGTCTCAGGTACCATA    2339
                * **********************************************************

FRI_An-1        TTCTGTTCTCACTCGGTGAATTTCATTGCAAAGGTGGTTCCTTTTGTTGACATCATCGAC    2396
FRI_Pro-0       TTCTGTTCTCACTCGGTGAATTTCATTGCAAAGGTGGTTCCTTTTGTTGACATCATCGAC    2398
FRI_Bål-2       TTCTGTTCTCACTCGGTGAATTTCATTGCAAAGGTGGTTCCTTTTGTTGACATCATCGAC    2396
FRI_St-0        TTCTGTtCTCACTCGGTGAATTTCATTGCAAAGGTGGTTCCTTTTGTTGACATCATCGAC    2325
FRI_Bg-2        TTCTGT-CTCACTCGGTGAATTTCATTGCAAAGGTGGTTCCTTTTGTTGACATCATCGAC    2393
FRI_C24         TTCTGTTCTCACTCGGTGAATTTCATTGCAAAGGTGGTTCCTTTTGTTGACATCATCGAC    2394
FRI_Van-0       TTCTGTtCTCACTCGGTGAATTTCATTGCAAAGGTGGTTCCTTTTGTTGACATCATCGAC    2394
FRI_Ull-2-3     TTCTGTtCTCACTCGGTGAATTTCATTGCAAAGGTGGTTCCTTTTGTTGACATCATCGAC    2397
FRI_Lip-0       TTCTGTTCTCACTCGGTGAATTTCATTGCAAAGGTGGTTCCTTTTGTTGACATCATCGAC    2397
FRI_NFA-8       TTCTGTTCTCACTCGGTGAATTTCATTGCAAAGGTGGTTCCTTTTGTTGACATCATCGAC    2397
FRI_Edi-0       TTCTGTTCTCACTCGGTGAATTTCATTGCAAAGGTGGTTCCTTTTGTTGACATCATCGAC    2396
FRI_Ren-1       TTCTGTTCTCACTCGGTGAATTTCATTGCAAAGGTGGTTCCTTTTGTTGACATCATCGAC    2025
FRI_Nok-3       TTCTGTTCTCACTCGGTGAATTTCATTGCAAAGGTGGTTCCTTTTGTTGACATCATCGAC    2396
FRI_Sha         TTCTGTTCTCACTCGGTGAATTTCATTGCAAAGGTGGTTCCTTTTGTTGACATCATCGAC    2396
FRI_Wa-1        TTCTGTTCTCACTCGGTGAATTTCATTGCAAAGGTGGTTCCTTTTGTTGACATCATCGAC    2396
FRI_Spr-1-6     TTCTGTTCTCACTCGGTGAATTTCATTGCAAAGGTGGTTCCTTTTGTTGACATCATCGAC    2396
FRI_Bil-7       TTCTGTtCTCACTCGGTGAATTTCATTGCAAAGGTGGTTCCTTTTGTTGACATCATCGAC    2396
FRI_Alc-0       TTCTGTTCTCACTCGGTGAATTTCATTGCAAAGGTGGTTCCTTTTGTTGACATCATCGAC    2396
FRI_Pu-2-23     TTCTGTTCTCACTCGGTGAATTTCATTGCAAAGGTGGTTCCTTTTGTTGACATCATCGAC    2399
FRI_NFA-10      TTCTGTTCTCACTCGGTGAATTTCATTGCAAAGGTGGTTCCTTTTGTTGACATCATCGAC    2398
FRI_Cvi-0       TTCTGTTCTCACTCGGTGAATTTCATTGCAAAGGTGGTTCCTTTTGTTGACATCATCGAC    2399
FRI_Zdr-6       TTCTGTtCTCACTCGGTGAATTTCATTGCAAAGGTGGTTCCTTTTGTTGACATCATCGAC    2399
                ****** *****************************************************

FRI_An-1        CAACATCAAGTTCCATCTTTGTTTTTCGATAAGCTTGATGGTATAAACTAGGAGAGCACA    2456
FRI_Pro-0       CAACATCAAGTTCCATCTTTGTTTTTCGATAAGCTTGATGGTATAAACTAGGAGAGCACA    2458
FRI_Bål-2       CAACATCAAGTTCCATCTTTGTTTTTCGATAAGCTTGATGGTATAAACTAGGAGAGCACA    2456
FRI_St-0        CAACATCAAGTTCCATCTTTGTTTTTCGATAAGCTTGATGGTATAAACTAGGAGAGCACA    2385
FRI_Bg-2        CAACATCAAGTTCCATCTTTGTTTTTCGATAAGCTTGATGGTACAAACTAGGAGAGCACA    2453
FRI_C24         CAACATCAAGTTCCATCTTTGTTTTTCGATAAGCTTGATGGTATAAACTAGGAGAGCACA    2454
FRI_Van-0       CAACATCAAGTTCCATCTTTGTTTTTCGATAAGCTTGATGGTATAAACTAGGAGAGCACA    2454
FRI_Ull-2-3     CAACATCAAGTTCCATCTTTGTTTTTCGATAAGCTTGATGGTATAAACTAGGAGAGCACA    2457
FRI_Lip-0       CAACATCAAGTTCCATCTTTGTTTTTCGATAAGCTTGATGGTATAAACTAGGAGAGCACA    2457
FRI_NFA-8       CAACATCAAGTTCCATCTTTGTTTTTCGATAAGCTTGATGGTATAAACTAGGAGAGCACA    2457
FRI_Edi-0       CAACATCAAGTTCCATCTTTGTTTTTCGATAAGCTTGATGGTATAAACTAGGAGAGCACA    2456
FRI_Ren-1       CAACATCAAGTTCCATCTTTGTTTTTCGATAAGCTTGATGGTATAAACTAGGAGAGCACA    2085
FRI_Nok-3       CAACATCAAGTTCCATCTTTGTTTTTCGATAAGCTTGATGGTATAAACTAGGAGAGCACA    2456
FRI_Sha         CAACATCAAGTTCCATCTTTGTTTTTCGATAAGCTTGATGGTATAAACTAGGAGAGCACA    2456
FRI_Wa-1        CAACATCAAGTTCCATCTTTGTTTTTCGATAAGCTTGATGGTATAAACTAGGAGAGCACA    2456
FRI_Spr-1-6     CAACATCAAGTTCCATCTTTGTTTTTCGATAAGCTTGATGGTATAAACTAGGAGAGCACA    2456
FRI_Bil-7       CAACATCAAGTTCCATCTTTGTTTTTCGATAAGCTTGATGGTATAAACTAGGAGAGCACA    2456
FRI_Alc-0       CAACATCAAGTTCCATCTTTGTTTTTCGATAAGCTTGATGGTATAAACTAGGAGAGCACA    2456
FRI_Pu-2-23     CAACATCAAGTTCCATCTTTGTTTTTCGATAAGCTTGATGGTATAAACTAGGAGAGCACA    2459
FRI_NFA-10      CAACATCAAGTTCCATCTTTGTTTTTCGATAAGCTTGATGGTATAAACTAGGAGAGCACA    2458
FRI_Cvi-0       CAACATCAAGTTCCATCTTTGTTTTTCGATAAGCTTGATGGTATAAACTAGGAGAGCACA    2459
FRI_Zdr-6       CAACATCAAGTTCCATCTTTGTTTTTCGATAAGCTTGATGGTATAAACTAGGAGAGCACA    2459
                ******************************************* ****************

FRI_An-1        TCAAATATTTAGAGTGCAATGACTGATTGAGCCAAATCCTAGCTAGAAATTAATCTGGAA    2516
FRI_Pro-0       TCAAATATTTAGAGTGCAATGACTGATTGAGCCAAATCCTAGCTAGAAATTAATCTGGAA    2518
FRI_Bål-2       TCAAATATTTAGAGTGCAATGACTGATTGAGCCAAATCCTAGCTAGAAATTAATCTGGAA    2516
```

```
FRI_St-0         TCAAATATTTAGAGTGCAATGACTGATTGAGCCAAATCCTAGCTAGAAATTAATCTGGAA         2445
FRI_Bg-2         TCAAATATTTAGAGTGCAATGACTGATTGAGCCAAATCCTAGCTAGAAATTAATCTGGAA         2513
FRI_C24          TCAAATATTTAGAGTGCAATGACTGATTGAGCCAAATCCTAGCTAGAAATTAATCTGGAA         2514
FRI_Van-0        TCAAATATTTAGAGTGCAATGACTGATTGAGCCAAATCCTAGCTAGAAATTAATCTGGAA         2514
FRI_Ull-2-3      TCAAATATTTAGAGTGCAATGACTGATTGAGCCAAATCCTAGCTAGAAATTAATCTGGAA         2517
FRI_Lip-0        TCAAATATTTAGAGTGCAATGACTGATTGAGCCAAATCCTAGCTAGAAATTAATCTGGAA         2517
FRI_NFA-8        TCAAATATTTAGAGTGCAATGACTGATTGAGCCAAATCCTAGCTAGAAATTAATCTGGAA         2517
FRI_Edi-0        TCAAATATTTAGAGTGCAATGACTGATTGAGCCAAATCCTAGCTAGAAATTAATCTGGAA         2516
FRI_Ren-1        TCAAATATTTAGAGTGCAATGACTGATTGAGCCAAATCCTAGCTAGAAATTAATCTGGAA         2145
FRI_Nok-3        TCAAATATTTAGAGTGCAATGACTGATTGAGCCAAATCCTAGCTAGAAATTAATCTGGAA         2516
FRI_Sha          TCAAATATTTAGAGTGCAATGACTGATTGAGCCAAATCCTAGCTAGAAATTAATCTGGAA         2516
FRI_Wa-1         TCAAATATTTAGAGTGCAATGACTGATTGAGCCAAATCCTAGCTAGAAATTAATCTGGAA         2516
FRI_Spr-1-6      TCAAATATTTAGAGTGCAATGACTGATTGAGCCAAATCCTAGCTAGAAATTAATCTGGAA         2516
FRI_Bil-7        TCAAATATTTAGAGTGCAATGACTGATTGAGCCAAATCCTAGCTAGAAATTAATCTGGAA         2516
FRI_Alc-0        TCAAATATTTAGAGTGCAATGACTGATTGAGCCAAATCCTAGCTAGAAATTAATCTGGAA         2516
FRI_Pu-2-23      TCAAATATTTAGAGTGCAATGACTGATTGAGCCAAATCCTAGCTAGAAATTAATCTGGAA         2519
FRI_NFA-10       TCAAATATTTAGAGTGCAATGACTGATTGAGCCAAATCCTAGCTAGAAATTAATCTGGAA         2518
FRI_Cvi-0        TCAAATATTTAGAGTGCAATGACTGATTGAGCCAAATCCTAGCTAGAAATTAATCTGGAA         2519
FRI_Zdr-6        TCAAATATTTAGAGTGCAATGACTGATTGAGCCAAATCCTAGCTAGAAATTAATCTGGAA         2519
                 ************************************************************

FRI_An-1         AGAACTTGGAACTCTCAACCATAGGTTTTGGTACGAAATTGTTGCTTGTCAGAACCAAAT         2576
FRI_Pro-0        AGAACTTGGAACTCTCAACCATAGGTTTTGGTACGAAATTGTTGCTTGTCAGAACCAAAT         2578
FRI_Bå1-2        AGAACTTGGAACTCTCAACCATAGGTTTTGGTACGAAATTGTTGCTTGTCAGAACCAAAT         2576
FRI_St-0         AGAACTTGGAACTCTCAACCATAGGTTTTGGTACGAAATTGTTGCTTGTCAGAACCAAAT         2505
FRI_Bg-2         AGAACTTGGAACTCTCAACCATAGGTTTTGGTACGAAATTGTTGCTTGTCAGAACCAAAT         2573
FRI_C24          AGAACTTGGAACTCTCAACCATAGGTTTTGGTACGAAATTGTTGCTTGTCAGAACCAAAT         2574
FRI_Van-0        AGAACTTGGAACTCTCAACCATAGGTTTTGGTACGAAATTGTTGCTTGTCAGAACCAAAT         2574
FRI_Ull-2-3      AGAACTTGGAACTCTCAACCATAGGTTTTGGTACGAAATTGTTGCTTGTCAGAACCAAAT         2577
FRI_Lip-0        AGAACTTGGAACTCTCAACCATAGGTTTTGGTACGAAATTGTTGCTTGTCAGAACCAAAT         2577
FRI_NFA-8        AGAACTTGGAACTCTCAACCATAGGTTTTGGTACGAAATTGTTGCTTGTCAGAACCAAAT         2577
FRI_Edi-0        AGAACTTGGAACTCTCAACCATAGGTTTTGGTACGAAATTGTTGCTTGTCAGAACCAAAT         2576
FRI_Ren-1        AGAACTTGGAACTCTCAACCATAGGTTTTGGTACGAAATTGTTGCTTGTCAGAACCAAAT         2205
FRI_Nok-3        AGAACTTGGAACTCTCAACCATAGGTTTTGGTACGAAATTGTTGCTTGTCAGAACCAAAT         2576
FRI_Sha          AGAACTTGGAACTCTCAACCATAGGTTTTGGTACGAAATTGTTGCTTGTCAGAACCAAAT         2576
FRI_Wa-1         AGAACTTGGAACTCTCAACCATAGGTTTTGGTACGAAATTGTTGCTTGTCAGAACCAAAT         2576
FRI_Spr-1-6      AGAACTTGGAACTCTCAACCATAGGTTTTGGTACGAAATTGTTGCTTGTCAGAACCAAAT         2576
FRI_Bil-7        AGAACTTGGAACTCTCAACCATAGGTTTTGGTACGAAATTGTTGCTTGTCAGAACCAAAT         2576
FRI_Alc-0        AGAACTTGGAACTCTCAACCATAGGTTTTGGTACGAAATTGTTGCTTGTCAGAACCAAAT         2576
FRI_Pu-2-23      AGAACTTGGAACTCTCAACCATAGGTTTTGGTACGAAATTGTTGCTTGTCAGAACCAAAT         2579
FRI_NFA-10       AGAACTTGGAACTCTCAACCATAGGTTTTGGTACGAAATTGTTGCTTGTCAGAACCAAAT         2578
FRI_Cvi-0        AGAACTTGGAACTCTCAACCATAGGTTTTGGTACGAAATTGTTGCTTGTCAGAACCAAAT         2579
FRI_Zdr-6        AGAACTTGGAACTCTCAACCATAGGTTTTGGTACGAAATTGTTGCTTGTCAGAACCAAAT         2579
                 ************************************************************

FRI_An-1         GATAGGCTATTGCCTTGAAATAGTGTTTCTTGTGGTTTCCAATATTGGAAGTTAAAATCA         2636
FRI_Pro-0        GATAGGCTATTGCCTTGAAATAGTGTTTCTTGTGGTTTCCAATATTGGAAGTTAAAATCA         2638
FRI_Bå1-2        GATAGGCTATTGCCTTGAAATAGTGTTTCTTGTGGTTTCCAATATTGGAAGTTAAAATCG         2636
FRI_St-0         GATAGGCTATTGCCTTGAAATAGTGTTTCTTGTGGTTTCCAATATTGGAAGTTAAAATCA         2565
FRI_Bg-2         GATAGGCTATTGCCTTGAAATAGTGTTTCTTGTGGTTTCCAATATTGGAAGTTAAAATCA         2633
FRI_C24          GATAGGCTATTGCCTTGAAATAGTGTTTCTTGTGGTTTCCAATATTGGAAGTTAAAATCA         2634
FRI_Van-0        GATAGGCTATTGCCTTGAAATAGTGTTTCTTGTGGTTTCCAATATTGGAAGTTAAAATCA         2634
FRI_Ull-2-3      GATAGGCTATTGCCTTGAAATAGTGTTTCTTGTGGTTTCCAATATTGGAAGTTAAAATCA         2637
FRI_Lip-0        GATAGGCTATTGCCTTGAAATAGTGTTTCTTGTGGTTTCCAATATTGGAAGTTAAAATCA         2637
FRI_NFA-8        GATAGGCTATTGCCTTGAAATAGTGTTTCTTGTGGTTTCCAATATTGGAAGTTAAAATCA         2637
FRI_Edi-0        GATAGGCTATTGCCTTGAAATAGTGTTTCTTGTGGTTTCCAATATTGGAAGTTAAAATCA         2636
FRI_Ren-1        GATAGGCTATTGCCTTGAAATAGTGTTTCTTGTGGTTTCCAATATTGGAAGTTAAAATCA         2265
FRI_Nok-3        GATAGGCTATTGCCTTGAAATAGTGTTTCTTGTGGTTTCCAATATTGGAAGTTAAAATCA         2636
FRI_Sha          GATAGGCTATTGCCTTGAAATAGTGTTTCTTGTGGTTTCCAATATTGGAAGTTAAAATCA         2636
FRI_Wa-1         GATAGGCTATTGCCTTGAAATAGTGTTTCTTGTGGTTTCCAATATTGGAAGTTAAAATCA         2636
FRI_Spr-1-6      GATAGGCTATTGCCTTGAAATAGTGTTTCTTGTGGTTTCCAATATTGGAAGTTAAAATCA         2636
FRI_Bil-7        GATAGGCTATTGCCTTGAAATAGTGTTTCTTGTGGTTTCCAATATTGGAAGTTAAAATCG         2636
FRI_Alc-0        GATAGGCTATTGCCTTGAAATAGTGTTTCTTGTGGTTTCCAATATTGGAAGTTAAAATCG         2636
FRI_Pu-2-23      GATAGGCTATTGCCTTGAAATAGTGTTTCTTGTGGTTTCCAATATTGGAAGTTAAAATCA         2639
FRI_NFA-10       GATAGGCTATTGCCTTGAAATAGTGTTTCTTGTGGTTTCCAATATTGGAAGTTAAAATCA         2638
FRI_Cvi-0        GATAGGCTATTGCCTTGAAATAGTGTTTCTTGTGGTTTCCAATATTGGAAGTTAAAATCA         2639
FRI_Zdr-6        GATAGGCTATTGCCTTGAAATAGTGTTTCTTGTGGTTTCCAATATTGGAAGTTAAAATCA         2639
                 ***********************************************************

FRI_An-1         TATGACTTAGCTGTTGGATACTAATTAAGCTTAAGCAATGCCAACTCTAAGAAGTGGTAC         2696
FRI_Pro-0        TATGACTTAGCTGTTGGATACTAATTAAGCTTAAGCAATGCCAACTCTAAGAAGTGGTAC         2698
```

```
FRI_Bål-2        TATGACTTAGCTGTTGGATACTAATTAAGCTTAAGCAATGCCAACTCTAAGAAGTGGTAC        2696
FRI_St-0         TATGACTTAGCTGTTGGATACTAATTAAGCTTAAGCAATGCCAACTCTAAGAAGTGGTAC        2625
FRI_Bg-2         TATGACTTAGCTGTTGGATACTAATTAAGCTTAAGCAATGCCAACTCTAAGAAGTGGTAC        2693
FRI_C24          TATGACTTAGCTGTTGGATACTAATTAAGCTTAAGCAATGCCAACTCTAAGAAGTGGTAC        2694
FRI_Van-0        TATGACTTAGCTGTTGGATACTAATTAAGCTTAAGCAATGCCAACTCTAAGAAGTGGTAC        2694
FRI_Ull-2-3      TATGACTTAGCTGTTGGATACTAATTAAGCTTAAGCAATGCCAACTCTAAGAAGTGGTAC        2697
FRI_Lip-0        TATGACTTAGCTGTTGGATACTAATTAAGCTTAAGCAATGCCAACTCTAAGAAGTGGTAC        2697
FRI_NFA-8        TATGACTTAGCTGTTGGATACTAATTAAGCTTAAGCAATGCCAACTCTAAGAAGTGGTAC        2697
FRI_Edi-0        TATGACTTAGCTGTTGGATACTAATTAAGCTTAAGCAATGCCAACTCTAAGAAGTGGTAC        2696
FRI_Ren-1        TATGACTTAGCTGTTGGATACTAATTAAGCTTAAGCAATGCCAACTCTAAGAAGTGGTAC        2325
FRI_Nok-3        TATGACTTAGCTGTTGGATACTAATTAAGCTTAAGCAATGCCAACTCTAAGAAGTGGTAC        2696
FRI_Sha          TATGACTTAGCTGTTGGATACTAATTAAGCTTAAGCAATGCCAACTCTAAGAAGTGGTAC        2696
FRI_Wa-1         TATGACTTAGCTGTTGGATACTAATTAAGCTTAAGCAATGCCAACTCTAAGAAGTGGTAC        2696
FRI_Spr-1-6      TATGACTTAGCTGTTGGATACTAATTAAGCTTAAGCAATGCCAACTCTAAGAAGTGGTAC        2696
FRI_Bil-7        TATGACTTAGCTGTTGGATACTAATTAAGCTTAAGCAATGCCAACTCTAAGAAGTGGTAC        2696
FRI_Alc-0        TATGACTTAGCTGTTGGATACTAATTAAGCTTAAGCAATGCCAACTCTAAGAAGTGGTAC        2696
FRI_Pu-2-23      TATGACTTAGCTGTTGGATACTAATTAAGCTTAAGCAATGCCAACTCTAAGAAGTGGTAC        2699
FRI_NFA-10       TATGACTTAGCTGTTGGATACTAATTAAGCTTAAGCAATGCCAACTCTAAGAAGTGGTAC        2698
FRI_Cvi-0        TATGACTTAGCTGTTGGATACTAATTAAGCTTAAGCAATGCCAACTCTAAGAAGTGGTAC        2699
FRI_Zdr-6        TATGACTTAGCTGTTGGATACTAATTAAGCTTAAGCAATGCCAACTCTAAGAAGTGGTAC        2699
                 ************************************************************

FRI_An-1         TTACACAATATTCTATTGGTCATAGGTATAGTTGAATCAAGTATCAAGCGTGGAATGCAT        2756
FRI_Pro-0        TTACACAATATTCTATTGGTCATAGGTATAGTTGAATCAAGTATCAAGCGTGGAATGCAT        2758
FRI_Bål-2        TTACACAATATTCTATTGGTCATAGGTATAGTTGAATCAAGTATCAAGCGTGGAATGCAT        2756
FRI_St-0         TTACACAATATTCTATTGGTCATAGGTATAGTTGAATCAAGTATCAAGCGTGGAATGCAT        2685
FRI_Bg-2         TTACACAATATTCTATTGGTCATAGGTATAGTTGAATCAAGTATCAAGCGTGGAATGCAT        2753
FRI_C24          TTACACAATATTCTATTGGTCATAGGTATAGTTGAATCAAGTATCAAGCGTGGAATGCAT        2754
FRI_Van-0        TTACACAATATTCTATTGGTCATAGGTATAGTTGAATCAAGTATCAAGCGTGGAATGCAT        2754
FRI_Ull-2-3      TTACACAATATTCTATTGGTCATAGGTATAGTTGAATCAAGTATCAAGCGTGGAATGCAT        2757
FRI_Lip-0        TTACACAATATTCTATTGGTCATAGGTATAGTTGAATCAAGTATCAAGCGTGGAATGCAT        2757
FRI_NFA-8        TTACACAATATTCTATTGGTCATAGGTATAGTTGAATCAAGTATCAAGCGTGGAATGCAT        2757
FRI_Edi-0        TTACACAATATTCTATTGGTCATAGGTATAGTTGAATCAAGTATCAAGCGTGGAATGCAT        2756
FRI_Ren-1        TTACACAATATTCTATTGGTCATAGGTATAGTTGAATCAAGTATCAAGCGTGGAATGCAT        2385
FRI_Nok-3        TTACACAATATTCTATTGGTCATAGGTATAGTTGAATCAAGTATCAAGCGTGGAATGCAT        2756
FRI_Sha          TTACACAATATTCTATTGGTCATAGGTATAGTTGAATCAAGTATCAAGCGTGGAATGCAT        2756
FRI_Wa-1         TTACACAATATTCTATTGGTCATAGGTATAGTTGAATCAAGTATCAAGCGTGGAATGCAT        2756
FRI_Spr-1-6      TTACACAATATTCTATTGGTCATAGGTATAGTTGAATCAAGTATCAAGCGTGGAATGCAT        2756
FRI_Bil-7        TTACACAATATTCTATTGGTCATAGGTATAGTTGAATCAAGTATCAAGCGTGGAATGCAT        2756
FRI_Alc-0        TTACACAATATTCTATTGGTCATAGGTATAGTTGAATCAAGTATCAAGCGTGGAATGCAT        2756
FRI_Pu-2-23      TTACACAATATTCTATTGGTCATAGGTATAGTTGAATCAAGTATCAAGCGTGGAATGCAT        2759
FRI_NFA-10       TTACACAATATTCTATTGGTCATAGGTATAGTTGAATCAAGTATCAAGCGTGGAATGCAT        2758
FRI_Cvi-0        TTACACAATATTCTATTGGTCATAGGTATAGTTGAATCAAGTATCAAGCGTGGAATGCAT        2759
FRI_Zdr-6        TTACACAATATTCTATTGGTCATAGGTATAGTTGAATCAAGTATCAAGCGTGGAATGCAT        2759
                 ************************************************************

FRI_An-1         ATTGAAGCTCTTGAGATGGTTTATACCTTTGGCATGGAGGATAAGTTTTCAGCTGCTCTA        2816
FRI_Pro-0        ATTGAAGCTCTTGAGATGGTTTATACCTTTGGCATGGAGGATAAGTTTTCAGCTGCTCTA        2818
FRI_Bål-2        ATTGAAGCTCTTGAGATGGTTTATACCTTTGGCATGGAGGATAAGTTTTCAGCTGCTCTA        2816
FRI_St-0         ATTGAAGCTCTTGAAATGGTTTATACCTTTGGCATGGAGGATAAGTTTTCAGCTGCTCTA        2745
FRI_Bg-2         ATTGAAGCTCTTGAAATGGTTTATACCTTTGGCATGGAGGATAAGTTTTCAGCTGCTCTA        2813
FRI_C24          ATTGAAGCTCTTGAAATGGTTTATACCTTTGGCATGGAGGATAAGTTTTCAGCTGCTCTA        2814
FRI_Van-0        ATTGAAGCTCTTGAGATGGTTTATACCTTTGGCATGGAGGATAAGTTTTCAGCTGCTCTA        2814
FRI_Ull-2-3      ATTGAAGCTCTTGAGATGGTTTATACCTTTGGCATGGAGGATAAGTTTTCAGCTGCTCTA        2817
FRI_Lip-0        ATTGAAGCTCTTGAGATGGTTTATACCTTTGGCATGGAGGATAAGTTTTCAGCTGCTCTA        2817
FRI_NFA-8        ATTGAAGCTCTTGAAATGGTTTATACCTTTGGCATGGAGGATAAGTTTTCAGCTGCTCTA        2817
FRI_Edi-0        ATTGAAGCTCTTGAAATGGTTTATACCTTTGGCATGGAGGATAAGTTTTCAGCTGCTCTA        2816
FRI_Ren-1        ATTGAAGCTCTTGAAATGGTTTATACCTTTGGCATGGAGGATAAGTTTTCAGCTGCTCTA        2445
FRI_Nok-3        ATTGAAGCTCTTGAAATGGTTTATACCTTTGGCATGGAGGATAAGTTTTCAGCTGCTCTA        2816
FRI_Sha          ATTGAAGCTCTTGAGATGGTTTATACCTTTGGCATGGAGGATAAGTTTTCAGCTGCTCTA        2816
FRI_Wa-1         ATTGAAGCTCTTGAGATGGTTTATACCTTTGGCATGGAGGATAAGTTTTCAGCTGCTCTA        2816
FRI_Spr-1-6      ATTGAAGCTCTTGAGATGGTTTATACCTTTGGCATGGAGGATAAGTTTTCAGCTGCTCTA        2816
FRI_Bil-7        ATTGAAGCTCTTGAGATGGTTTATACCTTTGGCATGGAGGATAAGTTTTCAGCTGCTCTA        2816
FRI_Alc-0        ATTGAAGCTCTTGAGATGGTTTATACCTTTGGCATGGAGGATAAGTTTTCAGCTGCTCTA        2816
FRI_Pu-2-23      ATTGAAGCTCTTGAGATGGTTTATACCTTTGGCATGGAGGATAAGTTTTCAGCTGCTCTA        2819
FRI_NFA-10       ATTGAAGCTCTTGAGATGGTTTATACCTTTGGCATGGAGGATAAGTTTTCAGCTGCTCTA        2818
FRI_Cvi-0        ATTGAAGCTCTTGAGATGGTTTATACCTTTGGCATGGAGGATAAGTTTTCAGCTGCTCTA        2819
FRI_Zdr-6        ATTGAAGCTCTTGAGATGGTTTATACCTTTGGCATCGAGGATAAGTTTTCAGCTGCTCTA        2819
                 ************** ******************** ************************

FRI_An-1         GTTCTAACTTCATTCTTAAAGATGAGCAAGGAGTCATTTGAGAGGGCAAAACGGAAAGCC        2876
```

```
FRI_Pro-0       GTTCTAACTTCATTCTTAAAGATGAGCAAGGAGTCATTTGAGAGGGCAAAACGGAAAGCC        2878
FRI_Bål-2       GTTCTAACTTCATTCTTAAAGATGAGCAAGGAGTCATTTGAGAGGGCAAAACGGAAAGCC        2876
FRI_St-0        GTTCTAACTTCATTCTTAAAGATGAGCAAGGAGTCATTTGAGAGGGCAAAACGGAAAGCC        2805
FRI_Bg-2        GTTCTAACTTCATTCTTAAAGATGAGCAAGGAGTCAtTTGAGAGGGCAAAACGGAAAGCC        2873
FRI_C24         GTTCTAACTTCATTCTTAAAGATGAGCAAGGAGTCATTTGAGAGGGCAAAACGGAAAGCC        2874
FRI_Van-0       GTTCTAACTTCATTCTTAAAGATGAGCAAGGAGTCATTTGAGAGGGCAAAACGGAAAGCC        2874
FRI_Ull-2-3     GTTCTAACTTCATTCTTAAAGATGAGCAAGGAGTCATTTGAGAGGGCAAAACGGAAAGCC        2877
FRI_Lip-0       GTTCTAACTTCATTCTTAAAGATGAGCAAGGAGTCATTTGAGAGGGCAAAACGGAAAGCC        2877
FRI_NFA-8       GTTCTAACTTCATTCTTAAAGATGAGCAAGGAGTCATTTGAGAGGGCAAAACGGAAAGCC        2877
FRI_Edi-0       GTTCTAACTTCATTCTTAAAGATGAGCAAGGAGTCATTTGAGAGGGCAAAACGGAAAGCC        2876
FRI_Ren-1       GTTCTAACTTCATTCTTAAAGATGAGCAAGGAGTCATTTGAGAGGGCAAAACGGAAAGCC        2505
FRI_Nok-3       GTTCTAACTTCATTCTTAAAGATGAGCAAGGAGTCATTTGAGAGGGCAAAACGGAAAGCC        2876
FRI_Sha         GTTCTAACTTCATTCTTAAAGATGAGCAAGGAGTCATTTGAGAGGGCAAAACGGAAAGCC        2876
FRI_Wa-1        GTTCTAACTTCATTCTTAAAGATGAGCAAGGAGTCATTTGAGAGGGCAAAACGGAAAGCC        2876
FRI_Spr-1-6     GTTCTAACTTCATTCTTAAAGATGAGCAAGGAGTCATTTGAGAGGGCAAAACGGAAAGCC        2876
FRI_Bil-7       GTTCTAACTTCATTCTTAAAGATGAGCAAGGAGTCATTTGAGAGGGCAAAACGGAAAGCC        2876
FRI_Alc-0       GTTCTAACTTCATTCTTAAAGATGAGCAAGGAGTCATTTGAGAGGGCAAAACGGAAAGCC        2876
FRI_Pu-2-23     GTTCTAACTTCATTCTTAAAGATGAGCAAGGAATCATTTGAGAGGGCAAAACGGAAAGCC        2879
FRI_NFA-10      GTTCTAACTTCATTCTTAAAGATGAGCAAGGAGTCATTTGAGAGGGCAAAACGGAAAGCC        2878
FRI_Cvi-0       GTTCTAACTTCATTCTTAAAGATGAGCAAGGAGTCATTTGAGAGGGCAAAACGGAAAGCC        2879
FRI_Zdr-6       GTTCTAACTTCATTCTTAAAGATGAGCAAGGAGTCATTTGAGAGGGCAAAACGGAAAGCC        2879
                ******************************** ***************************

FRI_An-1        CAGTCACCGCTGGCATTTGTATGAACCCTTCCCTTGCACATTATGTACCTTTATGAACTC        2936
FRI_Pro-0       CAGTCACCGCTGGCATTTGTATGAACCCTTCCCTTGCACATTATGTACCTTTATGAACTC        2938
FRI_Bål-2       CAGTCACCGCTGGCATTTGTATGAACCCTTCCCTTGCACATTATGTACCTTTATGAACTC        2936
FRI_St-0        CAGTCACCGCTGGCATTTGTATGAACCCTTCCCTTGCACATTATGTACCTTTATGAACTC        2865
FRI_Bg-2        CAGTCACCGCTGGCATTTGTATGAACCCTTCCCTTGCACATTATGTACCTTTATGAACTC        2933
FRI_C24         CAGTCACCGCTGGCATTTGTATGAACCCTTCCCTTGCACATTATGTACCTTTATGAACTC        2934
FRI_Van-0       CAGTCACCGCTGGCATTTGTATGAACCCTTCCCTTGCACATTATGTACCTTTATGAACTC        2934
FRI_Ull-2-3     CAGTCACCGCTG----TTGTATGAACCCTTCCCTTGCACATTATGTACCTTTATGAACTC        2933
FRI_Lip-0       CAGTCACCGCTGGCATTTGTATGAACCCTTCCCTTGCACATTATGTACCTTTATGAACTC        2937
FRI_NFA-8       CAGTCACCGCTGGCATTTGTATGAACCCTTCCCTTGCACATTATGTACCTTTATGAACTC        2937
FRI_Edi-0       CAGTCACCGCTGGCATTTGTATGAACCCTTCCCTTGCACATTATGTACCTTTATGAACTC        2936
FRI_Ren-1       CAGTCACCGCTGGCATTTGTATGAACCCTTCCCTTGCACATTATGTACCTTTATGAACTC        2565
FRI_Nok-3       CAGTCACCGCTGGCATTTGTATGAACCCTTCCCTTGCACATTATGTACCTTTATGAACTC        2936
FRI_Sha         CAGTCACCGCTGGCATTTGTATGAACCCTTCCCTTGCACATTATGTACCTTTATGAACTC        2936
FRI_Wa-1        CAGTCACCGCTGGCATTTGTATGAACCCTTCCCTTGCACATTATGTACCTTTATGAACTC        2936
FRI_Spr-1-6     CAGTCACCGCTGGCATTTGTATGAACCCTTCCCTTGCACATTATGTACCTTTATGAACTC        2936
FRI_Bil-7       CAGTCACCGCTGGCATTTGTATGAACCCTTCCCTTGCACATTATGTACCTTTATGAACTC        2936
FRI_Alc-0       CAGTCACCGCTGGCATTTGTATGAACCCTTCCCTTGCACATTATGTACCTTTATGAACTC        2936
FRI_Pu-2-23     CAGTCACCGCTGGCATTTGTATGAACCCTTCCCTTGCACATTATGTACCTTTATGAACTC        2939
FRI_NFA-10      CAGTCACCGCTGGCATTTGTATGAACCCTTCCCTTGCACATTATGTACCTTTATGAACTC        2938
FRI_Cvi-0       CAGTCACCGCTGGCATTTGTATGAACCCTTCCCTTGCACATTATGTACCTTTATGAACTC        2939
FRI_Zdr-6       CAGTCACCGCTGGCATTTGTATGAACCCTTCCCTTGCACATTATGTACCTTTATGAACTC        2939
                ************    ********************************************

FRI_An-1        TTTATCATCATCTGAGTCTGACCATTGATATATTTATTTCTCAACAGAAAGAAGCGGCTA        2996
FRI_Pro-0       TTTATCATCATCTGAGTCTGACCATTGATATATTTATTTCTCAACAGAAAGAAGCGGCTA        2998
FRI_Bål-2       TTTATCATCATCTGAGTCTGACCATTGATATATTTATTTCTCAACAGAAAGAAGCGGCTA        2996
FRI_St-0        TTTATCATCATCTGAGTCTGACCATTGATATATTTATTTCTCAACAGAAAGAAGCGGCTA        2925
FRI_Bg-2        TTTATCATCATCTGAGTCTGACCATTGATATATTTATTTCTCAACAGAAAGAAGCGGCTA        2993
FRI_C24         TTTATCATCATCTGAGTCTGACCATTGATATATTTATTTCTCAACAGAAAGAAGCGGCTA        2994
FRI_Van-0       TTTATCATCATCTGAGTCTGACCATTGATATATTTATTTCTCAACAGAAAGAAGCGGCTA        2994
FRI_Ull-2-3     TTTATCATCATCTGAGTCTGACCATTGATATATTTATTTCTCAACAGAAAGAAGCGGCTA        2993
FRI_Lip-0       TTTATCATCATCTGAGTCTGACCATTGATATATTTATTTCTCAACAGAAAGAAGCGGCTA        2997
FRI_NFA-8       TTTATCATCATCTGAGTCTGACCATTGATATATTTATTTCTCAACAGAAAGAAGCGGCTA        2997
FRI_Edi-0       TTTATCATCATCTGAGTCTGACCATTGATATATTTATTTCTCAACAGAAAGAAGCGGCTA        2996
FRI_Ren-1       TTTATCATCATCTGAGTCTGACCATTGATATATTTATTTCTCAACAGAAAGAAGCGGCTA        2625
FRI_Nok-3       TTTATCATCATCTGAGTCTGACCATTGATATATTTATTTCTCAACAGAAAGAAGCGGCTA        2996
FRI_Sha         TTTATCATCATCTGAGTCTGACCATTGATATATTTATTTCTCAACAGAAAGAAGCGGCTA        2996
FRI_Wa-1        TTTATCATCATCTGAGTCTGACCATTGATATATTTATTTCTCAACAGAAAGAAGCGGCTA        2996
FRI_Spr-1-6     TTTATCATCATCTGAGTCTGACCATTGATATATTTATTTCTCAACAGAAAGAAGCGGCTA        2996
FRI_Bil-7       TTTATCATCATCTGAGTCTGACCATTGATATATTTATTTCTCAACAGAAAGAAGCGGCTA        2996
FRI_Alc-0       TTTATCATCATCTGAGTCTGACCATTGATATATTTATTTCTCAACAGAAAGAAGCGGCTA        2996
FRI_Pu-2-23     TTTATCATCATCTGAGTCTGACCATTGATATATTTATTTCTCAACAGAAAGAAGCGGCTA        2999
FRI_NFA-10      TTTATCATCATCTGAGTCTGACCATTGATATATTTATTTCTCAACAGAAAGAAGCGGCTA        2998
FRI_Cvi-0       TTTATCATCATCTGAGTCTGACCATTGATATATTTATTTCTCAACAGCAAGAAGCGGCTA        2999
FRI_Zdr-6       TTTATCATCATCTGAGTCTGACCATTGATATATTTATTTCTCAACAGAAAGAAGCGGCTA        2999
                *********************************************** ************
```

```
FRI_An-1        CAAAGCAGCTAGCTGTGTTATCATCAGTTATGCAGTGTATGGAGACTCACAAGTTAGATC    3056
FRI_Pro-0       CAAAGCAGCTAGCTGTGTTATCATCAGTTATGCAGTGTATGGAGACTCACAAGTTAGATC    3058
FRI_Bål-2       CAAAGCAGCTAGCTGTGTTATCATCAGTTATGCAGTGTATGGAGACTCACAAGTTAGATC    3056
FRI_St-0        CAAAGCAGCTAGCTGTGTTATCATCAGTTATGCAGTGTATGGAGACTCACAAGTTAGATC    2985
FRI_Bg-2        CAAAGCAGCTAGCTGTGTTATCATCAGTTATGCAGTGTATGGAGACTCACAAGTTAGATC    3053
FRI_C24         CAAAGCAGCTAGCTGTGTTATCATCAGTTATGCAGTGTATGGAGACTCACAAGTTAGATC    3054
FRI_Van-0       CAAAGCAGCTAGCTGTGTTATCATCAGTTATGCAGTGTATGGAGACTCACAAGTTAGATC    3054
FRI_Ull-2-3     CAAAGCAGCTAGCTGTGTTATCATCAGTTATGCAGTGTATGGAGACTCACAAGTTAGATC    3053
FRI_Lip-0       CAAAGCAGCTAGCTGTGTTATCATCAGTTATGCAGTGTATGGAGACTCACAAGTTAGATC    3057
FRI_NFA-8       CAAAGCAGCTAGCTGTGTTATCATCAGTTATGCAGTGTATGGAGACTCACAAGTTAGATC    3057
FRI_Edi-0       CAAAGCAGCTAGCTGTGTTATCATCAGTTATGCAGTGTATGGAGACTCACAAGTTAGATC    3056
FRI_Ren-1       CAAAGCAGCTAGCTGTGTTATCATCAGTTATGCAGTGTATGGAGACTCACAAGTTAGATC    2685
FRI_Nok-3       CAAAGCAGCTAGCTGTGTTATCATCAGTTATGCAGTGTATGGAGACTCACAAGTTAGATC    3056
FRI_Sha         CAAAGCAGCTAGCTGTGTTATCATCAGTTATGCAGTGTATGGAGACTCACAAGTTAGATC    3056
FRI_Wa-1        CAAAGCAGCTAGCTGTGTTATCATCAGTTATGCAGTGTATGGAGACTCACAAGTTAGATC    3056
FRI_Spr-1-6     CAAAGCAGCTAGCTGTGTTATCATCAGTTATGCAGTGTATGGAGACTCACAAGTTAGATC    3056
FRI_Bil-7       CAAAGCAGCTAGCTGTGTTATCATCAGTTATGCAGTGTATGGAGACTCACAAGTTAGATC    3056
FRI_Alc-0       CAAAGCAGCTAGCTGTGTTATCATCAGTTATGCAGTGTATGGAGACTCACAAGTTAGATC    3056
FRI_Pu-2-23     CAAAGCAGCTAGCTGTGTTATCATCAGTTATGCAGTGTATGGAGACTCACAAGTTAGATC    3059
FRI_NFA-10      CAAAGCAGCTAGCTGTGTTATCATCAGTTATGCAGTGTATGGAGACTCACAAGTTAGATC    3058
FRI_Cvi-0       CAAAGCAGCTAGCTGTGTTATCATCAGTTATGCAGTGTATGGAGACTCACAAGTTAGATC    3059
FRI_Zdr-6       CAAAGCAGCTAGCTGTGTTATCATCAGTTATGCAGTGTATGGAGACTCACAAGTTAGATC    3059
                ************************************************************

FRI_An-1        TTGCTTTCTCTTCCATCTCTTTGTCGAGCTGA-------------------AGAGT----    3093
FRI_Pro-0       TTGCTTTCTCTTCCATCTCTTTGTCGAGCTGA-------------------AGAGT----    3095
FRI_Bål-2       CTGCGAAAGAACTACCAGGATGGCAGATCAAAGAGCAAATTGTTAGCTTGGAGAAAGACA    3116
FRI_St-0        CTGCGAAAGAACTACCAGGATGGCAGATCAAAGAGCAAATTGTTAGCTTGGAGAAAGACA    3045
FRI_Bg-2        CTGCGAAAGAACTACCAGGATGGCAGATCAAAGAGCAAATTGTTAGCTTGGAGAAAGACA    3113
FRI_C24         CTGCGAAAGAACTACCAGGATGGCAGATCAAAGAGCAAATTGTTAGCTTGGAGAAAGACA    3114
FRI_Van-0       CTGCGAAAGAACTACCAGGATGGCAGATCAAAGAGCAAATTGTTAGCTTGGAGAAAGACA    3114
FRI_Ull-2-3     CTGCGAAAGAACTACCAGGATGGCAGATCAAAGAGCAAATTGTTAGCTTGGAGAAAGACA    3113
FRI_Lip-0       CTGCGAAAGAACTACCAGGATGGCAGATCAAAGAGCAAATTGTTAGCTTGGAGAAAGACA    3117
FRI_NFA-8       CTGCGAAAGAACTACCAGGATGGCAGATCAAAGAGCAAATTGTTAGCTTGGAGAAAGACA    3117
FRI_Edi-0       CTGCGAAAGAACTACCAGGATGGCAGATCAAAGAGCAAATTGTTAGCTTGGAGAAAGACA    3116
FRI_Ren-1       CTGCGAAAGAACTACCAGGATGGCAGATCAAAGAGCAAATTGTTAGCTTGGAGAAAGACA    2745
FRI_Nok-3       CTGCGAAAGAACTACCAGGATGGCAGATCAAAGAGCAAATTGTTAGCTTGGAGAAAGACA    3116
FRI_Sha         CTGCGAAAGAACTACCAGGATGGCAGATCAAAGAGCAAATTGTTAGCTTGGAGAAAGACA    3116
FRI_Wa-1        CTGCGAAAGAACTACCAGGATGGCAGATCAAAGAGCAAATTGTTAGCTTGGAGAAAGACA    3116
FRI_Spr-1-6     CTGCGAAAGAACTACCAGGATGGCAGATCAAAGAGCAAATTGTTAGCTTGGAGAAAGACA    3116
FRI_Bil-7       CTGCGAAAGAACTACCAGGATGGCAGATCAAAGAGCAAATTGTTAGCTTGGAGAAAGACA    3116
FRI_Alc-0       CTGCGAAAGAACTACCAGGATGGCAGATCAAAGAGCAAATTGTTAGCTTGGAGAAAGACA    3116
FRI_Pu-2-23     CTGCGAAAGAACTACCAGGATGGCAGATCAAAGAGCAAATTGTTAGCTTGGAGAAAGACA    3119
FRI_NFA-10      CTGCGAAAGAACTACCAGGATGGCAGATCAAAGAGCAAATTGTTAGCTTGGAGAAAGACA    3118
FRI_Cvi-0       cTGCGAAAGAACTACCAGGATGGCAGATCAAAGAGCAAATTGTTAGCTTGGAGAAAGACA    3119
FRI_Zdr-6       CTGCGAAAGAACTACCAGGATGGCAGATCAAAGAGCAAATTGTTAGCTTGGAGAAAGACA    3119
                 ***                * *  ** *  *                   ***

FRI_An-1        ---------------GTCTTTCTCCAAGCTAACAATTTGCTCTCAGTTTAATGGAGGAAG    3138
FRI_Pro-0       ---------------GTCTTTCTCCAAGCTAACAATTTGCTCTCAGTTTAATGGAGGAAG    3140
FRI_Bål-2       CTCTTCAGCTCGACAAAGAGATGGAAGAGAAAGCAAGATCTCTCAGTTTAATGGAGGAAG    3176
FRI_St-0        CTCTTCAGCTCGACAAAGAGATGGAAGAGAAAGCAAGATCTCTCAGTTTAATGGAGGAAG    3105
FRI_Bg-2        CTCTTCAGCTCGACAAAGAGATGGAAGAGAAAGCAAGATCTCTCAGTTTAATGGAGGAAG    3173
FRI_C24         CTCTTCAGCTCGACAAAGAGATGGAAGAGAAAGCAAGATCTCTCAGTTTAATGGAGGAAG    3174
FRI_Van-0       CTCTTCAGCTCGACAAAGAGATGGAAGAGAAAGCAAGATCTCTCAGTTTAATGGAGGAAG    3174
FRI_Ull-2-3     CTCTTCAGCTCGACAAAGAGATGGAAGAGAAAGCAAGATCTCTCAGTTTAATGGAGGAAG    3173
FRI_Lip-0       CTCTTCAGCTCGACAAAGAGATGGAAGAGAAAGCAAGATCTCTCAGTTTAATGGAGGAAG    3177
FRI_NFA-8       CTCTTCAGCTCGACAAAGAGATGGAAGAGAAAGCAAGATCTCTCAGTTTAATGGAGGAAG    3177
FRI_Edi-0       CTCTTCAGCTCGACAAAGAGATGGAAGAGAAAGCAAGATCTCTCAGTTTAATGGAGGAAG    3176
FRI_Ren-1       CTCTTCAGCTCGACAAAGAGATGGAAGAGAAAGCAAGATCTCTCAGTTTAATGGAGGAAG    2805
FRI_Nok-3       CTCTTCAGCTCGACAAAGAGATGGAAGAGAAAGCAAGATCTCTCAGTTTAATGGAGGAAG    3176
FRI_Sha         CTCTTCAGCTCGACAAAGAGATGGAAGAGAAAGCAAGATCTCTCAGTTTAATGGAGGAAG    3176
FRI_Wa-1        CTCTTCAGCTCGACAAAGAGATGGAAGAGAAAGCAAGATCTCTCAGTTTAATGGAGGAAG    3176
FRI_Spr-1-6     CTCTTCAGCTCGACAAAGAGATGGAAGAGAAAGCAAGATCTCTCAGTTTAATGGAGGAAG    3176
FRI_Bil-7       CTCTTCAGCTCGACAAAGAGATGGAAGAGAAAGCAAGATCTCTCAGTTTAATGGAGGAAG    3176
FRI_Alc-0       CTCTTCAGCTCGACAAAGAGATGGAAGAGAAAGCAAGATCTCTCAGTTTAATGGAGGAAG    3176
FRI_Pu-2-23     CTCTTCAGCTCGACAAAGAGATGGAAGAGAAAGCAAGATCTCTCAGTTTAATGGAGGAAG    3179
FRI_NFA-10      CTCTTCAGCTCGACAAAGAGATGGAAGAGAAAGCAAGATCTCTCAGTTTAATGGAGGAAG    3178
FRI_Cvi-0       CTCTTCAGCTCGACAAAGAGATGGAAGAGAAAGCAAGATCTCTCAGTTTAATGGAGGAAG    3179
FRI_Zdr-6       CTCTTCAGCTCGACAAAGAGATGGAAGAGAAAGCAAGATCTCTCAGTTTAATGGAGGAAG    3179
                          *    **  *     ********************
```

```
FRI_An-1        CCGCACTTGCCAAGAGAATGTATAACCAACAGATAAAACGTCCAAGGTTGTCACCCATGG    3198
FRI_Pro-0       CCGCACTTGCCAAGAGAATGTATAACCAACAGATAAAACGTCCAAGGTTGTCACCCATGG    3200
FRI_Bål-2       CCGCACTTGCCAAGAGAATGTATAACCAACAGATAAAACGTCCAAGGTTGTCACCCATGG    3236
FRI_St-0        CCGCACTTGCCAAGAGAATGTATAACCAACAGATAAAACGTCCAAGGTTGTCACCCATGG    3165
FRI_Bg-2        CCGCACTTGCCAAGAGAATGTATAACCAACAGATAAAACGTCCAAGGTTGTCACCCATGG    3233
FRI_C24         CCGCACTTGCCAAGAGAATGTATAACCAACAGATAAAACGTCCAAGGTTGTCACCCATGG    3234
FRI_Van-0       CCGCACTTGCCAAGAGAATGTATAACCAACAGATAAAACGTCCAAGGTTGTCACCCATGG    3234
FRI_Ull-2-3     CCGCACTTGCCAAGAGAATGTATAACCAACAGATAAAACGTCCAAGGTTGTCACCCATGG    3233
FRI_Lip-0       CTGCACTTGCCAAGAGAATGTATAACCAACAGATAAAACGTCCAAGGTTGTCACCCATGG    3237
FRI_NFA-8       CCGCACTTGCCAAGAGAATGTATAACCAACAGATAAAACGTCCAAGGTTGTCACCCATGG    3237
FRI_Edi-0       CCGCACTTGCCAAGAGAATGTATAACCAACAGATAAAACGTCCAAGGTTATCACCCATGG    3236
FRI_Ren-1       CCGCACTTGCCAAGAGAATGTATAACCAACAGATAAAACGTCCAAGGTTGTCACCCATGG    2865
FRI_Nok-3       CCGCACTTGCCAAGAGAATGTATAACCAACAGATAAAACGTCCAAGGTTGTCACCCATGG    3236
FRI_Sha         CCGCACTTGCCAAGAGAATGTATAACCAACAGATAAAACGTCCAAGGTTGTCACCCATGG    3236
FRI_Wa-1        CCGCACTTGCCAAGAGAATGTATAACCAACAGATAAAACGTCCAAGGTTGTCACCCATGG    3236
FRI_Spr-1-6     CCGCACTTGCCAAGAGAATGTATAACCAACAGATAAAACGTCCAAGGTTGTCACCCATGG    3236
FRI_Bil-7       CCGCACTTGCCAAGAGAATGTATAACCAACAGATAAAACGTCCAAGGTTGTCACCCATGG    3236
FRI_Alc-0       CCGCACTTGCCAAGAGAATGTATAACCAACAGATAAAACGTCCAAGGTTGTCACCCATGG    3236
FRI_Pu-2-23     CCGCACTTGCCAAGAGAATGTATAACCAACAGATAAAACGTCCAAGGTTGTCACCCATGG    3239
FRI_NFA-10      CCGCACTTGCCAAGAGAATGTATAACCAACAGATAAAACGTCCAAGGTTGTCACCCATGG    3238
FRI_Cvi-0       CCGCACTTGCCAAGAGAATGTATAACCAACAGATAAAACGTCCAAGGTTGTCACCCATGG    3239
FRI_Zdr-6       CCGCACTTGCCAAGAGAATGTATAACCAACAGATAAAACGTCCAAGGTTGTCACCCATGG    3239
                * *********************************************** **********

FRI_An-1        AAATGCCACCAGTAACTTCTTCATCGTATTCTCCTATCTACCGTGATAGAAGCTTTCCTA    3258
FRI_Pro-0       AAATGCCACCAGTAACTTCTTCATCGTATTCTCCTATCTACCGTGATAGAAGCTTTCCTA    3260
FRI_Bål-2       AAATGCCACCAGTAACTTCTTCATCGTATTCTCCTATCTACCGTGATAGAAGCTTTCCTA    3296
FRI_St-0        AAATGCCACCAGTAACTTCTTCATCGTATTCTCCTATCTACCGTGATAGAAGCTTTCCTA    3225
FRI_Bg-2        AAATGCCACCAGTAACTTCTTCATCGTATTCTCCTATCTACCGTGATAGAAGCTTTCCTA    3293
FRI_C24         AAATGCCACCAGTAACTTCTTCATCGTATTCTCCTATCTACCGTGATAGAAGCTTTCCTA    3294
FRI_Van-0       AAATGCCACCAGTAACTTCTTCATCATATTCTCCTATCTACCGTGATAGAAGCTTTCCTA    3294
FRI_Ull-2-3     AAATGCCACCAGTAACTTCTTCATCGTATTCTCCTATCTACCGTGATAGAAGCTTTCCTA    3293
FRI_Lip-0       AAATGCCACCAGTAACTTCTTCATCGTATTCTCCTATCTACCGTGATAGAAGCTTTCCTA    3297
FRI_NFA-8       AAATGCCACCAGTAACTTCTTCATCGTATTCTCCTATCTACCGTGATAGAAGCTTTCCTA    3297
FRI_Edi-0       AAATGCCACCAGTAACTTCTTCATCGTATTCTCCTATCTACCGTGATAGAAGCTTTCCTA    3296
FRI_Ren-1       AAATGCCACCAGTAACTTCTTCATCGTATTCTCCTATCTACCGTGATAGAAGCTTTCCTA    2925
FRI_Nok-3       AAATGCCACCAGTAACTTCTTCATCGTATTCTCCTATCTACCGTGATAGAAGCTTTCCTA    3296
FRI_Sha         AAATGCCACCAGTAACTTCTTCATCGTATTCTCCTATCTACCGTGATAGAAGCTTTCCTA    3296
FRI_Wa-1        AAATGCCACCAGTAACTTCTTCATCGTATTCTCCTATCTACCGTGATAGAAGCTTTCCTA    3296
FRI_Spr-1-6     AAATGCCACCAGTAACTTCTTCATCGTATTCTCCTATCTACCGTGATAGAAGCTTTCCTA    3296
FRI_Bil-7       AAATGCCACCAGTAACTTCTTCATCGTATTCTCCTATCTACCGTGATAGAAGCTTTCCTA    3296
FRI_Alc-0       AAATGCCACCAGTAACTTCTTCATCGTATTCTCCTATCTACCGTGATAGAAGCTTTCCTA    3296
FRI_Pu-2-23     AAATGCCACCAGTAGCTTCTTCATCGTATTCTCCTATCTACCGTGATAGAAGCTTTCCTA    3299
FRI_NFA-10      AAATGCCACCAGTAACTTCTTCATCGTATTCTCCTATCTACCGTGATAGAAGCTTTCCTA    3298
FRI_Cvi-0       AAATGCCACCAGTAACTTCTTCATCGTATTCTCCTATCTACCGTGATAGAAGCTTTCCTA    3299
FRI_Zdr-6       AAATGCCACCAGTAACTTCTTCATCGTATTCTCCTATCTACCGTGATAGAAGCTTTCCTA    3299
                ************** ********** **********************************

FRI_An-1        GTCAAAGAGACGATGACCAAGATGAA-ATATCAGCTCTTGTGAGTAGTTACCTCGGCCCG    3317
FRI_Pro-0       GTCAAAGAGACGATGACCAAGATGAA-ATATCAGCTCTTGTGAGTAGTTACCTCGGCCCG    3319
FRI_Bål-2       GTCAAAGAGACGATGACCAAGATGAA-ATATCAGCTCTTGTGAGTAGTTACCTCGGCCCG    3355
FRI_St-0        GTCAAAGAGACGATGACCAAGATGAA-ATATCAGCTCTTGTGAGTAGTTACCTCGGCCCG    3284
FRI_Bg-2        GTCAAAGAGACGATGACCAAGATGAA-ATATCAGCTCTTGTGAGTAGTTACCTCGGCCCG    3352
FRI_C24         GTCAAAGAGACGATGACCAAGATGAA-ATATCAGCTCTTGTGAGTAGTTACCTCGGCCCG    3353
FRI_Van-0       GTCAAAGAGACGATGACCAAGATGAA-ATATCAGCTCTTGTGAGTAGTTACCTCGGCCCG    3353
FRI_Ull-2-3     GTCAAAGAGACGATGACCAAGATGAA-ATATCAGCTCTTGTGAGTAGTTACCTCGGCCCG    3352
FRI_Lip-0       GTCAAAGAGACGATGACCAAGATGAA-ATATCAGCTCTTGTGAGTAGTTACCTCGGCCCG    3356
FRI_NFA-8       GTCAAAGAGACGATGACCAAGATGAA-ATATCAGCTCTTGTGAGTAGTTACCTCGGCCCG    3356
FRI_Edi-0       GTCAAAGAGACGATGACCAAGATGAA-ATATCAGCTCTTGTGAGTAGTTACCTCGGCCCG    3355
FRI_Ren-1       GTCAAAGAGACGATGACCAAGATGAA-ATATCAGCTCTTGTGAGTAGTTACCTCGGCCCG    2984
FRI_Nok-3       GTCAAAGAGACGATGACCAAGATGAA-ATATCAGCTCTTGTGAGTAGTTACCTCGGCCCG    3355
FRI_Sha         GTCAAAGAGACGATGACCAAGATGAA-ATATCAGCTCTTGTGAGTAGTTACCTCGGCCCG    3355
FRI_Wa-1        GTCAAAGAGACGATGACCAAGATGAA-ATATCAGCTCTTGTGAGTAGTTACCTCGGCCCG    3355
FRI_Spr-1-6     GTCAAAGAGACGATGACCAAGATGAA-ATATCAGCTCTTGTGAGTAGTTACCTCGGCCCG    3355
FRI_Bil-7       GTCAAAGAGACGATGACCAAGATGAA-ATATCAGCTCTTGTGAGTAGTTACCTCGGCCCG    3355
FRI_Alc-0       GTCAAAGAGACGATGACCAAGATGAA-ATATCAGCTCTTGTGAGTAGTTACCTCGGCCCG    3355
FRI_Pu-2-23     GTCAAAGAGACGATGACCAAGATGAA-ATATCAGCTCTTGTGAGTAGTTACCTCGGCCCG    3358
FRI_NFA-10      GTCAAAGAGACGATGACCAAGATGAAATTATCAGCTCTTGTGAGTAGTTACCTCGGCCCG    3358
FRI_Cvi-0       GTCAAAGAGACGATGACCAAGATGAA-ATATCAGCTCTTGTGAGTAGTTACCTCGGCCCG    3358
FRI_Zdr-6       GTCAAAGAGACGATGACCAAGATGAA-ATATCAGCTCTTGTGAGTAGTTACCTCGGCCCG    3358
```

```
                 **************************  ********************************

FRI_An-1         TCAACATCTTTTCCTCATCGCTCAAGAAGATCCCCGGAATATATGGTTCCACTTCCACAT         3377
FRI_Pro-0        TCAACATCTTTTCCTCATCGCTCAAGAAGATCCCCGGAATATATGGTTCCACTTCCACAT         3379
FRI_Bål-2        TCAACATCTTTTCCTCATCGCTCAAGAAGATCCCCGGAATATATGGTTCCACTTCCACAT         3415
FRI_St-0         TCAACATCTTTTCCTCATCGCTCAAGAAGATCCCCGGAATATATGGTTCCACTTCCACAT         3344
FRI_Bg-2         TCAACATCTTTTCCTCATCGCTCAAGAAGATCCCCGGAATATATGGTTCCACTTCCACAT         3412
FRI_C24          TCAACATCTTTTCCTCATCGCTCAAGAAGATCCCCGGAATATATGGTTCCACTTCCACAT         3413
FRI_Van-0        TCAACATCTTTTCCTCATCGCTCAAGAAGATCCCCGGAATATATGGTTCCACTTCCACAT         3413
FRI_Ull-2-3      TCAACATCTTTTCCTCATCGCTCAAGAAGATCCCCGGAATATATGGTTCCACTTCCACAT         3412
FRI_Lip-0        TCAACATCTTTTCCTCATCGCTCAAGAAGATCCCCGGAATATATGGTTCCACTTCCACAT         3416
FRI_NFA-8        TCAACATCTTTTCCTCATCGCTCAAGAAGATCCCCGGAATATATGGTTCCACTTCCACAT         3416
FRI_Edi-0        TCAACATCTTTTCCTCATCGCTCAAGAAGATCCCCGGAATATATGGTTCCACTTCCACAT         3415
FRI_Ren-1        TCAACATCTTTTCCTCATCGCTCAAGAAGATCCCCGGAATATATGGTTCCACTTCCACAT         3044
FRI_Nok-3        TCAACATCTTTTCCTCATCGCTCAAGAAGATCCCCGGAATATATGGTTCCACTTCCACAT         3415
FRI_Sha          TCAACATCTTTTCCTCATCGCTCAAGAAGATCCCCGGAATATATGGTTCCACTTCCACAT         3415
FRI_Wa-1         TCAACATCTTTTCCTCATCGCTCAAGAAGATCCCCGGAATATATGGTTCCACTTCCACAT         3415
FRI_Spr-1-6      TCAACATCTTTTCCTCATCGCTCAAGAAGATCCCCGGAATATATGGTTCCACTTCCACAT         3415
FRI_Bil-7        TCAACATCTTTTCCTCATCGCTCAAGAAGATCCCCGGAATATATGGTTCCACTTCCACAT         3415
FRI_Alc-0        TCAACATCTTTTCCTCATCGCTCAAGAAGATCCCCGGAATATATGGTTCCACTTCCACAT         3415
FRI_Pu-2-23      TCAACATCTTTTCCTCATCGCTCAAGAAGATCCCCGGAATATATGGTTCCACTTCCACAT         3418
FRI_NFA-10       TCAACATCTTTTCCTCATCGCTCAAGAAGATCCCCGGAATATATGGTTCCACTTCCACAT         3418
FRI_Cvi-0        TCAACATCTTTTCCTCATCGCTCAAGAAGATCCCCGGAATATATGGTTCCACTTCCACAT         3418
FRI_Zdr-6        TCAACATCTTTTCCTCATCGCTCAAGAAGATCCCCGGAATATATGGTTCCACTTCCACAT         3418
                 ************************************************************

FRI_An-1         GGTGGGTTAGGAAGAAGTGTATATGCATATGAACATCTGGCCCCAAATTCATACTCTCCA         3437
FRI_Pro-0        GGTGGGTTAGGAAGAAGTGTATATGCATATGAACATCTGGCCCCAAATTCATACTCTCCA         3439
FRI_Bål-2        GGTGGGTTAGGAAGAAGTGTATATGCATATGAACATCTGGCCCCAAATTCATACTCTCCA         3475
FRI_St-0         GGTGGGTTAGGAAGAAGTGTATATGCATATGAACATCTGGCCCCAAATTCATATTCTCCA         3404
FRI_Bg-2         GGTGGGTTAGGAAGAAGTGTATATGCATATGAACATCTGGCCCCAAATTCATATTCTCCA         3472
FRI_C24          GGTGGGTTAGGAAGAAGTGTATATGCATATGAACATCTGGCCCCAAATTCATATTCTCCA         3473
FRI_Van-0        GGTGGGTTAGGAAGAAGTGTATATGCATATGAACATCTGGCCCCAAATTCATACTCTCCA         3473
FRI_Ull-2-3      GGTGGGTTAGGAAGAAGTGTATATGCATATGAACATCTGGCCCCAAATTCATACTCTCCA         3472
FRI_Lip-0        GGTGGGTTAGGAAGAAGTGTATATGCATATGAACATCTGGCCCCAAATTCATACTCTCCA         3476
FRI_NFA-8        GGTGGGTTAGGAAGAAGTGTATATGCATATGAACATCTGGCCCCAAATTCATATTCTCCA         3476
FRI_Edi-0        GGTGGGTTAGGAAGAAGTGTATATGCATATGAACATCTGGCCCCAAATTCATATTCTCCA         3475
FRI_Ren-1        GGTGGGTTAGGAAGAAGTGTATATGCATATGAACATCTGGCCCCAAATTCATACTCTCCA         3104
FRI_Nok-3        GGTGGGTTAGGAAGAAGTGTATATGCATATGAACATCTGGCCCCAAATTCATACTCTCCA         3475
FRI_Sha          GGTGGGTTAGGAAGAAGTGTATATGCATATGAACATCTGGCCCCAAATTCATACTCTCCA         3475
FRI_Wa-1         GGTGGGTTAGGAAGAAGTGTATATGCATATGAACATCTGGCCCCAAATTCATACTCTCCA         3475
FRI_Spr-1-6      GGTGGGTTAGGAAGAAGTGTATATGCATATGAACATCTGGCCCCAAATTCATACTCTCCA         3475
FRI_Bil-7        GGTGGGTTAGGAAGAAGTGTATATGCATATGAACATCTGGCCCCAAATTCATACTCTCCA         3475
FRI_Alc-0        GGTGGGTTAGGAAGAAGTGTATATGCATATGAACATCTGGCCCCAAATTCATACTCTCCA         3475
FRI_Pu-2-23      GGTGGGTTAGGAAGAAGTGTATATGCATATGAACATCTGGCCCCAAATTCATACTCTCCA         3478
FRI_NFA-10       GGTGGGTTAGGAAGAAGTGTATATGCATATGAACATCTGGCCCCAAATTCATACTCTCCA         3478
FRI_Cvi-0        GGTGGGTTAGGAAGAAGTGTATATGCATATGAACATCTGGCCCCAAATTCATACTCTCCA         3478
FRI_Zdr-6        GGTGGGTTAGGAAGAAGTGTATATGCATATGAACATCTGGCCCCAAATTCATACTCTCCA         3478
                 ***************************************************** ******

FRI_An-1         GGTCACGGACATAGACTTCATCGACAGTACTCTCCGTCTTTGGTTCACGGACAGAGACAT         3497
FRI_Pro-0        GGTCACGGACATAGACTTCATCGACAGTACTCTCCGTCTTTGGTTCACGGACAGAGACAT         3499
FRI_Bål-2        GGTCACGGACATAGACTTCATCGACAGTACTCTCCGTCTTTGGTTCACGGACAGAGACAT         3535
FRI_St-0         GGTCACGGACATAGACTTCATCGACAGTACTCTCCGTCTTTGGTTCACGGACAGAGACAT         3464
FRI_Bg-2         GGTCACGGACATAGACTTCATCGACAGTACTCTCCGTCTTTGGTTCACGGACAGAGACAT         3532
FRI_C24          GGTCACGGACATAGACTTCATCGACAGTACTCTCCGTCTTTGGTTCACGGACAGAGACAT         3533
FRI_Van-0        GGTCACGGACATAGACTTCATCGACAGTACTCTCCGTCTTTGGTTCACGGACAGAGACAT         3533
FRI_Ull-2-3      GGTCACGGACATAGACTTCATCGACAGTACTCTCCGTCTTTGGTTCACGGACAGAGACAT         3532
FRI_Lip-0        GGTCACGGACATAGACTTCATCGACAGTACTCTCCGTCTTTGGTTCACGGACAGAGACAT         3536
FRI_NFA-8        GGTCACGGACATAGACTTCATCGACAGTACTCTCCGTCTTTGGTTCACGGACAGAGACAT         3536
FRI_Edi-0        GGTCACGGACATAGACTTCATCGACAGTACTCTCCGTCTTTGGTTCACGGACAGAGACAT         3535
FRI_Ren-1        GGTCACGGACATAGACTTCATCGACAGTACTCTCCGTCTTTGGTTCACGGACAGAGACAT         3164
FRI_Nok-3        GGTCACGGACATAGACTTCATCGACAGTACTCTCCGTCTTTGGTTCACGGACAGAGACAT         3535
FRI_Sha          GGTCACGGACATAGACTTCATCGACAGTACTCTCCGTCTTTGGTTCACGGACAGAGACAT         3535
FRI_Wa-1         GGTCACGGACATAGACTTCATCGACAGTACTCTCCGTCTTTGGTTCACGGACAGAGACAT         3535
FRI_Spr-1-6      GGTCACGGACATAGACTTCATCGACAGTACTCTCCGTCTTTGGTTCACGGACAGAGACAT         3535
FRI_Bil-7        GGTCACGGACATAGACTTCATCGACAGTACTCTCCGTCTTTGGTTCACGGACAGAGACAT         3535
FRI_Alc-0        GGTCACGGACATAGACTTCATCGACAGTACTCTCCGTCTTTGGTTCACGGACAGAGACAT         3535
FRI_Pu-2-23      GGTCACGGACATAGACTTCATCGACAGTACTCTCCGTCTTTGGTTCACGGACAGAGACAT         3538
FRI_NFA-10       GGTCACGGACATAGACTTCATCGACAGTACTCTCCGTCTTTGGTTCACGGACAGAGACAT         3538
FRI_Cvi-0        GGTCACGGACATAGACTTCATCGACAGTACTCTCCGTCTTTGGTTCACGGACAGAGACAT         3538
```

```
FRI_Zdr-6       GGTCACGGACATAGACTTCATCGACAGTACTCTCCGTCTTTGGTTCACGGACAGAGACAT       3538
                ************************************************************

FRI_An-1        CCACTACAGTACTCTCCTCCAATTCATGGACAACAACAGTTACCATATGGTATACAAAGG       3557
FRI_Pro-0       CCACTACAGTACTCTCCTCCAATTCATGGACAACAACAGTTACCATATGGTATACAAAGG       3559
FRI_Bål-2       CCACTACAGTACTCTCCTCCAATTCATGGACAACAACAGTTACCATATGGTATACAAAGG       3595
FRI_St-0        CCACTACAGTACTCTCCTCCAATTCATGGACAACAACAGTTACCATATGGTATACAAAGG       3524
FRI_Bg-2        CCACTACAGTACTCTCCTCCAATTCATGGACAACAACAGTTACCATATGGTATACAAAGG       3592
FRI_C24         CCACTACAGTACTCTCCTCCAATTCATGGACAACAACAGTTACCATATGGTATACAAAGG       3593
FRI_Van-0       CCACTACAGTACTCTCCTCCAATTCATGGACAACAACAGTTACCATATGGTATACAAAGG       3593
FRI_Ull-2-3     CCACTACAGTACTCTCCTCCAATTCATGGACAACAACAGTTACCATATGGTATACAAAGG       3592
FRI_Lip-0       CCACTACAGTACTCTCCTCCAATTCATGGACAACAACAGTTACCATATGGTATACAAAGG       3596
FRI_NFA-8       CCACTACAGTACTCTCCTCCAATTCATGGACAACAACAGTTACCATATGGTATACAAAGG       3596
FRI_Edi-0       CCACTACAGTACTCTCCTCCAATTCATGGACAACAACAGTTACCATATGGTATACAAAGG       3595
FRI_Ren-1       CCACTACAGTACTCTCCTCCAATTCATGGACAACAACAGTTACCATATGGTATACAAAGG       3224
FRI_Nok-3       CCACTACAGTACTCTCCTCCAATTCATGGACAACAACAGTTACCATATGGTATACAAAGG       3595
FRI_Sha         CCACTACAGTACTCTCCTCCAATTCATGGACAACAACAGTTACCATATGGTATACAAAGG       3595
FRI_Wa-1        CCACTACAGTACTCTCCTCCAATTCATGGACAACAACAGTTACCATATGGTATACAAAGG       3595
FRI_Spr-1-6     CCACTACAGTACTCTCCTCCAATTCATGGACAACAACAGTTACCATATGGTATACAAAGG       3595
FRI_Bil-7       CCACTACAGTACTCTCCTCCAATTCATGGACAACAACAGTTACCATATGGTATACAAAGG       3595
FRI_Alc-0       CCACTACAGTACTCTCCTCCAATTCATGGACAACAACAGTTACCATATGGTATACAAAGG       3595
FRI_Pu-2-23     CCACTACAGTACTCTCCTCCAATTCATGGACAACAACAGTTACCATATTGTATACAAAGG       3598
FRI_NFA-10      CCACTACAGTACTCTCCTCCAATTCATGGACAACAACAGTTACCATATGGTATACAAAGG       3598
FRI_Cvi-0       CCACTACAGTACTCTCCTCCAATTCATGGACAACAACAGTTACCATATGGTATACAAAGG       3598
FRI_Zdr-6       CCACTACAGTACTCTCCTCCAATTCATGGACAACAACAGTTACCATATGGTATACAAAGG       3598
                ************************************************ ***********

FRI_An-1        GTTTACAGACATTCACCATCTGAAGAAAGATATTTGGGTTTATCCAATCAAAGGTCTCCT       3617
FRI_Pro-0       GTTTACAGACATTCACCATCTGAAGAAAGATATTTGGGTTTATCCAATCAAAGGTCTCCT       3619
FRI_Bål-2       GTTTACAGACATTCACCATCTGAAGAAAGATATTTGGGTTTATCCAATCAAAGGTCTCCT       3655
FRI_St-0        GTTTACAGACATTCACCATCTGAAGAAAGATATTTGGGTTTATCCAATCAAAGGTCTCCT       3584
FRI_Bg-2        GTTTACAGACATTCACCATCTGAAGAAAGATATTTGGGTTTATCCAATCAAAGGTCTCCT       3652
FRI_C24         GTTTACAGACATTCACCATCTGAAGAAAGATATTTGGGTTTATCCAATCAAAGGTCTCCT       3653
FRI_Van-0       GTTTACAGACATTCACCATCTGAAGAAAGATATTTGGGTTTATCCAATCAAAGGTCTCCT       3653
FRI_Ull-2-3     GtTTACAGACATTCACCATCTGAAGAAAGATATTTGGGTTTATCCAATCAAAGGTCTCCT       3652
FRI_Lip-0       GTTTACAGACATTCACCATCTGAAGAAAGATATTTGGGTTTATCCAATCAAAGGTCTCCT       3656
FRI_NFA-8       GTTTACAGACATTCACCATCTGAAGAAAGATATTTGGGTTtATCCAATCAAAGGTCTCCT       3656
FRI_Edi-0       GTTTACAGACATTCACCATCTGAAGAAAGATATTTGGGTTtATCCAATCAAAGGTCTCCT       3655
FRI_Ren-1       GTTTACAGACATTCACCATCTGAAGAAAGATATTTGGGTTtATCCAATCAAAGGTCTCCT       3284
FRI_Nok-3       GTTTACAGACATTCACCATCTGAAGAAAGATATTTGGGTTtATCCAATCAAAGGTCTCCT       3655
FRI_Sha         GTTTACAGACATTCACCATCTGAAGAAAGATATTTGGGTTTATCCAATCAAAGGTCTCCT       3655
FRI_Wa-1        GTTTACAGACATTCACCATCTGAAGAAAGATATTTGGGTTTATCCAATCAAAGGTCTCCT       3655
FRI_Spr-1-6     GTTTACAGACATTCACCATCTGAAGAAAGATATTTGGGTTTATCCAATCAAAGGTCTCCT       3655
FRI_Bil-7       GTTTACAGACATTCACCATCTGAAGAAAGATATTTGGGTTTATCCAATCAAAGGTCTCCT       3655
FRI_Alc-0       GTTTACAGACATTCACCATCTGAAGAAAGATATTTGGGTTTATCCAATCAAAGGTCTCCT       3655
FRI_Pu-2-23     GTTTACAGACATTCACCATCTGAAGAAAGATATTTGGGTTTATCCAATCAAAGGTCTCCT       3658
FRI_NFA-10      GTTTACAGACATTCACCATCTGAAGAAAGATATTTGGGTTtATCCAATCAAAGGTCTCCT       3658
FRI_Cvi-0       GtTTACAGACATTCACCATcTGAAGAAAGATATTTGGGTTTATCCAATCAAAGGTCTCCT       3658
FRI_Zdr-6       GTTTACAGACATTCACCATCTGAAGAAAGATATTTGGGTTTATCCAATCAAAGGTCTCCT       3658
                ************************************************************

FRI_An-1        CGCAGTAACTCATCATTAGACCCCAAATAGGAGGAATGTAAATTTGTAACAAAGCTTTTT       3677
FRI_Pro-0       CGCAGTAACTCATCATTAGACCCCAAATAGGAGGaATGTAAATTTGTAACAAAGCTTTTT       3679
FRI_Bål-2       CGCAGTAACTCATCATTAGACCCCAAATAGGAGGAATGTAAATTTGTAACAAAGCTTTTT       3715
FRI_St-0        CGCAGTAACTCATCATTAGACCCCAAATAGGAGGaATGTAAATTTGTAACAAAGCTTTTT       3644
FRI_Bg-2        CGCAGTAACTCATCATTAGACCCCAAATAGGAGGAATGTAAATTTGTAACAAAGCTTTTT       3712
FRI_C24         CGCAGTAACTCATCATTAGACCCCAAATAGGAGGAATGTAAATTTGTAACAAAGCTTTTT       3713
FRI_Van-0       CGCAGTAACTCATCATTAGACCCCAAATAGGAGGAATGTAAATTTGTAACAAAGCTTTTT       3713
FRI_Ull-2-3     CGCAGTAACTCATCATTAGACCCCAAATAGGAGGAATGTAAATTTGTAACAAAGCTTTTT       3712
FRI_Lip-0       CGCAGTAACTCATCATTAGACCCCAAATAGGAGGAATGTAAATTTGTAACAAAGCTTTTT       3716
FRI_NFA-8       CGCAGTAACTCATCATTAGACCCCAAATAGGAGGAATGTAAATTTGTAACAAAGCTTTTT       3716
FRI_Edi-0       CGCAGTAACTCATCATTAGACCCCAAATAGGAGGaATGTAAATTTGTAACAAAGCTTTTT       3715
FRI_Ren-1       CGCAGTAACTCATCATTAGACCCCAAATAGGAGGaATGTAAATTTGTAACAAAGCTTTTT       3344
FRI_Nok-3       CGCAGTAACTCATCATTAGACCCCAAATAGGAGGaATGTAAATTTGTAACAAAGCTTTTT       3715
FRI_Sha         CGCAGTAACTCATCATTAGACCCCAAATAGGAGGAATGTAAATTTGTAACAAAGCTTTTT       3715
FRI_Wa-1        CGCAGTAACTCATCATTAGACCCCAAATAGGAGGAATGTAAATTTGTAACAAAGCTTTTT       3715
FRI_Spr-1-6     CGCAGTAACTCATCATTAGACCCCAAATAGGAGGAATGTAAATTTGTAACAAAGCTTTTT       3715
FRI_Bil-7       CGCAGTAACTCATCATTAGACCCCAAATAGGAGGAATGTAAATTTGTAACAAAGCTTTTT       3715
FRI_Alc-0       CGCAGTAACTCATCATTAGACCCCAAATAGGAGGAATGTAAATTTGTAACAAAGCTTTTT       3715
FRI_Pu-2-23     CGCAGTAACTCATCATTAGACCCCAAATAGGAGGAATGTAAATTTGTAACAAAGCTTTTT       3718
FRI_NFA-10      CGCAGTAACTCATCATTAGACCCCAAATAGGAGGAATGTAAATTTGTAACAAAGCTTTTT       3718
```

```
FRI_Cvi-0       CGCAGTAACTCATCATTAGACCCCAAATAGGAGGAATGTAAATTTGTAACAAAGCTTTTT       3718
FRI_Zdr-6       CGCAGTAACTCATCATTAGACCCCAAATAGGAGGAATGTAAATTTGTAACAAAGCTTTTT       3718
                ************************************************************

FRI_An-1        GTTTTTGCTTAAGTTAGTCATTTATTTAACTCCCAACAGTCTCAAAATTTAATTTAATGT       3737
FRI_Pro-0       GTTTTTGCTTAAGTTAGTCATTTATTTAACTCCCAACAGTCTCAAAATTTAATTTAATGT       3739
FRI_Bå1-2       GTTTTTGCTTAAGTTAGTCATTTATTTAACTCCCAACAGTCTCAAAATTTAATTTAATGT       3775
FRI_St-0        GTTTTTGCTTAAGTTAGTCATTTATTTAACTCCCAACAGTCTCAAAATTTAATTTAATGT       3704
FRI_Bg-2        GTTTTTGCTTAAGTTAGTCATTTATTTAACTCCCAACAGTCTCAAAATTTAATTTAATGT       3772
FRI_C24         GTTTTTGCTTAAGTTAGTCATTTATTTAACTCCCAACAGTCTCAAAATTTAATTTAATGT       3773
FRI_Van-0       GTTTTTGCTTAAGTTAGTCATTTATTTAACTCCCAACAGTCTCAAAATTTAATTTAATGT       3773
FRI_Ull-2-3     GTTTTTGCTTAAGTTAGTCATTTATTTAACTCCCAACAGTCTCAAAATTTAATTTAATGT       3772
FRI_Lip-0       GTTTTTGCTTAAGTTAGTCATTTATTTAACTCCCAACAGTCTCAAAATTTAATTTAATGT       3776
FRI_NFA-8       GTTTTTGCTTAAGTTAGTCATTTATTTAACTCCCAACAGTCTCAAAATTTAATTTAATGT       3776
FRI_Edi-0       GTTTTTGCTTAAGTTAGTCATTTATTTAACTCCCAACAGTCTCAAAATTTAATTTAATGT       3775
FRI_Ren-1       GTTTTTGCTTAAGTTAGTCATTTATTTAACTCCCAACAGTCTCAAAATTTAATTTAATGT       3404
FRI_Nok-3       GTTTTTGCTTAAGTTAGTCATTTATTTAACTCCCAACAGTCTCAAAATTTAATTTAATGT       3775
FRI_Sha         GTTTTTGCTTAAGTTAGTCATTTATTTAACTCCCAACAGTCTCAAAATTTAATTTAATGT       3775
FRI_Wa-1        GTTTTTGCTTAAGTTAGTCATTTATTTAACTCCCAACAGTCTCAAAATTTAATTTAATGT       3775
FRI_Spr-1-6     GTTTTTGCTTAAGTTAGTCATTTATTTAACTCCCAACAGTCTCAAAATTTAATTTAATGT       3775
FRI_Bil-7       GTTTTTGCTTAAGTTAGTCATTTATTTAACTCCCAACAGTCTCAAAATTTAATTTAATGT       3775
FRI_Alc-0       GTTTTTGCTTAAGTTAGTCATTTATTTAACTCCCAACAGTCTCAAAATTTAATTTAATGT       3775
FRI_Pu-2-23     GTTTTTGCTTAAGTTAGTCATTTATTTAACTCCCAACAGTCTCAAAATTTAATTTAATGT       3778
FRI_NFA-10      GTTTTTGCTTAAGTTAGTCATTTATTTAACTCCCAACAGTCTCAAAATTTAATTTAATGT       3778
FRI_Cvi-0       GTTTTTGCTTAAGTTAGTCATTTATTTAACTCCCAACAGTCTCAAAATTTAATTTAATGT       3778
FRI_Zdr-6       GTTTTTGCTTAAGTTAGTCATTTATTTAACTCCCAACAGTCTCAAAATTTAATTTAATGT       3778
                ************************************************************

FRI_An-1        TTGGGGCTTAAGAATGCAAATTTTTTTGCTCCTGTAATTGACATTTAAGATGCTAATGTT       3797
FRI_Pro-0       TTGGGGCTTAAGAATGCAAATTTTTTTGCTCCTGTAATTGACATTTAAGATGCTAATGTT       3799
FRI_Bå1-2       TTGGGGCTTAAGAATGCAAATTTTTTTGCTCCTGTAATTGACATTTAAGATGCTAATGTT       3835
FRI_St-0        TTGGGGCTTAAGAATGCAAATTTTTTTGCTCCTGTAATTGACATTTAAGATGCTAATGTT       3764
FRI_Bg-2        TTGGGGCTTAAGAATGCAAATTTTTTTGCTCCTGTAATTGACATTTAAGATGCTAATGTT       3832
FRI_C24         TTGGGGCTTAAGAATGCAAATTTTTTTGCTCCTGTAATTGACATTTAAGATGCTAATGTT       3833
FRI_Van-0       TTGGGGCTTAAGAATGCAAATTTTTTTGCTCCTGTAATTGACATTTAAGATGCTAATGTT       3833
FRI_Ull-2-3     TTGGGGCTTAAGAATGCAAATTTTTTTGCTCCTGTAATTGACATTTAAGATGCTAATGTT       3832
FRI_Lip-0       TTGGGGCTTAAGAATGCAAATTTTTTTGCTCCTGTAATTGACATTTAAGATGCTAATGTT       3836
FRI_NFA-8       TTGGGGCTTAAGAATGCAAATTTTTTTGCTCCTGTAATTGACATTTAAGATGCTAATGTT       3836
FRI_Edi-0       TTGGGGCTTAAGAATGCAAATTTTTTTGCTCCTGTAATTGACATTTAAGATGCTAATGTT       3835
FRI_Ren-1       TTGGGGCTTAAGAATGCAAATTTTTTTGCTCCTGTAATTGACATTTAAGATGCTAATGTT       3464
FRI_Nok-3       TTGGGGCTTAAGAATGCAAATTTTTTTGCTCCTGTAATTGACATTTAAGATGCTAATGTT       3835
FRI_Sha         TTGGGGCTTAAGAATGCAAATTTTTTTGCTCCTGTAATTGACATTTAAGATGCTAATGTT       3835
FRI_Wa-1        TTGGGGCTTAAGAATGCAAATTTTTTTGCTCCTGTAATTGACATTTAAGATGCTAATGTT       3835
FRI_Spr-1-6     TTGGGGCTTAAGAATGCAAATTTTTTTGCTCCTGTAATTGACATTTAAGATGCTAATGTT       3835
FRI_Bil-7       TTGGGGCTTAAGAATGCAAATTTTTTTGCTCCTGTAATTGACATTTAAGATGCTAATGTT       3835
FRI_Alc-0       TTGGGGCTTAAGAATGCAAATTTTTTTGCTCCTGTAATTGACATTTAAGATGCTAATGTT       3835
FRI_Pu-2-23     TTGGGGCTTAAGAATGCAAATTTTTTTGCTCCTGTAATTGACATTTAAGATGCTAATGTT       3838
FRI_NFA-10      TTGGGGCTTAAGAATGCAAATTTTTTTGCTCCTGTAATTGACATTTAAGATGCTAATGTT       3838
FRI_Cvi-0       TTGGGGCTTAAGAATGCAAATTTTTTTGCTCCTGTAATTGACATTTAAGATGCTAATGTT       3838
FRI_Zdr-6       TTGGGGCTTAAGAATGCAAATTTTTTTGCTCCTGTAATTGACATTTAAGATGCTAATGTT       3838
                ************************************************************

FRI_An-1        ATTGCTTCAGAGGTTTTAGTCAACCTCAGATACATCGATATCACTATCTAAATAGACCTC       3857
FRI_Pro-0       ATTGCTTCAGAGGTTTTAGTCAACCTCAGATACATCGATATCACTATCTAAATAGACCTC       3859
FRI_Bå1-2       ATTGCTTCAGAGGTTTTAGTCAACCTCAGATACATCGATATCACTATCTAAAATAGACCT       3895
FRI_St-0        ATTGCTTCAGAGGTTTTAGTCAACCTCAGATACATCGATATCACTATCTAAATAGACCTC       3824
FRI_Bg-2        ATTGCTTCAGAGGTTTTAGTCAACCTCAGATACATCGATATCACTATCTAAATAGACCTC       3892
FRI_C24         ATTGCTTCAGAGGTTTTAGTCAACCTCAGATACATCGATATCACTATCTAAATAGACCTC       3893
FRI_Van-0       ATTGCTTCAGAGGTTTTAGTCAACCTCAGATACATCGATATCACTATCTAAATAGACCTC       3893
FRI_Ull-2-3     ATTGCTTCAGAGGTTTTAGTCAACCTCAGATACATCGATATCACTATCTAAATAGACCTC       3892
FRI_Lip-0       ATTGCTTCAGAGGTTTTAGTCAACCTCAGATACATCGATATCACTATCTAAATAGACCTC       3896
FRI_NFA-8       ATTGCTTCAGAGGTTTTAGTCAACCTCAGATACATCGATATCACTATCTAAATAGACCTC       3896
FRI_Edi-0       ATTGCTTCAGAGGTTTTAGTCAACCTCAGATACATCGATATCACTATCTAAATAGACCTC       3895
FRI_Ren-1       ATTGCTTCAGAGGTTTTAGTCAACCTCAGATACATCGATATCACTATCTAAATAGACCTC       3524
FRI_Nok-3       ATTGCTTCAGAGGTTTTAGTCAACCTCAGATACATCGATATCACTATCTAAATAGACCTC       3895
FRI_Sha         ATTGCTTCAGAGGTTTTAGTCAACCTCAGATACATCGATATCACTATCTAAATAGACCTC       3895
FRI_Wa-1        ATTGCTTCAGAGGTTTTAGTCAACCTCAGATACATCGATATCACTATCTAAATAGACCTC       3895
FRI_Spr-1-6     ATTGCTTCAGAGGTTTTAGTCAACCTCAGATACATCGATATCACTATCTAAATAGACCTC       3895
FRI_Bil-7       ATTGCTTCAGAGGTTTTAGTCAACCTCAGATACATCGATATCACTATCTAAATAGACCTC       3895
FRI_Alc-0       ATTGCTTCAGAGGTTTTAGTCAACCTCAGATACATCGATATCACTATCTAAATAGACCTC       3895
FRI_Pu-2-23     ATTGCTTCAGAGGTTTTAGTCAACCTCAGATACATCGATATCACTATCTAAATAGACCTC       3898
```

```
FRI_NFA-10       ATTGCTTCAGAGGTTTTAGTCAACCTCAGATACATCGATATCACTATCTAAATAGACCTC         3898
FRI_Cvi-0        ATTGCTTCAGAGGTTTTAGTCAACCTCAGATACATCGATATCACTATCTAAATAGACCTC         3898
FRI_Zdr-6        ATTGCTTCAGAGGTTTTAGTCAACCTCAGATACATCGATATCACTATCTAAATAGACCTC         3898
                 ****************************************************     *

FRI_An-1         TGGCTCTTGGTCATC-T--GGATTCTCTTCATCTTCTGT-CTCTGTTC----CTTCTTGT         3909
FRI_Pro-0        TGGCTCTTGGTCATC-T--GGATTCTCTTCATCTTCTGT-CTCTGTTC----CTTCTTGT         3911
FRI_Bå1-2        CTGGCTCTTGGGTCATCTGGGATTCTCTTCATCCTCTGGTCTCGGTTCCTTCTTGGTTCT         3955
FRI_St-0         TG-GCTCTTGGTCATCT--GGATTCTCTTCATCTTCTGT-CTCTGTTCCTTCTTGTTCT-         3879
FRI_Bg-2         TG-GCTCTTGGTCATCT--GGATTCTCTTCATCTTCTGT-CTCTGTTCCTTCTTGTTCT-         3947
FRI_C24          TG-GCTCTTGGTCATCT--GGATTCTCTTCATCTTCTGT-CTCTGTTCCTTCTTGTTCT-         3948
FRI_Van-0        TG-GCTCTTGGTCATCT--GGATTCTCTTCATCTTCTGT-CTCTGTTCCTTCTTGTTCT-         3948
FRI_Ull-2-3      TG-GCTCTTGGTCATCT--GGATTCTCTTCATCTTCTGT-CTCTGTTCCTTCTTGTTCT-         3947
FRI_Lip-0        TG-GCTCTTGGTCATCT--GGATTCTCTTCATCTTCTGT-CTCTGTTCCTTCTTGTTCT-         3951
FRI_NFA-8        TG-GCTCTTGGTCATCT--GGATTCTCTTCATCTTCTGT-CTCTGTTCCTTCTTGTTCT-         3951
FRI_Edi-0        TG-GCTCTTGGTCATCT--GGATTCTCTTCATCTTCTGT-CTCTGTTCCTTCTTGTTCT-         3950
FRI_Ren-1        TG-GCTCTTGGTCATCT--GGATTCTCTTCATCTTCTGT-CTCTGTTCCTTCTTGTTCT-         3579
FRI_Nok-3        TG-GCTCTTGGTCATCT--GGATTCTCTTCATCTTCTGT-CTCTGTTCCTTCTTGTTCT-         3950
FRI_Sha          TG-GCTCTTGGTCATCT--GGATTCTCTTCATCTTCTGT-CTCTGTTCCTTCTTGTTCT-         3950
FRI_Wa-1         TG-GCTCTTGGTCATCT--GGATTCTCTTCATCTTCTGT-CTCTGTTCCTTCTTGTTCT-         3950
FRI_Spr-1-6      TG-GCTCTTGGTCATCT--GGATTCTCTTCATCTTCTGT-CTCTGTTCCTTCTTGTTCT-         3950
FRI_Bil-7        TG-GCTCTTGGTCATCT--GGATTCTCTTCATCTTCTGT-CTCTGTTCCTTCTTGTTCT-         3950
FRI_Alc-0        TG-GCTCTTGGTCATCT--GGATTCTCTTCATCTTCTGT-CTCTGTTCCTTCTTGTTCT-         3950
FRI_Pu-2-23      TG-GCTCTTGGTCATCT--GGATTCTCTTCATCTTCTGT-CTCTGTTCCTTCTTGTTCT-         3953
FRI_NFA-10       TG-GCTCTTGGTCATCT--GGATTCTCTTCATCTTCTGT-CTCTGTTCCTTCTTGTTCT-         3953
FRI_Cvi-0        TG-GCTCTTGGTCATCT--GGATTCTCTTCATCTTCTGT-CTCTGTTCCTTCTTGTTCT-         3953
FRI_Zdr-6        TG-GCTCTTGGTCATCT--GGATTCTCTTCATCTTCTGT-CTCTGTTCCTTCTTGTTCT-         3953
                        * *         ************** ****  *** ****     *  *

FRI_An-1         TCTCGTTGCACTGCTCGAGCAATTGCGGATTCCAACCTTGTGCTTACAGTTTCCCATGAC         3969
FRI_Pro-0        TCTCGTTGCACTGCTCGAGCAATTGCGGATTCCAACCTTGTGCTTACAGTTTCCCATGAC         3971
FRI_Bå1-2        CGTTGGCACTGCCTCGAAGCAATTGGCGGGA-----------------------------         3986
FRI_St-0         CGTTGCACTG---CTCGAGCAATTGCGGATTCCAACCTTGTGCTTACAGTTTCCCATGAC         3936
FRI_Bg-2         CGTTGCACTG---CTCGAGCAATTGCGGATTCCAACCTTGTGCTTACAGTTTCCCATGAC         4004
FRI_C24          CGTTGCACTG---CTCGAGCAATTGCGGATTCCAACCTTGTGCTTACAGTTTCCCATGAC         4005
FRI_Van-0        CGTTGCACTG---CTCGAGCAATTGCGGATTCCAACCTTGTGCTTACAGTTTCCCATGAC         4005
FRI_Ull-2-3      CGTTGCACTG---CTCGAGCAATTGCGGATTCCAACCTTGTGCTTACAGTTTCCCATGAC         4004
FRI_Lip-0        CGTTGCACTG---CTCGAGCAATTGCGGATTCCAACCTTGTGCTTACAGTTTCCCATGAC         4008
FRI_NFA-8        CGTTGCACTG---CTCGAGCAATTGCGGATTCCAACCTTGTGCTTACAGTTTCCCATGAC         4008
FRI_Edi-0        CGTTGCACTG---CTCGAGCAATTGCGGATTCCAACCTTGTGCTTACAGTTTCCCATGAC         4007
FRI_Ren-1        CGTTGCACTG---CTCGAGCAATTGCGGATTCCAACCTTGTGCTTACAGTTTCCCATGAC         3636
FRI_Nok-3        CGTTGCACTG---CTCGAGCAATTGCGGATTCCAACCTTGTGCTTACAGTTTCCCATGAC         4007
FRI_Sha          CGTTGCACTG---CTCGAGCAATTGCGGATTCCAACCTTGTGCTTACAGTTTCCCATGAC         4007
FRI_Wa-1         CGTTGCACTG---CTCGAGCAATTGCGGATTCCAACCTTGTGCTTACAGTTTCCCATGAC         4007
FRI_Spr-1-6      CGTTGCACTG---CTCGAGCAATTGCGGATTCCAACCTTGTGCTTACAGTTTCCCATGAC         4007
FRI_Bil-7        CGTTGCACTG---CTCGAGCAATTGCGGATTCCAACCTTGTGCTTACAGTTTCCCATGAC         4007
FRI_Alc-0        CGTTGCACTG---CTCGAGCAATTGCGGATTCCAACCTTGTGCTTACAGTTTCCCATGAC         4007
FRI_Pu-2-23      CGTTGCACTG---CTCGAGCAATTGCGGATTCCAACCTTGTGCTTACAGTTTCCCATGAC         4010
FRI_NFA-10       CGTTGCACTG---CTCGAGCAATTGCGGATTCCAACCTTGTGCTTACAGTTTCCCATGAC         4010
FRI_Cvi-0        CGTTGCACTG---CTCGAGCAATTGCGGATTCCAACCTTGTGCTTACAGTTTCCCATGAC         4010
FRI_Zdr-6        CGTTGCACTG---CTCGAGCAATTGCGGATTCCAACCTTGTGCTTACAGTTTCCCATGAC         4010
                   * *            ********  *

FRI_An-1         ACAAGCTTTTCCATGAATGTATTTATGTCCGCCTTCTTATCTTTCTTGAGGAAGATGAAT         4029
FRI_Pro-0        ACAAGCTTTTCCATGAATGTATTTATGTCCGCCTTCTTATCTTTCTTGAGGAAGATGAAT         4031
FRI_Bå1-2        ------------------------------------------------------------         3986
FRI_St-0         ACAAGCTTTTCCATGAATGTATTTATGTCCGCCTTCTTATCTTTCTTGAGGAAGATGAAT         3996
FRI_Bg-2         ACAAGCTTTTCCATGAATGTATTTATGTCCGCCTTCTTATCTTTCTTGAGGAAGATGAAT         4064
FRI_C24          ACAAGCTTTTCCATGAATGTATTTATGTCCGCCTTCTTATCTTTCTTGAGGAAGA-----         4060
FRI_Van-0        ACAAGCTTTTCCATGAATGTATTTATGTCCGCCTTCTTATCTTTCTTGAGGAAGATGAAT         4065
FRI_Ull-2-3      ACAAGCTTTTCCATGAATGTATTTATGTCCGCCTTCTTATCTTTCTTGAGGAAGATGAAT         4064
FRI_Lip-0        ACAAGCTTTTCCATGAATGTATTTATGTCCGCCTTCTTATCTTTCTTGAGGAAGATGAAT         4068
FRI_NFA-8        ACAAGCTTTTCCATGAATGTATTTATGTCCGCCTTCTTATCTTTCTTGAGGAAGATGAAT         4068
FRI_Edi-0        ACAAGCTTTTCCATGAATGTATTTATGTCCGCCTTCTTATCTTTCTTGAGGAAGATGAAT         4067
FRI_Ren-1        ACAaGCTTTTCCATGAATGTATTTATGTCCGCCTTCTTATCTTTCTTGAGGAAGATGAAT         3696
FRI_Nok-3        ACAAGCTTTTCCATGAATGTATTTATGTCCGCCTTCTTATCTTTCTTGAGGAAGATGAAT         4067
FRI_Sha          ACAAGCTTTTCCATGAATGTATTTATGTCCGCCTTCTTATCTTTCTTGAGGAAGATGAAT         4067
FRI_Wa-1         ACAAGCTTTTCCATGAATGTATTTATGTCCGCCTTCTTATCTTTCTTGAGGAAGATGAAT         4067
FRI_Spr-1-6      ACAAGCTTTTCCATGAATGTATTTATGTCCGCCTTCTTATCTTTCTTGAGGAAGATGAAT         4067
FRI_Bil-7        ACAAGCTTTTCCATGAATGTATTTATGTCCGCCTTCTTATCTTTCTTGAGGAaGATGAAT         4067
FRI_Alc-0        ACAAGCTTTTCCATGAATGTATTTATGTCCGCCTTCTTATCTTTCTTGAGGAAGATGAAT         4067
```

```
FRI_Pu-2-23      ACAAGCTTTTCCATGAATGTATTTATGTCCGCCTTCTTATCTTTCTTGAGGAAGATGAAT         4070
FRI_NFA-10       ACAAGCTTTTCCATGAATGTATTTATGTCCGCCTTCTTATCTTTCTTGAGGAAGATGAAT         4070
FRI_Cvi-0        ACAAGCTTTTCCATGAATGTATTTATGTCCGCCTTCTTATCTTTCTTGAGGAAGATGAAT         4070
FRI_Zdr-6        ACAAGCTTTTCCATGAATGTATTTATGTCCGCCTTCTTATCTTTCTTGAGGAAGATGAAT         4070

FRI_An-1         TC          4031
FRI_Pro-0        TC          4033
FRI_Bål-2        --          3986
FRI_St-0         TC          3998
FRI_Bg-2         TC          4066
FRI_C24          --          4060
FRI_Van-0        TC          4067
FRI_Ull-2-3      TC          4066
FRI_Lip-0        T-          4069
FRI_NFA-8        TC          4070
FRI_Edi-0        TC          4069
FRI_Ren-1        TC          3698
FRI_Nok-3        TC          4069
FRI_Sha          TC          4069
FRI_Wa-1         TC          4069
FRI_Spr-1-6      TC          4069
FRI_Bil-7        TC          4069
FRI_Alc-0        TC          4069
FRI_Pu-2-23      TC          4072
FRI_NFA-10       TC          4072
FRI_Cvi-0        TC          4072
FRI_Zdr-6        TC          4072
```
